# metan: an R package for multi-environment trial analysis

**DOI:** 10.1101/2020.01.14.906750

**Authors:** Tiago Olivoto, Alessandro Dal’Col Lúcio

## Abstract

1. Multi-environment trials (MET) are crucial steps in plant breeding programs that aim increasing crop productivity to ensure global food security. The analysis of MET data requires the combination of several approaches including data manipulation, visualization, and modeling. As new methods are proposed, analyzing MET data correctly and completely remains a challenge, often intractable with existing tools.
2. Here we describe the `metan R` package, a collection of functions that implement a workflow-based approach to (a) check, manipulate and summarise typical MET data; (b) analyze individual environments using both fixed and mixed-effect models; (c) compute parametric and non-parametric stability statistics; (c) implement biometrical models widely used in MET analysis; and (d) plot typical MET data quickly.
3. In this paper, we present a summary of the functions implemented in `metan` and how they integrate into a workflow to explore and analyze MET data. We guide the user along a gentle learning curve and show how adding only a few commands or options at a time, powerfull analyzes can be implemented.
4. `metan` offers a flexible, intuitive, and richly documented working environment with tools that will facilitate the implementation of a complete analysis of MET data sets.

## 1 Introduction

In 50 years (1967-2017) the world average of cereal yields has increased by 64%, from 1.68 to 2.76 t ha^−1^. In the same period, the total production of cereals has raised from 1.305 × 10^9^ to 3.6 × 10^9^ t, an increase of 175%, while the cultivated area increased by only 7.9% in the same period (FAOSTAT, 2019). These unparallel increases have been possible due to the improved cultivation techniques in combination with superior cultivars. For maize, for example, 50% of the increase in yield was due to breeding (Duvick, 2005). Plant breeding programs have been developing new cultivars for adaptation to new locations, management practices, or growing conditions, in a clear and crucial example of exploitation of genotype-vs-environment interaction (GEI).

The breeders’ desire to modeling the GEI appropriately has led to the development of the so-called stability analyses, which includes ANOVA-based methods (Yates & Cochran, 1938; Wricke, 1965; Shukla, 1972; Annicchiarico, 1992); regression-based methods (Eberhart & Russell, 1966); non-parametric methods (Huehn, 1979; Lin & Binns, 1988; Fox, Skovmand, Thompson, Braun, & Cormier, 1990; Thennarasu, 1995) and some methods that combines different statistical techniques, such as the Additive Main Effect and Multiplicative Interaction, AMMI, (Gauch, 2013), and Genotype plus Genotype-vs-Environment interaction, GGE, (Yan & Kang, 2003). Then, it is no surprise that scientific production related to multi-environment trial analysis has been growing fast in the last decades. A bibliometric survey in the SCOPUS database revealed that in the last half-century (1969–2019) 6590 documents were published in 902 sources (Journals, books, etc.) by 19.351 authors. In this period, the number of publications has been increased on average by 11.22% year^−1^ but were in the last ten years that the largest amount (∼64%) of the documents were published (See Appendix S1, item 1 for more details).

Linear Mixed-effect Models (LMM) has been more frequently used to analyze MET data. For example, between 2013 and 2015, the larger number of papers proposing methods to deal with GEI were related to the Best Linear Unbiased Prediction (BLUP) in LMMs (Eeuwijk, Bustos-Korts, & Malosetti, 2016). Recent advances in this field showed that BLUP is more predictively accurate than AMMI and that the main advantages of these methods can be combined to help researchers to select or recommend stable and high productive genotypes (T. Olivoto, Lúcio, et al., 2019). Thus, the rapid spread of these methods to users around the world can be facilitated if these procedures are implemented in specific software.

In most cases, analyzing MET data involves manual checking of the data subset(s) to identify possible outliers, using some biometrical model to explore the relationships between traits(or groups of traits), computing a within-environment ANOVA, computing a joint-ANOVA, and, in case of a significant GEI, applying some stability method to explore it. While a spreadsheet program (e.g. Microsoft Excel) may be used to perform a visual check for outliers, an integrated development environment (IDE, e.g. R, SAS, or Matlab) is often required to process the complex matrix operations required in some stability methods. IDEs, however, require a certain degree of expertise to use and have steep learning curves, which sometimes prevents that a coding layman implements certain methods. In this sense, R (Team, 2019) packages have been making easier the life of hundreds of thousands of researchers by providing freely collections of functions developed by the community.

Some open-source R software packages that are designed –or are suitable– for analyzing MET data are available. The `stability` package (https://CRAN.R-project.org/package=stability) contains a collection of functions to perform stability analysis. The `ammistability` package (https://CRAN.R-project.org/package=ammistability) computes multiple AMMI-based stability parameters. The `gge` (https://CRAN.R-project.org/package=gge) and `GGEBiplots` (https://CRAN.R-project.org/package=GGEBiplots) packages may be used to perform a GGE model. The R packages `agricolae` (https://CRAN.R-project.org/package=agricolae) and `plantbreeding` (http://plantbreeding.r-forge.r-project.org/), while not specifically coded for MET analysis provides useful functions for computing parametric and nonparametric stability statistics. Although useful, these packages do not offer options to perform a complete analysis of MET data, i.e., to provide tools for all steps of the analysis (check, manipulation, analysis, and visualization of data). For example, `GGEBiplots` requires as input data a two-way table containing genotype by environment means with genotypes in rows and environments in columns, but doesn’t provide any function to create quickly such table from data that often is in a “*long*” format in R. In addition, several studies often compare different stability methods (e.g., Woyann et al., 2018; Scapim et al., 2010; Bornhofen et al., 2017; Freiria et al., 2018; Shahbazi, 2019; Teodoro et al., 2019). This requires a range of different packages to be used, making it the coding tedious and difficult to follow. Thus, it seems to be value the creation of an R package that presents an easy workflow, and incorporates the most used stability statistics, as well as recent introduced stability methods (T. Olivoto, Lúcio, et al., 2019; T. Olivoto et al., 2019) in addition to options for computing cross-validation (Piepho, 1994) and BLUP-based stability statistics (Colombari Filho et al., 2013), features frequently used but not yet implemented in any other R package for MET analysis.

Here, we describe the `metan` (**m**ulti-**e**nvironment **t**rial **an**alysis) package, an open-source R package designed to provide an efficient and reproducible workflow for the analysis of MET data. Our main aim in this paper is to describe the features of `metan` and how this collection of functions can be useful for an intuitive and complete analysis of MET data.

## 2 The metan package

The conceptual focus of `metan` is centered on five components (Fig. 1): (a) check, manipulate and summarise typical MET data; (b) performs within-environment analysis of variance; (c) compute parametric and non-parametric stability analysis; (d) compute biometrical models widely used in plant MET analysis of plant breeding trials; and (e) quickly create typical plots for two-way data considering any combination of qualitative and quantitative factors.

**Fig. 1.**
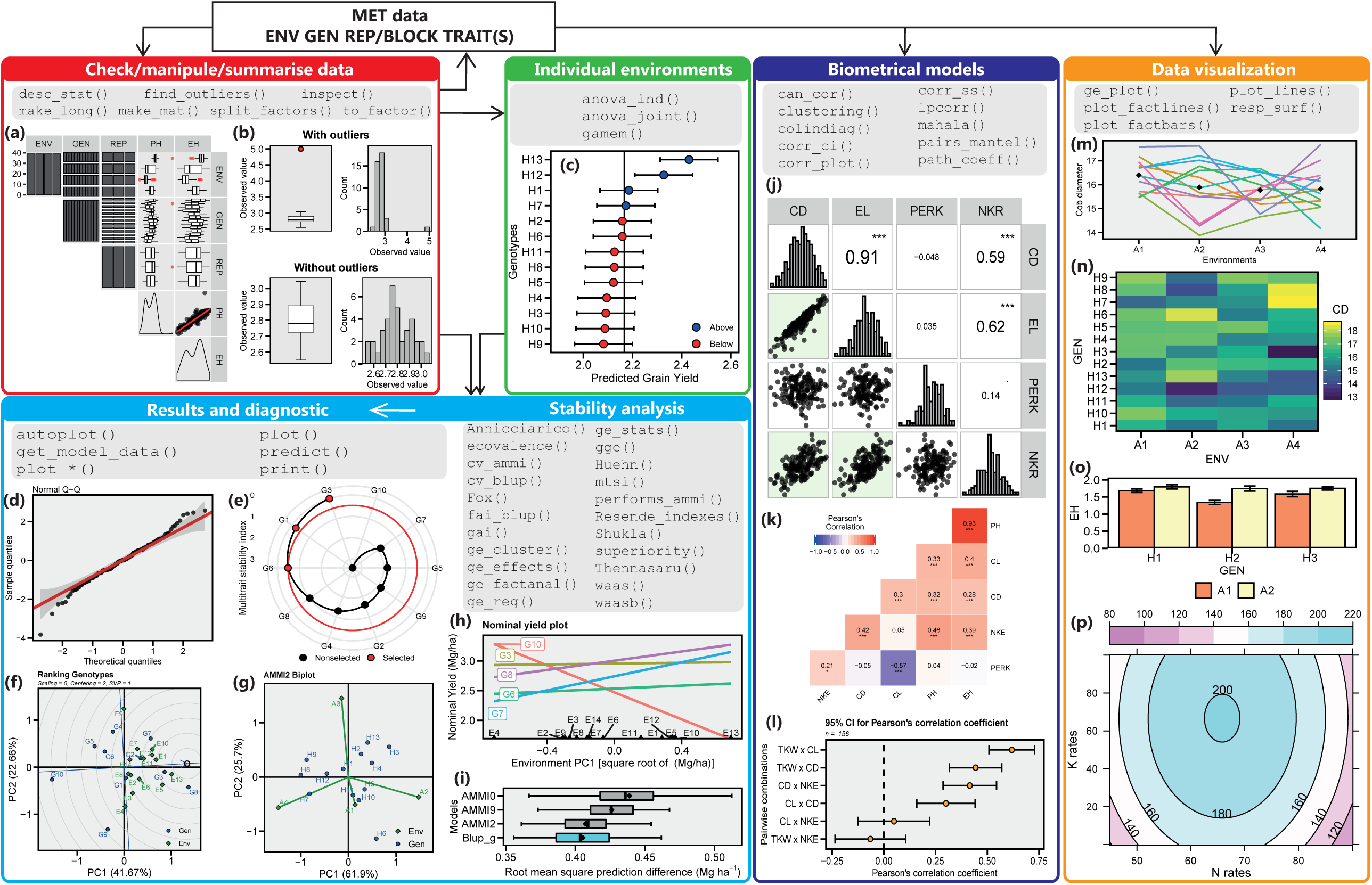
Diagram showing steps in a typical workflow in the analysis of multi-environment trial data using `metan`. (a) inspect plot made with `inspect();` (b) outlier check plot made with `find_outliers()`; (c) blups for genotypes made with `plot_blup()`; (d) model diagnostic made with `plot.*()`; (e) radar plot showing the multi-trait stability index made with `plot.mtsi()`;(f) a gge biplot made with `plot.gge()`; (g-h) an AMMI2 biplot and a nominal yield plot, respectively, made with `plot_scores()`; (i) results for a cross validation procedure made with `plot.cv_ammif()`; (j-k) visualization of correlation matrices with `corr_plot()` and `plot.corr_coef()`, respectively; (l) nonparametric confidence intervals for correlation made with `plot.corr_ci()`; (m-n) genotype-vs-environment plot made with `ge_plot()`; (o) a barplot created with `plot_factbars()`; (p) a contour plot created with `plot.resp_surf()`.

The development version of `metan` is available on Github (https://github.com/TiagoOlivoto/metan) and can be installed directly via the R console using `devtools`:

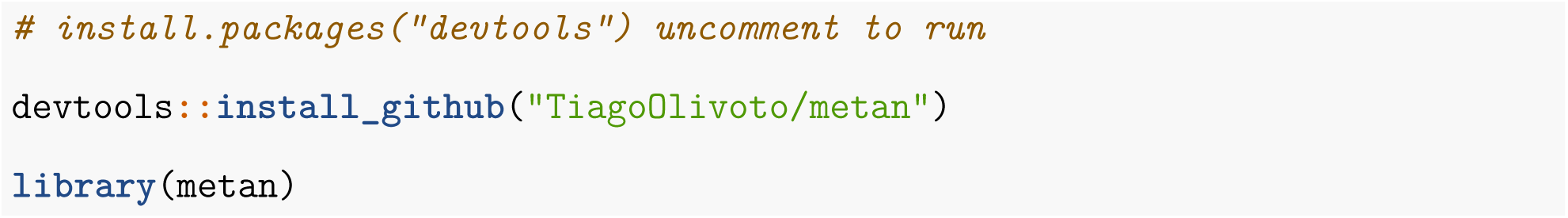

To illustrate the main features of the package, six example datasets (`data_alpha, data_g, data_ge, data_ge2, int.effects,` and `meansGxE`) are distributed with `metan`. Comprehensive details and examples of the functionality of `metan` are available in our online documentation (https://tiagoolivoto.github.io/metan/). Indeed, we strongly encourage readers to refer to the vignettes as the primary source for information on `metan`’s functionality since they are updated with every package release.

The `metan` package is constructed on an object-oriented approach, which allows for -among other things-the reliable use of S3 generic functions such as `plot(), predict()`and `print()`. These functions can be called any time to inspect and visualize a model. All functions in `metan` have a non-standard evaluation, where the expressions are evaluated in the specified data frame rather than in the current or global environments, thus avoiding ambiguity in input data. This makes it possible to evaluate code in non-standard ways. Basically, we can pass the argument as an expression rather than a value, reducing the amount of typing.

In `metan`, all functions have as first argument the input data. So, all of them work naturally with the forward-pipe operator `%>%` (Bache & Wickham, 2014), which makes the typing cleaner and more logical. Most of MET analyze more than one trait in each genotype. Thus, when possible, functions in `metan` analyze a vector of variables and return the results into a list, saving a lot of time and code when several variables need to be analyzed. In `metan`, if we want to compute the AMMI stability value (Purchase, Hatting, & Deventer, 2000) for several traits, we can combine the functions `performs_ammi(), AMMI_indexes(),` and `get_model_data()` with `%>%` to get a two-way table with the statistic for each genotype and traits (see an example in Appendix S1, item 8.5.4). To our current knowledge, no other package designed for MET analysis presents these features.

Sometimes in MET, a certain analysis needs to be run for each level of a factor, e.g., compute a path analysis or check outliers for each environment of the trial. The R base function `subset()` could be useful, but worryingly tedious if a large number of levels need to be evaluated. Users of `metan` can count with the function `split_factors()`, which split the original data into *n* subsets according to the grouping variable(s), where *n* is the total number of combinations of the factors used. The object of class `split_factors` can be passed on to several functions `%>%`. If a function recognizes such class of data them it will take care of details and compute what is required for each one of the *n* levels (See an example in Appendix S1, item 6.3).

### 2.1 Checking data

It is assumed that MET data has the following structure (columns): **ENV**, a factor with *e* levels, being *e* the number of environments; **GEN** a factor with *g* levels, being *g* the number of genotypes; **REP** a factor with *r* levels, being *r* the number of replicates within each environment; and at least one numeric variable, e.g., grain yield. The expected number of rows in a typical MET data is then *e* × *g* × *r*.

The function `inspect()` scans all columns of a data frame object for errors that may affect the use of functions in `metan` and return a warning if (i) the data has less than three columns as factor; (ii) the data has less than the expected number of rows based on the levels of factor variables; (iii) any variable has missing values; (iv) any possible outliers is detected. Running `inspect()` is an optional and exploratory step that flags potential issues before analysis. Error check results are summarised in the R console as warnings while a plot (Fig. 1a) can also be created by using the argument `plot = TRUE` in the function (See more details in Appendix S1, item 6.1).

Outliers may violate the assumption of identically distributed errors in ANOVA models. Anomalous values tend to increase the estimate of sample variance, thus lowering the chance of rejecting the null hypothesis. In this regard, we strongly recommend checking for outliers, especially if the function `inspect()` returned a warning about them. Users of `metan` can use the function `find_outliers()` to check for possible outliers in a numeric variable, returning a summary in the console (Appendix S1, item 6.2) and a plot (Fig. 1b) if `plots = TRUE` is used.

Descriptive statistics help researchers to describe and understand the structure of a MET data. The function `desc_stat()` computes a total of 30 statistics and when combined with `split_factors()` can be used to implement a descriptive analysis for each level of a factor, e.g., for each genotype (See more details in Appendix S1, item 6.3).

Frequently in MET analysis two-way tables (e.g., genotypes in rows and environments in columns) need to be created to serve as data input in some procedure, for example, in the R package `GGEBiplots`. The function `make_mat()` can be used to create such a table. You inform the data frame in the “*long*” format, the two variables to be mapped to rows and columns and one numeric variable from which the values will fill the table and `make_mat()` take care of the details. Conversely, `make_long()` can be used to quickly convert a “*wide*” table to a “*long*” data frame (See an example in Appendix S1, item 6.4).

### 2.2 Analyzing individual environments

Individual analysis performed within each environment gives to researchers important information regarding the performance of genotypes in such environments. Provided that a typical MET data is available, the function `anova_ind()` can be used to compute, for each environment, a fixed-effect ANOVA considering a Randomized Complete Block design. The function returns the significance of factors, coefficient of variation, heritability, and accuracy of selection (See a numeric example in Appendix S1, item 7).

The function `gamem()` is used to specifically analyze genotypes using a mixed-effect model considering both a randomized complete block design or an alpha-lattice design (Patterson & Williams, 1976). The function `get_model_data()` can be used to extract the model information such as variance components, genetic parameters, and *P*-values for the Likelihood ratio test for random effects. By using the function `plot_blup()` with an object of class `gamem` the plot in Fig. 1c is produced.

### 2.3 Stability analysis

After inspecting data, checking for outliers and possibly analyzing individual environments, a quick visual inspection of the genotype–environment interaction can be performed with the function `ge_plot()`, which will generate the plots in Fig. 1m-n. Statistically, GEI can be checked in a joint analysis of variance performed with the function `anova_joint()` (Appendix S1, item 7). If GEI is significant, then it is reasonable to proceed with some stability analysis to explore such interaction. `metan` provides a collection of functions to implement widely used methods for stability analysis in the evaluation of multi-environment trials (Table 1).

**Table 1.**
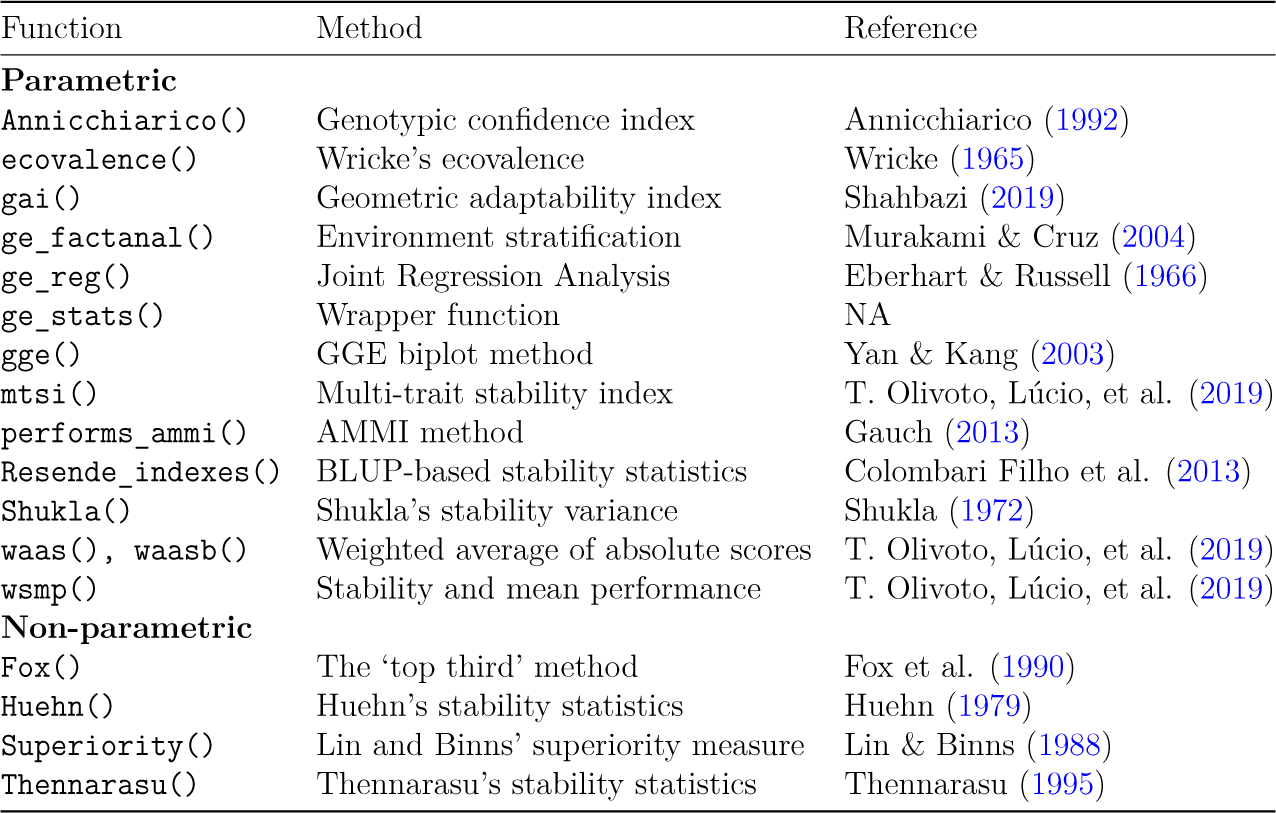
Functions available in metan version 1.1.0 for computing stability analysis.

After fitting a model, users can obtain custom plots to interpret the GEI. By invoking `plot()` in an object of class `performs_ammi()` residual plots (Fig. 1d) can be obtained. In AMMI analysis, biplots (Fig. 1f) are produced with the function `plot_scores()`, provided that an object of class `performs_ammi, waas` or `waasb` is available in the Global Environment (See Appendix S1, item 8.5.3 for more details). In GGE models, fitted with the function `gge()`, 10 types of biplots (Yan & Kang, 2003) can be created. Fig. 1g shows the biplot type 8, used for ranking genotypes. All plots are produced with package `ggplot2` (Wickham, 2016). So, users of `metan` can count on the high level of personalization provided by `ggplot2` to change any non-data elements of your plot (See an example in Appendix S1, item 7.5.3).

Users who research the associations between stability indexes (e.g., Woyann et al., 2018; Bornhofen et al., 2017; Freiria et al., 2018; Shahbazi, 2019) often find difficulties in computing the set of statistics and binding them into a “*ready-to-read*” file. `metan` provides an efficient solution for doing that. The function `ge_stats()` is a wrapper function and can be used to compute all the stability methods shown in Table 1 at once. Then, users can use `get_model_data()` to extract either the statistics or the ranks related to each genotype in each index and variable –if multiple variables are used in `ge_stat()`–, or `corr_stab_ind()`, to compute a Spearman’s rank correlation matrix between the computed stability indexes (See Appendix S1, item 8.9 for more details).

### 2.4 Biometrical models

Multi-environment trials often generate data on several traits, and this data should be exploited. In breeding trials (as well as in many other areas), indirect selection helps geneticists and breeders to select superior genotypes (Meira et al., 2017; T. Olivoto, Nardino, et al., 2017; T. Olivoto, Souza, et al., 2017; Ferrari et al., 2018; Santos et al., 2018; Fonseca, Lima, Dardengo, Silva, & Xavier, 2019; Gediya et al., 2019; Lopes Costa, Melo, & Oliveira Mano, 2019); thus, any tool that facilitates this work is welcome. `metan` provides useful functions for implementing biometrical models easily. This includes the functions `corr_coef()` for computing Pearson product-moment correlation with *P*-values, `lpcor()` for computing partial correlation coefficients; `covcor_design()` for computing phenotypic, genotypic, and residual (co)variance/correlation matrices based on designed experiments; `can_cor()` for computing canonical correlation analysis; `path_coeff()` for computing path coefficients; `corr_ss()` for sample size planning; `corr_plot()` for a mixed (text and plot) visualization of a correlation matrix (Fig. 1j); `corr_ci()` for computing nonparametric confidence intervals of Pearson’s correlation (Fig. 1k); and `clustering()` for clustering analysis (Fig. 1l).

Since `metan` was conceived for multi-environment trial analysis, the function `split_factors()` can be used to pass grouped data allowing, for example, that a path analysis or a canonical correlation be computed within each level of a factor, as shown in Santos et al. (2018). For more details, please, refer to Appendix S1, item 7.

### 2.5 Data visualization

`metan` provides useful functions for creating quickly typical plots of two-way data, such as those observed in MET data. The function `ge_plot()` can be used for a visual inspection of the GEI (Fig. 1m-n). The function `plot_factbars()` is used to create bar plots with two factor variables (Fig. 1o). `plot_factbars()` has as mandatory arguments only the data, the factors 1 and 2, and the response variable. Similarly, line plots with options for fitting different polynomial degrees can be made with the function `plot_factlines()`. In an experiment with two quantitative factors, the function `resp_surf()` can be used to fit a response surface model; Then a surface plot (Fig. 1p) can be created with `plot()` (See more details in Appendix S1, item 10).

## 3 Concluding remarks and future improvments

The package `metan` was designed to facilitate the analysis of multi-environment trials, allowing for more effective and less time-consuming handling and processing of MET datasets that have been increasing rapidly in the last years. Users will find in `metan` a complete framework to implement the most used parametric and non-parametric stability statistics for MET analysis. The package implements stability methods not available in any other R package, including the estimation of BLUP-based stability statistics (Colombari Filho et al., 2013), newer stability methods such as the weighted average of absolute scores from the (T. Olivoto, Lúcio, et al., 2019), the multi-trait stability index (Olivoto et al., 2019), and the implementation of cross-validation procedures for AMMI and BLUP models (Piepho, 1994). `metan` can also be useful for to a lot of other researchers since it provides options for implementing worldwide used multivariate statistics, e.g., path analysis, linear, partial and canonical correlations, thus allowing exploiting the maximum of (good or bad) information that a data set can offer. The estimation of stability indexes for several variables at once and the estimation of biometrical models for each level of a factor makes `metan` to outperform already published R packages for MET analysis. These features will reduce the amount of coding and save the precious time of the researchers when running their analyzes. The `metan` package is (and will always be) extensively documented online, with transparent and fully reproducible examples. `metan` is currently under active development; so, new functions will be implemented in the near future. Our next efforts will be focused on implementing cross-validation procedures for GGE models, allowing cross-validation to run in parallel, and increasing the number of stability methods available.

## Supporting information

available at

## 4 Acknowledgment

We thank the National Council for Scientific and Technological Development (CNPq) and Coordination for the Improvement of Higher Education Personnel (CAPES) for fellowships and grants to the authors. The authors have no conflicts of interest to declare.

## 5 Author’s contributions

T.O conceived the ideas, authored the software and manuscript; A.D.L assisted in the implementation of methods, and critically revised the manuscript; both authors gave final approval for publication.

## 6 Data accessibility

The `metan` R package is open-source and available on GitHub (https://github.com/TiagoOlivoto/metan). Package vignettes are also open-source, accessible at https://tiagoolivoto.github.io/metan/. Installing and loading `metan` will automatically load all example data used in this paper. Since the package is updated regularly, all code, data, and documentation used in this manuscript have been archived at https://doi.org/10.5281/zenodo.3548917 as `metan` version 1.1.0.

## 7 Supporting information

Additional supporting information may be found online in the Supporting Information section at the end of the article.

