## Supplementary material for "metan: an R package for multi-environment trial analysis": available at: colindiag.R

```
#' Collinearity Diagnostics
#'
#' Perform a (multi)collinearity diagnostic of a correlation matrix of predictor
#' variables using several indicators, as shown by Olivoto et al. (2017).
#'
#'
#' @param .data The data to be analyzed. Must be a symmetric correlation matrix
#'   or, a dataframe containing the predictor variables, or an object of class
#'   \code{split_factors}.
#' @param ... Variables to use in the correlation. If \code{...} is null then
#'   all the numeric variables from \code{.data} are used. It must be a single
#'   variable name or a comma-separated list of unquoted variables names.
#' @param by One variable (factor) to split the data into subsets. The function
#'   is then applied to each subset and returns a list where each element
#'   contains the results for one level of the variable in \code{by}. To split
#'   the data by more than one factor variable, use the function
#'   \code{\link{split_factors}} to pass subsetted data to \code{.data}.
#' @param n If a correlation matrix is provided, then \code{n} is the number of
#'   objects used to compute the correlation coefficients.
#' @param verbose If \code{verbose = TRUE} then some results are shown in the
#'   console.
#' @return
#'
#' The following values are returned. Please, note that if a grouping variable
#' is used, then the results are returned into a list.
#'
#' * \strong{cormat} A symmetric Pearson's coefficient correlation matrix
#' between the variables
#'
#' * \strong{corlist} A hypothesis testing for each of the correlation
#' coefficients
#'
#' * \strong{evalevet} The eigenvalues with associated eigenvectors of the
#' correlation matrix
#'
#' * \strong{VIF} The Variance Inflation Factors, being the diagonal elements of
#' the inverse of the correlation matrix.
#'
#' * \strong{CN} The Condition Number of the correlation matrix, given by the
#' ratio between the largest and smallest eigenvalue.
#'
#' * \strong{det} The determinant of the correlation matrix.
#'
#' * \strong{largest_corr} The largest correlation (in absolute value) observed.
#'
#' * \strong{smallest_corr} The smallest correlation (in absolute value)
#' observed.
#'
#' * \strong{weight_var} The variables with largest eigenvector (largest weight)
#' in the eigenvalue of smallest value, sorted in decreasing order.
#' @md
#' @author Tiago Olivoto \email{tiagoolivoto@@gmail.com}
#' @references Olivoto, T., V.Q. Souza, M. Nardino, I.R. Carvalho, M. Ferrari,
#'   A.J. Pelegrin, V.J. Szareski, and D. Schmidt. 2017. Multicollinearity in
#'   path analysis: a simple method to reduce its effects. Agron. J.
#'   109:131-142. doi:10.2134/agronj2016.04.0196.
#'   \href{https://dl.sciencesocieties.org/publications/aj/abstracts/109/1/131}{doi:10.2134/agronj2016.04.0196}
#'
#' @references Olivoto, T., M. Nardino, I.I.R. Carvalho, D.N. Follmann, M.
#'   Ferrari, A.J. de Pelegrin, V.J. Szareski, A.C. de Oliveira, B.O. Caron, and
#'   V.Q. de Souza. 2017. Optimal sample size and data arrangement method in
#'   estimating correlation matrices with lesser collinearity: A statistical
#'   focus in maize breeding. African J. Agric. Res. 12:93-103.
#'   \href{https://academicjournals.org/journal/AJAR/article-abstract/5DB554B62359}{doi:10.5897/AJAR2016.11799}.
#'
#' @export
#' @examples
#'\donttest{
#' # Using the correlation matrix
#' library(metan)
#'
#' cor_iris <- cor(iris[,1:4])
#' n <- nrow(iris)
#'
#' col_diag <- colindiag(cor_iris, n = n)
#'
#'
#' # Using a data frame
#' col_diag_gen <- data_ge2 %>%
#'                 split_factors(GEN) %>%
#'                 colindiag()
#'
#' # Diagnostic by levels of a factor selecting desired variables
#' col_diag_gen <- data_ge2 %>%
#'                 split_factors(GEN) %>%
#'                 colindiag(EH, PH, CD, CL)
#'}
colindiag <- function(.data,
                      ...,
                      by = NULL,
                      n = NULL,
                      verbose = TRUE) {
  if (!any(class(.data) %in% c("matrix", "data.frame", "split_factors",
                               "covcor_design", "tbl_df"))) {
    stop("The object 'x' must be a correlation matrix, a data.frame or an object of class split_factors")
  }
  if (is.matrix(.data) && is.null(n)) {
    stop("You have a matrix but the sample size used to compute the correlations (n) was not declared.")
  }
  if (is.data.frame(.data) && !is.null(n)) {
    stop("You cannot informe the sample size because a data frame was used as input.")
  }
  if (is.split_factors(.data) && !is.null(n)) {
    stop("You cannot informe the sample size because a data frame was used as input.")
  }
  internal <- function(x) {
    if (is.matrix(x)) {
      cor.x <- x
    }
    if (is.data.frame(x)) {
      cor.x <- cor(x)
      n <- nrow(x)
    }
    eigen <- eigen(cor.x)
    Det <- det(cor.x)
    NC <- max(eigen$values)/min(eigen$values)
    Aval <- data.frame(eigen$values)
    names(Aval) <- "Eigenvalues"
    Avet <- data.frame(t(eigen$vectors))
    names(Avet) <- colnames(x)
    AvAvet <- cbind(Aval, Avet)
    VIF <- data.frame(diag(solve_svd(cor.x)))
    names(VIF) <- "VIF"
    results <- data.frame(linear = as.vector(t(cor.x)[lower.tri(cor.x,
                                                                diag = F)]))
    results <- dplyr::mutate(results, t = linear * (sqrt(n -
                                                           2))/(sqrt(1 - linear^2)), prob = 2 * (1 - pt(abs(t),
                                                                                                        df = n - 2)))
    names <- colnames(x)
    combnam <- combn(names, 2, paste, collapse = " x ")
    rownames(results) <- names(sapply(combnam, names))
    largest_corr <- paste0(rownames(results)[which.max(abs(results$linear))],
                           " = ", round(results[which.max(abs(results$linear)),
                                                1], 3))
    smallest_corr <- paste0(rownames(results)[which.min(abs(results$linear))],
                            " = ", round(results[which.min(abs(results$linear)),
                                                 1], 3))
    ncorhigh <- sum(results$linear >= abs(0.8))
    if (verbose == TRUE) {
      if (NC > 1000) {
        cat(paste0("Severe multicollinearity in the matrix! Pay attention on the variables listed bellow\n",
                   "CN = ", round(NC, 3), "\n"))
      }
      if (NC < 100) {
        cat(paste0("Weak multicollinearity in the matrix\n",
                   "NC = ", round(NC, 3), "\n"))
      }
      if (NC > 100 & NC < 1000) {
        cat(paste0("The multicollinearity in the matrix should be investigated.\n",
                   "NC = ", round(NC, 3), "\n", "Largest VIF = ",
                   max(VIF), "\n"))
      }
    }
    ultimo <- data.frame(Peso = t(AvAvet[c(nrow(AvAvet)),
                                         ])[-c(1), ])
    abs <- data.frame(Peso = abs(ultimo[, "Peso"]))
    rownames(abs) <- rownames(ultimo)
    ultimo <- abs[order(abs[, "Peso"], decreasing = T), ,
                  drop = FALSE]
    pesovarname <- paste(rownames(ultimo), collapse = " > ")
    final <- list(cormat = data.frame(cor.x), corlist = results,
                  evalevet = AvAvet, VIF = VIF, CN = NC, det = Det,
                  largest_corr = largest_corr, smallest_corr = smallest_corr,
                  weight_var = pesovarname)
    if (verbose == TRUE) {
      cat(paste0("Matrix determinant: ", round(Det, 7)), "\n")
      cat(paste0("Largest correlation: ", largest_corr), "\n")
      cat(paste0("Smallest correlation: ", smallest_corr), "\n")
      cat(paste0("Number of VIFs > 10: ", length(which(VIF > 10))), "\n")
      cat(paste0("Number of correlations with r >= |0.8|: ", ncorhigh), "\n")
      cat(paste0("Variables with largest weight in the last eigenvalues: \n", pesovarname), "\n")
    }
    return(invisible(final))
  }

  if (is.matrix(.data)) {
    out <- internal(.data)
  }
  if (!missing(by)){
    if(length(as.list(substitute(by))[-1L]) != 0){
      stop("Only one grouping variable can be used in the argument 'by'.\nUse 'split_factors()' to pass '.data' grouped by more than one variable.", call. = FALSE)
    }
    .data <- split_factors(.data, {{by}}, verbose = FALSE, keep_factors = TRUE)
  }
  if (any(class(.data) %in% c("split_factors", "covcor_design"))) {
    if(!missing(...)){
      dfs <- lapply(.data[[1]], function(x){
        select_cols(x, ...) %>%
          select_numeric_cols()
      })
    } else{
      dfs <- lapply(.data[[1]], function(x){
        select_numeric_cols(x)
      })
    }
    out <- lapply(seq_along(dfs), function(x) {
      if (verbose == TRUE) {
        cat("\n----------------------------------------------------------------------------\n")
        cat("Level:", names(dfs)[[x]], "\n")
        cat("----------------------------------------------------------------------------\n")
      }
      internal(dfs[[x]])
    })
    names(out) <- names(dfs)
  }
  if (is.data.frame(.data)) {
    if(!missing(...)){
      dfs <-  select_cols(.data, ...) %>%
        select_numeric_cols()
    } else{
      if (verbose == TRUE) {
        if (sum(lapply(.data, is.factor) == TRUE) > 0) {
          warning("The factors ", paste0(collapse = " ", names(.data[, unlist(lapply(.data, is.factor))])),
                  " where ignored.  Use 'split_factors()' to perform an analysis for each level of a factor.")
        }
      }
      dfs <- select_numeric_cols(.data)
    }
    out <- internal(dfs)
  }
  invisible(out)
}
```
