## Supplementary material for "metan: an R package for multi-environment trial analysis": available at: comb_vars.R

```
#' Pairwise combinations of variables
#'
#' Pairwise combinations of variables that will be the result of a function
#' applied to each combination.
#'
#'
#' @param .data A matrix of data with, say, p columns.
#' @param order The order on how the results will appear in the output. Default
#'   is \code{order = 'first'}. In this case, assuming that .data has four
#'   columns, namely, \code{V1, V2, V3, V4}, the order of columns in the output
#'   will be \code{V1.V2, V1.V3, V1.V4, V2.V3, V2.V4, V3.V4}. If \code{order =
#'   'second'}, the result will be then \code{V1.V2, V1.V3, V2.V3, V1.V4, V2.V4,
#'   V3.V4}.
#' @param FUN The function that will be applied to each combination. The default
#'   is \code{+}, i.e., V1 + V2.
#' @param verbose Logical argument. If \code{verbose = FALSE} the code will run
#'   silently.
#' @return A data frame containing all possible combination of variables. Each
#'   combination is the result of the function in \code{FUN} applied to the two
#'   variables.
#' @author Tiago Olivoto \email{tiagoolivoto@@gmail.com}
#' @export
#' @examples
#'
#' library(metan)
#' data <- data.frame(A = rnorm(n = 5, mean = 10, sd = 3),
#'                   B = rnorm(n = 5, mean = 120, sd = 30),
#'                   C = rnorm(n = 5, mean = 40, sd = 10),
#'                   D = rnorm(n = 5, mean = 1100, sd = 200),
#'                   E = rnorm(n = 5, mean = 2, sd = 1))
#' comb1 <- comb_vars(data)
#' comb2 <- comb_vars(data, FUN = '*', order = 'second')
#'
comb_vars <- function(.data, order = "first", FUN = "+", verbose = TRUE) {
  FUN <- match.fun(FUN)
  if (!order %in% c("first", "second")) {
    stop("The orde must be one of 'first' or 'second'.", call. = FALSE)
  }
  if (verbose == TRUE) {
    if (sum(lapply(.data, is.factor) == TRUE) > 0) {
      warning("The columns factors ",
              paste0(collapse = " ",names(.data[, unlist(lapply(.data, is.factor))])),
              " where deleted. Only the numeric variables were used.")
    }
  }
  x <- .data[, unlist(lapply(.data, is.numeric))] %>% as.data.frame()
  cb <- data.frame(t(combn(ncol(x), 2)))
  if (order == "second") {
    cb %<>% arrange(X2)
  }
  nvars <- data.frame(matrix(nrow = nrow(x), ncol = nrow(cb)))
  for (i in 1:nrow(cb)) {
    nvars[, i] <- FUN(x[[cb[i, 1]]], x[[cb[i, 2]]])
    colnames(nvars)[[i]] <- paste(colnames(x[cb[i, 1]]), colnames(x[cb[i, 2]]), sep = "x")
  }
  return(as_tibble(nvars, rownames = NA))
}
```
