## Supplementary material for "metan: an R package for multi-environment trial analysis": available at: corr_stab_ind.R

```
#' Correlation between stability indexes
#'
#' Computes the Spearman's rank correlation between the parametric and
#' nonparametric stability indexes computed with the function
#' \code{\link{ge_stats}}.
#'
#' @param x An object of class \code{ge_stats}.
#' @param stats The statistics to compute the correlation. See the section
#'   \strong{Details} for more information.
#' @param plot Plot the heat map with the correlations? Defaults to \code{TRUE}.
#' @param ... Other arguments to be passed to the function
#'   \code{\link{plot.corr_coef}}.
#' @details The argument \code{stats} is used to chose the statistics to show
#'   the ranks. Allowed values are \code{"all"} (All statistics, default),
#'   \code{"par"} (Parametric statistics), \code{"nonpar"} (Non-parametric
#'   statistics), \code{"ammi"} (AMMI-based stability statistics), or the
#'   following values that can be combined into comma-separated character
#'   vector. \code{"Y"} (Response variable), \code{"Var"} (Genotype's variance),
#'   \code{"Shukla"} (Shukla's variance), \code{"Wi_g", "Wi_f", "Wi_u"}
#'   (Annichiarrico's genotypic confidence index for all, favorable and
#'   unfavorable environments, respectively), \code{"Ecoval"} (Wricke's
#'   ecovalence), \code{"Sij"} (Deviations from the joint-regression analysis),
#'   \code{"R2"} (R-squared from the joint-regression analysis), \code{"ASV"}
#'   (AMMI-stability value), \code{"SIPC"} (sum of the absolute values of the
#'   IPCA scores), \code{"EV"} (Average of the squared eigenvector values),
#'   \code{"ZA"} (Absolute values of the relative contributions of the IPCAs to
#'   the interaction), \code{"WAAS"} (Weighted Average of Absolute Scores),
#'   \code{"HMGV"} (Harmonic mean of the genotypic value), \code{"RPGV"}
#'   (Relative performance of the genotypic values), \code{"HMRPGV"} (Harmonic
#'   mean of the relative performance of the genotypic values), \code{"Pi_a",
#'   "Pi_f", "Pi_u"} (Superiority indexes for all, favorable and unfavorable
#'   environments, respectively), \code{"Gai"} (Geometric adaptability index),
#'   \code{"S1"} (mean of the absolute rank differences of a genotype over the n
#'   environments), \code{"S2"} (variance among the ranks over the k
#'   environments), \code{"S3"} (sum of the absolute deviations), \code{"S6"}
#'   (relative sum of squares of rank for each genotype), \code{"N1", "N2",
#'   "N3", "N4"} (Thennarasu"s statistics)).
#' @return A list with the data (ranks) correlation, p-values and a heat map showing the
#'   correlation coefficients.
#' @export
#' @author Tiago Olivoto \email{tiagoolivoto@@gmail.com}
#'
#' @examples
#' \donttest{
#' library(metan)
#' model <- ge_stats(data_ge, ENV, GEN, REP, GY)
#' a <- corr_stab_ind(model)
#' b <- corr_stab_ind(model, stats = "ammi")
#' c <- corr_stab_ind(model, stats = c("ASV, Sij, R2, WAAS, N1"))
#' }
#'
corr_stab_ind <- function(x, stats = "all", plot = TRUE, ...){
  all_s <- c("Y", "Var", "Shukla", "Wi_g", "Wi_f", "Wi_u", "Ecoval", "Sij", "R2", "ASV", "SIPC", "EV", "ZA", "WAAS", "HMGV", "RPGV", "HMRPGV", "Pi_a", "Pi_f", "Pi_u", "Gai", "S1", "S2", "S3", "S6", "N1", "N2", "N3", "N4")
  par_s <- c("Y", "Var", "Shukla", "Wi_g", "Wi_f", "Wi_u", "Ecoval", "Sij", "R2", "ASV", "SIPC", "EV", "ZA", "WAAS", "HMGV", "RPGV", "HMRPGV")
  nonpar_s <- c("Y", "Pi_a", "Pi_f", "Pi_u", "Gai", "S1", "S2", "S3", "S6", "N1", "N2", "N3", "N4" )
  ammi_s <- c("Y", "ASV", "SIPC", "EV", "ZA", "WAAS")
  if(!stats %in% c("all", "par", "nonpar", "ammi")){
    stats = unlist(strsplit(stats, split=", "))
  } else {
    if(any(stats == "all")){
      stats = all_s
    }
    if(any(stats == "par")){
      stats = par_s
    }
    if(any(stats == "nonpar")){
      stats = nonpar_s
    }
    if(any(stats == "ammi")){
      stats = ammi_s
    }
  }
  if(any(!stats %in% c("all", "par", "nonpar", "ammi", all_s)) == TRUE){
    stop("Argument 'stats' with invalid values. See ?corr_stab_ind for more details.", call. = FALSE)
  }
  bind <- do.call(
    cbind,
    lapply(x, function(x) {
      x %>% select(contains("_R"))
    })) %>%
    as_tibble() %>%
    mutate(gen = x[[1]][["GEN"]]) %>%
    pivot_longer(cols = contains(".")) %>%
    separate(name, into = c("var", "stat"), sep = "(\\.)") %>%
    separate(stat, into = "stat", sep = "_(?=[^_]*$)", extra = "drop") %>%
    pivot_wider(values_from = value, names_from = stat) %>%
    select(-c(var, gen))
  bind <- select(bind, stats)
  corr <- corr_coef(bind)
  p <- plot(corr, legend.title = "Speraman's\nCorrelation", ...)
  if(plot == TRUE){
    suppressWarnings(print(p))
  }
  invisible(structure(list(data = bind,
                        corr = corr$cor,
                        pval = corr$pval,
                        plot = p),
                   class = "corr_stab"))
}
```
