## Supplementary material for "metan: an R package for multi-environment trial analysis": available at: cv_ammi.R

```
#' Cross-validation procedure
#'
#' Cross-validation for estimation of AMMI models
#'
#' THe original dataset is split into two datasets: training set and validation
#' set. The 'training' set has all combinations (genotype x environment) with
#' N-1 replications. The 'validation' set has the remaining replication. The
#' splitting of the dataset into modeling and validation sets depends on the
#' design informed. For Completely Randomized Block Design (default), and
#' alpha-lattice design (declaring \code{block} arguments), complete replicates
#' are selected within environments. The remained replicate serves as validation
#' data. If \code{design = 'RCD'} is informed, completely randomly samples are
#' made for each genotype-by-environment combination (Olivoto et al. 2019). The
#' estimated values considering \code{naxis}-Interaction Principal Component
#' Axis are compared with the 'validation' data. The Root Mean Square Prediction
#' Difference (RMSPD) is computed. At the end of boots, a list is returned.
#'
#' @param .data The dataset containing the columns related to Environments,
#'   Genotypes, replication/block and response variable(s).
#' @param env The name of the column that contains the levels of the
#'   environments.
#' @param gen The name of the column that contains the levels of the genotypes.
#' @param rep The name of the column that contains the levels of the
#'   replications/blocks.
#' @param resp The response variable.
#' @param block Defaults to \code{NULL}. In this case, a randomized complete
#'   block design is considered. If block is informed, then a resolvable
#'   alpha-lattice design (Patterson and Williams, 1976) is employed.
#'   \strong{All effects are assumed to be fixed.}
#' @param nboot The number of resamples to be used in the cross-validation.
#'   Defaults to 200.
#' @param naxis The number of axis to be considered for estimation of GE
#'   effects.
#' @param design The experimental design. Defaults to \code{RCBD} (Randomized
#'   complete Block Design). For Completely Randomized Designs inform
#'   \code{design = 'CRD'}.
#' @param verbose A logical argument to define if a progress bar is shown.
#'   Default is \code{TRUE}.
#' @references Olivoto, T., A.D.C. L{\'{u}}cio, J.A.G. da silva, V.S. Marchioro,
#'   V.Q. de Souza, and E. Jost. 2019. Mean performance and stability in
#'   multi-environment trials I: Combining features of AMMI and BLUP techniques.
#'   Agron. J. 111:2949-2960.
#'   \href{https://dl.sciencesocieties.org/publications/aj/abstracts/0/0/agronj2019.03.0220?access=0&view=pdf}{doi:10.2134/agronj2019.03.0220}
#'
#' @references Patterson, H.D., and E.R. Williams. 1976. A new class of
#'   resolvable incomplete block designs. Biometrika 63:83-92.
#' @return An object of class \code{cv_ammi} with the following items: *
#' \strong{RMSPD}: A vector with nboot-estimates of the Root Mean Squared
#' Prediction Difference between predicted and validating data.
#'
#' * \strong{RMSPDmean}: The mean of RMSPDmean estimates.
#'
#' * \strong{Estimated}: A data frame that contain the values (predicted,
#' observed, validation) of the last loop.
#'
#' * \strong{Modeling}: The dataset used as modeling data in the last loop
#'
#' * \strong{Testing}: The dataset used as testing data in the last loop.
#' @md
#' @author Tiago Olivoto \email{tiagoolivoto@@gmail.com}
#' @seealso \code{\link{cv_ammif}, \link{cv_blup}}
#' @importFrom progress progress_bar
#' @export
#' @examples
#'
#' \donttest{
#' library(metan)
#' model <- cv_ammi(data_ge,
#'                 env = ENV,
#'                 gen = GEN,
#'                 rep = REP,
#'                 resp = GY,
#'                 nboot = 10,
#'                 naxis = 2)
#'
#' # Alternatively using the pipe operator %>%
#' model <- data_ge %>%
#'          cv_ammi(ENV, GEN, REP, GY)
#' }
#'
#'
cv_ammi <- function(.data, env, gen, rep, resp, block = NULL, naxis = 2, nboot = 200, design = "RCBD", verbose = TRUE) {
  if(missing(block)){
    if (!design %in% c("RCBD", "CRD")) {
      stop("Incorrect experimental design informed! Plesease inform RCBD for randomized complete block or CRD for completely randomized design.")
    }
    data <- .data %>%
      dplyr::select(ENV = {{env}},
                    GEN = {{gen}},
                    REP = {{rep}},
                    Y = {{resp}})%>%
      mutate_at(1:3, as.factor)
    RMSPDres <- data.frame(RMSPD = matrix(0, nboot, 1))
    data <- tibble::rowid_to_column(data)
    Nenv <- length(unique(data$ENV))
    Ngen <- length(unique(data$GEN))
    Nbloc <- length(unique(data$REP))
    nrepval <- Nbloc - 1
    minimo <- min(Nenv, Ngen) - 1
    if (naxis > minimo) {
      stop("The number of axis to be used must be lesser than or equal to ",
           minimo, " [min(GEN-1;ENV-1)]")
    }
    if (verbose == TRUE) {
      pb <- progress_bar$new(
        format = "Validating :current of :total sets [:bar] :percent (:elapsedfull -:eta left)",
        clear = FALSE, total = nboot, width = 80)
    }
    for (b in 1:nboot) {
      if (design == "CRD") {
        X <- sample(1:10000, 1)
        set.seed(X)
        modeling <- data %>%
          dplyr::group_by(ENV, GEN) %>%
          dplyr::sample_n(nrepval, replace = FALSE) %>%
          arrange(rowid) %>%
          as.data.frame()
        rownames(modeling) <- modeling$rowid
      }
      if (design == "RCBD") {
        tmp <- split_factors(data, ENV, keep_factors = TRUE, verbose = FALSE)
        modeling <- do.call(rbind, lapply(tmp[[1]], function(x) {
          X2 <- sample(unique(data$REP), nrepval, replace = FALSE)
          x %>% dplyr::group_by(GEN) %>% dplyr::filter(REP %in% X2)
        })) %>% as.data.frame()
        rownames(modeling) <- modeling$rowid
      }
      testing <- anti_join(data, modeling, by = c("ENV", "GEN", "REP", "Y", "rowid")) %>%
        arrange(ENV, GEN) %>%
        as.data.frame()
      MEDIAS <- modeling %>%
        group_by(ENV, GEN) %>%
        summarise(Y = mean(Y)) %>%
        as.data.frame()
      residual <- modeling %>%
        group_by(ENV, GEN) %>%
        summarise(Y = mean(Y)) %>%
        ungroup() %>%
        mutate(residuals = residuals(lm(Y ~ ENV + GEN, data = .))) %>%
        pull(residuals)
      s <- svd(t(matrix(residual, Nenv, byrow = T)))
      MGEN <- model.matrix(~factor(testing$GEN) - 1)
      MENV <- model.matrix(~factor(testing$ENV) - 1)
      MEDIAS %<>% mutate(Ypred = Y - residual,
                         ResAMMI = ((MGEN %*% s$u[, 1:naxis]) * (MENV %*% s$v[, 1:naxis])) %*% s$d[1:naxis],
                         YpredAMMI = Ypred + ResAMMI,
                         testing = testing$Y,
                         error = YpredAMMI - testing,
                         errrorAMMI0 = Ypred - testing)
      RMSPDres[, 1][b] <- ifelse(naxis == 0, sqrt(sum(MEDIAS$errrorAMMI0^2)/length(MEDIAS$errrorAMMI0)),
                                 sqrt(sum(MEDIAS$error^2)/length(MEDIAS$error)))
      if (verbose == TRUE) {
        pb$tick()
      }
    }
    RMSPDmean <- summarise(RMSPDres,
                           MODEL = paste("AMMI", naxis, sep = ""),
                           mean = mean(RMSPD),
                           sd = sd(RMSPD),
                           se = sd(RMSPD)/sqrt(n()),
                           Q2.5 = quantile(RMSPD, 0.025),
                           Q97.5 = quantile(RMSPD, 0.975))
    RMSPDres <- RMSPDres %>%
      mutate(MODEL = paste("AMMI", naxis, sep = "")) %>%
      dplyr::select(MODEL, everything())
  }

  if(!missing(block)){
    data <- .data %>%
      dplyr::select(ENV = {{env}},
                    GEN = {{gen}},
                    REP = {{rep}},
                    BLOCK = {{block}},
                    Y = {{resp}})
    RMSPDres <- data.frame(RMSPD = matrix(0, nboot, 1))
    data <- tibble::rowid_to_column(data)
    Nenv <- nlevels(data$ENV)
    Ngen <- nlevels(data$GEN)
    Nbloc <- nlevels(data$REP)
    nrepval <- Nbloc - 1
    minimo <- min(Nenv, Ngen) - 1
    if (naxis > minimo) {
      stop("The number of axis to be used must be lesser than or equal to ",
           minimo, " [min(GEN-1;ENV-1)]")
    }
    if (verbose == TRUE) {
      pb <- progress_bar$new(
        format = "Validating :current of :total sets [:bar] :percent (:elapsedfull -:eta left)",
        clear = FALSE, total = nboot, width = 80)
    }
    for (b in 1:nboot) {
      tmp <- split_factors(data, ENV, keep_factors = TRUE, verbose = FALSE)
      modeling <- do.call(rbind, lapply(tmp[[1]], function(x) {
        X2 <- sample(unique(data$REP), nrepval, replace = FALSE)
        x %>% dplyr::group_by(GEN) %>% dplyr::filter(REP %in% X2)
      })) %>% as.data.frame()
      rownames(modeling) <- modeling$rowid
      testing <- anti_join(data, modeling, by = c("ENV", "GEN", "REP", "BLOCK", "Y", "rowid")) %>%
        arrange(ENV, GEN) %>%
        as.data.frame()
      MEDIAS <- modeling %>%
        group_by(ENV, GEN) %>%
        summarise(Y = mean(Y)) %>%
        as.data.frame()
      residual <- modeling %>%
        mutate(residuals = residuals(lm(Y ~ ENV/REP + REP:BLOCK + ENV +  GEN, data = .))) %>%
        group_by(ENV, GEN) %>%
        summarise(residuals = mean(residuals)) %>%
        ungroup() %>%
        pull(residuals)
      s <- svd(t(matrix(residual, Nenv, byrow = T)))
      MGEN <- model.matrix(~factor(testing$GEN) - 1)
      MENV <- model.matrix(~factor(testing$ENV) - 1)
      MEDIAS %<>% mutate(Ypred = Y - residual,
                         ResAMMI = ((MGEN %*% s$u[, 1:naxis]) * (MENV %*% s$v[, 1:naxis])) %*% s$d[1:naxis],
                         YpredAMMI = Ypred + ResAMMI,
                         testing = testing$Y,
                         error = YpredAMMI - testing,
                         errrorAMMI0 = Ypred - testing)
      RMSPDres[, 1][b] <- ifelse(naxis == 0, sqrt(sum(MEDIAS$errrorAMMI0^2)/length(MEDIAS$errrorAMMI0)),
                                 sqrt(sum(MEDIAS$error^2)/length(MEDIAS$error)))
      if (verbose == TRUE) {
        pb$tick()
      }
    }
    RMSPDmean <- summarise(RMSPDres,
                           MODEL = paste("a_AMMI", naxis, sep = ""),
                           mean = mean(RMSPD),
                           sd = sd(RMSPD),
                           se = sd(RMSPD)/sqrt(n()),
                           Q2.5 = quantile(RMSPD, 0.025),
                           Q97.5 = quantile(RMSPD, 0.975))
    RMSPDres <- RMSPDres %>%
      mutate(MODEL = paste("a_AMMI", naxis, sep = "")) %>%
      dplyr::select(MODEL, everything())
  }

  return(structure(list(RMSPD = RMSPDres, RMSPDmean = RMSPDmean,
                        Estimated = MEDIAS, Modeling = modeling, Testing = testing),
                   class = "cvalidation"))
}
```
