## Supplementary material for "metan: an R package for multi-environment trial analysis": available at: ecovalence.R

```
#' Stability analysis based on Wricke's model
#'
#' The function computes the ecovalence (Wricke, 1965) for stability analysis.
#'
#'
#' @param .data The dataset containing the columns related to Environments,
#'   Genotypes, replication/block and response variable(s).
#' @param env The name of the column that contains the levels of the
#'   environments.
#' @param gen The name of the column that contains the levels of the genotypes.
#' @param rep The name of the column that contains the levels of the
#'   replications/blocks.
#' @param resp The response variable(s). To analyze multiple variables in a
#'   single procedure use, for example, \code{resp = c(var1, var2, var3)}.
#' @param verbose Logical argument. If \code{verbose = FALSE} the code will run
#'   silently.
#' @return An object of class \code{ecovalence} containing the results for each
#'   variable used in the argument \code{resp}.
#' @author Tiago Olivoto \email{tiagoolivoto@@gmail.com}
#' @references Wricke, G. 1965. Zur berechnung der okovalenz bei sommerweizen
#'   und hafer. Z. Pflanzenzuchtg 52:127-138.
#' @export
#' @examples
#'
#' library(metan)
#'out <- ecovalence(data_ge2,
#'                  env = ENV,
#'                  gen = GEN,
#'                  rep = REP,
#'                  resp = PH)
#'
ecovalence <- function(.data, env, gen, rep, resp, verbose = TRUE) {
  factors  <- .data %>%
    select(ENV = {{env}},
           GEN = {{gen}},
           REP = {{rep}}) %>%
    mutate_all(as.factor)
  vars <- .data %>%
    select({{resp}}) %>%
    select_numeric_cols()
  listres <- list()
  nvar <- ncol(vars)
  for (var in 1:nvar) {
    data <- factors %>%
      mutate(mean = vars[[var]])
    data2 <- data %>%
      group_by(ENV, GEN) %>%
      summarise(mean = mean(mean)) %>%
      as.data.frame()
    data3 <- mutate(data2,
                    ge = residuals(lm(mean ~ ENV + GEN, data = data2)))
    ge_effect <- make_mat(data3, GEN, ENV, ge)
    Ecoval <- rowSums(ge_effect^2 * nlevels(data$REP))
    Ecov_perc <- (Ecoval/sum(Ecoval)) * 100
    rank <- rank(Ecoval)
    temp <- cbind(ge_effect, Ecoval, Ecov_perc, rank) %>%
      as_tibble(rownames = NA) %>%
      rownames_to_column("GEN")
    if (nvar > 1) {
      listres[[paste(names(vars[var]))]] <- temp
      if (verbose == TRUE) {
        cat("Evaluating variable", paste(names(vars[var])),
            round((var - 1)/(length(vars) - 1) * 100, 1), "%", "\n")
      }
    } else {
      listres[[paste(names(vars[var]))]] <- temp
    }
  }
  return(structure(listres, class = "ecovalence"))
}
```
