## Supplementary material for "metan: an R package for multi-environment trial analysis": available at: env_dissimilarity.R

```
#' Dissimilarity between environments
#'
#' Computes the dissimilarity between environments based on several approaches.
#' See the section \strong{details} for more details.
#'
#' @param .data The dataset containing the columns related to Environments,
#'   Genotypes, replication/block and response variable(s).
#' @param env The name of the column that contains the levels of the
#'   environments.
#' @param gen The name of the column that contains the levels of the genotypes.
#' @param rep The name of the column that contains the levels of the
#'   replications/blocks.
#' @param resp The response variable(s). To analyze multiple variables in a
#'   single procedure a vector of variables may be used. For example \code{resp
#'   = c(var1, var2, var3)}. Select helpers are also allowed.
#' @return A list with the following matrices:
#' * \code{SPART_CC}: The percentage of the single (non cross-over) part of the
#' interaction between genotypes and pairs of environments according to the
#' method proposed by Cruz and Castoldi (1991).
#' * \code{CPART_CC}: The percentage of the complex (cross-over) part of the
#' interaction between genotypes and pairs of environments according to the
#' method proposed by Cruz and Castoldi (1991).
#' * \code{SPART_RO}: The percentage of the single (non cross-over) part of the
#' interaction between genotypes and pairs of environments according to the
#' method proposed by Robertson (1959).
#' * \code{CPART_RO}: The percentage of the complex (cross-over) part of the
#' interaction between genotypes and pairs of environments according to the
#' method proposed by Robertson (1959).
#' * \code{MSGE}: Interaction mean square between genotypes and pairs of
#' environments.
#' * \code{SSGE}: Interaction sum of square between genotypes and pairs of
#' environments.
#' * \code{correlation}: Correlation coefficients between genotypes's average in
#' each pair of environment.
#' @md
#' @details Roberteson (1959) proposed the partition of the mean square of the
#'   genotype-environment interaction  (MS_GE) into single (S) and complex (C)
#'   parts, where \eqn{S = \frac{1}{2}(\sqrt{Q1}-\sqrt{Q2})^2)} and \eqn{C =
#'   (1-r)\sqrt{Q1-Q2}}, being \emph{r} the correlation between the genotype's
#'   average in the two environments; and \emph{Q1} and \emph{Q2} the genotype
#'   mean square in the environments 1 and 2, respectively. Cruz and Castoldi
#'   (1991) proposed a new decomposition of the MS_GE, in which the complex part
#'   is given by \eqn{C = \sqrt{(1-r)^3\times Q1\times Q2}}.
#' @references
#' Cruz, C.D., Castoldi, F. (1991). Decomposicao da interacao genotipos x
#' ambientes em partes simples e complexa. Ceres, 38:422-430. Available at:
#' \url{http://www.ceres.ufv.br/ojs/index.php/ceres/article/view/2165}.
#'
#' Robertson, A. (1959). Experimental design on the measurement of
#' heritabilities and genetic correlations. biometrical genetics. New York:
#' Pergamon Press.
#' @export
#' @author Tiago Olivoto \email{tiagoolivoto@@gmail.com}
#' @examples
#' mod <- env_dissimilarity(data_ge, ENV, GEN, REP, GY)
#' print(mod)
env_dissimilarity <- function(.data,
                              env,
                              gen,
                              rep,
                              resp){
  factors  <- .data %>%
    select(ENV = {{env}},
           GEN = {{gen}},
           REP = {{rep}}) %>%
    mutate_all(as.factor)
  vars <- .data %>%
    select({{resp}}, -!!colnames(factors)) %>%
    select_numeric_cols()
  listres <- list()
  nvar <- ncol(vars)
  for (var in 1:nvar) {
    data <- factors %>%
      mutate(mean = vars[[var]])
    mat <- make_mat(data, GEN, ENV, mean)
    ind_anova <- anova_ind(data, ENV, GEN, REP, mean)
    NG <- nrow(mat)
    NE <- ncol(mat)
    NR <- nlevels(data$REP)
    index <- data.frame(t(combn(NE, 2)))
    # Mean square of the GE
    dist <- as.numeric(dist(t(mat))^2)
    sumg <- colSums(mat)
    SQGA <- 0
    MSGE <- matrix(NA, NE, NE)
    SSGE <- matrix(NA, NE, NE)
    for (i in 1:length(dist)){
      SQGA[i] <- 0.5 * (dist[i] - 1/NG *(sumg[index[i, 1]] - sumg[index[i, 2]])^2)
    }
    SSGE[lower.tri(SSGE, diag = F)] <- SQGA
    MSGE[lower.tri(MSGE, diag = F)] <- SQGA / (NG - 1)
    cmat <- cor(mat)
    dist_r <- as.vector(t(cor(mat))[lower.tri(cor(mat), diag = F)])
    MS_GEN <- ind_anova[["mean"]][["individual"]][["MSG"]] / NR
    C <- 0
    R <- 0
    CPARTCC <- matrix(NA, NE, NE)
    CPARTRO<- matrix(NA, NE, NE)
    for (i in 1:length(dist_r)){
      # Cruz and Castoldi (1991)
      C[i] <- sqrt((1- dist_r[i])^3 * MS_GEN[index[i, 1]] * MS_GEN[index[i, 2]])
      #Robertson (1959)
      R[i] <- (1 - dist_r[i]) * sqrt(MS_GEN[index[i, 1]] * MS_GEN[index[i, 2]])
    }
    CPARTCC[lower.tri(CPARTCC, diag = F)] <- C
    CPARTRO[lower.tri(CPARTRO, diag = F)] <- R
    CPARTCC <- t(CPARTCC)
    CPARTRO <- t(CPARTRO)
    PCINT_CC <- 0
    PCINT_RO <- 0
    CPART_PER_CC <- matrix(NA, NE, NE)
    CPART_PER_RO <- matrix(NA, NE, NE)
    for (i in 1:length(C)){
      PCINT_RO[i] <-  100 * R[i] / (SQGA[i] / (NG - 1))
      PCINT_CC[i] <- 100 * C[i] / (SQGA[i] / (NG - 1))
    }
    CPART_PER_RO[lower.tri(CPART_PER_RO, diag = F)] <- PCINT_RO
    CPART_PER_CC[lower.tri(CPART_PER_CC, diag = F)] <- PCINT_CC
    SPART_PER_CC <- 100 - CPART_PER_CC
    SPART_PER_RO <- 100 - CPART_PER_RO
    SPART_PER_CC <- make_sym(SPART_PER_CC, diag = 0)
    CPART_PER_CC <- make_sym(CPART_PER_CC, diag = 0)
    SPART_PER_RO <- make_sym(SPART_PER_RO, diag = 0)
    CPART_PER_RO <- make_sym(CPART_PER_RO, diag = 0)
    SSGE <- make_sym(SSGE, diag = 0)
    MSGE <- make_sym(MSGE, diag = 0)
    rownames(SPART_PER_CC) <- colnames(SPART_PER_CC) <- colnames(mat)
    rownames(CPART_PER_CC) <- colnames(CPART_PER_CC) <- colnames(mat)
    rownames(SPART_PER_RO) <- colnames(SPART_PER_RO) <- colnames(mat)
    rownames(CPART_PER_RO) <- colnames(CPART_PER_RO) <- colnames(mat)
    rownames(cmat) <- colnames(cmat) <- colnames(mat)
    rownames(SSGE) <- colnames(SSGE) <- colnames(mat)
    rownames(MSGE) <- colnames(MSGE) <- colnames(mat)
    listres[[paste(names(vars[var]))]] <- list(SPART_CC = SPART_PER_CC,
                                               CPART_CC = CPART_PER_CC,
                                               SPART_RO = SPART_PER_RO,
                                               CPART_RO = CPART_PER_RO,
                                               MSGE = MSGE,
                                               SSGE = SSGE,
                                               correlation = cmat)
  }
  return(structure(listres, class = "env_dissimilarity"))
}


#' Print an object of class env_dissimilarity
#'
#' Print the \code{env_dissimilarity} object in two ways. By default, the results
#' are shown in the R console. The results can also be exported to the directory
#' into a *.txt file.
#'
#' @param x An object of class \code{env_dissimilarity}.
#' @param export A logical argument. If \code{TRUE}, a *.txt file is exported to
#'   the working directory.
#' @param file.name The name of the file if \code{export = TRUE}
#' @param digits The significant digits to be shown.
#' @param ... Currently not used.
#' @author Tiago Olivoto \email{tiagoolivoto@@gmail.com}
#' @method print env_dissimilarity
#' @export
#' @examples
#'
#' library(metan)
#' mod <- env_dissimilarity(data_ge, ENV, GEN, REP, GY)
#' print(mod)
print.env_dissimilarity <- function(x, export = FALSE, file.name = NULL, digits = 3, ...) {
  if (!class(x) == "env_dissimilarity") {
    stop("The object must be of class 'env_dissimilarity'")
  }
  if (export == TRUE) {
    file.name <- ifelse(is.null(file.name) == TRUE, "Env_dissimilarity print", file.name)
    sink(paste0(file.name, ".txt"))
  }
  for (i in 1:length(x)) {
    var <- x[[i]]
    cat("Variable", names(x)[i], "\n")
    cat("----------------------------------------------------------------------\n")
    cat("Pearson's correlation coefficient\n")
    cat("----------------------------------------------------------------------\n")
    mod <- round(var$correlation, digits = digits)
    print(mod)
    cat("----------------------------------------------------------------------\n")
    lt_mod <- make_lower_tri(mod)
    index_min <- which(lt_mod == min(lt_mod, na.rm = TRUE), arr.ind = TRUE)
    index_max <- which(lt_mod == max(lt_mod, na.rm = TRUE), arr.ind = TRUE)
    cat("Minimum correlation = ",   round(min(lt_mod, na.rm = TRUE),3), paste("between environments", index_min[1], 'and', index_min[2]), "\n")
    cat("Maximum correlation = ",   round(max(lt_mod, na.rm = TRUE),3), paste("between environments", index_max[1], 'and', index_max[2]), "\n")
    cat("----------------------------------------------------------------------\n")

    cat("Mean square GxEjj'\n")
    cat("----------------------------------------------------------------------\n")
    mod <- round(var$MSGE, digits = digits)
    print(mod)
    cat("----------------------------------------------------------------------\n")
    lt_mod <- make_lower_tri(mod)
    index_min <- which(lt_mod == min(lt_mod, na.rm = TRUE), arr.ind = TRUE)
    index_max <- which(lt_mod == max(lt_mod, na.rm = TRUE), arr.ind = TRUE)
    cat("Total mean square = ",   round(sum(lt_mod, na.rm = TRUE),3), "\n")
    cat("Minimum = ",   round(min(lt_mod, na.rm = TRUE),3), paste("between environments", index_min[1], 'and', index_min[2]), "\n")
    cat("Maximum = ",   round(max(lt_mod, na.rm = TRUE),3), paste("between environments", index_max[1], 'and', index_max[2]), "\n")
    cat("----------------------------------------------------------------------\n")

    cat("% Of the single part of MS GxEjj' (Robertson, 1959)\n")
    cat("----------------------------------------------------------------------\n")
    mod <- round(var$SPART_RO, digits = digits)
    print(mod)
    cat("----------------------------------------------------------------------\n")
    lt_mod <- make_lower_tri(mod)
    index_min <- which(lt_mod == min(lt_mod, na.rm = TRUE), arr.ind = TRUE)
    index_max <- which(lt_mod == max(lt_mod, na.rm = TRUE), arr.ind = TRUE)
    cat("Average = ",   round(mean(lt_mod, na.rm = TRUE),3), "\n")
    cat("Minimum = ",   round(min(lt_mod, na.rm = TRUE),3), paste("between environments", index_min[1], 'and', index_min[2]), "\n")
    cat("Maximum = ",   round(max(lt_mod, na.rm = TRUE),3), paste("between environments", index_max[1], 'and', index_max[2]), "\n")
    cat("----------------------------------------------------------------------\n")

    cat("% Of the complex part of MS GxEjj' (Robertson, 1959)\n")
    cat("----------------------------------------------------------------------\n")
    mod <- round(var$CPART_RO, digits = digits)
    print(mod)
    cat("----------------------------------------------------------------------\n")
    lt_mod <- make_lower_tri(mod)
    index_min <- which(lt_mod == min(lt_mod, na.rm = TRUE), arr.ind = TRUE)
    index_max <- which(lt_mod == max(lt_mod, na.rm = TRUE), arr.ind = TRUE)
    cat("Average = ",   round(mean(lt_mod, na.rm = TRUE),3), "\n")
    cat("Minimum = ",   round(min(lt_mod, na.rm = TRUE),3), paste("between environments", index_min[1], 'and', index_min[2]), "\n")
    cat("Maximum = ",   round(max(lt_mod, na.rm = TRUE),3), paste("between environments", index_max[1], 'and', index_max[2]), "\n")
    cat("----------------------------------------------------------------------\n")

    cat("% Of the single part of MS GxEjj' (Cruz and Castoldi, 1991)\n")
    cat("----------------------------------------------------------------------\n")
    mod <- round(var$SPART_CC, digits = digits)
    print(mod)
    cat("----------------------------------------------------------------------\n")
    lt_mod <- make_lower_tri(mod)
    index_min <- which(lt_mod == min(lt_mod, na.rm = TRUE), arr.ind = TRUE)
    index_max <- which(lt_mod == max(lt_mod, na.rm = TRUE), arr.ind = TRUE)
    cat("Average = ",   round(mean(lt_mod, na.rm = TRUE),3), "\n")
    cat("Minimum = ",   round(min(lt_mod, na.rm = TRUE),3), paste("between environments", index_min[1], 'and', index_min[2]), "\n")
    cat("Maximum = ",   round(max(lt_mod, na.rm = TRUE),3), paste("between environments", index_max[1], 'and', index_max[2]), "\n")
    cat("----------------------------------------------------------------------\n")

    cat("% Of the complex part of MS GxEjj' (Cruz and Castoldi, 1991)\n")
    cat("----------------------------------------------------------------------\n")
    mod <- round(var$CPART_CC, digits = digits)
    print(mod)
    cat("----------------------------------------------------------------------\n")
    lt_mod <- make_lower_tri(mod)
    index_min <- which(lt_mod == min(lt_mod, na.rm = TRUE), arr.ind = TRUE)
    index_max <- which(lt_mod == max(lt_mod, na.rm = TRUE), arr.ind = TRUE)
    cat("Average = ",   round(mean(lt_mod, na.rm = TRUE),3), "\n")
    cat("Minimum = ",   round(min(lt_mod, na.rm = TRUE),3), paste("between environments", index_min[1], 'and', index_min[2]), "\n")
    cat("Maximum = ",   round(max(lt_mod, na.rm = TRUE),3), paste("between environments", index_max[1], 'and', index_max[2]), "\n")
    cat("----------------------------------------------------------------------\n")
    cat("\n\n\n")
  }
  if (export == TRUE) {
    sink()
  }
}


#' Plot an object of class env_dissimilarity
#'
#' Create dendrograms to show the dissimilarity between environments.
#'
#'
#' @param x An object of class \code{env_dissimilarity}
#' @param var The variable to plot. Defaults to \code{var = 1} the first
#'   variable of \code{x}.
#' @param nclust The number of clusters to show.
#' @param ... Other arguments bo be passed to the function \code{\link[stats]{hclust}}.
#' @method plot env_dissimilarity
#' @export
#' @author Tiago Olivoto \email{tiagoolivoto@@gmail.com}
#' @examples
#'
#' library(metan)
#' mod <- env_dissimilarity(data_ge, ENV, GEN, REP, GY)
#' plot(mod)
plot.env_dissimilarity <- function(x, var = 1, nclust = NULL, ...){
  x <- x[[var]]
  opar <- par(mfrow = c(3, 2),
              mar = c(1, 4, 3, 1))
  on.exit(par(opar))
  hc <- x$SPART_CC %>%
    as.dist() %>%
    hclust()
    plot(hc,
         main = "Single part\n (Cruz and Castoldi, 1991)",
         ylab = "Distance", sub = "", ...)
  if(!missing(nclust)){
    rect.hclust(hc, k = nclust, border = "red")
  }
    hc <- x$CPART_CC %>%
      as.dist() %>%
      hclust()
    plot(hc,
         main = "Complex part\n (Cruz and Castoldi, 1991)",
         ylab = "Distance", sub = "", ...)
    if(!missing(nclust)){
      rect.hclust(hc, k = nclust, border = "red")
    }
    hc <- x$SPART_RO %>%
      as.dist() %>%
      hclust()
    plot(hc,
         main = "Single part\n (Robertson, 1959)",
         ylab = "Distance", sub = "", ...)
    if(!missing(nclust)){
      rect.hclust(hc, k = nclust, border = "red")
    }
    hc <- x$CPART_RO %>%
      as.dist() %>%
      hclust()
    plot(hc,
         main = "Complex part\n (Robertson, 1959)",
         ylab = "Distance", sub = "", ...)
    if(!missing(nclust)){
      rect.hclust(hc, k = nclust, border = "red")
    }
    hc <- x$MSGE %>%
      as.dist() %>%
      hclust()
    plot(hc,
         main = "Mean Square of the Interaction\n genotypes x pairs of environments",
         ylab = "Distance", sub = "", ...)
    if(!missing(nclust)){
      rect.hclust(hc, k = nclust, border = "red")
    }
    hc <- x$correlation %>%
      as.dist() %>%
      hclust()
    plot(hc,
         main = "Pearson's correlation\n genotype's performance",
         ylab = "Distance", sub = "", ...)
    if(!missing(nclust)){
      rect.hclust(hc, k = nclust, border = "red")
    }
}
```
