## Supplementary material for "metan: an R package for multi-environment trial analysis": available at: Fox.R

```
#' Fox's stability function
#'
#' Performs a stability analysis based on the criteria of Fox et al. (1990),
#' using the statistical "TOP third" only. A stratified ranking of the genotypes
#' at each environment is done. The proportion of locations at which the
#' genotype occurred in the top third are expressed in the output.
#'
#'
#' @param .data The dataset containing the columns related to Environments,
#'   Genotypes, replication/block and response variable(s).
#' @param env The name of the column that contains the levels of the
#'   environments.
#' @param gen The name of the column that contains the levels of the genotypes.
#' @param rep The name of the column that contains the levels of the
#'   replications/blocks.
#' @param resp The response variable(s). To analyze multiple variables in a
#'   single procedure use, for example, \code{resp = c(var1, var2, var3)}.
#' @param verbose Logical argument. If \code{verbose = FALSE} the code will run
#'   silently.
#' @return An object of class \code{Fox}, which is a list containing the results
#'   for each variable used in the argument \code{resp}. For each variable, a
#'   tibble with the following columns is returned.
#' * \strong{GEN} the genotype's code.
#' * \strong{mean} the mean for the response variable.
#' * \strong{TOP} The proportion of locations at which the
#' @md
#' @author Tiago Olivoto \email{tiagoolivoto@@gmail.com}
#' @references Fox, P.N., B. Skovmand, B.K. Thompson, H.J. Braun, and R.
#'   Cormier. 1990. Yield and adaptation of hexaploid spring triticale.
#'   Euphytica 47:57-64.
#'   \href{https://link.springer.com/article/10.1007/BF00040364}{doi:10.1007/BF00040364}.
#'
#' @export
#' @examples
#'
#' library(metan)
#' out = Fox(data_ge2,
#'           env = ENV,
#'           gen = GEN,
#'           rep = REP,
#'           resp = PH)
#'
Fox <- function(.data, env, gen, rep, resp, verbose = TRUE) {
  factors  <- .data %>%
    select(ENV = {{env}},
           GEN = {{gen}},
           REP = {{rep}}) %>%
    mutate_all(as.factor)
  vars <- .data %>%
    select({{resp}}) %>%
    select_numeric_cols()
  listres <- list()
  nvar <- ncol(vars)
  for (var in 1:nvar) {
    data <- factors %>%
      mutate(Y = vars[[var]])
    temp <- data %>%
      group_by(ENV) %>%
      mutate(grank = rank(-Y)) %>%
      group_by(GEN) %>%
      summarise(Y = mean(Y),
                TOP = sum(grank <= 3)) %>%
      as_tibble()
    if (nvar > 1) {
      listres[[paste(names(vars[var]))]] <- temp
      if (verbose == TRUE) {
        cat("Evaluating variable", paste(names(vars[var])),
            round((var - 1)/(length(vars) - 1) * 100, 1), "%", "\n")
      }
    } else {
      listres[[paste(names(vars[var]))]] <- temp
    }
  }
  return(structure(listres, class = "Fox"))
}
```
