## Supplementary material for "metan: an R package for multi-environment trial analysis": available at: gamem.R

```
#' Genotype analysis by mixed-effect models
#'
#' Analysis of genotypes in single experiments using mixed-effect models with
#' estimation of genetic parameters.
#'
#'
#' @param .data The dataset containing the columns related to, Genotypes,
#' replication/block and response variable(s).
#' @param gen The name of the column that contains the levels of the genotypes, that will
#' be treated as random effect.
#' @param rep The name of the column that contains the levels of the replications
#' (assumed to be fixed).
#' @param resp The response variable(s). To analyze multiple variables in a
#' single procedure a vector of variables may be used. For example \code{resp =
#' c(var1, var2, var3)}. Select helpers are also allowed.
#' @param block Defaults to \code{NULL}. In this case, a randomized complete block design is considered.
#'  If block is informed, then an alpha-lattice design is employed considering block as random to make use
#'  of inter-block information, whereas the complete replicate effect is always taken as fixed,
#'  as no inter-replicate information was to be recovered (Mohring et al., 2015).
#' @param prob The probability for estimating confidence interval for BLUP's
#' prediction.
#' @param verbose Logical argument. If \code{verbose = FALSE} the code are run
#' silently.
#' @references Mohring, J., E. Williams, and H.-P. Piepho. 2015. Inter-block information:
#' to recover or not to recover it? TAG. Theor. Appl. Genet. 128:1541-54.
#'  \href{http://www.ncbi.nlm.nih.gov/pubmed/25972114}{doi:10.1007/s00122-015-2530-0}

#' @return An object of class \code{gamem}, which is a list with the following items for each
#' element (variable):
#'  * \strong{fixed:} Test for fixed effects.
#'
#'  * \strong{random:} Variance components for random effects.
#'
#'  * \strong{LRT:} The Likelihood Ratio Test for the random effects.
#'
#'  * \strong{blupGEN:} The estimated BLUPS for genotypes
#'
#'  * \strong{Details:} A tibble with the following data: \code{Ngen}, the number of genotypes;
#'    \code{OVmean}, the grand mean; \code{Min}, the minimum observed (returning the genotype and replication/block);
#'    \code{Max} the maximum observed, \code{MinGEN} the winner genotype,
#'    \code{MaxGEN}, the loser genotype.
#'
#'  * \strong{ESTIMATES:} A tibble with the values for the genotypic variance, block-within-replicate
#' variance (if an alpha-lattice design is used by informing the block in \code{block}), the residual
#' variance and their respective contribution to the phenotypic variance; broad-sence heritability,
#' heritability on the entry-mean basis, genotypic coefficient of variation residual coefficient of variation
#' and ratio between genotypic and residual coefficient of variation.
#'
#'  * \strong{residuals:} The residuals of the model.
#'
#' @details \code{gamem} analyses data from a one-way genotype testing experiment.
#' By default, a randomized complete block design is used according to the following model:
#' \deqn{Y_{ij} = m + g_i + r_j + e_{ij}}
#' where \eqn{Y_{ij}} is the response variable of the ith genotype in the \emph{j}th block;
#'  \emph{m} is the grand mean (fixed); \eqn{g_i} is the effect of the \emph{i}th genotype
#'  (assumed to be random); \eqn{r_j} is the effect of the \emph{j}th replicate (assumed to be fixed);
#'  and \eqn{e_{ij}} is the random error.
#'
#' When \code{block} is informed, then a resolvable alpha design is implemented, according to the following model:
#'
#' \deqn{Y_{ijk} = m + g_i + r_j + b_{jk} + e_{ijk}}
#' where where \eqn{y_{ijk}} is the response variable of the \emph{i}th genotype in the
#' \emph{k}th block of the \emph{j}th replicate; \emph{m} is the intercept, \eqn{t_i} is
#'  the effect for the \emph{i}th genotype \eqn{r_j} is the effect of the \emph{j}th
#'  replicate, \eqn{b_{jk}} is the effect of the \emph{k}th incomplete block of
#'  the \emph{j}th replicate, and \eqn{e_{ijk}} is the plot error effect
#'  corresponding to \eqn{y_{ijk}}.
#'
#' @md
#' @importFrom dplyr summarise_at
#' @author Tiago Olivoto \email{tiagoolivoto@@gmail.com}
#' @seealso \code{\link{get_model_data}} \code{\link{waasb}}
#' @export
#' @examples
#'\donttest{
#' library(metan)
#'
#' # fitting the model considering an RCBD
#' # Genotype as random effects
#'
#'rcbd <- gamem(data_g,
#'              gen = GEN,
#'              rep = REP,
#'              resp = c(PH, ED, EL, CL, CW, KW, NR, TKW, NKE))
#'
#' # Likelihood ratio test for random effects
#' ## Statistic
#' get_model_data(rcbd, "lrt")
#'
#' ## P-value
#' get_model_data(rcbd, "pval_lrt")
#'
#' # Variance components
#' get_model_data(rcbd, "vcomp")
#'
#' # Genetic parameters
#' get_model_data(rcbd, "genpar")
#'
#' # BLUPs for genotypes
#' get_model_data(rcbd)
#'
#' # fitting the model considering an alpha-lattice design
#' # Genotype and block-within-replicate as random effects
#' # Note that block effect was now informed.
#'
#' alpha <- gamem(data_alpha,
#'                gen = GEN,
#'                rep = REP,
#'                block = BLOCK,
#'                resp = YIELD)
#' # Use the function  get_model_data() to easely extract the model values.
#' get_model_data(alpha, "genpar")
#'}
#'
gamem <- function(.data, gen, rep, resp, block = NULL, prob = 0.05, verbose = TRUE) {
  # RCBD
  if (missing(block) == TRUE) {
    factors  <- .data %>%
      select(GEN = {{gen}},
             REP = {{rep}}) %>%
      mutate_all(as.factor)
    vars <- .data %>%
      select({{resp}}) %>%
      select_numeric_cols()
    listres <- list()
    nvar <- ncol(vars)
    for (var in 1:nvar) {
      data <- factors %>%
        mutate(Y = vars[[var]])
      Ngen <- nlevels(data$GEN)
      Nbloc <- nlevels(data$REP)
      ovmean <- mean(data$Y)
      Complete <- suppressWarnings(suppressMessages(lmerTest::lmer(Y ~ REP + (1 | GEN), data = data)))
      LRT <- lmerTest::ranova(Complete, reduce.terms = FALSE) %>%
        mutate(model = c("Complete", "Genotype")) %>%
        select(model, everything())
      fixed <- anova(Complete)
      random <- lme4::VarCorr(Complete) %>%
        as.data.frame() %>%
        select(1, 4) %>%
        arrange(grp) %>%
        rename(Group = grp, Variance = vcov)
      GV <- as.numeric(random[1, 2])
      RV <- as.numeric(random[2, 2])
      FV <- GV + RV
      h2g <- GV / FV
      h2mg <- GV / (GV + RV / (Nbloc))
      AccuGen <- sqrt(h2mg)
      CVg <- (sqrt(GV) / ovmean) * 100
      CVr <- (sqrt(RV) / ovmean) * 100
      CVratio <- CVg / CVr
      PROB <- ((1 - (1 - prob)) / 2) + (1 - prob)
      t <- qt(PROB, Nbloc)
      Limits <- t * sqrt(((1 - AccuGen) * GV))
      GVper <- (GV / FV) * 100
      RVper <- (RV / FV) * 100
      ESTIMATES <- tibble(
        Parameters = c(
          "Gen_var", "Gen (%)", "Res_var",
          "Res (%)", "Phen_var", "H2", "H2mg",
          "Accuracy", "CVg", "CVr", "CV ratio"
        ),
        Values = c(GV, GVper, RV, RVper, FV, h2g, h2mg, AccuGen, CVg, CVr, CVratio)
      )

      blups <- tibble(
        GEN = rownames(ranef(Complete)[[1]]),
        BLUPg = ranef(Complete)[[1]]$`(Intercept)`,
        Predicted = BLUPg + ovmean,
        LL = Predicted - Limits,
        UL = Predicted + Limits
      )
      min_gen <- data %>%
        group_by(GEN) %>%
        summarise(Y = mean(Y)) %>%
        top_n(1, -Y) %>%
        select(GEN, Y) %>%
        slice(1)
      max_gen <- data %>%
        group_by(GEN) %>%
        summarise(Y = mean(Y)) %>%
        top_n(1, Y) %>%
        select(GEN, Y) %>%
        slice(1)
      max <- data %>%
        top_n(1, Y) %>%
        slice(1)
      min <- data %>%
        top_n(1, -Y) %>%
        slice(1)
      Details <- tibble(
        Parameters = c("Ngen", "OVmean", "Min", "Max", "MinGEN", "MaxGEN"),
        Values = c(
          Ngen,
          round(mean(data$Y), 4),
          paste0(round(min$Y, 4), " (", min$GEN, " in ", min$REP, ")"),
          paste0(round(max$Y, 4), " (", max$GEN, " in ", max$REP, ")"),
          paste0(round(min_gen[1, 2], 4), " (", min_gen$GEN, ")"),
          paste0(round(max_gen[1, 2], 4), " (", max_gen$GEN, ")")
        )
      )
      residuals <- data.frame(fortify.merMod(Complete))
      temp <- structure(list(
        fixed = fixed %>% rownames_to_column("SOURCE") %>% as_tibble(),
        random = as_tibble(random),
        LRT = as_tibble(LRT),
        blupGEN = as_tibble(blups),
        Details = as_tibble(Details),
        ESTIMATES = as_tibble(ESTIMATES),
        residuals = as_tibble(residuals)
      ),
      class = "gamem"
      )
      if (nvar > 1) {
        listres[[paste(names(vars[var]))]] <- temp
        if (verbose == TRUE) {
          cat("Evaluating variable", paste(names(vars[var])),
              round((var - 1)/(length(vars) - 1) * 100, 1), "%", "\n")
        }
      } else {
        listres[[paste(names(vars[var]))]] <- temp
      }
    }
  }
  # ALPHA-LATTICE
  if (missing(block) == FALSE) {
    factors  <- .data %>%
      select(GEN = {{gen}},
             REP = {{rep}},
             BLOCK = {{block}}) %>%
      mutate_all(as.factor)
    vars <- .data %>%
      select({{resp}}) %>%
      select_numeric_cols()
    listres <- list()
    nvar <- ncol(vars)
    for (var in 1:nvar) {
      data <- factors %>%
        mutate(Y = vars[[var]])
      Ngen <- nlevels(data$GEN)
      Nbloc <- nlevels(data$REP)
      ovmean <- mean(data$Y)
      Complete <- suppressWarnings(suppressMessages(lmerTest::lmer(Y ~ (1 | GEN) + REP + (1 | REP:BLOCK), data = data)))
      LRT <- lmerTest::ranova(Complete, reduce.terms = FALSE) %>%
        mutate(model = c("Complete", "Genotype", "rep:block")) %>%
        select(model, everything())
      fixed <- anova(Complete)
      random <- lme4::VarCorr(Complete) %>%
        as.data.frame() %>%
        select(1, 4) %>%
        arrange(grp) %>%
        rename(Group = grp, Variance = vcov)
      GV <- as.numeric(random[1, 2])
      BRV <- as.numeric(random[2, 2])
      RV <- as.numeric(random[3, 2])
      FV <- GV + RV + BRV
      h2g <- GV / FV
      regen <- ranef(Complete, condVar = TRUE)
      vv <- attr(regen$GEN, "postVar")
      vblup <- 2 * mean(vv)
      sg2 <- c(lme4::VarCorr(Complete)[["GEN"]])
      # H^2 measure proposed by Cullis
      h2mg <- 1 - (vblup / 2 / sg2)
      AccuGen <- sqrt(h2mg)
      CVg <- (sqrt(GV) / ovmean) * 100
      CVr <- (sqrt(RV) / ovmean) * 100
      CVratio <- CVg / CVr
      PROB <- ((1 - (1 - prob)) / 2) + (1 - prob)
      t <- qt(PROB, Nbloc)
      Limits <- t * sqrt(((1 - AccuGen) * GV))
      GVper <- (GV / FV) * 100
      BRper <- (BRV / FV) * 100
      RVper <- (RV / FV) * 100
      ESTIMATES <- tibble(
        Parameters = c(
          "Gen_var", "Gen (%)", "rep:block_var", "rep:block (%)", "Res_var",
          "Res (%)", "Phen_var", "H2", "H2mg", "Accuracy", "CVg", "CVr", "CV ratio"
        ),
        Values = c(GV, GVper, BRV, BRper, RV, RVper, FV, h2g, h2mg, AccuGen, CVg, CVr, CVratio)
      )
      blups <- fortify.merMod(Complete) %>%
        group_by(GEN) %>%
        summarise_at(vars(observed = Y, Predicted = .fitted, resid = .resid), funs(mean)) %>%
        mutate(LL = Predicted - Limits,
               UL = Predicted + Limits)

      min_gen <- data %>%
        group_by(GEN) %>%
        summarise(Y = mean(Y)) %>%
        top_n(1, -Y) %>%
        select(GEN, Y) %>%
        slice(1)
      max_gen <- data %>%
        group_by(GEN) %>%
        summarise(Y = mean(Y)) %>%
        top_n(1, Y) %>%
        select(GEN, Y) %>%
        slice(1)
      max <- data %>%
        top_n(1, Y) %>%
        slice(1)
      min <- data %>%
        top_n(1, -Y) %>%
        slice(1)
      Details <- tibble(
        Parameters = c("Ngen", "OVmean", "Min", "Max", "MinGEN", "MaxGEN"),
        Values = c(
          Ngen,
          round(mean(data$Y), 4),
          paste0(round(min$Y, 4), " (", min$GEN, " in ", min$BLOCK, " of ", min$REP, ")"),
          paste0(round(max$Y, 4), " (", max$GEN, " in ", max$BLOCK, " of ", max$REP, ")"),
          paste0(round(min_gen[1, 2], 4), " (", min_gen$GEN, ")"),
          paste0(round(max_gen[1, 2], 4), " (", max_gen$GEN, ")")
        )
      )
      residuals <- as_tibble(fortify.merMod(Complete))
      temp <- structure(list(
        fixed = fixed %>% rownames_to_column("SOURCE") %>% as_tibble(),
        random = as_tibble(random),
        LRT = as_tibble(LRT),
        blupGEN = as_tibble(blups),
        Details = as_tibble(Details),
        ESTIMATES = as_tibble(ESTIMATES),
        residuals = as_tibble(residuals)
      ),
      class = "gamem"
      )
      if (nvar > 1) {
        listres[[paste(names(vars[var]))]] <- temp
        if (verbose == TRUE) {
          cat("Evaluating variable", paste(names(vars[var])),
              round((var - 1)/(length(vars) - 1) * 100, 1), "%", "\n")
        }
      } else {
        listres[[paste(names(vars[var]))]] <- temp
      }
    }
  }
  if (verbose == TRUE) {
    if (length(which(unlist(lapply(listres, function(x) {
      x[["LRT"]] %>%
        dplyr::filter(model == "Genotype") %>%
        pull(`Pr(>Chisq)`)
    })) > prob)) > 0) {
      cat("------------------------------------------------------------\n")
      cat("Variables with nonsignificant genotype effect\n")
      cat(names(which(unlist(lapply(listres, function(x) {
        x[["LRT"]] %>%
          dplyr::filter(model == "Genotype") %>%
          pull(`Pr(>Chisq)`)
      })) > prob)), "\n")
      cat("------------------------------------------------------------\n")
    }
    cat("Done!\n")
  }
  invisible(structure(listres, class = "gamem"))
}
```
