## Supplementary material for "metan: an R package for multi-environment trial analysis": available at: ge_cluster.R

```
#' Cluster genotypes or environments
#'
#' Performs clustering for genotypes or tester environments based on a dissimilarity matrix.
#'

#' @param .data The dataset containing the columns related to Environments, Genotypes
#' and the response variable. It is also possible to use a two-way table with genotypes
#' in lines and environments in columns as input. In this case you must use \code{table = TRUE}.
#' @param env The name of the column that contains the levels of the environments. Defaults to \code{NULL},
#' in case of the input data is a two-way table.
#' @param gen The name of the column that contains the levels of the genotypes. Defaults to \code{NULL},
#' in case of the input data is a two-way table.
#' @param resp The response variable(s). Defaults to \code{NULL}, in case of the input data is a two-way table.
#' @param table Logical values indicating if the input data is a two-way table with genotypes
#' in the rows and environments in the columns. Defaults to \code{FALSE}.
#' @param distmethod The distance measure to be used. This must be one of
#' \code{'euclidean'}, \code{'maximum'}, \code{'manhattan'}, \code{'canberra'},
#' \code{'binary'}, or \code{'minkowski'}.
#' @param clustmethod The agglomeration method to be used. This should be one
#' of \code{'ward.D'} (Default), \code{'ward.D2'}, \code{'single'}, \code{'complete'},
#' \code{'average'} (= UPGMA), \code{'mcquitty'} (= WPGMA), \code{'median'} (=
#' WPGMC) or \code{'centroid'} (= UPGMC).
#' @param scale Should the data be scaled befor computing the distances? Set to
#' TRUE. Let \eqn{Y_{ij}} be the yield of Hybrid \emph{i} in Location \emph{j},
#' \eqn{\bar Y_{.j}} be the mean yield, and \eqn{S_j} be the standard deviation of
#'  Location \emph{j}. The standardized yield (Zij) is computed as (Ouyang et al. 1995):
#'  \eqn{Z_{ij} = (Y_{ij} - Y_{.j}) / S_j}.
#'
#' @param cluster What should be clustered? Defaults to \code{cluster = "env"} (cluster environments).
#'  To cluster the genotypes use \code{cluster = "gen"}.
#' @param nclust The number of clust to be formed. Set to \code{NULL}.
#'
#' @return
#' * \strong{data} The data that was used to compute the distances.
#'
#' * \strong{cutpoint} The cutpoint of the dendrogram according to Mojena (1977).
#'
#' * \strong{distance} The matrix with the distances.
#'
#' * \strong{de} The distances in an object of class \code{dist}.
#'
#' * \strong{hc} The hierarchical clustering.
#'
#' * \strong{cophenetic} The cophenetic correlation coefficient between distance matrix
#' and cophenetic matrix
#'
#' * \strong{Sqt} The total sum of squares.
#'
#' * \strong{tab} A table with the clusters and similarity.
#'
#' * \strong{clusters} The sum of square and the mean of the clusters for each
#' genotype (if \code{cluster = "env"} or environment (if \code{cluster = "gen"}).
#'
#' * \strong{labclust The labels} of genotypes/environments within each cluster.
#' @md
#' @author Tiago Olivoto \email{tiagoolivoto@@gmail.com}
#' @references Mojena, R. 2015. Hierarchical grouping methods and stopping
#'   rules: an evaluation. Comput. J. 20:359-363. doi:10.1093/comjnl/20.4.359
#'
#' @references Ouyang, Z., R.P. Mowers, A. Jensen, S. Wang, and S. Zheng. 1995.
#'  Cluster analysis for genotype x environment interaction with unbalanced data. Crop Sci. 35:1300-1305.
#'  \href{https://dl.sciencesocieties.org/publications/cs/abstracts/35/5/CS0350051300?access=0&view=pdf}{doi:10.2135/cropsci1995.0011183X003500050008x}
#' @export
#' @examples
#'
#' library(metan)
#'
#' d1 = ge_cluster(data_ge, ENV, GEN, GY, nclust = 3)
#' plot(d1, nclust = 3)
#'
ge_cluster <- function(.data, env = NULL, gen = NULL, resp = NULL,
                       table = FALSE, distmethod = "euclidean",
                       clustmethod = "ward.D", scale = TRUE, cluster =  "env", nclust = NULL) {
  if(!cluster %in% c("env", "gen")){
    stop("The argument 'cluster' must use either 'env' or 'gen'.")
  }
  if(table == FALSE & missing(gen) & missing(env) & missing(resp)){
    stop("Invalid input. If the input data is a two-way table then you must set the argument 'table' to TRUE.")
  }
  if (!distmethod %in% c("euclidean", "maximum", "manhattan",
                         "canberra", "binary", "minkowski")) {
    stop("The argument 'distmethod' is incorrect. It should be one of the 'euclidean', 'maximum', 'manhattan', 'canberra', 'binary', or 'minkowski'.")
  }
  if (!clustmethod %in% c("complete", "ward.D", "ward.D2",
                          "single", "average", "mcquitty", "median", "centroid")) {
    stop("The argument 'distmethod' is incorrect. It should be one of the 'ward.D', 'ward.D2', 'single', 'average', 'mcquitty', 'median' or 'centroid'.")
  }
  if (table == FALSE) {
    data <- as.matrix(make_mat(.data, {{gen}}, {{env}}, {{resp}}))
  }
  if (table == TRUE) {
    if(any(sapply(.data, is.numeric) == FALSE)){
      stop("All columns must be numeric. Please check and fix.")
    }
    data <- as.matrix(.data)
  }
  if(scale == TRUE){
    data = sweep(sweep(data, 2, colMeans(data), FUN = "-"), 2, apply(data, 2, sd), FUN = "/")
  }
  if(cluster == "env"){
    data = t(data)
  }
  de <- dist(data, method = distmethod, diag = T, upper = T)
  mat <- as.matrix(de)
  hc <- hclust(de, method = clustmethod)
  d2 <- cophenetic(hc)
  cof <- cor(d2, de)
  k <- 1.25
  pcorte <- mean(hc$height) + k * sd(hc$height)
  if (!missing(nclust)) {
    groups <- cutree(hc, k = nclust)
    Mgroups <- cbind(data, groups)
    distance <- hc$height[(length(hc$height) - nclust):length(hc$height)]
    Sim <- (1 - distance/max(de))
    Passos <- 1:length(Sim)
    Simgroups <- length(Sim):1
    similarity <- Sim * 100
    Tab <- cbind(Passos, Simgroups, round(similarity,
                                          3), round(distance, 2))
    colnames(Tab) <- c("Steps", "Groups", "Similarity",
                       "Distance")
    TabResgroups <- NULL
    MGr <- cbind(data, groups)
    for (i in 1:nclust) {
      NewGroups <- subset(MGr, groups == i)
      GrupCalc <- NewGroups[, 1:(ncol(NewGroups) - 1)]
      Qtd.Elementos <- nrow(NewGroups)
      if (Qtd.Elementos == 1){
        Media <- GrupCalc
        SqG <- 0
        } else {
          Media <- apply(GrupCalc, 2, mean)
          SqG <- sum(sweep(GrupCalc, 2, Media)^2)
        }
      TabResgroups <- rbind(TabResgroups, c(i, Qtd.Elementos, SqG, Media))
    }
    if(cluster == "env"){
    colnames(TabResgroups) <- c("Cluster", "Number of Environments",
                                "Sum_sq", paste(colnames(TabResgroups[, 4:(ncol(TabResgroups))])))
    } else{
      colnames(TabResgroups) <- c("Cluster", "Number of Genotypes",
                                  "Sum_sq", paste(colnames(TabResgroups[, 4:(ncol(TabResgroups))])))
    }
  } else {
    TabResgroups <- NULL
    Tab <- NULL
  }
  Sqt <- sum(sweep(data, 2, apply(data, 2, mean))^2)
  labels = groups %>%
    as.data.frame() %>%
    rownames_to_column("Code") %>%
    rename(Cluster = ".") %>%
    arrange(Cluster)

  return(structure(list(data = data, cutpoint = pcorte,
                        distance = mat, de = de, hc = hc,
                        cophenetic = cof, Sqt = Sqt, tab = as.data.frame(Tab),
                        clusters = as.data.frame(TabResgroups), labclust = labels),
                   class = "ge_cluster"))
}
```
