## Supplementary material for "metan: an R package for multi-environment trial analysis": available at: ge_details.R

```
#' Details for genotype-environment trials
#'
#' Details for genotype-environment trials
#'
#' @param .data The dataset containing the columns related to Environments,
#'   Genotypes, replication/block and response variable(s).
#' @param env The name of the column that contains the levels of the
#'   environments.
#' @param gen The name of the column that contains the levels of the genotypes.
#' @param resp The response variable(s). To analyze multiple variables in a
#'   single procedure a vector of variables may be used. For example \code{resp
#'   = c(var1, var2, var3)}. Select helpers are also allowed.
#'
#' @return A tibble with the following results for each variable:
#' * \code{Mean}: The grand mean.
#' * \code{Std_err}: The standard error of the mean
#' * \code{Desv_pad}: The standard deviation.
#' * \code{CV}: The coefficient of variation.
#' * \code{Min,Max}: The minimum and maximum value, indicating the genotype and environment of occurence.
#' * \code{MinENV, MinGEN}: The environment and genotype with the lower mean.
#' * \code{MaxENV, MaxGEN}: The environment and genotype with the higher mean.
#' @md
#' @export
#' @author Tiago Olivoto \email{tiagoolivoto@@gmail.com}
#' @examples
#' \donttest{
#' library(metan)
#' details <- ge_details(data_ge2, ENV, GEN, everything())
#' print(details)
#' }
ge_details <- function(.data, env, gen, resp){
  factors  <- .data %>%
    select(ENV = {{env}},
           GEN = {{gen}}) %>%
    mutate_all(as.factor)
  vars <- .data %>%
    select({{resp}}) %>%
    select_numeric_cols()
  listres <- list()
  nvar <- ncol(vars)
  for (var in 1:nvar) {
    data <- factors %>%
      mutate(Y = vars[[var]])
    env_data <- means_by(data, {{env}}) %>%
      add_cols(TYPE = "Env") %>%
      rename(CODE = {{env}})
    gen_data <- means_by(data, {{gen}}) %>%
      add_cols(TYPE = "Gen")%>%
      rename(CODE = {{gen}})
    df_bind <-
      rbind(env_data, gen_data) %>%
      column_to_first(TYPE, CODE)
    min_group <-
      df_bind %>%
      group_by(TYPE) %>%
      top_n(1, -Y) %>%
      select(TYPE, CODE, Y) %>%
      slice(1) %>%
      as.data.frame()
    max_group <-
      df_bind %>%
      group_by(TYPE) %>%
      top_n(1, Y) %>%
      select(TYPE, CODE, Y) %>%
      slice(1) %>%
      as.data.frame()
    min <- data %>% top_n(1, -Y) %>% select(ENV, GEN, Y) %>% slice(1)
    max <- data %>% top_n(1, Y) %>% select(ENV, GEN, Y) %>% slice(1)
    desc_st <- desc_stat(data, stats = c("mean, SE.mean, SD.pop, CV"), verbose = FALSE)
    temp <- tibble(Parameters = c("Mean", "Std_err", "Desv_pad", "CV", "Min", "Max", "MinENV", "MaxENV", "MinGEN", "MaxGEN"),
                   Values = c(round(desc_st[1, 2], 2),
                              round(desc_st[2, 2], 2),
                              round(desc_st[3, 2], 2),
                              round(desc_st[4, 2], 2),
                              paste0(round(min[3], 2), " (", min$GEN, " in ", min$ENV,")"),
                              paste0(round(max$Y, 2), " (", max$GEN, " in ", max$ENV,")"),
                              paste0(min_group[1,2], " (", round(min_group[1,3], 2),")"),
                              paste0(max_group[1,2], " (", round(max_group[1,3], 2),")"),
                              paste0(min_group[2,2], " (", round(min_group[2,3], 2), ") "),
                              paste0(max_group[2,2], " (", round(max_group[2,3], 2), ") ")))
    listres[[paste(names(vars[var]))]] <- temp
  }
  bind <-
    do.call(cbind, lapply(listres, function(x) {
    val <- x[["Values"]] %>% as.character()
  })) %>%
    as_tibble() %>%
    mutate(Parameters = listres[[1]][["Parameters"]]) %>%
    select(Parameters, everything())
  return(bind)
}
```
