## Supplementary material for "metan: an R package for multi-environment trial analysis": available at: ge_plot.R

```
#' Graphical analysis of genotype-vs-environment interaction
#'
#' This function produces a line plot for a graphical interpretation of the
#' genotype-vs-environment interaction. By default, environments are in the x
#' axis whereas the genotypes are depicted by different lines. The y axis
#' contains the value of the selected variable. A heatmap can also be created.
#'
#' @param .data The dataset containing the columns related to Environments,
#'   Genotypes, replication/block and response variable(s).
#' @param env The name of the column that contains the levels of the
#'   environments
#' @param gen The name of the column that contains the levels of the genotypes.
#' @param resp The response variable.
#' @param type The type of plot \code{type = 1} for a heatmap or \code{type = 2}
#'   for a line plot.
#' @param plot_theme The graphical theme of the plot. Default is
#'   \code{plot_theme = theme_metan()}. For more details,see
#'   \code{\link[ggplot2]{theme}}.
#' @param colour Logical argument. If \code{FALSE} then the plot will not be
#'   colored.
#' @return An object of class \code{gg, ggplot}.
#' @author Tiago Olivoto \email{tiagoolivoto@@gmail.com}
#' @export
#' @examples
#' library(metan)
#' library(ggplot2)
#' ge_plot(data_ge2, ENV, GEN, PH)
#' ge_plot(data_ge, ENV, GEN, GY, type = 2)
#'
ge_plot <- function(.data,
                    env,
                    gen,
                    resp,
                    type = 1,
                    plot_theme = theme_metan(),
                    colour = TRUE) {
  if(type == 1){
    p <-
      ggplot(.data, aes({{env}}, {{gen}}, fill= {{resp}})) +
      geom_tile()+
      scale_y_discrete(expand = expand_scale(mult = c(0,0)))+
      scale_x_discrete(expand = expand_scale(mult = c(0,0)))+
      scale_fill_viridis_c()+
      guides(fill = guide_colourbar(label = TRUE,
                                    draw.ulim = TRUE,
                                    draw.llim = TRUE,
                                    frame.colour = "black",
                                    ticks = TRUE,
                                    nbin = 10,
                                    label.position = "right",
                                    barwidth = 1.3,
                                    barheight = 10,
                                    direction = 'vertical'))+
      plot_theme %+replace%
      theme(legend.position = "right",
            legend.title = element_text())

  }
  if(type == 2){
  p <- ggplot(.data, aes(x = {{env}}, y = {{resp}}))
  if (colour == TRUE) {
    p <- p +
      stat_summary(aes(colour = {{gen}},
                       group = {{gen}}),
                   fun.y = mean,
                   geom = "line")
  } else {
    p <- p +
      stat_summary(aes(group = {{gen}}),
                   fun.y = mean,
                   geom = "line",
                   colour = "black")
  }
  p <- p + geom_point(stat = "summary",
                      fun.y = mean,
                      size = 3,
                      shape = 18) +
    plot_theme %+replace%
    theme(legend.position = "right")
  }

  return(p)
}
```
