## Supplementary material for "metan: an R package for multi-environment trial analysis": available at: ge_stats.R

```
#' Statistics for genotype-vs-environment interaction
#'
#' Computes \strong{(i)} within-environment analysis of variance, GEI effect,
#' GEI means, and genotype plus GEI effects; \strong{(ii)} parametric statistics
#' including AMMI-based indexes, Annicchiarico's genotypic confidence index
#' (1992), Ecovalence (Wricke, 1965), regression-based stability (Eberhart and
#' Russell., 1966), Shukla's stability variance parameter (1972); and
#' \strong{(iii)} nonparametric statistics including Fox's stability function
#' (Fox et al. 1990), superiority index (Lin and Binns, 1988), Huehn's stability
#' statistics (Huehn, 1979), and Thennarasu (1995) statistics.
#'
#' @param .data The dataset containing the columns related to Environments,
#'   Genotypes, replication/block and response variable(s).
#' @param env The name of the column that contains the levels of the
#'   environments.
#' @param gen The name of the column that contains the levels of the genotypes.
#' @param rep The name of the column that contains the levels of the
#'   replications/blocks.
#' @param resp The response variable(s). To analyze multiple variables in a
#'   single procedure use, for example, \code{resp = c(var1, var2, var3)}.
#' @param verbose Logical argument. If \code{verbose = FALSE} the code will run
#'   silently.
#' @param prob The probability error assumed.
#' @details The function computes the statistics and ranks for the following
#'   stability indexes. \code{"Y"} (Response variable), \code{"CV"} (coefficient
#'   of variation), \code{"Var"} (Genotype's variance), \code{"Shukla"}
#'   (Shukla's variance, calling \code{\link{Shukla}} internally), \code{"Wi_g",
#'   "Wi_f", "Wi_u"} (Annichiarrico's genotypic confidence index for all,
#'   favorable and unfavorable environments, respectively, calling
#'   \code{\link{Annicchiarico}} internally ), \code{"Ecoval"} (Wricke's
#'   ecovalence, \code{\link{ecovalence}} internally), \code{"Sij"} (Deviations
#'   from the joint-regression analysis) and \code{"R2"} (R-squared from the
#'   joint-regression analysis, calling \code{\link{ge_reg}} internally),
#'   \code{"ASV"} (AMMI-stability value), \code{"SIPC"} (sum of the absolute
#'   values of the IPCA scores), \code{"EV"} (Average of the squared eigenvector
#'   values), \code{"ZA"} (Absolute values of the relative contributions of the
#'   IPCAs to the interaction), and \code{"WAAS"} (Weighted Average of Absolute
#'   Scores), by calling \code{\link{AMMI_indexes}} internally; \code{"HMGV"}
#'   (Harmonic mean of the genotypic value), \code{"RPGV"} (Relative performance
#'   of the genotypic values), \code{"HMRPGV"} (Harmonic mean of the relative
#'   performance of the genotypic values), by calling
#'   \code{\link{Resende_indexes}} internally; \code{"Pi_a", "Pi_f", "Pi_u"}
#'   (Superiority indexes for all, favorable and unfavorable environments,
#'   respectively, calling \code{\link{superiority}} internally), \code{"Gai"}
#'   (Geometric adaptability index, calling \code{\link{gai}} internally),
#'   \code{"S1"} (mean of the absolute rank differences of a genotype over the n
#'   environments), \code{"S2"} (variance among the ranks over the k
#'   environments), \code{"S3"} (sum of the absolute deviations), \code{"S6"}
#'   (relative sum of squares of rank for each genotype), by calling
#'   \code{\link{Huehn}} internally; and  \code{"N1", "N2", "N3", "N4"}
#'   (Thennarasu"s statistics, calling \code{\link{Thennarasu}} internally ).
#' @return An object of class \code{ge_stats} which is a list with one data
#'   frame for each variable containing the computed indexes.
#' @author Tiago Olivoto \email{tiagoolivoto@@gmail.com}
#' @references
#'   Annicchiarico, P. 1992. Cultivar adaptation and recommendation from alfalfa
#'   trials in Northern Italy. Journal of Genetic \& Breeding, 46:269-278
#'
#'   Eberhart, S.A., and W.A. Russell. 1966. Stability parameters for
#'   comparing Varieties. Crop Sci. 6:36-40.
#'   \href{https://www.crops.org/publications/cs/abstracts/6/1/CS0060010036}{doi:10.2135/cropsci1966.0011183X000600010011x}
#'
#'   Fox, P.N., B. Skovmand, B.K. Thompson, H.J. Braun, and R.
#'   Cormier. 1990. Yield and adaptation of hexaploid spring triticale.
#'   Euphytica 47:57-64.
#'   \href{https://link.springer.com/article/10.1007/BF00040364}{doi:10.1007/BF00040364}.
#'
#'   Huehn, V.M. 1979. Beitrage zur erfassung der phanotypischen
#'   stabilitat. EDV Med. Biol. 10:112.
#'
#'   Kang, M.S., and H.N. Pham. 1991. Simultaneous Selection for High
#'   Yielding and Stable Crop Genotypes. Agron. J. 83:161.
#'   \href{https://dl.sciencesocieties.org/publications/aj/abstracts/83/1/AJ0830010161}{doi:10.2134/agronj1991.00021962008300010037x}.
#'
#'   Lin, C.S., and M.R. Binns. 1988. A superiority measure of cultivar
#'   performance for cultivar x location data. Can. J. Plant Sci. 68:193-198.
#'   \href{http://pubs.aic.ca/doi/abs/10.4141/cjps88-018}{doi:10.4141/cjps88-018}
#'
#'   Olivoto, T., A.D.C. L{\'{u}}cio, J.A.G. da silva, V.S. Marchioro,
#'   V.Q. de Souza, and E. Jost. 2019a. Mean performance and stability in
#'   multi-environment trials I: Combining features of AMMI and BLUP techniques.
#'   Agron. J. 111:2949-2960.
#'   \href{https://dl.sciencesocieties.org/publications/aj/abstracts/0/0/agronj2019.03.0220?access=0&view=pdf}{doi:10.2134/agronj2019.03.0220}
#'
#'   Shahbazi, E. 2019. Genotype selection and stability analysis for
#'   seed yield of Nigella sativa using parametric and non-parametric
#'   statistics. Sci. Hortic. (Amsterdam). 253:172-179.
#'   \href{https://www.sciencedirect.com/science/article/pii/S0304423819303012}{doi:10.1016/j.scienta.2019.04.047}.
#'
#'   Shukla, G.K. 1972. Some statistical aspects of partitioning
#'   genotype-environmental components of variability. Heredity. 29:238-245.
#'   \href{http://www.nature.com/articles/hdy197287}{doi:10.1038/hdy.1972.87}.
#'
#'   Thennarasu, K. 1995. On certain nonparametric procedures for
#'   studying genotype x environment interactions and yield stability. Ph.D.
#'   thesis. P.J. School, IARI, New Delhi, India.
#'
#'   Wricke, G. 1965. Zur berechnung der okovalenz bei sommerweizen und hafer.
#'   Z. Pflanzenzuchtg 52:127-138.
#'
#'
#' @export
#' @examples
#' \donttest{
#' library(metan)
#'
#' model <- ge_stats(data_ge, ENV, GEN, REP, GY)
#' get_model_data(model, "stats")
#' }
#'
#'
ge_stats = function(.data,
                    env,
                    gen,
                    rep,
                    resp,
                    verbose = TRUE,
                    prob = 0.05){
  factors  <- .data %>%
    select(ENV = {{env}},
           GEN = {{gen}},
           REP = {{rep}}) %>%
    mutate_all(as.factor)
  vars <- .data %>%
    select({{resp}}) %>%
    select_numeric_cols()
  listres <- list()
  nvar <- ncol(vars)
  for (var in 1:nvar) {
    data <- factors %>%
      mutate(mean = vars[[var]])
individual <- data %>% anova_ind(ENV, GEN, REP, mean)
ge_mean = make_mat(data, GEN, ENV, mean)
ge_effect = ge_effects(data, ENV, GEN, REP, mean)[[1]]
gge_effect = ge_effects(data, ENV, GEN, REP, mean, type = "gge")[[1]]
Mean = apply(ge_mean, 1, mean)
Variance = rowSums(apply(ge_mean, 2, function(x) (x - Mean)^2))
CV <- apply(ge_mean, 1, function(x) (sd(x) / mean(x) * 100))
GENSS = (rowSums(ge_mean^2) - (rowSums(ge_mean)^2)/nlevels(data$ENV))*nlevels(data$REP)
mod = anova(lm(mean ~ REP/ENV + ENV * GEN, data = data))
GENMS = GENSS / mod[2, 1]
Fcal =  GENMS / mod[6, 3]
pval = pf(Fcal, mod[2, 1], mod[6, 1], lower.tail = FALSE)
er_mod <- ge_reg(data, ENV, GEN, REP, mean, verbose = FALSE)
ec_mod <- ecovalence(data, ENV, GEN, REP, mean, verbose = FALSE)[[1]]
an_mod <- Annicchiarico(data, ENV, GEN, REP, mean, verbose = FALSE, prob = prob)
shu_mod <- Shukla(data, ENV, GEN, REP, mean, verbose = FALSE)[[1]]
fox_mod <- Fox(data, ENV, GEN, REP, mean, verbose = FALSE)[[1]]
gai_mod <- gai(data, ENV, GEN, REP, mean, verbose = FALSE)[[1]]
hue_mod <- Huehn(data, ENV, GEN, REP, mean, verbose = FALSE)[[1]]
lb_mod <- superiority(data, ENV, GEN, REP, mean, verbose = FALSE)[[1]]
then_mod <- Thennarasu(data, ENV, GEN, REP, mean, verbose = FALSE)[[1]]
ammm_mod <- performs_ammi(data, ENV, GEN, REP, mean, verbose = FALSE)
ammm_mod <- AMMI_indexes(ammm_mod)[[1]]
blup_mod <- waasb(data, ENV, GEN, REP, mean, verbose = FALSE)
blup_mod <- Resende_indexes(blup_mod)[[1]]
temp <- tibble(GEN = an_mod[[1]]$general$GEN,
               Y = Mean,
               Y_R = rank(-Mean),
               CV = CV,
               CV_R = rank(CV),
               Var = Variance,
               Var_R = rank(Variance),
               Shukla = shu_mod$ShuklaVar,
               Shukla_R = shu_mod$rShukaVar,
               Wi_g = an_mod[[1]]$general$Wi,
               Wi_g_R = an_mod[[1]]$general$rank,
               Wi_f = an_mod[[1]]$favorable$Wi,
               Wi_f_R = an_mod[[1]]$favorable$rank,
               Wi_u = an_mod[[1]]$unfavorable$Wi,
               Wi_u_R = an_mod[[1]]$unfavorable$rank,
               Ecoval = ec_mod$Ecoval,
               Ecoval_R = ec_mod$rank,
               bij = er_mod[[1]]$regression$slope,
               Sij = er_mod[[1]]$regression$deviations,
               Sij_R = rank(abs(er_mod[[1]]$regression$deviations)),
               R2 = er_mod[[1]]$regression$R2,
               R2_R = rank(-er_mod[[1]]$regression$R2),
               ASV = ammm_mod$ASV,
               ASV_R = ammm_mod$ASV_R,
               SIPC = ammm_mod$SIPC,
               SIPC_R = ammm_mod$SIPC_R,
               EV = ammm_mod$EV,
               EV_R = ammm_mod$EV_R,
               ZA = ammm_mod$ZA,
               ZA_R = ammm_mod$ZA_R,
               WAAS = ammm_mod$WAAS,
               WAAS_R = ammm_mod$WAAS_R,
               HMGV = blup_mod$HMGV,
               HMGV_R = blup_mod$HMGV_R,
               RPGV = blup_mod$RPGV,
               RPGV_R = blup_mod$RPGV_R,
               HMRPGV = blup_mod$HMRPGV,
               HMRPGV_R = blup_mod$HMRPGV_R,
               Pi_a = lb_mod$index$Pi_a,
               Pi_a_R = lb_mod$index$R_a,
               Pi_f = lb_mod$index$Pi_f,
               Pi_f_R = lb_mod$index$R_f,
               Pi_u = lb_mod$index$Pi_u,
               Pi_u_R = lb_mod$index$R_u,
               Gai = gai_mod$GAI,
               Gai_R = gai_mod$GAI_R,
               S1 = hue_mod$S1,
               S1_R = hue_mod$S1_R,
               S2 = hue_mod$S2,
               S2_R = hue_mod$S2_R,
               S3 = hue_mod$S3,
               S3_R = hue_mod$S3_R,
               S6 = hue_mod$S6,
               S6_R = hue_mod$S6_R,
               N1 = then_mod$N1,
               N1_R = then_mod$N1_R,
               N2 = then_mod$N2,
               N2_R = then_mod$N2_R,
               N3 = then_mod$N3,
               N3_R = then_mod$N3_R,
               N4 = then_mod$N4,
               N4_R = then_mod$N4_R)
if (nvar > 1) {
  listres[[paste(names(vars[var]))]] <- temp
  if (verbose == TRUE) {
    cat("Evaluating variable", paste(names(vars[var])),
        round((var - 1)/(length(vars) - 1) * 100, 1), "%", "\n")
  }
} else {
  listres[[paste(names(vars[var]))]] <- temp
}
  }
  return(structure(listres, class = "ge_stats"))
}
```
