## Supplementary material for "metan: an R package for multi-environment trial analysis": available at: get_model_data.R

```
#' Get data from a model easily
#'
#' Easily get data from some objects generated in the \strong{metan} package
#' such as the WAASB and WAASBY indexes  (Olivoto et al., 2019a, 2019b) BLUPs,
#' variance components, details of AMMI models and AMMI-based stability
#' statistics.
#'
#'
#' @param x An object created with the functions \code{\link{AMMI_indexes}},
#'   \code{\link{ecovalence}},  \code{\link{Fox}}, \code{\link{gai}},
#'   \code{\link{gamem}}, \code{\link{ge_means}}, \code{\link{ge_reg}},
#'   \code{\link{performs_ammi}}, \code{\link{Resende_indexes}},
#'   \code{\link{Shukla}}, \code{\link{superiority}}, \code{\link{waas}} or
#'   \code{\link{waasb}}.
#' @param what What should be captured from the model. See more in section
#'   \strong{Details}.
#' @param type Chose if the statistics must be show by genotype (\code{type =
#'   "GEN"}, default) or environment (\code{type = "ENV"}), when possible.
#' @return A tibble showing the values of the variable chosen in argument
#'   \code{what}.
#' @details
#' Bellow are listed the options allowed in the argument \code{what} depending
#' on the class of the object
#'
#'  \strong{Objects of class \code{AMMI_indexes}:}
#' * \code{"ASV"} AMMI stability value.
#' * \code{"EV"} Averages of the squared eigenvector values.
#' * \code{"SIPC"} Sums of the absolute value of the IPCA scores.
#' * \code{"WAAS"} Weighted average of absolute scores (default).
#' * \code{"ZA"} Absolute value of the relative contribution of IPCAs to the
#' interaction.
#'
#'  \strong{Objects of class \code{Annicchiarico} and \code{Schmildt}:}
#' * \code{"Sem_rp"} The standard error of the relative mean performance (Schmildt).
#' * \code{"Mean_rp"} The relative performance of the mean.
#' * \code{"rank"} The rank for genotypic confidence index.
#' * \code{"Wi"} The genotypic confidence index.
#'
#'  \strong{Objects of class \code{ecovalence}:}
#' * \code{"Ecoval"} Ecovalence value (default).
#' * \code{"Ecov_perc"} Ecovalence in percentage value.
#' * \code{"rank"} Rank for ecovalence.
#'
#'  \strong{Objects of class \code{ge_reg}:}
#' * \code{"deviations"} The deviations from regression.
#' * \code{"RMSE"} The Root Mean Square Error.
#' * \code{"R2"} The r-square of the regression.
#' * \code{"slope"} The sloop of the regression (default).
#'
#'
#'  \strong{Objects of class \code{ge_effects}:}
#' * For objects of class \code{ge_effects} no argument \code{what} is required.
#'
#'  \strong{Objects of class \code{ge_means}:}
#' * \code{"ge_means"} Genotype-environment interaction means (default).
#' * \code{"env_means"} Environment means.
#' * \code{"gen_means"} Genotype means.
#'
#'  \strong{Objects of class \code{Shukla}:}
#' * \code{"rMean"} Rank for the mean.
#' * \code{"ShuklaVar"} Shukla's stablity variance (default).
#' * \code{"rShukaVar"} Rank for Shukla's stablity variance.
#' * \code{"ssiShukaVar"} Simultaneous selection index.
#'
#'  \strong{Objects of class \code{Fox}:}
#' * \code{"TOP"} The proportion of locations at which the genotype occurred in
#' the top third (default).
#'
#'  \strong{Objects of class \code{gai}:}
#' * \code{"GAI"} The geometric adaptability index (default).
#' * \code{"GAI_R"} The rank for the GAI values.
#'
#'  \strong{Objects of class \code{superiority}:}
#' * \code{"Pi_a"} The superiority measure for all environments (default).
#' * \code{"R_a"} The rank for Pi_a.
#' * \code{"Pi_f"} The superiority measure for favorable environments.
#' * \code{"R_f"} The rank for Pi_f.
#' * \code{"Pi_u"} The superiority measure for unfavorable environments.
#' * \code{"R_u"} The rank for Pi_u.
#'
#'  \strong{Objects of class \code{Huehn}:}
#' * \code{"S1"} Mean of the absolute rank differences of a genotype over the n
#' environments (default).
#' * \code{"S2"} variance among the ranks over the k environments.
#' * \code{"S3"} Sum of the absolute deviations.
#' * \code{"S6"} Relative sum of squares of rank for each genotype.
#' * \code{"S1_R"}, \code{"S2_R"}, \code{"S3_R"}, and  \code{"S6_R"}, the ranks
#' for S1, S2, S3, and S6, respectively.
#'
#'  \strong{Objects of class \code{Thennarasu}:}
#' * \code{"N1"} First statistic (default).
#' * \code{"N2"} Second statistic.
#' * \code{"N3"} Third statistic.
#' * \code{"N4"} Fourth statistic.
#' * \code{"N1_R"}, \code{"N2_R"}, \code{"N3_R"}, and \code{"N4_R"}, The ranks
#' for the statistics.
#'
#'
#'  \strong{Objects of class \code{performs_ammi}:}
#' * \code{"PC1", "PC2", ..., "PCn"} The values for the nth interaction
#' principal component axis.
#' * \code{"ipca_ss"} Sum of square for each IPCA.
#' * \code{"ipca_ms"} Mean square for each IPCA.
#' * \code{"ipca_fval"} F value for each IPCA.
#' * \code{"ipca_pval"} P-value for for each IPCA.
#' * \code{"ipca_expl"}  Explained sum of square for each IPCA (default).
#' * \code{"ipca_accum"} Accumulated explained sum of square.
#'
#'
#' \strong{Objects of class \code{waas}, \code{waas_means}, and \code{waasb}:}
#' * \code{"PC1", "PC2", ..., "PCn"} The values for the nth interaction
#' principal component axis.
#' * \code{"WAASB"}  The weighted average of the absolute scores (default for
#' objects of class \code{waas}).
#' * \code{"PctResp"} The rescaled values of the response variable.
#' * \code{"PctWAASB"} The rescaled values of the WAASB.
#' * \code{"wResp"} The weight for the response variable.
#' * \code{"wWAASB"} The weight for the stability.
#' * \code{"OrResp"} The ranking regarding the response variable.
#' * \code{"OrWAASB"} The ranking regarding the WAASB.
#' * \code{"OrPC1"} The ranking regarding the first principal component axix.
#' * \code{"WAASBY"} The superiority index WAASBY.
#' * \code{"OrWAASBY"} The ranking regarding the superiority index.
#'
#'  \strong{Objects of class \code{waasb} or \code{gamem}:}
#' * \code{"blupg"} For genotype's predicted mean.
#' * \code{"blupge"} for genotype-vs-environment's predicted mean (only for
#' objects of class \code{waasb}).
#' * \code{"genpar"} Genetic parameters (default).
#' * \code{"lrt"} The statistic for the likelihood-ratio test for random effects.
#' * \code{"pval_lrt"}  The p-values for the likelihood-ratio test.
#' * \code{"vcomp"} The variance components for random effects.
#'
#'  \strong{Objects of class \code{Res_ind}}
#' * \code{"HMGV"} For harmonic mean of genotypic values.
#' * \code{"RPGV or RPGV_Y"} For relative performance of genotypic values
#' * \code{"HMRPGV"} For harmonic mean of relative performance of genotypic values
#'
#'
#' @md
#' @importFrom dplyr starts_with matches case_when
#' @author Tiago Olivoto \email{tiagoolivoto@@gmail.com}
#' @export
#' @references
#'
#' Annicchiarico, P. 1992. Cultivar adaptation and recommendation from alfalfa
#' trials in Northern Italy. J. Genet. Breed. 46:269-278.
#'
#' Dias, P.C., A. Xavier, M.D.V. de Resende, M.H.P. Barbosa, F.A. Biernaski,
#' R.A. Estopa. 2018. Genetic evaluation of Pinus taeda clones from somatic
#' embryogenesis and their genotype x environment interaction. Crop Breed. Appl.
#' Biotechnol. 18:55-64.
#' \href{http://www.scielo.br/scielo.php?script=sci_arttext&pid=S1984-70332018000100055&lng=en&tlng=en}{doi:10.1590/1984-70332018v18n1a8}
#'
#' Azevedo Peixoto, L. de, P.E. Teodoro, L.A. Silva, E.V. Rodrigues, B.G.
#' Laviola, and L.L. Bhering. 2018. Jatropha half-sib family selection with high
#' adaptability and genotypic stability. PLoS One 13:e0199880.
#' \href{https://dx.plos.org/10.1371/journal.pone.0199880}{doi:10.1371/journal.pone.0199880}
#'
#' Eberhart, S.A., and W.A. Russell. 1966. Stability parameters for comparing
#' Varieties. Crop Sci. 6:36-40.
#' \href{https://www.crops.org/publications/cs/abstracts/6/1/CS0060010036}{doi:10.2135/cropsci1966.0011183X000600010011x}.
#'
#' Fox, P.N., B. Skovmand, B.K. Thompson, H.J. Braun, and R. Cormier. 1990.
#' Yield and adaptation of hexaploid spring triticale. Euphytica 47:57-64.
#' \href{https://link.springer.com/article/10.1007/BF00040364}{doi:10.1007/BF00040364}.
#'
#' Huehn, V.M. 1979. Beitrage zur erfassung der phanotypischen stabilitat. EDV
#' Med. Biol. 10:112.
#'
#' Olivoto, T., A.D.C. L{\'{u}}cio, J.A.G. da silva, V.S. Marchioro, V.Q. de
#' Souza, and E. Jost. 2019a. Mean performance and stability in
#' multi-environment trials I: Combining features of AMMI and BLUP techniques.
#' Agron. J. 111:2949-2960.
#' \href{https://dl.sciencesocieties.org/publications/aj/abstracts/0/0/agronj2019.03.0220}{doi:10.2134/agronj2019.03.0220}
#'
#' Olivoto, T., A.D.C. L{\'{u}}cio, J.A.G. da silva, B.G. Sari, and M.I. Diel.
#' 2019b. Mean performance and stability in multi-environment trials II:
#' Selection based on multiple traits. Agron. J. 111:2961-2969.
#' \href{https://dl.sciencesocieties.org/publications/aj/abstracts/0/0/agronj2019.03.0221}{doi:10.2134/agronj2019.03.0221}
#'
#' Purchase, J.L., H. Hatting, and C.S. van Deventer. 2000.
#' Genotype vs environment interaction of winter wheat (Triticum aestivum L.)
#' in South Africa: II. Stability analysis of yield performance. South African
#' J. Plant Soil 17:101-107.
#' \href{http://doi.org/10.1080/02571862.2000.10634878}{doi:10.1080/02571862.2000.10634878}
#'
#' Resende MDV (2007) Matematica e estatistica na analise de experimentos e no
#' melhoramento genetico. Embrapa Florestas, Colombo
#'
#' Sneller, C.H., L. Kilgore-Norquest, and D. Dombek. 1997. Repeatability of
#' Yield Stability Statistics in Soybean. Crop Sci. 37:383-390.
#' \href{http://doi.org/10.2135/cropsci1997.0011183X003700020013x}{doi:10.2135/cropsci1997.0011183X003700020013x}
#'
#' Shahbazi, E. 2019. Genotype selection and stability analysis for seed yield
#' of Nigella sativa using parametric and non-parametric statistics. Sci.
#' Hortic. (Amsterdam). 253:172-179.
#' \href{https://www.sciencedirect.com/science/article/pii/S0304423819303012}{doi:10.1016/j.scienta.2019.04.047}.
#'
#' Wricke, G. 1965. Zur berechnung der okovalenz bei sommerweizen und hafer. Z.
#' Pflanzenzuchtg 52:127-138.
#'
#' Zali, H., E. Farshadfar, S.H. Sabaghpour, and R. Karimizadeh. 2012.
#' Evaluation of genotype vs environment interaction in chickpea using measures
#' of stability from AMMI model. Ann. Biol. Res. 3:3126-3136.
#' \href{http://eprints.icrisat.ac.in/id/eprint/7173}{http://eprints.icrisat.ac.in/id/eprint/7173}
#'
#'
#' @examples
#' \donttest{
#' library(metan)
#'
#' #################### joint-regression analysis #####################
#' ge_r <- ge_reg(data_ge2, ENV, GEN, REP,
#'                resp = c(PH, EH, CD, CL, ED))
#' get_model_data(ge_r)
#' get_model_data(ge_r, "deviations")
#'
#'
#' #################### AMMI model #####################
#' # Fit an AMMI model for 7 variables.
#' AMMI <- data_ge2 %>%
#'  performs_ammi(ENV, GEN, REP,
#'                resp = c(PH, ED, TKW, NKR, CD, CL, CW))
#'
#' # Sum of squares
#' get_model_data(AMMI, "ipca_ss")
#'
#' # Mean squares
#' get_model_data(AMMI, "ipca_ms")
#'
#' # Examine the significance (p-value) of the IPCAs
#' get_model_data(AMMI, "ipca_pval")
#'
#' # Explained sum of square for each IPCA
#' get_model_data(AMMI)
#'
#' # Accumulated sum of square
#' get_model_data(AMMI, "ipca_accum")
#'
#' ### AMMI-based stability statistics ###
#' # Get the AMMI stability value
#' AMMI %>%
#' AMMI_indexes() %>%
#' get_model_data("ASV")
#'
#'
#' #################### WAASB model #####################
#' # Fitting the WAAS index
#' AMMI <- waas(data_ge2, ENV, GEN, REP,
#'              resp = c(PH, ED, TKW, NKR))
#'
#' # Getting the weighted average of absolute scores
#' get_model_data(AMMI, what = "WAASB")
#'
#' # And the rank for the WAASB index.
#' get_model_data(AMMI, what = "OrWAASB")
#'
#'
#' #################### BLUP model #####################
#' # Fitting a mixed-effect model
#' blup <- waasb(data_ge2, ENV, GEN, REP,
#'               resp = c(PH, ED, TKW, NKR))
#'
#' # Getting p-values for likelihood-ratio test
#' get_model_data(blup, what = "pval_lrt")
#'
#' # Getting the variance components
#' get_model_data(blup, what = "vcomp")
#'
#' # Getting the genetic parameters
#' get_model_data(blup)
#'
#' ### BLUP-based stability indexes ###
#' blup %>%
#' Resende_indexes() %>%
#' get_model_data()
#'
#'
#' #################### Stability indexes #####################
#' stats_ge <- ge_stats(data_ge, ENV, GEN, REP, everything())
#' get_model_data(stats_ge)
#'}
#'
get_model_data <- function(x, what = NULL, type = "GEN") {
  call_f <- match.call()
  if (!class(x) %in% c("waasb", "waas","waas_means", "gamem", "performs_ammi", "Res_ind",
                       "AMMI_indexes", "ecovalence", "ge_reg", "Fox", "Shukla",
                       "superiority", "ge_effects", "gai", "Huehn", "Thennarasu",
                       "ge_stats", "Annicchiarico", "Schmildt", "ge_means")) {
    stop("Invalid class in object ", call_f[["x"]], ". See ?get_model_data for more information.")
  }
  if (!is.null(what) && substr(what, 1, 2) == "PC") {
    npc <- ncol(x[[1]][["model"]] %>%
                  select(starts_with("PC")) %>%
                  select(matches("PC\\d+")))
    npcwhat <- as.numeric(substr(what, 3, nchar(what)))
    if (npcwhat > npc) {
      stop("The number of principal components informed seems to be larger than those in the model informed in 'x'.")
    }
  }
  check <- c(
    "blupg", "blupge", "Y", "WAASB", "PctResp", "PctWAASB", "wRes", "wWAASB", "OrResp", "OrWAASB",
    "OrPC1", "WAASBY", "OrWAASBY", "vcomp", "lrt", "details", "genpar", "pval_lrt")
  check2 <- paste("PC", 1:200, sep = "")
  check3 <- c("blupg", "blupge", "vcomp", "lrt", "genpar", "pval_lrt", "details")
  check4 <- c("Y", "WAASB", "PctResp", "PctWAASB", "wRes", "wWAASB",
              "OrResp", "OrWAASB", "OrPC1", "WAASBY", "OrWAASBY")
  check5 <- c("ipca_ss", "ipca_ms", "ipca_fval", "ipca_pval", "ipca_expl", "ipca_accum")
  check6 <- c("HMGV", "HMGV_R", "RPGV", "RPGV_Y", "RPGV_R", "HMRPGV", "HMRPGV_Y", "HMRPGV_R")
  check7 <- c("ASV", "SIPC", "EV", "ZA", "WAAS", "ASV_R", "SIPC_R", "EV_R", "ZA_R", "WAAS_R", "ASV_SSI", "SIPC_SSI", "EV_SSI", "ZA_SSI", "WAAS_SSI")
  check8 <- c("Ecoval", "Ecov_perc", "rank")
  check9 <- c("slope", "deviations", "RMSE", "R2")
  check10 <- c("TOP")
  check11 <- c("ShuklaVar", "rMean", "rShukaVar", "ssiShukaVar")
  check12 <- c("Pi_a", "R_a", "Pi_f", "R_f", "Pi_u", "R_u")
  check13 <- c("GAI", "GAI_R")
  check14 <- c("S1","S1_R", "S2", "S2_R", "S3", "S3_R", "S6", "S6_R")
  check15 <- c("N1", "N1_R", "N2", "N2_R", "N3", "N3_R", "N4", "N4_R")
  check16 <- c("stats", "ranks")
  check17 <- c("Mean_rp", "Sd_rp", "Wi", "rank")
  check18 <- c("Mean_rp", "Sem_rp", "Wi", "rank")
  check19 <- c("ge_means", "env_means", "gen_means")
  if (!is.null(what) && !what %in% c(check, check2, check5, check6, check7, check8, check9, check10,
                                     check11, check12, check13, check14, check15, check16, check17,
                                     check18, check19)) {
    stop("The argument 'what' is invalid. Please, check the function help (?get_model_data) for more details.")
  }
  if (!is.null(what) && what %in% check3 && !class(x) %in% c("waasb", "gamem")) {
    stop("Invalid argument 'what'. It can only be used with an object of class 'waasb' or 'gamem'. Please, check and fix.")
  }
  if (!type %in% c("GEN", "ENV")) {
    stop("Argument 'type' invalid. It must be either 'GEN' or 'ENV'.")
  }
  if(class(x) == "ge_means"){
    if (is.null(what)){
      what <- "ge_means"
    }
    if (!what %in% c(check19)) {
      stop("Invalid value in 'what' for object of class 'ge_means'. Allowed are ", paste(check19, collapse = ", "), call. = FALSE)
    }
    if(what == "ge_means"){
    bind <- do.call(
      cbind,
      lapply(x, function(x) {
        x[["ge_means_long"]][["Mean"]]
      })) %>%
      as_tibble() %>%
      add_cols(ENV = x[[1]][["ge_means_long"]][["ENV"]],
               GEN = x[[1]][["ge_means_long"]][["GEN"]]) %>%
      column_to_first(ENV, GEN)
    }
    if(what == "env_means"){
      bind <- do.call(
        cbind,
        lapply(x, function(x) {
          x[["env_means"]][["Mean"]]
        })) %>%
        as_tibble() %>%
        add_cols(ENV = x[[1]][["env_means"]][["ENV"]]) %>%
        column_to_first(ENV)
    }
    if(what == "gen_means"){
      bind <- do.call(
        cbind,
        lapply(x, function(x) {
          x[["gen_means"]][["Mean"]]
        })) %>%
        as_tibble() %>%
        add_cols(GEN = x[[1]][["gen_means"]][["GEN"]]) %>%
        column_to_first(GEN)
    }
  }
  if (class(x) == "Annicchiarico") {
    if (is.null(what)){
      what <- "Wi"
    }
    if (!what %in% c(check17)) {
      stop("Invalid value in 'what' for object of class 'Annicchiarico'. Allowed are ", paste(check17, collapse = ", "), call. = FALSE)
    }
    bind <- do.call(
      cbind,
      lapply(x, function(x) {
        x[["general"]][[what]]
      })) %>%
      as_tibble() %>%
      mutate(gen = x[[1]][["general"]][["GEN"]]) %>%
      select(gen, everything())
  }
  if (class(x) == "Schmildt") {
    if (is.null(what)){
      what <- "Wi"
    }
    if (!what %in% c(check18)) {
      stop("Invalid value in 'what' for object of class 'Schmildt'. Allowed are ", paste(check18, collapse = ", "), call. = FALSE)
    }
    bind <- do.call(
      cbind,
      lapply(x, function(x) {
        x[["general"]][[what]]
      })) %>%
      as_tibble() %>%
      mutate(gen = x[[1]][["general"]][["GEN"]]) %>%
      select(gen, everything())
  }
  if (class(x) == "ge_stats") {
    if (is.null(what)){
      what <- "stats"
    }
    if (!what  %in%  check16) {
      stop("Invalid value in 'what' for object of class 'ge_stats'. Allowed are ", paste(check16, collapse = ", "), call. = FALSE)
    }
    bind <- do.call(
      cbind,
      lapply(x, function(x) {
        if(what == "stats"){
          x %>% select(-contains("_R"), -contains("GEN"))
        } else{
          x %>% select(contains("_R"))
        }
      })) %>%
      as_tibble() %>%
      mutate(gen = x[[1]][["GEN"]]) %>%
      pivot_longer(cols = contains(".")) %>%
      separate(name, into = c("var", "stat"), sep = "\\.") %>%
      pivot_wider(values_from = value, names_from = stat) %>%
      select(var, everything()) %>%
      arrange(var)
  }
  if (class(x) == "Thennarasu") {
    if (is.null(what)){
      what <- "N1"
    }
    if (!what %in% c(check15)) {
      stop("Invalid value in 'what' for object of class 'Thennarasu'. Allowed are ", paste(check15, collapse = ", "), call. = FALSE)
    }
    bind <- do.call(
      cbind,
      lapply(x, function(x) {
        x[[what]]
      })) %>%
      as_tibble() %>%
      mutate(gen = x[[1]][["GEN"]]) %>%
      select(gen, everything())
  }
  if (class(x) == "Huehn") {
    if (is.null(what)){
      what <- "S1"
    }
    if (!what %in% c(check14)) {
      stop("Invalid value in 'what' for object of class 'Huehn'. Allowed are ", paste(check14, collapse = ", "), call. = FALSE)
    }
    bind <- do.call(
      cbind,
      lapply(x, function(x) {
        x[[what]]
      })) %>%
      as_tibble() %>%
      mutate(gen = x[[1]][["GEN"]]) %>%
      select(gen, everything())
  }
  if (class(x) == "gai") {
    if (is.null(what)){
      what <- "GAI"
    }
    if (!what %in% check13) {
      stop("Invalid value in 'what' for object of class 'gai'. Allowed are ", paste(check13, collapse = ", "), call. = FALSE)
    }
    bind <- do.call(
      cbind,
      lapply(x, function(x) {
        x[[what]]
      })) %>%
      as_tibble() %>%
      mutate(gen = x[[1]][["GEN"]]) %>%
      select(gen, everything())
  }
  if (class(x) == "ge_effects") {
    bind <- do.call(
      cbind,
      lapply(x, function(x) {
        make_long(x)[[3]]
      })) %>%
      as_tibble()
    factors <- x[[1]] %>%
      make_long() %>%
      select(1:2)
    bind <- cbind(factors, bind)
  }
  if (class(x) == "superiority") {
    if (is.null(what)){
      what <- "Pi_a"
    }
    if (!what %in% c(check12)) {
      stop("Invalid value in 'what' for object of class 'superiority'. Allowed are ", paste(check12, collapse = ", "), call. = FALSE)
    }
    bind <- do.call(
      cbind,
      lapply(x, function(x) {
        x[["index"]][[what]]
      })) %>%
      as_tibble() %>%
      mutate(gen = x[[1]][["index"]][["GEN"]]) %>%
      select(gen, everything())
  }
  if (class(x) == "Shukla") {
    if (is.null(what)){
      what <- "ShuklaVar"
    }
    if (!what %in% c(check11)) {
      stop("Invalid value in 'what' for object of class 'Shukla'. Allowed are ", paste(check11, collapse = ", "), call. = FALSE)
    }
    bind <- do.call(
      cbind,
      lapply(x, function(x) {
        x[[what]]
      })) %>%
      as_tibble() %>%
      mutate(gen = x[[1]][["GEN"]]) %>%
      select(gen, everything())
  }
  if (class(x) == "Fox") {
    if (is.null(what)){
      what <- "TOP"
    }
    if (!what %in% c(check10)) {
      stop("Invalid value in 'what' for object of class 'Fox'. Allowed are ", paste(check10, collapse = ", "), call. = FALSE)
    }
    bind <- do.call(
      cbind,
      lapply(x, function(x) {
        x[[what]]
      })) %>%
      as_tibble() %>%
      mutate(gen = x[[1]][["GEN"]]) %>%
      select(gen, everything())
  }
  if (class(x) == "ge_reg") {
    if (is.null(what)){
      what <- "slope"
    }
    if (!what %in% c(check9)) {
      stop("Invalid value in 'what' for object of class 'ge_reg'. Allowed are ", paste(check9, collapse = ", "), call. = FALSE)
    }
    bind <- do.call(
      cbind,
      lapply(x, function(x) {
        x[["regression"]][[what]]
      })) %>%
      as_tibble() %>%
      mutate(gen = x[[1]][["regression"]][["GEN"]]) %>%
      select(gen, everything())
  }
  if (class(x) == "ecovalence") {
    if (is.null(what)){
      what <- "Ecoval"
    }
    if (!what %in% check8) {
      stop("Invalid value in 'what' for object of class 'ecovalence'. Allowed are ", paste(check8, collapse = ", "), call. = FALSE)
    }
    bind <- do.call(
      cbind,
      lapply(x, function(x) {
        x[[what]]
      })) %>%
      as_tibble() %>%
      mutate(gen = x[[1]][["GEN"]]) %>%
      select(gen, everything())
  }
  if (class(x) == "AMMI_indexes") {
    if (is.null(what)){
      what <- "WAAS"
    }
    if (!what %in% c(check7)) {
      stop("Invalid value in 'what' for object of class 'AMMI_indexes'. Allowed are ", paste(check7, collapse = ", "), call. = FALSE)
    }
    bind <- do.call(
      cbind,
      lapply(x, function(x) {
        x[[what]]
      })) %>%
      as_tibble() %>%
      mutate(gen = x[[1]][["GEN"]]) %>%
      select(gen, everything())
  }
  if (class(x) == "Res_ind") {
    if (is.null(what)){
      what <- "HMRPGV"
    }
    if (!what %in% c(check6)) {
      stop("Invalid value in 'what' for object of class 'Res_ind'. Allowed are ", paste(check6, collapse = ", "), call. = FALSE)
    }
    bind <- do.call(
      cbind,
      lapply(x, function(x) {
        x[[what]]
      })) %>%
      as_tibble() %>%
      mutate(gen = x[[1]][["GEN"]]) %>%
      select(gen, everything())
  }
  if (class(x) == "performs_ammi") {
    if (is.null(what)){
      what <- "ipca_expl"
    }
    if (!what %in% c("Y", check2, check5)) {
      stop("Invalid value in 'what' for object of class 'performs_ammi'.")
    }
    if (what == "Y" | what %in% check2) {
      bind <- do.call(
        cbind,
        lapply(x, function(x) {
          x$model[[what]]
        })) %>%
        as_tibble() %>%
        mutate(gen = x[[1]][["model"]][["Code"]],
               type = x[[1]][["model"]][["type"]]) %>%
        dplyr::filter(type == {{type}}) %>%
        select(-type) %>%
        select(gen, everything())
    }
    if (what  %in% check5) {
      what <- case_when(
        what == "ipca_ss" ~ "Sum Sq",
        what == "ipca_ms" ~ "Mean Sq",
        what == "ipca_fval" ~ "F value",
        what == "ipca_pval" ~ "Pr(>F)",
        what == "ipca_expl" ~ "Percent",
        what == "ipca_accum" ~ "Accumul"
      )
      bind <- do.call(cbind, lapply(x, function(x) {
        val <- x[["PCA"]][[what]]
      })) %>%
        as_tibble() %>%
        mutate(PC = x[[1]][["PCA"]][["PC"]],
               DF = x[[1]][["PCA"]][["Df"]]) %>%
        select(PC, DF, everything())
    }
  }
  if (any(class(x) %in% c("waas", "waas_means"))){
    if (is.null(what)){
      what <- "WAASB"
    }
    if (what %in% c("WAASB", "PctWAASB", "wWAASB", "OrWAASB", "WAASBY", "OrWAASBY")) {
      what <- case_when(
        what == "WAASB" ~ "WAAS",
        what == "PctWAASB" ~ "PctWAAS",
        what == "wWAASB" ~ "wWAAS",
        what == "OrWAASB" ~ "OrWAAS",
        what == "WAASBY" ~ "WAASY",
        what == "OrWAASBY" ~ "OrWAASY"
      )
    }
    if (what %in% c("Y", "WAAS", "PctResp", "PctWAAS", "wRes", "wWAAS",
                    "OrResp", "OrWAAS", "OrPC1", "WAASY", "OrWAASY") | what  %in% check2) {
      bind <- do.call(
        cbind,
        lapply(x, function(x) {
          x$model[[what]]
        })) %>%
        as_tibble() %>%
        mutate(gen = x[[1]][["model"]][["Code"]],
               type = x[[1]][["model"]][["type"]]) %>%
        dplyr::filter(type == {{type}}) %>%
        select(-type) %>%
        select(gen, everything())
    }
    if (what == "details") {
      bind <- do.call(cbind, lapply(x, function(x) {
        val <- x[["Details"]][["Values"]] %>% as.character()
      })) %>%
        as_tibble() %>%
        mutate(Parameters = x[[1]][["Details"]][["Parameters"]]) %>%
        select(Parameters, everything())
    }
  }
  if (class(x)  %in% c("waasb", "gamem")) {
    if (is.null(what)){
      what <- "genpar"
    }
    if(class(x) == "gamem" & !what %in% check3){
      stop("Invalid value in 'what' for an object of class 'gamem'")
    }
    if (class(x) == "waasb" & what %in% check4) {
      bind <- do.call(
        cbind,
        lapply(x, function(x) {
          x$model[[what]]
        })) %>%
        as_tibble() %>%
        mutate(gen = x[[1]][["model"]][["Code"]],
               type = x[[1]][["model"]][["type"]]) %>%
        dplyr::filter(type == {{type}}) %>%
        select(-type) %>%
        select(gen, everything())
    }
    if (what == "vcomp") {
      bind <- do.call(cbind, lapply(x, function(x) {
        val <- x[["random"]][["Variance"]]
      })) %>%
        as_tibble() %>%
        mutate(Group = x[[1]][["random"]][["Group"]]) %>%
        select(Group, everything())
    }
    if (what == "genpar") {
      bind <- do.call(cbind, lapply(x, function(x) {
        val <- x[["ESTIMATES"]][["Values"]]
      })) %>%
        as_tibble() %>%
        mutate(Parameters = x[[1]][["ESTIMATES"]][["Parameters"]]) %>%
        select(Parameters, everything())
    }
    if (what == "details") {
      bind <- do.call(cbind, lapply(x, function(x) {
        val <- x[["Details"]][["Values"]] %>% as.character()
      })) %>%
        as_tibble() %>%
        mutate(Parameters = x[[1]][["Details"]][["Parameters"]]) %>%
        select(Parameters, everything())
    }
    if (what == "pval_lrt") {
      bind <- do.call(cbind, lapply(x, function(x) {
        val <- x[["LRT"]][["Pr(>Chisq)"]]
      })) %>%
        as_tibble() %>%
        mutate(model = x[[1]][["LRT"]][["model"]]) %>%
        select(model, everything())
    }
    if (what == "lrt") {
      bind <- do.call(cbind, lapply(x, function(x) {
        val <- x[["LRT"]][["LRT"]]
      })) %>%
        as_tibble() %>%
        mutate(model = x[[1]][["LRT"]][["model"]]) %>%
        select(model, everything())
    }
    if (what %in% c("blupg", "blupge")) {
      if (what == "blupg") {
        datt <- x[[1]][["blupGEN"]]
        bind <- do.call(cbind, lapply(x, function(x) {
          val <- arrange(x[["blupGEN"]], GEN)$Predicted
        })) %>%
          as_tibble() %>%
          mutate(gen = datt %>% arrange(GEN) %>% pull(GEN)) %>%
          select(gen, everything())
      }
      if (what == "blupge") {
        bind <- do.call(cbind, lapply(x, function(x) {
          val <- x[["BLUPgge"]][["Predicted"]]
        })) %>%
          as_tibble() %>%
          mutate(ENV = x[[1]][["BLUPgge"]][["ENV"]],
                 GEN = x[[1]][["BLUPgge"]][["GEN"]]) %>%
          select(ENV, GEN, everything())
      }
    }
  }
  return(bind)
}
```
