## Supplementary material for "metan: an R package for multi-environment trial analysis": available at: gm_mean.R

```
#' Geometric mean
#'
#' Helper function to compute the geometric mean. The geometric mean is the \emph{n}th
#' root of \emph{n} products.
#' @param x A numeric vector or a data frame.
#' @param ... Variables to compute the geometric mean. If no variable is informed
#'   and \code{x} is a data frame, all the numeric variables will be used.
#' @param na.rm A logical value indicating whether \code{NA} values should be
#'   stripped before computation proceeds.
#' @seealso \code{\link{hm_mean}}
#' @note Not useful if there are elements with values \code{<=0}.
#' @return The geometric mean(s) of \code{x}. If \code{x} is a numeric vector, the
#'   function returns a numeric value. If a data frame is used then a numeric
#'   vector with the geometric mean for each variable is returned.
#' @export
#'
#' @examples
#' num <- c(1:10, 50)
#' gm_mean(num)
#'
#' num_df <- make_mat(data_ge, ENV, GEN, GY)
#' gm_mean(num_df)
gm_mean <- function(x, ..., na.rm = TRUE){
  if(is.null(nrow(x))){
    exp(sum(log(x[x > 0]), na.rm = na.rm) / length(x))
  } else{
    if(missing(...)){
      df <- select_numeric_cols(x)
    } else{
      df <- select(x, ...) %>%
        select_numeric_cols()
    }
    exp(apply(log(df), 2, mean, na.rm = na.rm))
  }
}
```
