## Supplementary material for "metan: an R package for multi-environment trial analysis": available at: pairs_mantel.R

```
#' Mantel test for a set of correlation matrices
#'
#' This function generate a pairwise matrix of plots to compare the similarity
#' of two or more correlation matrices. In the upper diagonal are presented the
#' plots and in the lower diagonal the result of Mantel test based on
#' permutations.
#'
#'
#' @param ... The input matrices. May be an output generated by the function
#'   \code{lpcor} or a coerced list generated by the function \code{as.lpcor}
#' @param type The type of correlation if an obect generated by the function
#'   \code{lpcor} is used. 1 = Linear correlation matrices, or 2 = partial
#'   correlation matrices.
#' @param nrepet The number of permutations. Default is 1000
#' @param names An optional vector of names of the same length of \code{...} .
#' @param prob The error probability for Mantel test.
#' @param diag Logical argument. If \code{TRUE}, the Kernel density is shown in
#'   the diagonal of plot.
#' @param export Logical argument. If \code{TRUE}, then the plot is exported to
#'   the current directory.
#' @param main The title of the plot, set to 'auto'.
#' @param file.type The format of the file if \code{export = TRUE}.  Set to
#'   \code{'pdf'}. Other possible values are \code{*.tiff} using \code{file.type
#'   = 'tiff'}.
#' @param file.name The name of the plot when exported. Set to \code{NULL},
#'   i.e., automatically.
#' @param width The width of the plot, set to \code{8}.
#' @param height The height of the plot, set to \code{7}.
#' @param resolution The resolution of the plot if \code{file.type = 'tiff'} is
#'   used. Set to \code{300} (300 dpi).
#' @param size.point The size of the points in the plot. Set to \code{0.5}.
#' @param shape.point The shape of the point, set to \code{ 19}.
#' @param alpha.point The value for transparency of the points: 1 = full color.
#' @param fill.point The color to fill the points. Valid argument if points are
#'   between 21 and 25.
#' @param col.point The color for the edge of the point, set to \code{black}.
#' @param minsize The size of the letter that will represent the smallest
#'   correlation coefficient.
#' @param maxsize The size of the letter that will represent the largest
#'   correlation coefficient.
#' @param signcol The colour that indicate significant correlations (based on
#'   the \code{prob} value.), set to 'green'.
#' @param alpha The value for transparency of the color informed in
#'   \code{signcol}, when 1 = full color. Set to 0.15.
#' @param diagcol The color in the kernel distribution. Set to 'gray'.
#' @param col.up.panel,col.lw.panel,col.dia.panel The color for the opper, lower
#'   and diagonal pannels. Set to 'gray', 'gray', and 'gray', respectively.
#' @param pan.spacing The space between the pannels. Set to 0.15.
#' @param digits The number of digits to show in the plot.
#' @return An object of class \code{gg, ggmatrix}.
#' @author Tiago Olivoto \email{tiagoolivoto@@gmail.com}
#' @export
#' @examples
#'\donttest{
#' library(metan)
#' library(dplyr)
#' # iris dataset
#' lpc <- iris %>%
#'        split_factors(Species) %>%
#'        lpcor() %>%
#'        pairs_mantel(names = c('setosa', 'versicolor', 'virginica'))
#'
#'
#' # mtcars dataset
#' mt_num <- mtcars %>% select_if(., is.numeric)
#' lpdata <- as.lpcor(cor(mt_num[1:5]),
#'                    cor(mt_num[1:5]),
#'                    cor(mt_num[2:6]),
#'                    cor(mt_num[4:8])) %>%
#'           pairs_mantel()
#'}
pairs_mantel <- function(..., type = 1, nrepet = 1000, names = NULL,
                         prob = 0.05, diag = FALSE, export = FALSE, main = "auto",
                         file.type = "pdf", file.name = NULL, width = 8, height = 7,
                         resolution = 300, size.point = 0.5, shape.point = 19, alpha.point = 1,
                         fill.point = NULL, col.point = "black", minsize = 2, maxsize = 3,
                         signcol = "green", alpha = 0.15, diagcol = "gray", col.up.panel = "gray",
                         col.lw.panel = "gray", col.dia.panel = "gray", pan.spacing = 0.15,
                         digits = 2) {
  class <- list(...)
  if (!type %in% c(1:2)) {
    stop("The argument type must be 1 (linear correlation) or 2 (partial correlation).")
  }
  if (sum(lapply(class, function(x) !class(x) %in% c("lpcor_group",
                                                     "lpcor", "mahala_group", "covcor_design", "group_clustering") ==
                 TRUE) > 0)) {
    stop("The object must be of the class lpcor. Please use 'as.lpcorr' to convert correlation matrices into the correct format.")
  }
  if (class(...) == "lpcor_group") {
    data <- ...[[1]]
    data <- lapply(data, function(x) {
      x[[1]]
    })
  }
  if (class(...) == "group_clustering") {
    data <- lapply(..., function(x) {
      x$distance
    })
  }
  if (!class(...) %in% c("lpcor_group", "group_clustering")) {
    data <- lapply(..., function(x) {
      x
    })
  }
  w <- c(21:25)
  if (is.null(fill.point) == TRUE && any(w == shape.point)) {
    stop(call. = FALSE, "If 'shape.point' is a value between 21 and 25, you must provide a color for fill the shape using the argument 'fill.point.'")
  }
  for (i in 1:length(data)) {
    if (i == 1) {
      Dataset <- data.frame(var = as.vector(t(data[[1]])[lower.tri(data[[1]], diag = FALSE)]))
      if (is.null(names)) {
        names(Dataset)[which(colnames(Dataset) == "var")] <- paste0("Matrix 1")
      } else {
        names(Dataset)[which(colnames(Dataset) == "var")] <- names[1]
      }
    }
    if (i >= 2) {
      Dataset <- mutate(Dataset, var = as.vector(t(data[[i]])[lower.tri(data[[i]], diag = FALSE)]))
      if (is.null(names)) {
        names(Dataset)[which(colnames(Dataset) == "var")] <- paste0("Matrix ", i)
      } else {
        names(Dataset)[which(colnames(Dataset) == "var")] <- names[i]
      }
    }
  }
  dim <- nrow(data[[1]])
  my_custom_cor <- function(data, mapping, color = I("black"),
                            sizeRange = c(minsize, maxsize), ...) {
    # get the x and y data to use the other code
    x <- GGally::eval_data_col(data, mapping$x)
    y <- GGally::eval_data_col(data, mapping$y)
    D <- matrix(nrow = dim, ncol = dim)
    D[lower.tri(D, diag = FALSE)] <- x
    D[upper.tri(D, diag = FALSE)] <- x
    diag(D) <- 0
    D2 <- matrix(nrow = dim, ncol = dim)
    D2[lower.tri(D2, diag = FALSE)] <- y
    D2[upper.tri(D2, diag = FALSE)] <- y
    diag(D2) <- 0
    ct <- ade4::mantel.randtest(as.dist(D), as.dist(D2), nrepet = nrepet)
    sig <- symnum(ct$pvalue,
                  corr = FALSE,
                  na = FALSE,
                  cutpoints = c(0, 0.001, 0.01, 0.05, 1),
                  symbols = c("***", "**", "*", ""))
    r <- unname(ct$obs)
    rt <- format(r, digits = digits)[1]
    cex <- max(sizeRange)
    percent_of_range <- function(percent, range) {
      percent * diff(range) + min(range, na.rm = TRUE)
    }
    GGally::ggally_text(label = as.character(rt),
                        mapping = aes(),
                        xP = 0.5,
                        yP = 0.5,
                        size = I(percent_of_range(cex * abs(r), sizeRange)),
                        color = color, ...) +
      geom_text(aes_string(x = 0.8, y = 0.8),
                label = sig,
                size = I(cex),
                color = color, ...) +
      theme_classic() +
      theme(panel.background = element_rect(color = col.lw.panel),
            axis.line = element_blank(),
            axis.ticks = element_blank(),
            axis.text.y = element_blank(),
            axis.text.x = element_blank())
  }
  my_custom_smooth <- function(data, mapping, ...) {
    x <- GGally::eval_data_col(data, mapping$x)
    y <- GGally::eval_data_col(data, mapping$y)
    D <- matrix(nrow = dim, ncol = dim)
    D[lower.tri(D, diag = FALSE)] <- x
    D[upper.tri(D, diag = FALSE)] <- x
    diag(D) <- 0
    D2 <- matrix(nrow = dim, ncol = dim)
    D2[lower.tri(D2, diag = FALSE)] <- y
    D2[upper.tri(D2, diag = FALSE)] <- y
    diag(D2) <- 0
    ct <- ade4::mantel.randtest(as.dist(D), as.dist(D2), nrepet = nrepet)
    pval <- unname(ct$pvalue)
    p <- ggplot(data = data, mapping = mapping)
    if (is.null(fill.point) == FALSE) {
      p <- p +
        geom_point(color = I(col.point),
                   fill = fill.point,
                   shape = shape.point,
                   size = size.point,
                   alpha = alpha.point)
    } else {
      p <- p +
        geom_point(color = I(col.point),
                   shape = shape.point,
                   size = size.point,
                   alpha = alpha.point)
    }
    p <- p +
      theme_classic() +
      theme(panel.background = element_rect(fill = "white", color = col.up.panel),
            axis.line = element_blank(),
            axis.ticks = element_blank(),
            axis.text.y = element_blank(),
            axis.text.x = element_blank())
    if (pval < prob) {
      p <- p +
        theme(panel.background = element_rect(fill = ggplot2::alpha(signcol, alpha)))
    }
    p
  }
  ggally_mysmooth <- function(data, mapping, ...) {
    ggplot(data = data, mapping = mapping) +
      geom_density(fill = alpha(diagcol, 1)) +
      theme_classic() +
      theme(panel.background = element_rect(fill = alpha("white", 1), color = col.dia.panel))
  }
  if (main == "auto") {
    title <- paste0("Mantel's test with ", nrepet, " resamples")
  } else {
    title <- main
  }
  if (diag == TRUE) {
    diag <- list(continuous = ggally_mysmooth)
  } else {
    diag <- NULL
  }
  p1 <- GGally::ggpairs(Dataset,
                        title = title,
                        diag = diag,
                        lower = list(continuous = my_custom_cor),
                        upper = list(continuous = my_custom_smooth),
                        axisLabels = "none") +
  theme(panel.spacing = grid::unit(pan.spacing, "lines"))
  if (export == FALSE) {
    return(p1)
  } else if (file.type == "pdf") {
    if (is.null(file.name)) {
      pdf("Pairs of Mantel's test.pdf", width = width,
          height = height)
    } else pdf(paste0(file.name, ".pdf"), width = width, height = height)
    print(p1)
    dev.off()
  }
  if (file.type == "tiff") {
    if (is.null(file.name)) {
      tiff(filename = "Pairs of Mantel's test.tiff", width = width,
           height = height, units = "in", compression = "lzw",
           res = resolution)
    } else tiff(filename = paste0(file.name, ".tiff"), width = width,
                height = height, units = "in", compression = "lzw",
                res = resolution)
    print(p1)
    dev.off()
  }
}
```
