## Supplementary material for "metan: an R package for multi-environment trial analysis": available at: path_coeff.R

```
#' Path coefficients with minimal multicollinearity
#'
#' Computes direct and indirect effects in path analysis. An algorithm to select
#' a set of predictors with minimal multicollinearity and high explanatory power
#' is implemented.
#'
#' When \code{brutstep = TRUE}, first, the algorithm will select a set of
#' predictors with minimal multicollinearity. The selection is based on the
#' variance inflation factor (VIF). An iterative process is performed until the
#' maximum VIF observed is less than \code{maxvif}. The variables selected in
#' this iterative process are then used in a series of stepwise-based
#' regressions. The first model is fitted and p-1 predictor variables are
#' retained (p is the number of variables selected in the iterative process. The
#' second model adjusts a regression considering p-2 selected variables, and so
#' on until the last model, which considers only two variables. Three objects
#' are created. \code{Summary}, with the process summary, \code{Models},
#' containing the aforementioned values for all the adjusted models; and
#' \code{Selectedpred}, a vector with the name of the selected variables in the
#' iterative process.
#'
#' @param .data The data. Must be a dataframe or an object of class
#'   \code{split_factors}.
#' @param resp The dependent variable.
#' @param by One variable (factor) to split the data into subsets. The function
#'   is then applied to each subset and returns a list where each element
#'   contains the results for one level of the variable in \code{by}. To split
#'   the data by more than one factor variable, use the function
#'   \code{\link{split_factors}} to pass subsetted data to \code{.data}.
#' @param pred The predictor variables, set to \code{NULL}, i.e., the predictor
#'   variables are all the numeric variables in the data except that in
#'   \code{resp}.
#' @param exclude Logical argument, set to false. If \code{exclude = TRUE}, then
#'   the variables in \code{pred} are deleted from the data, and the analysis
#'   will use as predictor those that remained, except that in \code{resp}.
#' @param correction Set to \code{NULL}. A correction value (k) that will be
#'   added into the diagonal elements of the \bold{X'X} matrix aiming at
#'   reducing the harmful problems of the multicollinearity in path analysis
#'   (Olivoto et al., 2017)
#' @param knumber When \code{correction = NULL}, a plot showing the values of
#'   direct effects in a set of different k values (0-1) is produced.
#'   \code{knumber} is the number of k values used in the range of 0 to 1.
#' @param brutstep Logical argument, set to \code{FALSE}. If true, then an
#'   algorithm will select a subset of variables with minimal multicollinearity
#'   and fit a set of possible models. See the \bold{Details} section for more
#'   information.
#' @param maxvif The maximum value for the Variance Inflation Factor (cut point)
#'   that will be accepted. See the \bold{Details} section for more information.
#' @param missingval How to deal with missing values. For more information,
#'   please see \code{?cor}.
#' @param verbose If \code{verbose = TRUE} then some results are shown in the
#'   console.
#' @return An object of class \code{path_coeff, group_path, or brute_path} with
#'   the following items:
#' * \strong{Corr.x} A correlation matrix between the predictor variables.
#' * \strong{Corr.y} A vector of correlations between each predictor variable with
#' the dependent variable.
#' * \strong{Coefficients} The path coefficients. Direct effects are the diagonal
#' elements, and the indirect effects those in the off-diagonal elements
#' (column)
#' * \strong{Eigen} Eigenvectors and eigenvalues of the \code{Corr.x.}
#' * \strong{VIF} The Variance Inflation Factors.
#' * \strong{plot} A ggplot2-based graphic showing the direct effects in 21
#' different k values.
#' * \strong{Predictors} The predictor variables used in the model.
#' * \strong{CN} The Condition Number, i.e., the ratio between the highest and
#' lowest eigenvalue.
#' * \strong{Det} The matrix determinant of the \code{Corr.x.}.
#' * \strong{R2} The coefficient of determination of the model.
#' * \strong{Residual} The residual effect of the model.
#' * \strong{Response} The response variable.
#' * \strong{weightvar} The order of the predictor variables with the highest weight
#' (highest eigenvector) in the lowest eigenvalue.
#'
#' If \code{.data} is an object of class \code{split_factors} then a list is
#' returned with the above items for each level of the grouping variable in the
#' function \code{split_factors}.
#' @md
#' @author Tiago Olivoto \email{tiagoolivoto@@gmail.com}
#' @references Olivoto, T., V.Q. Souza, M. Nardino, I.R. Carvalho, M. Ferrari,
#'   A.J. Pelegrin, V.J. Szareski, and D. Schmidt. 2017. Multicollinearity in
#'   path analysis: a simple method to reduce its effects. Agron. J.
#'   109:131-142. doi:10.2134/agronj2016.04.0196.
#'   \href{https://dl.sciencesocieties.org/publications/aj/abstracts/109/1/131}{10.2134/agronj2016.04.0196}.
#'
#' @export
#' @examples
#' \donttest{
#' library(metan)
#'
#' # Using KW as the response variable and all other ones as predictors
#' pcoeff <- path_coeff(data_ge2, resp = KW)
#'
#'
#' # Declaring the predictors
#' pcoeff2 <- path_coeff(data_ge2,
#'                      resp = KW,
#'                      pred = c(PH, EH, NKE, TKW))
#'
#'
#' # Selecting variables to be excluded from the analysis
#'pcoeff3 <- path_coeff(data_ge2,
#'                     resp = KW,
#'                     pred = c(NKR, PERK, KW, NKE),
#'                     exclude = TRUE)
#'
#'
#' # Selecting a set of predictors with minimal multicollinearity
#' # Maximum variance Inflation factor of 5
#'pcoeff4 <- path_coeff(data_ge2,
#'                     resp = KW,
#'                     brutstep = TRUE,
#'                     maxvif = 5)
#'
#'
#' # When one analysis should be carried out for each environment
#' # Using the forward-pipe operator %>%
#'pcoeff5 <- path_coeff(data_ge2, resp = KW, by = ENV)
#'}
#'
#'
path_coeff <- function(.data,
                       resp,
                       by = NULL,
                       pred = NULL,
                       exclude = FALSE,
                       correction = NULL,
                       knumber = 50,
                       brutstep = FALSE,
                       maxvif = 10,
                       missingval = "pairwise.complete.obs",
                       verbose = TRUE) {
  if (missing(resp) == TRUE) {
    stop("A response variable (dependent) must be declared.")
  }
  if (!missing(pred) && brutstep == TRUE) {
    stop("Multiple arguments to select the predictors. Set 'pred' to NULL or 'brutstep' to FALSE.")
  }
  kincrement <- 1/knumber
  if (!missing(by)){
    if(length(as.list(substitute(by))[-1L]) != 0){
      stop("Only one grouping variable can be used in the argument 'by'.\nUse 'split_factors()' to pass '.data' grouped by more than one variable.", call. = FALSE)
    }
    .data <- split_factors(.data, {{by}}, verbose = FALSE, keep_factors = TRUE)
  }
  if (any(class(.data) == "split_factors")) {
    dfs <- list()
    for (k in 1:length(.data[[1]])) {
      data <- .data[[1]][[k]] %>%
        select_numeric_cols() %>%
        as.data.frame()
      nam <- names(.data[[1]][k])
      if (brutstep == FALSE) {
        if (missing(pred)) {
          pr <- data %>% dplyr::select(-{{resp}})
        } else {
          if (exclude == TRUE) {
            pr <- data %>% select(-{{pred}}, -{{resp}})
          } else {
            pr <- data %>% select({{pred}}, -{{resp}})
          }
        }
        names <- colnames(pr)
        y <- data %>% select({{resp}})
        nam_resp <- names(y)
        cor.y <- cor(pr, y, use = missingval)
        cor.x <- cor(pr, use = missingval)
        if (is.null(correction) == FALSE) {
          diag(cor.x) <- diag(cor.x) + correction
        } else {
          cor.x <- cor(pr, use = missingval)
        }
        if (is.null(correction) == TRUE) {
          betas <- data.frame(matrix(nrow = knumber,
                                     ncol = length(pr) + 1))
          cc <- 0
          nvar <- length(pr) + 1
          for (i in 1:knumber) {
            cor.x2 <- cor.x
            diag(cor.x2) <- diag(cor.x2) + cc
            betas[i, 1] <- cc
            betas[i, 2:nvar] <- t(solve_svd(cor.x2, cor.y))
            cc <- cc + kincrement
          }
          names(betas) <- paste0(c("K", names(pr)))
          xx <- betas$K
          yy <- colnames(betas)
          fila <- knumber
          col <- length(yy)
          total <- fila * (col - 1)
          x <- character(length = total)
          y <- character(length = total)
          z <- numeric(length = total)
          k <- 0
          for (i in 1:fila) {
            for (j in 2:col) {
              k <- k + 1
              x[k] <- xx[i]
              y[k] <- yy[j]
              z[k] <- betas[i, j]
            }
          }
          x <- as.numeric(x)
          betas <- data.frame(K = x, VAR = y, direct = z)
          p1 <- ggplot2::ggplot(betas, ggplot2::aes(K,
                                                    direct, col = VAR)) + ggplot2::geom_line(size = 1) +
            ggplot2::theme_bw() + ggplot2::theme(axis.ticks.length = unit(0.2,
                                                                          "cm"), axis.text = element_text(size = 12,
                                                                                                          colour = "black"), axis.title = element_text(size = 12,
                                                                                                                                                       colour = "black"), axis.ticks = element_line(colour = "black"),
                                                 legend.position = "bottom", plot.margin = margin(0.1,
                                                                                                  0.1, 0.1, 0.1, "cm"), legend.title = element_blank(),
                                                 panel.border = element_rect(colour = "black",
                                                                             fill = NA, size = 1), panel.grid.major.x = element_blank(),
                                                 panel.grid.major.y = element_blank(), panel.grid.minor.x = element_blank(),
                                                 panel.grid.minor.y = element_blank()) + ggplot2::labs(x = "k values",
                                                                                                       y = "Beta values") + ggplot2::geom_abline(intercept = 0,
                                                                                                                                                 slope = 0) + ggplot2::scale_x_continuous(limits = c(0,
                                                                                                                                                                                                     1), breaks = seq(0, 1, by = 0.1))
        } else {
          p1 <- "No graphic generated due to correction value"
        }
        eigen <- eigen(cor.x)
        Det <- det(cor.x)
        NC <- max(eigen$values)/min(eigen$values)
        Aval <- data.frame(eigen$values)
        names(Aval) <- "Eigenvalues"
        Avet <- data.frame(t(eigen$vectors))
        names(Avet) <- names
        AvAvet <- cbind(Aval, Avet)
        Direct <- solve(cor.x, cor.y)
        n <- ncol(cor.x)
        Coeff <- data.frame(cor.x)
        for (i in 1:n) {
          for (j in 1:n) {
            Coeff[i, j] <- Direct[j] * cor.x[i, j]
          }
        }
        Residual <- 1 - t(Direct) %*% cor.y
        R2 <- t(Direct) %*% cor.y
        VIF <- data.frame(diag(solve_svd(cor.x)))
        if (verbose == TRUE) {
          names(VIF) <- "VIF"
          cat("\n----------------------------------------------------------------------------\n")
          cat("Level", nam, "\n")
          cat("----------------------------------------------------------------------------\n")
          if (NC > 1000) {
            cat(paste0("Severe multicollinearity. \n",
                       "Condition Number = ", round(NC, 3), "\n",
                       "Please, consider using a correction factor, or use 'brutstep = TRUE'. \n"))
          }
          if (NC < 100) {
            cat(paste0("Weak multicollinearity. \n",
                       "Condition Number = ", round(NC, 3), "\n",
                       "You will probably have path coefficients close to being unbiased. \n"))
          }
          if (NC > 100 & NC < 1000) {
            cat(paste0("Moderate multicollinearity! \n",
                       "Condition Number = ", round(NC, 3), "\n",
                       "Please, cautiosely evaluate the VIF and matrix determinant.",
                       "\n"))
          }
        }
        last <- data.frame(weight = t(AvAvet[c(nrow(AvAvet)),
                                             ])[-c(1), ])
        abs <- data.frame(weight = abs(last[, "weight"]))
        rownames(abs) <- rownames(last)
        last <- abs[order(abs[, "weight"], decreasing = T),
                    , drop = FALSE]
        weightvarname <- paste(rownames(last), collapse = " > ")
        temp <- structure(list(Corr.x = as_tibble(cor.x, rownames = NA),
                               Corr.y = as_tibble(cor.y, rownames = NA),
                               Coefficients = as_tibble(t(Coeff), rownames = NA),
                               Eigen = as_tibble(AvAvet),
                               VIF =  rownames_to_column(VIF, "VAR") %>% as_tibble(),
                               plot = p1,
                               Predictors = names(pr),
                               CN = NC,
                               Det = Det,
                               R2 = R2,
                               Residual = Residual,
                               Response = nam_resp,
                               weightvar = weightvarname),
                          class = "path_coeff")
        dfs[[paste(nam)]] <- temp
      }
      if (brutstep == TRUE) {
        yyy <- data %>% dplyr::select({{resp}}) %>% as.data.frame()
        nam_resp <- names(yyy)
        xxx <- data %>% dplyr::select(-{{resp}}) %>% as.data.frame()
        cor.xx <- cor(xxx, use = missingval)
        VIF <- data.frame(VIF = diag(solve_svd(cor.xx))) %>%
          rownames_to_column("VAR") %>%
          arrange(VIF)
        repeat {
          VIF2 <- slice(VIF, -n())
          xxx2 <- data[VIF2$VAR]
          VIF3 <- data.frame(VIF = diag(solve_svd(cor(xxx2, use = missingval))))%>%
            rownames_to_column("VAR") %>%
            arrange(VIF)
          if (max(VIF3$VIF) <= maxvif)
            break
          VIF <- VIF3
        }
        xxx <- data[VIF3$VAR] %>% as.data.frame()
        selectedpred <- VIF3$VAR
        npred <- ncol(xxx) - 1
        statistics <- data.frame(matrix(nrow = npred -
                                          1, ncol = 8))
        ModelEstimates <- list()
        modelcode <- 1
        nproced <- npred - 1
        if (verbose == TRUE) {
          cat("Level", nam, "\n")
          cat("----------------------------------------------------------------------------\n")
          cat(paste("The algorithm has selected a set of ",
                    nrow(VIF3), " predictors with largest VIF = ",
                    round(max(VIF3$VIF), 3), ".", sep = ""), "\n")
          cat("Selected predictors:", paste0(selectedpred),"\n")
          cat(paste("A forward stepwise-based selection procedure will fit",
                    nproced, "models.", "\n"))
          cat("--------------------------------------------------------------------------\n")
        }

        for (i in 1:nproced) {

          FDSel <- FWDselect::selection(x = xxx, y = unlist(yyy),
                                        q = npred, method = "lm", criterion = "aic",
                                        cluster = FALSE)
          predstw <- FDSel$Variable_names
          x <- data[, c(predstw)]
          names <- colnames(x)
          y <- data %>% dplyr::select({{resp}})
          nam_resp <- names(y)
          cor.y <- cor(x, y, use = missingval)
          cor.x <- cor(x, use = missingval)
          if (is.null(correction) == FALSE) {
            diag(cor.x) <- diag(cor.x) + correction
          } else {
            cor.x <- cor(x, use = missingval)
          }
          if (is.null(correction) == T) {
            betas <- data.frame(matrix(nrow = knumber,
                                       ncol = length(predstw) + 1))
            cc <- 0
            nvar <- length(predstw) + 1
            for (i in 1:knumber) {
              cor.x2 <- cor.x
              diag(cor.x2) <- diag(cor.x2) + cc
              betas[i, 1] <- cc
              betas[i, 2:nvar] <- t(solve_svd(cor.x2, cor.y))
              cc <- cc + kincrement
            }
            names(betas) <- paste0(c("K", predstw))
            xx <- betas$K
            yy <- colnames(betas)
            fila <- knumber
            col <- length(yy)
            total <- fila * (col - 1)
            x <- character(length = total)
            y <- character(length = total)
            z <- numeric(length = total)
            k <- 0
            for (i in 1:fila) {
              for (j in 2:col) {
                k <- k + 1
                x[k] <- xx[i]
                y[k] <- yy[j]
                z[k] <- betas[i, j]
              }
            }
            x <- as.numeric(x)
            betas <- data.frame(K = x, VAR = y, direct = z)
            p1 <- ggplot2::ggplot(betas, ggplot2::aes(K,
                                                      direct, col = VAR)) + ggplot2::geom_line(size = 1) +
              ggplot2::theme_bw() + ggplot2::theme(axis.ticks.length = unit(0.2,
                                                                            "cm"), axis.text = element_text(size = 12,
                                                                                                            colour = "black"), axis.title = element_text(size = 12,
                                                                                                                                                         colour = "black"), axis.ticks = element_line(colour = "black"),
                                                   legend.position = "bottom", plot.margin = margin(0.1,
                                                                                                    0.1, 0.1, 0.1, "cm"), legend.title = element_blank(),
                                                   panel.border = element_rect(colour = "black",
                                                                               fill = NA, size = 1), panel.grid.major.x = element_blank(),
                                                   panel.grid.major.y = element_blank(), panel.grid.minor.x = element_blank(),
                                                   panel.grid.minor.y = element_blank()) +
              ggplot2::labs(x = "k values", y = expression(paste(beta,
                                                                 " values"))) + ggplot2::geom_abline(intercept = 0,
                                                                                                     slope = 0) + ggplot2::scale_x_continuous(breaks = seq(0,
                                                                                                                                                           1, by = 0.1))
          } else {
            p1 <- "No graphic generated due to correction value"
          }
          eigen <- eigen(cor.x)
          Det <- det(cor.x)
          NC <- max(eigen$values)/min(eigen$values)
          Aval <- as.data.frame(eigen$values)
          names(Aval) <- "Eigenvalues"
          Avet <- as.data.frame(eigen$vectors)
          names(Avet) <- names
          AvAvet <- cbind(Aval, Avet)
          Direct <- solve(cor.x, cor.y)
          n <- ncol(cor.x)
          Coeff <- data.frame(cor.x)
          for (i in 1:n) {
            for (j in 1:n) {
              Coeff[i, j] <- Direct[j] * cor.x[i, j]
            }
          }
          Residual <- 1 - t(Direct) %*% cor.y
          R2 <- t(Direct) %*% cor.y
          VIF <- data.frame(diag(solve_svd(cor.x)))
          names(VIF) <- "VIF"
          last <- data.frame(weight = t(AvAvet[c(nrow(AvAvet)),
                                               ])[-c(1), ])
          abs <- data.frame(weight = abs(last[, "weight"]))
          rownames(abs) <- rownames(last)
          last <- abs[order(abs[, "weight"], decreasing = T),
                      , drop = FALSE]
          weightvarname <- paste(rownames(last), collapse = " > ")
          cor.y <- data.frame(cor.y)
          Results <- list(Corr.x = data.frame(cor.x),
                          Corr.y = data.frame(cor.y),
                          Coefficients = data.frame(t(Coeff)),
                          Eigen = AvAvet,
                          VIF = VIF,
                          plot = p1,
                          Predictors = names,
                          CN = NC,
                          Det = Det,
                          R2 = R2,
                          Residual = Residual,
                          Response = nam_resp,
                          weightvar = weightvarname)
          ModelEstimates[[paste("Model", modelcode)]] <- Results
          statistics[i, 1] <- paste("Model", modelcode, sep = "")
          statistics[i, 2] <- FDSel$Information_Criterion
          statistics[i, 3] <- npred
          statistics[i, 4] <- NC
          statistics[i, 5] <- Det
          statistics[i, 6] <- R2
          statistics[i, 7] <- Residual
          statistics[i, 8] <- max(VIF)
          if(verbose == TRUE){
          cat(paste("Adjusting the model", modelcode,
                    "with", npred, "predictor variables (",round(modelcode/nproced * 100, 2), "% concluded)",
                    "\n"), sep = "")
            }
          npred <- npred - 1
          modelcode <- modelcode + 1
        }
        statistics <- statistics[-c(1), ]
        names(statistics) <- c("Model", "AIC", "Numpred",
                               "CN", "Determinant", "R2", "Residual", "maxVIF")
        if (verbose == TRUE) {
          cat("Done!\n")
          cat("\n\n")
          cat("--------------------------------------------------------------------------\n")
          cat("Summary of the adjusted models", "\n")
          cat("--------------------------------------------------------------------------\n")
          print(statistics, digits = 3, row.names = FALSE)
          cat("--------------------------------------------------------------------------\n")
        }
        temp <- structure(list(Models = ModelEstimates,
                               Summary = statistics, Selectedpred = selectedpred),
                          class = "brute_path")
        dfs[[paste(nam)]] <- temp
      }
    }
    return(structure(dfs, class = "group_path"))
  } else {
    if (sum(lapply(.data, is.factor) == TRUE) > 0) {
      if (verbose == TRUE) {
        message("The factors ", paste0(collapse = " ",
                                       names(.data[, unlist(lapply(.data, is.factor))])),
                " where excluded to perform the analysis. If you want to perform an analysis for each level of a factor, use the function 'split_factors() before.' ")
      }
    }
    data <- select_numeric_cols(.data)
    nam <- names(.data)
    if (brutstep == FALSE) {
      if (missing(pred)) {
        pr <- data %>% dplyr::select(-{{resp}})
      } else {
        if (exclude == TRUE) {
          pr <- data %>% select(-{{pred}}, -{{resp}})
        } else {
          pr <- data %>% select({{pred}}, -{{resp}})
        }
      }
      names <- colnames(pr)
      y <- data %>% select({{resp}})
      nam_resp <- names(y)
      cor.y <- cor(pr, y, use = missingval)
      cor.x <- cor(pr, use = missingval)
      if (is.null(correction) == FALSE) {
        diag(cor.x) <- diag(cor.x) + correction
      } else {
        cor.x <- cor(pr, use = missingval)
      }
      if (is.null(correction) == TRUE) {
        betas <- data.frame(matrix(nrow = knumber, ncol = length(pr) +
                                     1))
        cc <- 0
        nvar <- length(pr) + 1
        for (i in 1:knumber) {
          cor.x2 <- cor.x
          diag(cor.x2) <- diag(cor.x2) + cc
          betas[i, 1] <- cc
          betas[i, 2:nvar] <- t(solve_svd(cor.x2, cor.y))
          cc <- cc + kincrement
        }
        names(betas) <- paste0(c("K", names(pr)))
        xx <- betas$K
        yy <- colnames(betas)
        fila <- knumber
        col <- length(yy)
        total <- fila * (col - 1)
        x <- character(length = total)
        y <- character(length = total)
        z <- numeric(length = total)
        k <- 0
        for (i in 1:fila) {
          for (j in 2:col) {
            k <- k + 1
            x[k] <- xx[i]
            y[k] <- yy[j]
            z[k] <- betas[i, j]
          }
        }
        x <- as.numeric(x)
        betas <- data.frame(K = x, VAR = y, direct = z)
        p1 <- ggplot2::ggplot(betas, ggplot2::aes(K,
                                                  direct, col = VAR)) + ggplot2::geom_line(size = 1) +
          ggplot2::theme_bw() + ggplot2::theme(axis.ticks.length = unit(0.2,
                                                                        "cm"), axis.text = element_text(size = 12,
                                                                                                        colour = "black"), axis.title = element_text(size = 12,
                                                                                                                                                     colour = "black"), axis.ticks = element_line(colour = "black"),
                                               legend.position = "bottom", plot.margin = margin(0.1,
                                                                                                0.1, 0.1, 0.1, "cm"), legend.title = element_blank(),
                                               panel.border = element_rect(colour = "black",
                                                                           fill = NA, size = 1), panel.grid.major.x = element_blank(),
                                               panel.grid.major.y = element_blank(), panel.grid.minor.x = element_blank(),
                                               panel.grid.minor.y = element_blank()) + ggplot2::labs(x = "k values",
                                                                                                     y = "Beta values") + ggplot2::geom_abline(intercept = 0,
                                                                                                                                               slope = 0) + ggplot2::scale_x_continuous(breaks = seq(0,
                                                                                                                                                                                                     1, by = 0.1))
      } else {
        p1 <- "No graphic generated due to correction value"
      }
      eigen <- eigen(cor.x)
      Det <- det(cor.x)
      NC <- max(eigen$values)/min(eigen$values)
      Aval <- data.frame(eigen$values)
      names(Aval) <- "Eigenvalues"
      Avet <- data.frame(t(eigen$vectors))
      names(Avet) <- names
      AvAvet <- cbind(Aval, Avet)
      Direct <- solve(cor.x, cor.y)
      n <- ncol(cor.x)
      Coeff <- data.frame(cor.x)
      for (i in 1:n) {
        for (j in 1:n) {
          Coeff[i, j] <- Direct[j] * cor.x[i, j]
        }
      }
      Residual <- 1 - t(Direct) %*% cor.y
      R2 <- t(Direct) %*% cor.y
      VIF <- data.frame(diag(solve_svd(cor.x)))
      names(VIF) <- "VIF"
      if (verbose == TRUE) {
        if (NC > 1000) {
          cat(paste0("Severe multicollinearity. \n",
                     "Condition Number = ", round(NC, 3), "\n",
                     "Please, consider using a correction factor, or use 'brutstep = TRUE'. \n"))
        }
        if (NC < 100) {
          cat(paste0("Weak multicollinearity. \n", "Condition Number = ",
                     round(NC, 3), "\n", "You will probably have path coefficients close to being unbiased. \n"))
        }
        if (NC > 100 & NC < 1000) {
          cat(paste0("Moderate multicollinearity! \n",
                     "Condition Number = ", round(NC, 3), "\n",
                     "Please, cautiosely evaluate the VIF and matrix determinant.",
                     "\n"))
        }
      }
      last <- data.frame(weight = t(AvAvet[c(nrow(AvAvet)),
                                           ])[-c(1), ])
      abs <- data.frame(weight = abs(last[, "weight"]))
      rownames(abs) <- rownames(last)
      last <- abs[order(abs[, "weight"], decreasing = T),
                  , drop = FALSE]
      weightvarname <- paste(rownames(last), collapse = " > ")
      temp <- structure(list(Corr.x = as_tibble(cor.x, rownames = NA),
                             Corr.y = as_tibble(cor.y, rownames = NA),
                             Coefficients = as_tibble(t(Coeff), rownames = NA),
                             Eigen = as_tibble(AvAvet),
                             VIF =  rownames_to_column(VIF, "VAR") %>% as_tibble(),
                             plot = p1,
                             Predictors = names(pr),
                             CN = NC,
                             Det = Det,
                             R2 = R2,
                             Residual = Residual,
                             Response = nam_resp,
                             weightvar = weightvarname),
                        class = "path_coeff")
      return(temp)
    }
    if (brutstep == TRUE) {
      yyy <- data %>% dplyr::select({{resp}}) %>% as.data.frame()
      nam_resp <- names(yyy)
      xxx <- data %>% dplyr::select(-{{resp}}) %>% as.data.frame()
      cor.xx <- cor(xxx, use = missingval)
      VIF <- data.frame(VIF = diag(solve_svd(cor.xx))) %>%
        rownames_to_column("VAR") %>%
        arrange(VIF)
      repeat {
        VIF2 <- slice(VIF, -n())
        xxx2 <- data[VIF2$VAR]
        VIF3 <- data.frame(VIF = diag(solve_svd(cor(xxx2, use = missingval))))%>%
          rownames_to_column("VAR") %>%
          arrange(VIF)
        if (max(VIF3$VIF) <= maxvif)
          break
        VIF <- VIF3
      }
      xxx <- data[VIF3$VAR] %>% as.data.frame()
      selectedpred <- VIF3$VAR
      npred <- ncol(xxx) - 1
      statistics <- data.frame(matrix(nrow = npred - 1,
                                      ncol = 8))
      ModelEstimates <- list()
      modelcode <- 1
      nproced <- npred - 1
      if (verbose == TRUE) {
        cat("--------------------------------------------------------------------------\n")
        cat(paste("The algorithm has selected a set of ",
                  nrow(VIF3), " predictors with largest VIF = ",
                  round(max(VIF3$VIF), 3), ".", sep = ""), "\n")
        cat("Selected predictors:", paste0(selectedpred),"\n")
        cat(paste("A forward stepwise-based selection procedure will fit", nproced, "models.\n"))
        cat("--------------------------------------------------------------------------\n")
      }
      for (i in 1:nproced) {
        FDSel <- FWDselect::selection(x = xxx, y = unlist(yyy),
                                      q = npred, method = "lm", criterion = "aic",
                                      cluster = FALSE)
        predstw <- FDSel$Variable_names
        x <- data[, c(predstw)]
        names <- colnames(x)
        y <- data %>% dplyr::select({{resp}})
        nam_resp <- names(y)
        cor.y <- cor(x, y, use = missingval)
        cor.x <- cor(x, use = missingval)
        if (is.null(correction) == FALSE) {
          diag(cor.x) <- diag(cor.x) + correction
        } else {
          cor.x <- cor(x, use = missingval)
        }
        if (is.null(correction) == T) {
          betas <- data.frame(matrix(nrow = knumber,
                                     ncol = length(predstw) + 1))
          cc <- 0
          nvar <- length(predstw) + 1
          for (i in 1:knumber) {
            cor.x2 <- cor.x
            diag(cor.x2) <- diag(cor.x2) + cc
            betas[i, 1] <- cc
            betas[i, 2:nvar] <- t(solve_svd(cor.x2, cor.y))
            cc <- cc + kincrement
          }
          names(betas) <- paste0(c("K", predstw))
          xx <- betas$K
          yy <- colnames(betas)
          fila <- knumber
          col <- length(yy)
          total <- fila * (col - 1)
          x <- character(length = total)
          y <- character(length = total)
          z <- numeric(length = total)
          k <- 0
          for (i in 1:fila) {
            for (j in 2:col) {
              k <- k + 1
              x[k] <- xx[i]
              y[k] <- yy[j]
              z[k] <- betas[i, j]
            }
          }
          x <- as.numeric(x)
          betas <- data.frame(K = x, VAR = y, direct = z)
          p1 <- ggplot2::ggplot(betas, ggplot2::aes(K,
                                                    direct, col = VAR)) + ggplot2::geom_line(size = 1) +
            ggplot2::theme_bw() + ggplot2::theme(axis.ticks.length = unit(0.2,
                                                                          "cm"), axis.text = element_text(size = 12,
                                                                                                          colour = "black"), axis.title = element_text(size = 12,
                                                                                                                                                       colour = "black"), axis.ticks = element_line(colour = "black"),
                                                 legend.position = "bottom", plot.margin = margin(0.1,
                                                                                                  0.1, 0.1, 0.1, "cm"), legend.title = element_blank(),
                                                 panel.border = element_rect(colour = "black",
                                                                             fill = NA, size = 1), panel.grid.major.x = element_blank(),
                                                 panel.grid.major.y = element_blank(), panel.grid.minor.x = element_blank(),
                                                 panel.grid.minor.y = element_blank()) + ggplot2::labs(x = "k values",
                                                                                                       y = expression(paste(beta, " values"))) +
            ggplot2::geom_abline(intercept = 0, slope = 0) +
            ggplot2::scale_x_continuous(breaks = seq(0,
                                                     1, by = 0.1))
        } else {
          p1 <- "No graphic generated due to correction value"
        }
        eigen <- eigen(cor.x)
        Det <- det(cor.x)
        NC <- max(eigen$values)/min(eigen$values)
        Aval <- as.data.frame(eigen$values)
        names(Aval) <- "Eigenvalues"
        Avet <- as.data.frame(eigen$vectors)
        names(Avet) <- names
        AvAvet <- cbind(Aval, Avet)
        Direct <- solve(cor.x, cor.y)
        n <- ncol(cor.x)
        Coeff <- data.frame(cor.x)
        for (i in 1:n) {
          for (j in 1:n) {
            Coeff[i, j] <- Direct[j] * cor.x[i, j]
          }
        }
        Residual <- 1 - t(Direct) %*% cor.y
        R2 <- t(Direct) %*% cor.y
        VIF <- data.frame(diag(solve_svd(cor.x)))
        names(VIF) <- "VIF"
        last <- data.frame(weight = t(AvAvet[c(nrow(AvAvet)),
                                             ])[-c(1), ])
        abs <- data.frame(weight = abs(last[, "weight"]))
        rownames(abs) <- rownames(last)
        last <- abs[order(abs[, "weight"], decreasing = T),
                    , drop = FALSE]
        weightvarname <- paste(rownames(last), collapse = " > ")
        cor.y <- data.frame(cor.y)
        Results <- list(Corr.x = data.frame(cor.x),
                        Corr.y = data.frame(cor.y),
                        Coefficients = data.frame(t(Coeff)),
                        Eigen = AvAvet,
                        VIF = VIF,
                        plot = p1,
                        Predictors = predstw,
                        CN = NC,
                        Det = Det,
                        R2 = R2,
                        Residual = Residual,
                        Response = nam_resp,
                        weightvar = weightvarname)
        ModelEstimates[[paste("Model", modelcode)]] <- Results
        statistics[i, 1] <- paste("Model", modelcode, sep = "")
        statistics[i, 2] <- FDSel$Information_Criterion
        statistics[i, 3] <- npred
        statistics[i, 4] <- NC
        statistics[i, 5] <- Det
        statistics[i, 6] <- R2
        statistics[i, 7] <- Residual
        statistics[i, 8] <- max(VIF)
        if (verbose == TRUE) {
          cat(paste("Adjusting the model ", modelcode,
                    " with ", npred, " predictors (", round(modelcode/nproced *
                                                              100, 2), "% concluded)", "\n", sep = ""))
        }
        npred <- npred - 1
        modelcode <- modelcode + 1
      }
      statistics <- statistics[-c(1), ]
      names(statistics) <- c("Model", "AIC", "Numpred",
                             "CN", "Determinant", "R2", "Residual", "maxVIF")
      if (verbose == TRUE) {
        cat("Done!\n")
        cat("--------------------------------------------------------------------------\n")
        cat("Summary of the adjusted models", "\n")
        cat("--------------------------------------------------------------------------\n")
        print(statistics, digits = 3, row.names = FALSE)
        cat("--------------------------------------------------------------------------\n")
      }
      temp <- list(Models = ModelEstimates, Summary = statistics,
                   Selectedpred = selectedpred)
      return(structure(temp, class = "brute_path"))
    }
  }
}
```
