## Supplementary material for "metan: an R package for multi-environment trial analysis": available at: performs_ammi.R

```
#' Additive Main effects and Multiplicative Interaction
#'
#' Compute the Additive Main effects and Multiplicative interaction. This
#' function also serves as a helper function for other procedures performed in
#' the \pkg{metan} package such as \code{\link{waas}} and \code{\link{wsmp}}
#'
#' @param .data The dataset containing the columns related to Environments,
#'   Genotypes, replication/block and response variable(s).
#' @param env The name of the column that contains the levels of the
#'   environments
#' @param gen The name of the column that contains the levels of the genotypes
#' @param rep The name of the column that contains the levels of the
#'   replications/blocks
#' @param resp The response variable(s). To analyze multiple variables in a
#'   single procedure a vector of variables may be used. For example \code{resp
#'   = c(var1, var2, var3)}.
#' @param verbose Logical argument. If \code{verbose = FALSE} the code will run
#'   silently.
#' @return
#' * \strong{ANOVA} The analysis of variance for the AMMI model.
#'
#' * \strong{PCA} The principal component analysis
#'
#' * \strong{MeansGxE} The means of genotypes in the environments
#'
#' * \strong{model} scores for genotypes and environments in all the possible
#' axes.
#' @md
#' @author Tiago Olivoto \email{tiagoolivoto@@gmail.com}
#' @export
#' @examples
#'\donttest{
#' library(metan)
#' ammi_model = performs_ammi(data_ge, ENV, GEN, REP,
#'                            resp = c(GY, HM))
#'
#' # GY x PC1 (variable GY)
#' plot_scores(ammi_model,
#'             col.env = 'olivedrab',
#'             col.gen = 'orange2',
#'             x.lab = 'My own x label')
#'
#' # PC1 x PC2 (variable HM)
#' plot_scores(ammi_model,
#'             type = 2)
#'
#' # PC1 x PC2 (variable HM)
#' # Draw a convex hull polygon
#' plot_scores(ammi_model,
#'             type = 2,
#'             polygon = TRUE)
#'
#'}
performs_ammi <- function(.data, env, gen, rep, resp, verbose = TRUE) {
    factors  <- .data %>%
        select(ENV = {{env}},
               GEN = {{gen}},
               REP = {{rep}}) %>%
        mutate_all(as.factor)
    vars <- .data %>%
        select({{resp}}) %>%
        select_numeric_cols()
    listres <- list()
    nvar <- ncol(vars)
    for (var in 1:nvar) {
        data <- factors %>%
            mutate(mean = vars[[var]])
        nenv <- nlevels(data$ENV)
        ngen <- nlevels(data$GEN)
        nrep <- nlevels(data$REP)
        minimo <- min(ngen, nenv) - 1
        model <- aov(mean ~ ENV + REP %in% ENV + GEN + ENV:GEN, data = data)
        df <- fortify(model)
        datares <- model$model
        datares$factors <- paste(datares$ENV, datares$GEN)
        residuals <- cbind(datares, df %>%
                               select(fitted = .fitted,
                                      resid = .resid,
                                      stdres = .stdresid))
        if (minimo < 2) {
            stop("The analysis AMMI is not possible. Both genotypes and environments must have more than two levels.")
        }
        mm <- anova(model)
        nn <- mm[2, ]
        mm[2, ] <- mm[3, ]
        mm[3, ] <- nn
        row.names(mm)[2] <- "REP(ENV)"
        row.names(mm)[3] <- "GEN     "
        mm[1, 4] <- mm[1, 3]/mm[2, 3]
        mm[1, 5] <- 1 - pf(mm[1, 4], mm[1, 1], mm[2, 1])
        anova <- mm
        probint <- anova[4, 5]
        DFE <- df.residual(model)
        MSE <- deviance(model)/DFE
        MEANS <- data %>%
            group_by(ENV, GEN) %>%
            summarise(Y = mean(mean)) %>%
            ungroup()
        residual <- residuals(lm(Y ~ ENV + GEN, data = MEANS))
        MEANS %<>% mutate(RESIDUAL = residual)
        s <- svd(t(matrix(residual, nenv, byrow = T)))
        U <- s$u[, 1:minimo]
        LL <- diag(s$d[1:minimo])
        V <- s$v[, 1:minimo]
        SS <- (s$d[1:minimo]^2) * nrep
        SUMA <- sum(SS)
        percent <- (1/SUMA) * SS * 100
        DFAMMI <- replicate(minimo, 0)
        acum <- DFAMMI
        MSAMMI <- DFAMMI
        F.AMMI <- DFAMMI
        PROBF <- DFAMMI
        acumula <- 0
        for (i in 1:(minimo)) {
            DF <- (ngen - 1) + (nenv - 1) - (2 * i - 1)
            if (DF <= 0)
                break
            DFAMMI[i] <- DF
            acumula <- acumula + percent[i]
            acum[i] <- acum[i] + acumula
            MSAMMI[i] <- SS[i]/DFAMMI[i]
            if (MSE > 0)
                F.AMMI[i] <- round(MSAMMI[i]/MSE, 2) else F.AMMI[i] <- NA
            if (DFE > 0)
                PROBF[i] <- round(1 - pf(F.AMMI[i], DFAMMI[i], DFE),
                                  4) else PROBF[i] <- NA
        }
        percent <- round(percent, 1)
        acum <- round(acum, 1)
        SS <- round(SS, 5)
        MSAMMI <- round(MSAMMI, 5)
        SSAMMI <- data.frame(percent, acum,
                             Df = DFAMMI,
                             `Sum Sq` = SS,
                             `Mean Sq` = MSAMMI,
                             `F value` = F.AMMI,
                             Pr.F = PROBF)
        nssammi <- nrow(SSAMMI)
        SSAMMI <- SSAMMI[SSAMMI$Df > 0, ]
        nss <- nrow(SSAMMI)
        row.names(SSAMMI) <- paste("PC", 1:nss, sep = "")
        SCOREG <- U %*% LL^0.5
        SCOREE <- V %*% LL^0.5
        colnames(SCOREG) <- colnames(SCOREE) <- paste("PC", 1:minimo, sep = "")
        bplot <- MEANS %>%
            group_by(GEN) %>%
            summarise(Y = mean(Y)) %>%
            mutate(type = "GEN") %>%
            rename(Code = GEN) %>%
            cbind(., SCOREG) %>%
            rbind(., MEANS %>%
                      group_by(ENV) %>%
                      summarise(Y = mean(Y)) %>%
                      mutate(type = "ENV") %>%
                      rename(Code = ENV) %>%
                      cbind(., SCOREE)) %>%
            select(type, Code, everything())
        PC <- SSAMMI %>% select(-percent, -acum, everything())
        resid <- as.data.frame(anova[nrow(anova), ])
        rownames(resid) <- "Residuals"
        sum <- as.data.frame(anova[nrow(anova), ])
        sum$Df <- sum(anova$Df)
        sum$`Sum Sq` <- sum(anova$`Sum Sq`)
        sum$`Mean Sq` <- sum$`Sum Sq`/sum$Df
        rownames(sum) <- "Total"
        ERRO <- rbind(resid, sum)
        names(PC) <- paste(c("Df", "Sum Sq", "Mean Sq", "F value",
                             "Pr(>F)", "Percent", "Accumul"))
        anova <- rbind_fill(mm[-nrow(mm), ], PC, ERRO) %>%
            rownames_to_column("Source") %>%
            as_tibble()
        MeansGxE <- MEANS[, 1:3]
        EscGEN <- subset(bplot, type == "GEN")
        names(EscGEN)[2] <- "GEN"
        names(EscGEN)[3] <- "y"
        EscENV <- subset(bplot, type == "ENV")
        names(EscENV)[2] <- "ENV"
        MeansGxE <- suppressMessages(
            suppressWarnings(
                mutate(MeansGxE,
                       envPC1 = left_join(MeansGxE, EscENV %>% select(ENV, PC1))$PC1,
                       genPC1 = left_join(MeansGxE, EscGEN %>% select(GEN, PC1))$PC1,
                       nominal = left_join(MeansGxE, EscGEN %>% select(GEN, y))$y + genPC1 * envPC1)
            )
        )
        listres[[paste(names(vars[var]))]] <-
            structure(list(ANOVA = anova,
                           PCA = PC %>% rownames_to_column("PC") %>% as_tibble(),
                           MeansGxE = MeansGxE,
                           model = bplot %>% as_tibble(),
                           residuals = residuals %>% as_tibble(),
                           probint = probint),
                      class = "performs_ammi")
        if (verbose == TRUE) {
            cat("variable", paste(names(vars[var])),"\n")
            cat("---------------------------------------------------------------------------\n")
            cat("AMMI analysis table\n")
            cat("---------------------------------------------------------------------------\n")
            print(as.data.frame(anova), digits = 3, row.names = FALSE)
            cat("---------------------------------------------------------------------------\n\n")
        }
    }
    if (verbose == TRUE) {
        if (length(which(unlist(lapply(listres, function(x) {
            x[["probint"]]
        })) > 0.05)) > 0) {
            cat("------------------------------------------------------------\n")
            cat("Variables with nonsignificant GxE interaction\n")
            cat(names(which(unlist(lapply(listres, function(x) {
                x[["probint"]]
            })) > 0.05)), "\n")
            cat("------------------------------------------------------------\n")
        } else {
            cat("All variables with significant (p < 0.05) genotype-vs-environment interaction\n")
        }
        cat("Done!\n")
    }
    return(structure(listres, class = "performs_ammi"))
}
```
