## Supplementary material for "metan: an R package for multi-environment trial analysis": available at: plot.cvalidation.R

```
#' Plot the RMSPD of a cross-validation procedure
#'
#' Boxplot showing the Root Means Square Prediction Difference of of a cross
#' validation procedure.
#'
#' Five statistics are shown in this type of plot. The lower and upper hinges
#' correspond to the first and third quartiles (the 25th and 75th percentiles).
#' The upper whisker extends from the hinge to the largest value no further than
#' 1.5 * IQR from the hinge (where IQR is the inter-quartile range). The lower
#' whisker extends from the hinge to the smallest value at most 1.5 * IQR of the
#' hinge. Data beyond the end of the whiskers are considered outlying points.
#'
#' @param x An object of class \code{cvalidation} fitted with the functions
#'   \code{\link{cv_ammi}}, \code{\link{cv_ammif}}, \code{\link{cv_blup}}, or a
#'   bound object fitted with \code{\link{bind_cv}}.
#' @param violin Define if a violin plot is used with boxplot. Default is 'TRUE'
#' @param export Export (or not) the plot. Default is \code{T}.
#' @param order_box Logical argument. If \code{TRUE} then the boxplots will be
#'   ordered according to the values of the RMSPD.
#' @param x.lab The label of x-axis. New arguments can be inserted as
#'   \code{x.lab = 'my x label'}.
#' @param y.lab The label of y-axis. New arguments can be inserted as
#'   \code{y.lab = 'my y label'}.
#' @param size.tex.lab The size of the text in axis text and labels.
#' @param file.type The type of file to be exported. Default is \code{pdf},
#'   Graphic can also be exported in \code{*.tiff} format by declaring
#'   \code{file.type = 'tiff'}.
#' @param file.name The name of the file for exportation, default is
#'   \code{NULL}, i.e. the files are automatically named.
#' @param plot_theme The graphical theme of the plot. Default is
#'   \code{plot_theme = theme_metan()}. For more details,see
#'   \code{\link[ggplot2]{theme}}.
#' @param width The width 'inch' of the plot. Default is \code{6}.
#' @param height The height 'inch' of the plot. Default is \code{6}.
#' @param resolution The resolution of the plot. Parameter valid if
#'   \code{file.type = 'tiff'} is used. Default is \code{300} (300 dpi)
#' @param col.violin Parameter valid if \code{violin = T}. Define the color of
#'   the violin plot. Default is 'gray90.
#' @param col.boxplot Define the color for boxplot. Default is 'gray70'.
#' @param col.boxplot.win Define the color for boxplot of the best model.
#'   Default is 'cyan'.
#' @param width.boxplot The width of boxplots. Default is \code{0.2}.
#' @param x.lim The range of x-axis. Default is \code{NULL} (maximum and minimum
#'   values of the data set). New arguments can be inserted as \code{x.lim =
#'   c(x.min, x.max)}.
#' @param x.breaks The breaks to be plotted in the x-axis. Default is
#'   \code{authomatic breaks}. New arguments can be inserted as \code{x.breaks =
#'   c(breaks)}
#' @param ... Currently not used.
#' @return An object of class \code{gg, ggplot}.
#' @author Tiago Olivoto \email{tiagoolivoto@@gmail.com}
#' @method plot cvalidation
#' @export
#' @examples
#'
#' \donttest{
#' validation <- cv_ammif(data_ge,
#'                        resp = GY,
#'                        gen = GEN,
#'                        env = ENV,
#'                        rep = REP,
#'                        nboot = 5)
#' plot(validation)
#' }
#'
#'
plot.cvalidation <- function(x, violin = FALSE, export = FALSE, order_box =  FALSE,
                          x.lab = NULL, y.lab = NULL, size.tex.lab = 12, file.type = "pdf",
                          file.name = NULL, plot_theme = theme_metan(), width = 6, height = 6,
                          resolution = 300, col.violin = "gray90", col.boxplot = "gray70",
                          col.boxplot.win = "cyan", width.boxplot = 0.6, x.lim = NULL,
                          x.breaks = waiver(), ...) {
    if (!class(x) == "cvalidation") {
        stop("The object 'x' must be of class 'cvalidation'.")
    }
    y.lab <- ifelse(missing(y.lab),
                    expression(paste("Root mean square prediction difference (Mg ha"^-1, ")")),
                    y.lab)
    x.lab <- ifelse(missing(x.lab), "Model", x.lab)
    a <- suppressMessages(x$RMSPD %>%
                              group_by(MODEL) %>%
                              summarise(RMSPD = mean(RMSPD)) %>%
                              top_n(-1))
    mod <- paste(a$MODEL[1])
    if (violin == TRUE) {
        dodge <- position_dodge(width = 1)
        if(order_box == TRUE){
            p1 <- ggplot(x$RMSPD, aes(x = reorder(MODEL, RMSPD), y = RMSPD))
        } else {
            p1 <- ggplot(x$RMSPD, aes(x = MODEL, y = RMSPD))
        }
        p1 = p1 +
            geom_violin(position = dodge, fill = col.violin) +
            stat_boxplot(geom = "errorbar", width=width.boxplot/2, na.rm = TRUE)+
            geom_boxplot(width = width.boxplot,
                         position = dodge,
                         color = "black",
                         outlier.alpha = 0.75,
                         outlier.color = "black",
                         fill = col.boxplot) +
            geom_boxplot(data = x$RMSPD[x$RMSPD$MODEL == mod, ],
                         aes(x = MODEL, y = RMSPD),
                         width = width.boxplot,
                         position = dodge,
                         color = "black",
                         outlier.alpha = 0.75,
                         outlier.color = "black",
                         fill = col.boxplot.win) +
            stat_summary(fun.y = mean,
                         geom = "point",
                         shape = 23,
                         fill = "black") +
            plot_theme %+replace%
            theme(axis.text = element_text(size = size.tex.lab, colour = "black"),
                  axis.title = element_text(size = size.tex.lab, colour = "black")) +
            coord_flip() +
            scale_y_continuous(limits = x.lim, breaks = x.breaks) +
            labs(x = x.lab, y = y.lab)
    } else {
        dodge <- position_dodge(width = 1)
        if(order_box == TRUE){
            p1 <- ggplot(x$RMSPD, aes(x = reorder(MODEL, RMSPD), y = RMSPD))
        } else {
            p1 <- ggplot(x$RMSPD, aes(x = MODEL, y = RMSPD))
        }
        p1 = p1 +
            stat_boxplot(geom = "errorbar", width=width.boxplot/2, na.rm = TRUE)+
            geom_boxplot(width = width.boxplot,
                         position = dodge,
                         color = "black",
                         outlier.alpha = 0.75,
                         outlier.color = "black",
                         fill = col.boxplot) +
            geom_boxplot(data = x$RMSPD[x$RMSPD$MODEL == mod, ],
                         aes(x = MODEL, y = RMSPD),
                         width = width.boxplot,
                         position = dodge,
                         outlier.alpha = 0.75,
                         color = "black",
                         outlier.color = "black",
                         fill = col.boxplot.win) +
            stat_summary(fun.y = mean,
                         geom = "point",
                         shape = 23,
                         fill = "black") +
            plot_theme %+replace%
            theme(axis.text = element_text(size = size.tex.lab, colour = "black"),
                  axis.title = element_text(size = size.tex.lab, colour = "black")) +
            coord_flip() +
            scale_y_continuous(limits = x.lim, breaks = x.breaks) +
            labs(x = x.lab, y = y.lab)
    }
    if (export == FALSE) {
        return(p1)
    } else if (file.type == "pdf") {
        if (is.null(file.name)) {
            pdf("RMSPD validation.pdf", width = width, height = height)
        } else pdf(paste0(file.name, ".pdf"), width = width, height = height)
        plot(p1)
        dev.off()
    }
    if (file.type == "tiff") {
        if (is.null(file.name)) {
            tiff(filename = "RMSPD validation.tiff", width = width,
                 height = height, units = "in", compression = "lzw",
                 res = resolution)
        } else tiff(filename = paste0(file.name, ".tiff"), width = width,
                    height = height, units = "in", compression = "lzw",
                    res = resolution)
        plot(p1)
        dev.off()
    }
}
```
