## Supplementary material for "metan: an R package for multi-environment trial analysis": available at: plot.FAI_BLUP.R

```
#' Multi-trait selection index
#'
#' Plot the multitrait index based on factor analysis and ideotype-design
#' proposed by Rocha et al. (2018).
#'
#'
#' @param x An object of class \code{waasb}
#' @param ideotype The ideotype to be plotted. Default is 1.
#' @param SI An integer [0-100]. The selection intensity in percentage of the
#'   total number of genotypes.
#' @param radar Logical argument. If true (default) a radar plot is generated
#'   after using \code{coord_polar()}.
#' @param arrange.label Logical argument. If \code{TRUE}, the labels are
#'   arranged to avoid text overlapping. This becomes useful when the number of
#'   genotypes is large, say, more than 30.
#' @param size.point The size of the point in graphic.
#' @param col.sel The colour for selected genotypes.
#' @param col.nonsel The colour for nonselected genotypes.
#' @param size.text The size for the text in the plot. Defaults to 10.
#' @param ... Other arguments to be passed from ggplot2::theme().
#' @author Tiago Olivoto \email{tiagoolivoto@@gmail.com}
#' @references Rocha, J.R.A.S.C.R, J.C. Machado, and P.C.S. Carneiro. 2018.
#'   Multitrait index based on factor analysis and ideotype-design: proposal and
#'   application on elephant grass breeding for bioenergy. GCB Bioenergy
#'   10:52-60. doi:
#'   \href{https://onlinelibrary.wiley.com/doi/full/10.1111/gcbb.12443}{doi:10.1111/gcbb.12443}.
#'
#' @method plot fai_blup
#' @export
#' @return An object of class \code{gg, ggplot}.
#' @examples
#' \donttest{
#' library(metan)
#'
#' FAI = data_ge2 %>%
#'       waasb(ENV, GEN, REP, c(KW, NKE, PH, EH)) %>%
#'       fai_blup(DI = c('max, max, max, min'),
#'                UI = c('min, min, min, max'),
#'                SI = 15)
#' plot(FAI)
#' }
#'
plot.fai_blup <- function(x, ideotype = 1, SI = 15, radar = TRUE, arrange.label = FALSE,
                          size.point = 2, col.sel = "red", col.nonsel = "black", size.text = 10,
                          ...) {

    if (!class(x) == "fai_blup") {
        stop("The object 'x' is not of class 'fai_blup'")
    }
    data <- tibble(FAI = x$FAI[[ideotype]], Genotype = names(x$FAI[[ideotype]]),
                   sel = "Selected")
    data[["sel"]][(round(nrow(data) * (SI/100), 0) + 1):nrow(data)] <- "Nonselected"
    cutpoint <- min(subset(data, sel == "Selected")$FAI)
    p <- ggplot(data = data, aes(x = reorder(Genotype, FAI),
                                 y = FAI)) + geom_hline(yintercept = cutpoint, col = "red") +
        geom_path(colour = "black", group = 1) + geom_point(size = size.point,
                                                            aes(fill = sel), shape = 21, colour = "black") + scale_x_discrete() +
        theme_minimal() + theme(legend.position = "bottom", legend.title = element_blank(),
                                axis.title.x = element_blank(), panel.border = element_blank(),
                                axis.text = element_text(colour = "black"), text = element_text(size = size.text)) +
        labs(x = "", y = "FAI-BLUP") + scale_fill_manual(values = c(col.nonsel,
                                                                    col.sel))
    if (radar == TRUE) {
        if(arrange.label == TRUE){
        tot_gen <- length(unique(data$Genotype))
        fseq <- c(1:(tot_gen/2))
        sseq <- c((tot_gen/2 + 1):tot_gen)
        fang <- c(90 - 180/length(fseq) * fseq)
        sang <- c(-90 - 180/length(sseq) * sseq)
        p <- p + coord_polar() + theme(axis.text.x = element_text(angle = c(fang,
                                                                            sang)), legend.margin = margin(-120, 0, 0, 0), ...)
        } else{
            p <- p + coord_polar()
        }
    }
    return(p)
}
```
