## Supplementary material for "metan: an R package for multi-environment trial analysis": available at: plot.ge_cluster.R

```
#' Plot an object of class ge_cluster
#'
#' Plot an object of class ge_cluster
#'
#'
#' @param x An object of class \code{ge_cluster}
#' @param nclust The number of clusters to show.
#' @param xlab The label of the x axis.
#' @param ... Other arguments passed from the function \code{plot.hclust}.
#' @method plot ge_cluster
#' @export
#' @author Tiago Olivoto \email{tiagoolivoto@@gmail.com}
#'
plot.ge_cluster <- function(x, nclust = NULL, xlab = "", ...){
    plot(x$hc, hang = -1, xlab = xlab, sub = "", ...)
  if(!missing(nclust)){
  rect.hclust(x$hc, k = nclust, border = "red")
}
}
```
