## Supplementary material for "metan: an R package for multi-environment trial analysis": available at: plot.wsmp.R

```
#' Plot heat maps with genotype ranking
#'
#' Plot heat maps with genotype ranking in two ways.
#'
#' The first type of heatmap shows the genotype ranking depending on the number
#' of principal component axis used for estimating the WAASB index. An euclidean
#' distance-based dendrogram is used for grouping the genotype ranking for both
#' genotypes and principal component axis. The second type of heatmap shows the
#' genotype ranking depending on the WAASB/GY ratio. The ranks obtained with a
#' ratio of 100/0 considers exclusively the stability for the genotype ranking.
#' On the other hand, a ratio of 0/100 considers exclusively the productivity
#' for the genotype ranking.  Four clusters are estimated (1) unproductive and
#' unstable genotypes; (2) productive, but unstable genotypes; (3) stable, but
#' unproductive genotypes; and (4), productive and stable genotypes.
#'
#' @param x The object returned by the function \code{wsmp}.
#' @param var The variable to plot. Defaults to \code{var = 1} the first
#'   variable of \code{x}.
#' @param type \code{1 = Heat map Ranks}: this graphic shows the genotype
#'   ranking considering the WAAS estimated with different numbers of Principal
#'   Components; \code{2 = Heat map WAASY-GY ratio}: this graphic shows the
#'   genotype ranking considering the different combinations in the WAAS/GY
#'   ratio.
#' @param export Export (or not) the plot. Default is \code{FALSE}.
#' @param file.type If \code{export = TRUE} define the type of file to be
#'   exported. Default is \code{pdf}, Graphic can also be exported in
#'   \code{*.tiff} format by declaring \code{file.type = 'tiff'}.
#' @param file.name The name of the file for exportation, default is
#'   \code{NULL}, i.e. the files are automatically named.
#' @param width The width 'inch' of the plot. Default is \code{8}.
#' @param height The height 'inch' of the plot. Default is \code{7}.
#' @param size.lab The label size of the plot. It is suggested attribute 1
#' @param margins Numeric vector of length 2 containing the margins for column
#'   and row names, respectively. Default is \code{c(5, 4)}.
#' @param y.lab The label of y axis. Default is 'Genotypes'.
#' @param x.lab The label of x axis. Default is 'Number of axes'.
#' @param key.lab The label of color key. Default is 'Genotype ranking'.
#' @param resolution Valid parameter if \code{file.type = 'tiff'}. Define the
#'   resolution of the plot. Default is '300'.
#' @param ... Currently not used.
#' @author Tiago Olivoto \email{tiagoolivoto@@gmail.com}
#' @method plot wsmp
#' @export
#' @examples
#' \donttest{
#' library(metan)
#' model <- waas(data_ge2, ENV, GEN, REP, PH) %>%
#'          wsmp()
#' plot(model)
#' }
#'
plot.wsmp <- function(x, var = 1, type = 2, export = FALSE, file.type = "pdf",
                      file.name = NULL, width = 6, height = 5, size.lab = 1, margins = c(5, 4), y.lab = NULL, x.lab = NULL, key.lab = "Genotype ranking",
                      resolution = 300, ...) {
    data <- x[[var]]
    if (type == 1) {
        if (is.null(x.lab)) {
            x.lab <- "Number of axes"
        } else x.lab <- x.lab
        if (is.null(y.lab)) {
            y.lab <- "Genotypes"
        } else y.lab <- y.lab
        if (export == FALSE) {
            mat <- as.matrix(data$hetdata)
            Rowv <- mat %>% dist %>% hclust %>% as.dendrogram %>%
                dendextend::set("branches_k_color", k = 4) %>%
                dendextend::set("branches_lwd", 2) %>% sort(type = "nodes")
            Colv <- mat %>% t %>% dist %>% hclust %>% as.dendrogram %>%
                dendextend::set("branches_k_color", k = 3) %>%
                dendextend::set("branches_lwd", 2) %>% sort(type = "nodes")
            colfunc <- grDevices::colorRampPalette(c("green",
                                                     "yellow", "red"))
            gplots::heatmap.2(mat, dendrogram = "both", col = colfunc(nrow(subset(data$WAASY,
                                                                                  Code = "GEN"))), Rowv = Rowv, Colv = Colv, density.info = ("none"),
                              offsetRow = -0.4, offsetCol = -0.4, adjCol = c(1,
                                                                             0.5), cexCol = size.lab, cexRow = size.lab,
                              trace = "none", key.title = "", key.xlab = key.lab,
                              key.ylab = "", key.par = list(mgp = c(1.5, 0.5,
                                                                    0), mar = c(3, 0.2, 0.2, 0.2)), lhei = c(1,
                                                                                                             6), xlab = x.lab, ylab = y.lab, margins = margins)
        } else if (file.type == "pdf") {
            if (is.null(file.name)) {
                pdf("Heat Ranks PCA.pdf", width = width, height = height)
            } else pdf(paste0(file.name, ".pdf"), width = width,
                       height = height)
            Rowv <- mat %>% dist %>% hclust %>% as.dendrogram %>%
                dendextend::set("branches_k_color", k = 4) %>%
                dendextend::set("branches_lwd", 2) %>% sort(type = "nodes")
            Colv <- mat %>% t %>% dist %>% hclust %>% as.dendrogram %>%
                dendextend::set("branches_k_color", k = 3) %>%
                dendextend::set("branches_lwd", 2) %>% sort(type = "nodes")
            colfunc <- grDevices::colorRampPalette(c("green",
                                                     "yellow", "red"))
            gplots::heatmap.2(mat, dendrogram = "both", col = colfunc(nrow(subset(data$WAASY,
                                                                                  Code = "GEN"))), Rowv = Rowv, Colv = Colv, density.info = ("none"),
                              offsetRow = -0.4, offsetCol = -0.4, adjCol = c(1,
                                                                             0.5), cexCol = size.lab, cexRow = size.lab,
                              trace = "none", key.title = "", key.xlab = key.lab,
                              key.ylab = "", key.par = list(mgp = c(1.5, 0.5,
                                                                    0), mar = c(3, 0.2, 0.2, 0.2)), lhei = c(1,
                                                                                                             6), xlab = x.lab, ylab = y.lab, margins = margins)
            dev.off()
        }
        if (file.type == "tiff") {
            if (is.null(file.name)) {
                tiff(filename = "Heat Ranks PCA.tiff", width = width,
                     height = height, units = "in", compression = "lzw",
                     res = resolution)
            } else tiff(filename = paste0(file.name, ".tiff"),
                        width = width, height = height, units = "in",
                        compression = "lzw", res = resolution)
            Rowv <- mat %>% dist %>% hclust %>% as.dendrogram %>%
                dendextend::set("branches_k_color", k = 4) %>%
                dendextend::set("branches_lwd", 2) %>% sort(type = "nodes")
            Colv <- mat %>% t %>% dist %>% hclust %>% as.dendrogram %>%
                dendextend::set("branches_k_color", k = 3) %>%
                dendextend::set("branches_lwd", 2) %>% sort(type = "nodes")
            colfunc <- grDevices::colorRampPalette(c("green",
                                                     "yellow", "red"))
            gplots::heatmap.2(mat, dendrogram = "both", col = colfunc(nrow(subset(data$WAASY,
                                                                                  Code = "GEN"))), Rowv = Rowv, Colv = Colv, density.info = ("none"),
                              offsetRow = -0.4, offsetCol = -0.4, adjCol = c(1,
                                                                             0.5), cexCol = size.lab, cexRow = size.lab,
                              trace = "none", key.title = "", key.xlab = key.lab,
                              key.ylab = "", key.par = list(mgp = c(1.5, 0.5,
                                                                    0), mar = c(3, 0.2, 0.2, 0.2)), lhei = c(1,
                                                                                                             6), xlab = x.lab, ylab = y.lab, margins = margins)
            dev.off()
        }
    }
    if (type == 2) {
        if (is.null(x.lab)) {
            x.lab <- "WAASB/GY ratio"
        } else x.lab <- x.lab
        if (is.null(y.lab)) {
            y.lab <- "Genotypes"
        } else y.lab <- y.lab
        mat2 <- as.matrix(data$hetcomb)
        if (export == FALSE) {
            Rowv <- mat2 %>% dist %>% hclust %>% as.dendrogram %>%
                dendextend::set("branches_k_color", k = 4) %>%
                dendextend::set("branches_lwd", 2) %>% dendextend::rotate_DendSer(ser_weight = dist(mat2)) %>%
                sort(type = "nodes")
            colfunc <- grDevices::colorRampPalette(c("green",
                                                     "yellow", "red"))
            gplots::heatmap.2(mat2, dendrogram = "row", col = colfunc(nrow(subset(data$WAASY,
                                                                                  Code = "GEN"))), Rowv = Rowv, Colv = F, density.info = ("none"),
                              offsetRow = -0.4, offsetCol = -0.4, adjCol = c(1,
                                                                             0.5), trace = "none", key.title = "", key.xlab = key.lab,
                              key.ylab = "", key.par = list(mgp = c(1.5, 0.5,
                                                                    0), mar = c(3, 0.2, 0.2, 0.2)), lhei = c(1,
                                                                                                             6), xlab = x.lab, ylab = y.lab, cexCol = size.lab,
                              cexRow = size.lab, margins = margins)
        } else if (file.type == "pdf") {
            if (is.null(file.name)) {
                pdf("Heat map Ranks WAAS-GY.pdf", width = width,
                    height = height)
            } else pdf(paste0(file.name, ".pdf"), width = width,
                       height = height)
            Rowv <- mat2 %>% dist %>% hclust %>% as.dendrogram %>%
                dendextend::set("branches_k_color", k = 4) %>%
                dendextend::set("branches_lwd", 2) %>% dendextend::rotate_DendSer(ser_weight = dist(mat2)) %>%
                sort(type = "nodes")
            colfunc <- grDevices::colorRampPalette(c("green",
                                                     "yellow", "red"))
            gplots::heatmap.2(mat2, dendrogram = "row", col = colfunc(nrow(subset(data$WAASY,
                                                                                  Code = "GEN"))), Rowv = Rowv, Colv = F, density.info = ("none"),
                              offsetRow = -0.4, offsetCol = -0.4, adjCol = c(1,
                                                                             0.5), trace = "none", key.title = "", key.xlab = key.lab,
                              key.ylab = "", key.par = list(mgp = c(1.5, 0.5,
                                                                    0), mar = c(3, 0.2, 0.2, 0.2)), lhei = c(1,
                                                                                                             6), xlab = x.lab, ylab = y.lab, cexCol = size.lab,
                              cexRow = size.lab, margins = margins)
            dev.off()
        }
        if (file.type == "tiff") {
            if (is.null(file.name)) {
                tiff(filename = "Heat map Ranks WAAS-GY.tiff",
                     width = width, height = height, units = "in",
                     compression = "lzw", res = resolution)
            } else tiff(filename = paste0(file.name, ".tiff"),
                        width = width, height = height, units = "in",
                        compression = "lzw", res = resolution)
            Rowv <- mat2 %>% dist %>% hclust %>% as.dendrogram %>%
                dendextend::set("branches_k_color", k = 4) %>%
                dendextend::set("branches_lwd", 2) %>% dendextend::rotate_DendSer(ser_weight = dist(mat2)) %>%
                sort(type = "nodes")
            colfunc <- grDevices::colorRampPalette(c("green",
                                                     "yellow", "red"))
            gplots::heatmap.2(mat2, dendrogram = "row", col = colfunc(nrow(subset(data$WAASY,
                                                                                  Code = "GEN"))), Rowv = Rowv, Colv = F, density.info = ("none"),
                              offsetRow = -0.4, offsetCol = -0.4, adjCol = c(1,
                                                                             0.5), trace = "none", key.title = "", key.xlab = key.lab,
                              key.ylab = "", key.par = list(mgp = c(1.5, 0.5,
                                                                    0), mar = c(3, 0.2, 0.2, 0.2)), lhei = c(1,
                                                                                                             6), xlab = x.lab, ylab = y.lab, cexCol = size.lab,
                              cexRow = size.lab, margins = margins)
            dev.off()
        }
    }
}
```
