## Supplementary material for "metan: an R package for multi-environment trial analysis": available at: plot_blup.R

```
#' Plot the BLUPs for genotypes
#'
#' Plot the predicted BLUP of the genotypes.
#'
#'
#' @param x The \code{waasb object}
#' @param var The variable to plot. Defaults to \code{var = 1} the first
#'   variable of \code{x}.
#' @param prob The probability error for constructing confidence interval.
#' @param export Export (or not) the plot. Default is \code{TRUE}.
#' @param file.type If \code{export = TRUE}, define the type of file to be
#'   exported. Default is \code{pdf}, Graphic can also be exported in
#'   \code{*.tiff} format by declaring \code{file.type = "tiff"}.
#' @param file.name The name of the file for exportation, default is
#'   \code{NULL}, i.e. the files are automatically named.
#' @param plot_theme The graphical theme of the plot. Default is
#'   \code{plot_theme = theme_metan()}. For more details, see
#'   \code{\link[ggplot2]{theme}}.
#' @param width The width "inch" of the plot. Default is \code{6}.
#' @param height The height "inch" of the plot. Default is \code{6}.
#' @param size.err.bar The size of the error bar for the plot. Default is
#'   \code{0.5}.
#' @param size.shape The size of the shape (both for genotypes and
#'   environments). Default is \code{3.5}.
#' @param size.tex.lab The size of the text in axis text and labels.
#' @param height.err.bar The height for error bar. Default is \code{0.3}.
#' @param x.lim The range of x-axis. Default is \code{NULL} (maximum and minimum
#'   values of the data set). New arguments can be inserted as \code{x.lim =
#'   c(x.min, x.max)}.
#' @param x.breaks The breaks to be plotted in the x-axis. Default is
#'   \code{authomatic breaks}. New arguments can be inserted as \code{x.breaks =
#'   c(breaks)}
#' @param col.shape A vector of length 2 that contains the color of shapes for
#'   genotypes above and below of the mean, respectively. Default is
#'   \code{c("blue", "red")}.
#' @param x.lab The label of the x-axis in the plot. Default is \code{"Predicted
#'   Grain Yield"}.
#' @param y.lab The label of the y-axis in the plot. Default is
#'   \code{"Genotypes"}.
#' @param resolution The resolution of the plot. Parameter valid if
#'   \code{file.type = "tiff"} is used. Default is \code{300} (300 dpi)
#' @param ... Currently not used.
#' @return An object of class \code{gg, ggplot}.
#' @author Tiago Olivoto \email{tiagoolivoto@@gmail.com}
#' @seealso \code{\link{plot_scores}}, \code{\link{plot_waasby}}
#' @export
#' @examples
#'\donttest{
#' library(metan)
#' BLUP <- waasb(data_ge,
#'               resp = c(GY, HM),
#'               gen = GEN,
#'               env = ENV,
#'               rep = REP)
#' plot_blup(BLUP)
#'
#'}
#'
#'
plot_blup <- function(x, var = 1, prob = 0.05, export = FALSE, file.type = "pdf", file.name = NULL,
                      plot_theme = theme_metan(), width = 6, height = 6, size.err.bar = 0.5, size.shape = 3.5,
                      size.tex.lab = 12, height.err.bar = 0.3, x.lim = NULL, x.breaks = waiver(),
                      col.shape = c("blue", "red"), y.lab = "Genotypes", x.lab = "Predicted Grain Yield",
                      resolution = 300, ...) {
    x <- x[[var]]
    if(!class(x)  %in% c("waasb", "gamem")){
        stop("The object 'x' must be of class 'waasb' or 'gamem'.")
    }
    if(class(x) == "gamem"){
        PROB <- ((1 - (1 - prob))/2) + (1 - prob)
        t <- qt(PROB, nlevels(x$residuals$REP))
        GV <- as.numeric(x$ESTIMATES[1, 2])
        AccuGen <- as.numeric(x$ESTIMATES[8, 2])
        Limits <- t * sqrt(((1 - AccuGen) * GV))
        blup <- x$blupGEN %>%
            mutate(LL = Predicted - Limits,
                   UL = Predicted + Limits,
                   Mean = ifelse(Predicted < mean(Predicted), "below", "above")) %>%
            arrange(Predicted)
    }
    if(class(x) ==  "waasb"){
        PROB <- ((1 - (1 - prob))/2) + (1 - prob)
        t <- qt(PROB, nlevels(x$residuals$REP))
        GV <- as.numeric(x$ESTIMATES[3, 2])
        AccuGen <- as.numeric(x$ESTIMATES[11, 2])
        Limits <- t * sqrt(((1 - AccuGen) * GV))
        blup <- x$blupGEN %>%
            mutate(LL = Predicted - Limits,
                   UL = Predicted + Limits,
                   Mean = ifelse(Predicted < mean(Predicted), "below", "above")) %>%
            arrange(Predicted)
    }
    p1 <-ggplot(blup, aes(x = Predicted, y = reorder(GEN, Predicted))) +
        geom_vline(xintercept = mean(blup$Predicted)) +
        geom_errorbarh(aes(xmin = LL, xmax = UL), size = size.err.bar, height = height.err.bar) +
        geom_point(stat = "identity", aes(fill = Mean), shape = 21, size = size.shape) +
        scale_fill_manual(name = "Average", values = col.shape, labels = c("Above", "Below")) +
        labs(x = x.lab, y = y.lab) +
        scale_x_continuous(limits = x.lim, breaks = x.breaks) +
        plot_theme %+replace% theme(axis.text = element_text(size = size.tex.lab,colour = "black"),
                               axis.title = element_text(size = size.tex.lab, colour = "black"))


    if (export == FALSE) {
        return(p1)
    } else if (file.type == "pdf") {
        if (is.null(file.name)) {
            pdf("BLUPs genotypes.pdf", width = width, height = height)
        } else pdf(paste0(file.name, ".pdf"), width = width, height = height)
        plot(p1)
        dev.off()
    }
    if (file.type == "tiff") {
        if (is.null(file.name)) {
            tiff(filename = "BLUPs genotypes.tiff", width = width, height = height,
                 units = "in", compression = "lzw", res = resolution)
        } else tiff(filename = paste0(file.name, ".tiff"), width = width, height = height,
                    units = "in", compression = "lzw", res = resolution)
        plot(p1)
        dev.off()
    }
}
```
