## Supplementary material for "metan: an R package for multi-environment trial analysis": available at: plot_eigen.R

```
#' Plot the eigenvalues
#'
#' Plot the eigenvalues for from singular value decomposition of BLUP
#' interaction effects matrix.
#'
#'
#' @param x The \code{waasb object}
#' @param var The variable to plot. Defaults to \code{var = 1} the first
#'   variable of \code{x}.
#' @param export Export (or not) the plot. Default is \code{TRUE}.
#' @param plot_theme The graphical theme of the plot. Default is
#'   \code{plot_theme = theme_metan()}. For more details, see
#'   \code{\link[ggplot2]{theme}}.
#' @param file.type If \code{export = TRUE}, define the type of file to be
#'   exported. Default is \code{pdf}, Graphic can also be exported in
#'   \code{*.tiff} format by declaring \code{file.type = "tiff"}.
#' @param file.name The name of the file for exportation, default is
#'   \code{NULL}, i.e. the files are automatically named.
#' @param width The width "inch" of the plot. Default is \code{6}.
#' @param height The height "inch" of the plot. Default is \code{6}.
#' @param size.shape The size of the shape. Default is \code{3.5}.
#' @param size.line The size of the line. Default is \code{1}.
#' @param size.tex.lab The size of the text in axis text and labels.
#' @param y.lab The label of the y-axis in the plot. Default is
#'   \code{"Eigenvalue"}.
#' @param y2.lab The label of the second y-axis in the plot. Default is
#'   \code{"Accumulated variance"}.
#' @param x.lab The label of the x-axis in the plot. Default is \code{"Number of
#'   multiplicative terms"}.
#' @param resolution The resolution of the plot. Parameter valid if
#'   \code{file.type = "tiff"} is used. Default is \code{300} (300 dpi)
#' @param ... Currently not used.
#' @return An object of class \code{gg, ggplot}.
#' @author Tiago Olivoto \email{tiagoolivoto@@gmail.com}
#' @seealso \code{\link{plot_scores}}, \code{\link{plot_waasby}}
#' @export
#' @examples
#' \donttest{
#' library(metan)
#' BLUP <- waasb(data_ge,
#'              resp = c(GY, HM),
#'              gen = GEN,
#'              env = ENV,
#'              rep = REP)
#' plot_eigen(BLUP)
#'
#'}
#'
plot_eigen <- function(x, var = 1, export = FALSE, plot_theme = theme_metan(), file.type = "pdf",
    file.name = NULL, width = 6, height = 6, size.shape = 3.5, size.line = 1, size.tex.lab = 12,
    y.lab = "Eigenvalue", y2.lab = "Accumulated variance", x.lab = "Number of multiplicative terms",
    resolution = 300, ...) {
    x <- x[[var]]
    class <- class(x)
    if(!class(x) ==  "waasb"){
        stop("The object 'x' must be of class 'waasb'.")
    }
    eigen <- x$PCA
    eigen$PC <- factor(eigen$PC, levels = eigen$PC)
    scaleFactor <- 100/max(eigen$Eigenvalue)

    p <- ggplot2::ggplot(eigen, aes(x = PC, group = 1)) + geom_line(aes(y = Eigenvalue,
        col = "Eigenvalue"), size = size.line) + geom_point(aes(y = Eigenvalue, col = "Eigenvalue"),
        size = size.shape) + geom_line(aes(y = Accumulated/scaleFactor, col = "Percentage"),
        size = size.line) + geom_point(aes(y = Accumulated/scaleFactor, col = "Percentage"),
        size = size.shape) + scale_y_continuous(sec.axis = sec_axis(~. * scaleFactor,
        name = y2.lab)) + labs(x = x.lab, y = y.lab) +
        plot_theme %+replace%
        theme(axis.text = element_text(size = size.tex.lab,
        colour = "black"), axis.title = element_text(size = size.tex.lab, colour = "black"),
        legend.position = c(0.15, 0.1))


    if (export == FALSE) {
        return(p)
    } else if (file.type == "pdf") {
        if (is.null(file.name)) {
            pdf("Eigenvalues.pdf", width = width, height = height)
        } else pdf(paste0(file.name, ".pdf"), width = width, height = height)
        plot(p)
        dev.off()
    }

    if (file.type == "tiff") {
        if (is.null(file.name)) {
            tiff(filename = "Eigenvalues.tiff", width = width, height = height, units = "in",
                compression = "lzw", res = resolution)
        } else tiff(filename = paste0(file.name, ".tiff"), width = width, height = height,
            units = "in", compression = "lzw", res = resolution)
        plot(p)
        dev.off()
    }
}
```
