## Supplementary material for "metan: an R package for multi-environment trial analysis": available at: plot_factlines.R

```
#' @title Fast way to create a line plot
#' @description Create a graphic to show a fitted line based on numerical
#'   variables and one grouping variable.
#' @param .data The data set
#' @param x The variable in data to be shown in the x axis
#' @param y The variable in data to be shown in the y axis
#' @param group The grouping variable
#' @param fit The polynomial degree to use. It must be an integer between 1
#'   (linear fit) to 4 (fourth-order polynomial regression.), or a numeric
#'   vector with the same length of the variable in \code{group}
#' @param level The fonfidence level
#' @param confidence Display confidence interval around smooth? (TRUE by
#'   default)
#' @param xlab The x label
#' @param ylab The y label
#' @param legend.position The position of the legend. Defaults to 'bottom'.
#' @param grid Logical argument. If \code{TRUE} then a grid will be created.
#' @param scales If \code{grid = TRUE} scales controls how the scales are in the
#'   plot. Possible values are 'free' (default), 'fixed', 'free_x' or 'free_y'.
#' @param col The colour to be used in the line plot and points
#' @param alpha The alpha for the color in confidence band
#' @param size.shape The size for the shape in plot
#' @param size.line The size for the line in the plot
#' @param size.text The size of the text
#' @param fontfam The family of the font text
#' @param plot_theme The graphical theme of the plot. Default is
#'   \code{plot_theme = theme_metan()}. For more details, see
#'   \code{\link[ggplot2]{theme}}.
#' @return An object of class \code{gg, ggplot}.
#' @export
#' @seealso \code{\link{plot_lines}}, \code{\link{plot_factbars}}
plot_factlines <- function(.data, x, y, group, fit, level = 0.95,
                           confidence = TRUE, xlab = NULL, ylab = NULL, legend.position = "bottom",
                           grid = FALSE, scales = "free", col = TRUE, alpha = 0.2, size.shape = 1.5,
                           size.line = 1, size.text = 12, fontfam = "sans", plot_theme = theme_metan()) {
  if (length(fit) == 1 & grid == TRUE) {
    stop("When grid is TRUE the argument fit must have the same length of the grouping variable.")
  }
  if (max(fit) >= 5) {
    stop("The maximum polynomial degree is 4.")
  }
  cl <- match.call()
  data2 <- .data %>% select(factors = !!enquo(group), x = !!enquo(x),
                            y = !!enquo(y))
  group <- as.factor(data2$factors)
  p_smooth <- list()
  levels <- levels(group)
  if (length(fit) > 1) {
    for (i in 1:length(levels)) {
      levelname <- levels[i]
      mycond <- quote(group == levelname)
      if (fit[i] == 1) {
        formula <- as.formula("y ~ x")
      }
      if (fit[i] == 2) {
        formula <- as.formula("y ~ poly(x, 2)")
      }
      if (fit[i] == 3) {
        formula <- as.formula("y ~ poly(x, 3)")
      }
      if (fit[i] == 4) {
        formula <- as.formula("y ~ poly(x, 4)")
      }
      if (col == FALSE) {
        linetype <- i
        p_smooth[[paste(levels[i])]] <- ggplot2::stat_smooth(method = "lm",
                                                             formula = formula, data = subset(data2, eval(mycond)),
                                                             level = level, linetype = linetype, alpha = alpha,
                                                             col = "black", size = size.line, se = confidence)
      } else {
        linetype <- 1
        p_smooth[[paste(levels[i])]] <- ggplot2::stat_smooth(method = "lm",
                                                             formula = formula, data = subset(data2, eval(mycond)),
                                                             level = level, linetype = linetype, alpha = alpha,
                                                             size = size.line, se = confidence)
      }
    }
  } else {
    if (fit == 1) {
      formula <- as.formula("y ~ x")
    }
    if (fit == 2) {
      formula <- as.formula("y ~ poly(x, 2)")
    }
    if (fit == 3) {
      formula <- as.formula("y ~ poly(x, 3)")
    }
    if (fit == 4) {
      formula <- as.formula("y ~ poly(x, 4)")
    }
    if (col == TRUE) {
      p_smooth <- ggplot2::stat_smooth(method = "lm", formula = formula,
                                       data = data2, level = level, linetype = 1, size = size.line,
                                       se = confidence)
    } else {
      p_smooth <- ggplot2::stat_smooth(method = "lm", formula = formula,
                                       data = data2, level = level, linetype = 1, col = "black",
                                       size = size.line, se = confidence)
    }
  }
  if (grid == TRUE) {
    legend.position <- "none"
  } else {
    legend.position <- legend.position
  }
if (is.null(ylab) == TRUE) {
    ylab <- cl$y
  } else {
    ylab <- ylab
  }
  if (is.null(xlab) == TRUE) {
    xlab <- cl$x
  } else {
    xlab <- xlab
  }
  if (col == FALSE) {
    p <- ggplot2::ggplot(data2, aes(x = x, y = y)) + ggplot2::geom_point(aes(shape = factors),
                                                                         size = size.shape)
  } else {
    if (length(fit) > 1) {
      p <- ggplot2::ggplot(data2, aes(x = x, y = y, colour = factors)) +
        ggplot2::geom_point(aes(colour = factors), size = size.shape)
    } else {
      p <- ggplot2::ggplot(data2, aes(x = x, y = y)) +
        ggplot2::geom_point(aes(colour = factors), size = size.shape)
    }
  }
  p <- p + p_smooth + plot_theme %+replace% theme(axis.ticks.length = unit(0.2,
                                                                      "cm"), axis.text = element_text(size = size.text, family = fontfam,
                                                                                                      colour = "black"), axis.title = element_text(size = size.text,
                                                                                                                                                   family = fontfam, colour = "black"), axis.ticks = element_line(colour = "black"),
                                             plot.margin = margin(0.2, 0.2, 0.2, 0.2, "cm"), legend.title = element_blank(),
                                             legend.position = legend.position, legend.text = element_text(size = size.text,
                                                                                                           family = fontfam)) + ggplot2::labs(y = ylab, x = xlab)
  if (grid == TRUE) {
    p <- p + ggplot2::facet_wrap(~factors, scales = scales)
  } else {
    p <- p
  }
  return(p)
}
```
