## Supplementary material for "metan: an R package for multi-environment trial analysis": available at: predict.gge.R

```
#' Predict a two-way table based on GGE model
#'
#' Predict the means for a genotype-vs-environment trial based on a Genotype
#' plus Genotype-vs-Environment interaction (GGE) model.
#'
#' This function is used to predict the response variable of a two-way table
#' (for examples the yielding of g genotypes in e environments) based on GGE
#' model. This prediction is based on the number of principal components used.
#' For more details see Yan and Kang (2007).
#'
#' @param object An object of class \code{gge}.
#' @param naxis The the number of principal components to be used in the
#'   prediction. Generally, two axis may be used. In this case, the estimated
#'   values will be those shown in the biplot.
#' @param output The type of output. It must be one of the \code{'long'}
#'   (default) returning a long-format table with the columns for environment
#'   (ENV), genotypes (GEN) and response variable (Y); or \code{'wide'} to
#'   return a two-way table with genotypes in the row, environments in the
#'   columns, filled by the estimated values.
#' @param ... Additional parameter for the function
#' @return A two-way table with genotypes in rows and environments in columns if
#'   \code{output = "wide"} or a long format (columns ENV, GEN and Y) if
#'   \code{output = "long"} with the predicted values by the GGE model.
#' @author Tiago Olivoto \email{tiagoolivoto@@gmail.com}
#' @references Yan, W., and M.S. Kang. 2003. GGE biplot analysis: a graphical
#'   tool for breeders, geneticists, and agronomists. CRC Press.
#' @method predict gge
#' @export
#' @examples
#'
#' library(metan)
#' mod <- gge(data_ge, GEN, ENV, c(GY, HM))
#' predict(mod)
#'
predict.gge <- function(object, naxis = 2, output = "wide", ...) {
  if (!class(object) == "gge") {
    stop("The object must be of class 'gge'.")
  }
  listres <- list()
  varin <- 1
  for (var in 1:length(object)) {
    objectin <- object[[var]]
  if (naxis > min(dim(objectin$coordenv))) {
    stop("The number of principal components cannot be greater than min(g, e), in this case ",
         min(dim(objectin$coordenv)))
  }
  # SVP
  if (objectin$svp == "environment" | objectin$svp == 2) {
    pred <- (objectin$coordgen[, 1:naxis] * (objectin$d)) %*%
      t(objectin$coordenv[, 1:naxis])
  }
  if (objectin$svp == "genotype" | objectin$svp == 1) {
    pred <- (objectin$coordgen[, 1:naxis] %*% t(objectin$coordenv[,
                                                              1:naxis] * (objectin$d)))
  }
  if (objectin$svp == "symmetrical" | objectin$svp == 3) {
    pred <- (objectin$coordgen[, 1:naxis] %*% t(objectin$coordenv[,
                                                              1:naxis]))
  }
  # Scaling
  if (objectin$scaling == "sd" | objectin$scaling == 1) {
    pred <- sweep(pred, 2, objectin$scale_val, FUN = "*")
  }
  # Centering
  if (objectin$centering == "global" | objectin$centering == 1) {
    pred <- pred + objectin$grand_mean
  }
  if (objectin$centering == "environment" | objectin$centering ==
      2) {
    pred <- sweep(pred, 2, objectin$mean_env, FUN = "+")
  }
  if (objectin$centering == "double" | objectin$centering == 3) {
    for (i in 1:nrow(pred)) {
      for (j in 1:ncol(pred)) {
        pred[i, j] <- pred[i, j] - objectin$grand_mean +
          objectin$mean_env[j] + objectin$mean_gen[i]
      }
    }
  }
  rownames(pred) <- objectin$labelgen
  colnames(pred) <- objectin$labelenv
  if (output == "wide") {
    temp <- as_tibble(pred, rownames = NA)
  }
  if (output == "long") {
    temp <-
    pred %>%
      make_long() %>%
      as_tibble()
  }
  listres[[paste(names(object[var]))]] <- temp
  }
  return(listres)
}
```
