## Supplementary material for "metan: an R package for multi-environment trial analysis": available at: print.Ammi_indexes.R

```
#' Print an object of class AMMI_indexes
#'
#' Print the \code{AMMI_indexes} object in two ways. By default, the results are shown
#' in the R console. The results can also be exported to the directory into a
#' *.txt file.
#'
#' @param x An object of class \code{AMMI_indexes}.
#' @param export A logical argument. If \code{TRUE}, a *.txt file is exported to
#'   the working directory.
#' @param file.name The name of the file if \code{export = TRUE}
#' @param digits The significant digits to be shown.
#' @param ... Options used by the tibble package to format the output. See
#'   \code{\link[tibble:formatting]{tibble::print()}} for more details.
#' @author Tiago Olivoto \email{tiagoolivoto@@gmail.com}
#' @method print AMMI_indexes
#' @export
#' @examples
#'
#' library(metan)
#' model <- performs_ammi(data_ge, ENV, GEN, REP, GY) %>%
#'          AMMI_indexes()
#' print(model)
print.AMMI_indexes <- function(x, export = FALSE, file.name = NULL, digits = 3, ...) {
  if (!class(x) == "AMMI_indexes") {
    stop("The object must be of class 'AMMI_indexes'")
  }
  opar <- options(pillar.sigfig = digits)
  on.exit(options(opar))
  if (export == TRUE) {
    file.name <- ifelse(is.null(file.name) == TRUE, "AMMI_indexes print", file.name)
    sink(paste0(file.name, ".txt"))
  }
  for (i in 1:length(x)) {
    var <- x[[i]]
    cat("Variable", names(x)[i], "\n")
    cat("---------------------------------------------------------------------------\n")
    cat("AMMI-based stability indexes\n")
    cat("---------------------------------------------------------------------------\n")
    print(var$statistics)
    cat("---------------------------------------------------------------------------\n")
    cat("Ranks for stability indexes\n")
    cat("---------------------------------------------------------------------------\n")
    print(var$ranks)
    cat("---------------------------------------------------------------------------\n")
    cat("Simultaneous selection indexes\n")
    cat("---------------------------------------------------------------------------\n")
    print(var$ssi)
    cat("\n\n\n")
  }
  if (export == TRUE) {
    sink()
  }
}
```
