## Supplementary material for "metan: an R package for multi-environment trial analysis": available at: print.gamem.R

```
#' Print an object of class gamem
#'
#' Print the \code{gamem} object in two ways. By default, the results are shown
#' in the R console. The results can also be exported to the directory.
#'
#'
#' @param x An object fitted with the function \code{\link{gamem}} .
#' @param export A logical argument. If \code{TRUE}, a *.txt file is exported to
#'   the working directory
#' @param file.name The name of the file if \code{export = TRUE}
#' @param digits The significant digits to be shown.
#' @param ... Options used by the tibble package to format the output. See
#'   \code{\link[tibble:formatting]{tibble::print()}} for more details.
#' @author Tiago Olivoto \email{tiagoolivoto@@gmail.com}
#' @method print gamem
#' @export
#' @examples
#'
#' library(metan)
#' alpha <- gamem(data_alpha,
#'   gen = GEN,
#'   rep = REP,
#'   block = BLOCK,
#'   resp = YIELD
#' )
#'
#' print(alpha)
print.gamem <- function(x, export = FALSE, file.name = NULL, digits = 4, ...) {
  if (!class(x) == "gamem") {
    stop("The object must be of class 'gamem'")
  }
  if (export == TRUE) {
    file.name <- ifelse(is.null(file.name) == TRUE, "gamem print", file.name)
    sink(paste0(file.name, ".txt"))
  }
  opar <- options(pillar.sigfig = digits)
  on.exit(options(opar))
  for (i in 1:length(x)) {
    var <- x[[i]]
    cat("Variable", names(x)[i], "\n")
    cat("---------------------------------------------------------------------------\n")
    cat("Fixed-effect anova table\n")
    cat("---------------------------------------------------------------------------\n")
    print(var$fixed, ...)
    cat("---------------------------------------------------------------------------\n")
    cat("Variance components for random effects\n")
    cat("---------------------------------------------------------------------------\n")
    print(var$random, ...)
    cat("---------------------------------------------------------------------------\n")
    cat("Likelihood ratio test for random effects\n")
    cat("---------------------------------------------------------------------------\n")
    print(var$LRT, ...)
    cat("---------------------------------------------------------------------------\n")
    cat("Details of the analysis\n")
    cat("---------------------------------------------------------------------------\n")
    print(var$Details, ...)
    cat("---------------------------------------------------------------------------\n")
    cat("Genetic parameters\n")
    cat("---------------------------------------------------------------------------\n")
    print(var$ESTIMATES, ...)
    cat("\n\n\n")
  }
  if (export == TRUE) {
    sink()
  }
}
```
