## Supplementary material for "metan: an R package for multi-environment trial analysis": available at: resca.R

```
#' Rescale a variable to have specified minimum and maximum values
#'
#' Helper function that rescales a continuous variable to have specified minimum
#' and maximum values.
#'
#' The function rescale a continuous variable as follows: \deqn{Rv_i = (Nmax -
#' Nmin)/(Omax - Omin) * (O_i - Omax) + Nmax} Where \eqn{Rv_i} is the rescaled
#' value of the ith position of the variable/ vector; \eqn{Nmax} and \eqn{Nmin}
#' are the new maximum and minimum values; \eqn{Omax and Omin} are the maximum
#' and minimum values of the original data, and \eqn{O_i} is the ith value of
#' the original data.
#'
#' There are basically two options to use \code{resca} to rescale a variable.
#' The first is passing a data frame to \code{.data} argument and selecting one
#' or more variables to be scaled using \code{...}. The function will return the
#' original variables in \code{.data} plus the rescaled variable(s) with the
#' prefix \code{_res}. By using the function \code{group_by} from \bold{dplyr}
#' package it is possible to rescale the variable(s) within each level of the
#' grouping factor. The second option is pass a numeric vector in the argument
#' \code{values}. The output, of course, will be a numeric vector of rescaled
#' values.
#'
#' @param .data The dataset. Grouped data is allowed.
#' @param ... Comma-separated list of unquoted variable names that will be
#'   rescaled.
#' @param values Optional vector of values to rescale
#' @param new_min The minimum value of the new scale. Default is 0.
#' @param new_max The maximum value of the new scale. Default is 100
#' @param keep Should all variables be kept after rescaling? If false, only
#'   rescaled variables will be kept.
#' @return A numeric vector if \code{values} is used as input data or a tibble
#'   if a data frame is used as input in \code{.data}.
#' @author Tiago Olivoto \email{tiagoolivoto@@gmail.com}
#' @export
#' @examples
#' library(metan)
#' library(dplyr)
#' # Rescale a numeric vector
#' resca(values = c(1:5))
#'
#'  # Using a data frame
#' head(
#'  resca(data_ge, GY, HM, new_min = 0, new_max = 1)
#' )
#'
#' # Rescale within factors;
#' # Select variables that stats with 'N' and ends with 'L';
#' # Compute the mean of these variables by ENV and GEN;
#' # Rescale the variables that ends with 'L' whithin ENV;
#' data_ge2 %>%
#'   select(ENV, GEN, starts_with("N"), ends_with("L")) %>%
#'   group_by(ENV, GEN) %>%
#'   summarise_all(mean) %>%
#'   group_by(ENV) %>%
#'   resca(ends_with("L")) %>%
#'   head(n = 13)
#'
#'
resca <- function(.data = NULL, ..., values = NULL, new_min = 0, new_max = 100, keep = TRUE) {
  if(!missing(.data) && !missing(values)){
    stop("You cannot inform a vector of values if a data frame is used as input.")
  }
  if(!missing(.data)){
    rescc <- function(x){
      (new_max - new_min)/(max(x) - min(x)) * (x - max(x)) + new_max
    }
    if(is_grouped_df(.data)){
      dplyr::do(.data, resca(., ...))
    }
    if (keep == TRUE){
    .data %>% mutate_at(.vars = vars(...), .funs = list(res = rescc)) %>% ungroup()
    } else {
      .data %<>% mutate_at(.vars = vars(...), .funs = list(res = rescc))  %>% ungroup()
        return(suppressMessages(select(.data, contains("res"))))
    }
  } else {
    new_v <- function(v) {
      (new_max - new_min)/(max(values) - min(values)) * (v - max(values)) + new_max
    }
    sapply(values, new_v)
  }
}
```
