## Supplementary material for "metan: an R package for multi-environment trial analysis": available at: resende_indexes.R

```
#' Stability indexes based on a mixed-effect model
#'
#' This function computes the following indexes proposed by Resende (2007): the
#' harmonic mean of genotypic values (HMGV), the relative performance of the
#' genotypic values (RPGV) and the harmonic mean of the relative performance of
#' genotypic values (HMRPGV).
#'
#' The indexes computed with this function have been used to select genotypes
#' with stability performance in a mixed-effect model framework. Some examples
#' are in Alves et al (2018), Azevedo Peixoto et al. (2018), Dias et al. (2018)
#' and Colombari Filho et al. (2013).
#'
#' The HMGV index is computed as \deqn{HMGV_i = \frac{1}{E}\sum\limits_{j = 1}^E
#' {\frac{1}{{G{v_{ij}}}}}} where \eqn{E} is the number of environments included
#' in the analysis, \eqn{Gv_{ij}} is the genotypic value (BLUP) for the ith
#' genotype in the jth environment.
#'
#' The RPGV index is computed as \deqn{RPGV_i = \frac{1}{E}{\sum\limits_{j =
#' 1}^E {BLUP_{ij}} /\mathop \mu \nolimits_j }}
#'
#' The HMRPGV index is computed as \deqn{HMRPGV_i = \frac{1}{E}\sum\limits_{j =
#' 1}^E {\frac{1}{{G{v_{ij}}/{\mu _j}}}}}
#'
#' @param .data An object of class \code{waasb}
#' @return
#'
#' A dataframe containing the indexes.
#' @author Tiago Olivoto \email{tiagoolivoto@@gmail.com}
#' @references Alves, R.S., L. de Azevedo Peixoto, P.E. Teodoro, L.A. Silva,
#'   E.V. Rodrigues, M.D.V. de Resende, B.G. Laviola, and L.L. Bhering. 2018.
#'   Selection of Jatropha curcas families based on temporal stability and
#'   adaptability of genetic values. Ind. Crops Prod. 119:290-293.
#'   \href{https://www.sciencedirect.com/science/article/pii/S0926669018303406}{doi:10.1016/J.INDCROP.2018.04.029}
#'
#'
#'   Colombari Filho, J.M., M.D.V. de Resende, O.P. de Morais, A.P. de Castro,
#'   E.P. Guimaraes, J.A. Pereira, M.M. Utumi, and F. Breseghello. 2013. Upland
#'   rice breeding in Brazil: a simultaneous genotypic evaluation of stability,
#'   adaptability and grain yield. Euphytica 192:117-129.
#'   \href{https://link.springer.com/article/10.1007/s10681-013-0922-2}{doi:10.1007/s10681-013-0922-2}
#'
#'
#'   Dias, P.C., A. Xavier, M.D.V. de Resende, M.H.P. Barbosa, F.A. Biernaski,
#'   R.A. Estopa. 2018. Genetic evaluation of Pinus taeda clones from somatic
#'   embryogenesis and their genotype x environment interaction. Crop Breed.
#'   Appl. Biotechnol. 18:55-64.
#'   \href{http://www.scielo.br/scielo.php?script=sci_arttext&pid=S1984-70332018000100055&lng=en&tlng=en}{doi:10.1590/1984-70332018v18n1a8}
#'
#'
#'   Azevedo Peixoto, L. de, P.E. Teodoro, L.A. Silva, E.V. Rodrigues, B.G.
#'   Laviola, and L.L. Bhering. 2018. Jatropha half-sib family selection with
#'   high adaptability and genotypic stability. PLoS One 13:e0199880.
#'   \href{https://dx.plos.org/10.1371/journal.pone.0199880}{doi:10.1371/journal.pone.0199880}
#'
#'
#'   Resende MDV (2007) Matematica e estatistica na analise de experimentos e no
#'   melhoramento genetico. Embrapa Florestas, Colombo
#' @export
#' @examples
#' \donttest{
#' library(metan)
#' res_ind <- waasb(data_ge,
#'                  env = ENV,
#'                  gen = GEN,
#'                  rep = REP,
#'                  resp = c(GY, HM))
#' model_indexes <- Resende_indexes(res_ind)
#'
#' # Alternatively using the pipe operator %>%
#' res_ind <- data_ge %>%
#'            waasb(ENV, GEN, REP, c(GY, HM)) %>%
#'            Resende_indexes()
#'}
#'
Resende_indexes <- function(.data) {
    if (!is(.data, "waasb")) {
        stop("The object '.data' must be an object of class \"waasb\"")
    }
    listres <- list()
    for (var in 1:length(.data)) {
        gge <- .data[[var]][["BLUPgge"]]
        # Harmonic mean
        GEPRED <- make_mat(gge, GEN, ENV, Predicted)
        HMGV <- data.frame(HMGV = apply(GEPRED, 1, FUN = hm_mean),
                           HMGV_R = rank(-apply(GEPRED, 1, FUN = hm_mean)))
        ## Relative performance
        y <- .data[[var]][["MeansGxE"]]
        GEMEAN <- make_mat(y, GEN, ENV, Y)
        ovmean <- mean(y$Y)
        mean_env <- apply(GEMEAN, 2, FUN = mean)
        RPGV <- data.frame(RPGV = apply(t(t(GEPRED)/mean_env), 1, mean))
        RPGV_data <- RPGV %>%
            mutate(RPGV_Y = RPGV * ovmean,
                   RPGV_R = rank(-RPGV_Y))
        ## Harmonic mean of Relative performance
        HMRPGV <- data.frame(HMRPGV = apply(t(t(GEPRED)/mean_env), 1, hm_mean))
        HMRPGV_data <- HMRPGV %>%
            mutate(HMRPGV_Y = HMRPGV * ovmean,
                   HMRPGV_R = rank(-HMRPGV_Y))
        Y <- .data[[var]][["model"]] %>%
            dplyr::filter(type == "GEN") %>%
            select(Y)
        listres[[paste(names(.data[var]))]] <- cbind(Y, HMGV, RPGV_data, HMRPGV_data) %>%
            rownames_to_column("GEN") %>%
            as_tibble()
    }
    invisible(structure(listres, class = "Res_ind"))
}
```
