## Supplementary material for "metan: an R package for multi-environment trial analysis": available at: solve_svd.R

```
#' Pseudoinverse of a square matrix
#'
#' This function computes the Moore-Penrose pseudoinverse of a square matrix
#' using singular value decomposition.
#'
#' @param x A square matrix
#' @param tolerance The tolerance to consider an eigenvalue equals to zero.
#' @author Tiago Olivoto, \email{tiagoolivoto@@gmail.com}
#' @return A matrix with the same dimension of \code{x}.
#' @export
#' @examples
#' library(metan)
#' mat = matrix(c(1, 4, 2, 8), ncol = 2)
#' det(mat)
#' solve_svd(mat)

solve_svd <- function(x, tolerance = 2.220446e-16) {
  if (dim(x)[1] - dim(x)[2] != 0) {
    stop("The object 'x' must be a square matrix")
  }
  s <- svd(as.matrix(x))
  posi <- s$d > max(tolerance * s$d[1], 0)
  if (all(posi)) {
    xsol <- s$v %*% (1/s$d * t(s$u))
    rownames(xsol) <- colnames(xsol) <- colnames(x)
  } else if (!any(posi)) {
    array(0, dim(x)[2L:1L])
  } else {
    xsol <- s$v[, posi, drop = FALSE] %*% ((1/s$d[posi]) * t(s$u[, posi, drop = FALSE]))
    rownames(xsol) <- colnames(xsol) <- colnames(x)
  }
  return(xsol)
}
```
