## Supplementary material for "metan: an R package for multi-environment trial analysis": available at: split_factors.R

```
#' Split a data frame by factors
#'
#' Split a data frame into subsets grouping by one or more factors.
#'
#' This function is used to split a data frame into a list where each element is
#' a level of the grouping variable (or combination of grouping variables). By
#' combining the function \code{split_factors} with the forward-pipe operator
#' %>%, it is possible to apply some functions of the metan package to each
#' element of the list.
#'
#' @param .data The data that will be split. Must contain at least one grouping
#'   variable.
#' @param ... Comma-separated list of unquoted variable names that will be used
#'   to group the data.
#' @param keep_factors If more than two factors are in the dataframe, should the
#'   columns of the non-grouping factors be kept?
#' @param verbose Logical argument. If \code{verbose = FALSE} the code will run
#'   silently.
#' @return A list where each element is a named level of the grouping factors.
#'   If more than one grouping variable is used, then each element is the
#'   combination of the grouping variables.
#' @author Tiago Olivoto \email{tiagoolivoto@@gmail.com}
#' @export
#' @importFrom rlang  eval_bare  expr set_names
#' @examples
#' library(metan)
#'
#' g1 = split_factors(iris, Species)
#' g2 = split_factors(data_ge, ENV, keep_factors = TRUE)
#'
split_factors <- function(.data, ..., keep_factors = FALSE, verbose = TRUE) {
  grouped <- group_by(.data, ...)
  names <- eval_bare(expr(paste(!!!group_keys(grouped), sep = " | ")))
  gd <- grouped %>%
    group_split(keep = TRUE) %>%
    set_names(names)
  if (keep_factors == FALSE) {
    if (verbose == TRUE) {
      if (sum(lapply(gd[[1]], is.factor) == TRUE) > 0) {
        warning("Use 'keep_factors = TRUE' to keep the columns ", paste0(collapse = " ", names(gd[[1]][, unlist(lapply(gd[[1]], is.factor))])), " in the grouped data. ", call. = FALSE)
      }
    }
    gd <- lapply(gd, function(x) {
      x[, unlist(lapply(x, is.numeric))] %>% as_tibble()
    })
  } else {
    gd <- lapply(gd, function(x) {
      as_tibble(x)
    })
  }
  return(structure(list(dfs = gd, names = group_vars(grouped)),
                   class = "split_factors"))
}
```
