## Supplementary material for "metan: an R package for multi-environment trial analysis": available at: superiority.R

```
#' Lin e Binns' superiority index
#'
#' Nonparametric stability analysis using the superiority index proposed by Lin
#' & Binns, (1992).
#'
#'
#' @param .data The dataset containing the columns related to Environments,
#'   Genotypes, replication/block and response variable(s)
#' @param env The name of the column that contains the levels of the
#'   environments.
#' @param gen The name of the column that contains the levels of the genotypes.
#' @param rep The name of the column that contains the levels of the
#'   replications/blocks
#' @param resp The response variable(s). To analyze multiple variables in a
#'   single procedure use, for example, \code{resp = c(var1, var2, var3)}.
#' @param verbose Logical argument. If \code{verbose = FALSE} the code will run
#'   silently.
#' @return An object of class \code{superiority} where each element is the
#'   result of one variable and contains the following items:
#'
#' * \strong{environments} The mean for each environment, the environment index
#' and classification as favorable and unfavorable environments.
#' * \strong{index} The superiority index computed for all, favorable and
#' unfavorable environments.
#'
#' @md
#' @author Tiago Olivoto, \email{tiagoolivoto@@gmail.com}
#' @seealso \code{\link{Annicchiarico}, \link{ecovalence}, \link{ge_stats}}
#' @references Lin, C.S., and M.R. Binns. 1988. A superiority measure of
#'   cultivar performance for cultivar x location data. Can. J. Plant Sci.
#'   68:193-198.
#'   \href{http://pubs.aic.ca/doi/abs/10.4141/cjps88-018}{doi:10.4141/cjps88-018}
#'
#' @export
#' @examples
#'
#' library(metan)
#' out <- superiority(data_ge2,
#'                    env = ENV,
#'                    gen = GEN,
#'                    rep = REP,
#'                    resp = PH)
#'
#'
superiority <- function(.data, env, gen, rep, resp, verbose = TRUE) {
  factors  <- .data %>%
    select(ENV = {{env}},
           GEN = {{gen}},
           REP = {{rep}}) %>%
    mutate_all(as.factor)
  vars <- .data %>%
    select({{resp}}) %>%
    select_numeric_cols()
  listres <- list()
  nvar <- ncol(vars)
  for (var in 1:nvar) {
    data <- factors %>%
      mutate(mean = vars[[var]])
    ge_mean <- data %>% dplyr::group_by(ENV, GEN) %>% dplyr::summarise(mean = mean(mean))
    environments <- data %>% dplyr::group_by(ENV) %>% dplyr::summarise(Mean = mean(mean))
    environments <- mutate(environments, index = Mean - mean(environments$Mean),
                           class = ifelse(index < 0, "unfavorable", "favorable")) %>%
      as.data.frame()
    data <- suppressMessages(left_join(data, environments %>%
                                         select(ENV, class)))
    lin_fun <- function(mat) {
      P <- apply(mat, 1, function(x) {
        sum((x - apply(mat, 2, max))^2)/(2 * length(x))
      })
      return(P)
    }
    mat_g <- make_mat(data, row = GEN, col = ENV, value = mean)
    ge_mf <- subset(data, class == "favorable")
    mat_f <- dplyr::select_if(make_mat(ge_mf, row = GEN,
                                       col = ENV, value = mean), function(x) !any(is.na(x)))
    ge_mu <- subset(data, class == "unfavorable")
    mat_u <- dplyr::select_if(make_mat(ge_mu, row = GEN,
                                       col = ENV, value = mean), function(x) !any(is.na(x)))
    temp <- list(environments = tibble(environments),
                 index = tibble(GEN = rownames(mat_g),
                                Y = apply(mat_g, 1, mean),
                                Pi_a = lin_fun(mat_g),
                                R_a = rank(lin_fun(mat_g)),
                                Pi_f = lin_fun(mat_f),
                                R_f = rank(lin_fun(mat_f)),
                                Pi_u = lin_fun(mat_u),
                                R_u = rank(lin_fun(mat_u))))
    rownames(temp) <- NULL
    if (nvar > 1) {
      listres[[paste(names(vars[var]))]] <- temp
      if (verbose == TRUE) {
        cat("Evaluating variable", paste(names(vars[var])),
            round((var - 1)/(length(vars) - 1) * 100, 1), "%", "\n")
      }
    } else {
      listres[[paste(names(vars[var]))]] <- temp
    }
  }
  return(structure(listres, class = "superiority"))
}
```
