## Supplementary material for "metan: an R package for multi-environment trial analysis": available at: utilities.R

```
#' @title Utilities for handling with numbers and strings
#' @name utils_num_str
#' @param .data A data frame
#' @param ... The argument depends on the function used.
#' * For \code{round_cols()} \code{...} are the variables to round. If no
#' variable is informed, all the numeric variables from \code{data} are used.
#' * For \code{remove_strings()} \code{...} are the variables to remove the
#' strings. If no variable is informed, the strings of all non-numeric variables
#' will be removed, keeping the numbers. If variables contains only strings,
#' they will be replaced with \code{NA}.
#' @param digits The number of significant figures.
#' @param var The variable to extract or replace numbers or strings.
#' @param new_var The name of the new variable containing the numbers or
#'   strings extracted or replaced. Defaults to \code{new_var}.
#' @param drop Logical argument. If \code{TRUE} keeps the new variable
#'   \code{new_var} and drops the existing ones. Defaults to \code{FALSE}.
#' @param pattern A string to be matched. Regular Expression Syntax is also
#'   allowed.
#' @param replacement A string for replacement.
#' @param pull Logical argument. If \code{TRUE}, returns the last column (on the
#'   assumption that's the column you've created most recently), as a vector.
#' @param .before,.after For \code{replace_sting()}, \code{replace_number()},
#'   \code{extract_string()}, ,and  \code{extract_number()} one-based column
#'   index or column name where to add the new columns.
#' @description
#' * \code{all_lower_case()}: Translate all non-numeric strings of a data frame
#' to lower case.
#' * \code{all_upper_case()}: Translate all non-numeric strings of a data frame
#' to upper case.
#' * \code{extract_number()}: Extract the number(s) of a string.
#' * \code{extract_string()}: Extract all strings, ignoring case.
#' * \code{remove_strings()}: Remove all strings of a variable.
#' * \code{replace_number()}: Replace numbers with a replacement.
#' * \code{replace_string()}: Replace all strings with a replacement, ignoring
#' case.
#' * \code{round_cols()}: Round a selected column or a whole data frame to
#' significant figures.
#' @md
#' @examples
#' \donttest{
#' library(metan)
#'
#' ################ Rounding numbers ###############
#' # All numeric columns
#' round_cols(data_ge2, digits = 1)
#'
#' # Round specific columns
#' round_cols(data_ge2, EP, digits = 1)
#'
#' ########### Extract or replace numbers ##########
#' # Extract numbers
#' extract_number(data_ge, GEN)
#' extract_number(data_ge,
#'                var = GEN,
#'                drop = TRUE,
#'                new_var = g_number)
#'
#' # Replace numbers
#'
#' replace_number(data_ge, GEN)
#' replace_number(data_ge,
#'                var = GEN,
#'                pattern = "1",
#'                replacement = "_one",
#'                pull = TRUE)
#'
#' ########## Extract, replace or remove strings ##########
#' # Extract strings
#' extract_string(data_ge, GEN)
#' extract_string(data_ge,
#'                var = GEN,
#'                drop = TRUE,
#'                new_var = g_name)
#'
#' # Replace strings
#' replace_string(data_ge, GEN)
#' replace_string(data_ge,
#'                var = GEN,
#'                new_var = GENOTYPE,
#'                pattern = "G",
#'                replacement = "GENOTYPE_")
#' # Remove strings
#' remove_strings(data_ge)
#' remove_strings(data_ge, ENV)
#' ############# upper and lower cases ############
#' all_lower_case("GENOTYPE")
#' lc <- all_lower_case(data_ge)
#' lc
#' all_lower_case("GENOTYPE")
#'
#' all_upper_case("Genotype")
#' all_upper_case(lc)
#' }
#' @export
all_upper_case <- function(.data){
  if (any(class(.data) %in% c("data.frame","tbl_df", "data.table"))){
    .data <- as.data.frame(.data)
    mutate_if(.data, ~!is.numeric(.x), toupper) %>%
      as_tibble(rownames = NA)
  } else{
    toupper(.data)
  }
}
#' @name utils_num_str
#' @export
all_lower_case <- function(.data){
  if (any(class(.data) %in% c("data.frame","tbl_df", "data.table"))){
    .data <- as.data.frame(.data)
    mutate_if(.data, ~!is.numeric(.x), tolower) %>%
      as_tibble(rownames = NA)
  } else{
    tolower(.data)
  }
}
#' @name utils_num_str
#' @export
extract_number <- function(.data,
                           var,
                           new_var = new_var,
                           drop = FALSE,
                           pull = FALSE,
                           .before = NULL,
                           .after  = NULL){
  if (drop == FALSE){
    results <- .data %>%
      mutate({{new_var}} :=   as.numeric(gsub("[^0-9.-]+", "", as.character({{var}}))))
    if (pull == TRUE){
      results <- pull(results)
    }
    if (!is.null(.before) | !is.null(.after)){
      results <- reorder_cols(results, {{new_var}}, .before = .before, .after = .after)
    }
  } else{
    results <- .data %>%
      transmute({{new_var}} :=   as.numeric(gsub("[^0-9.-]+", "", as.character({{var}}))))
    if(pull == TRUE){
      results <- pull(results)
    }
  }
  return(results)
}
#' @name utils_num_str
#' @export
extract_string <- function(.data,
                           var,
                           new_var = new_var,
                           drop = FALSE,
                           pull = FALSE,
                           .before = NULL,
                           .after  = NULL){
  if (drop == FALSE){
    results <- .data %>%
      mutate({{new_var}} := as.character(gsub("[^A-z.-]+", "", as.character({{var}}))))
    if(pull == TRUE){
      results <- pull(results)
    }
    if (!is.null(.before) | !is.null(.after)){
      results <- reorder_cols(results, {{new_var}}, .before = .before, .after = .after)
    }
  } else{
    results <- .data %>%
      transmute({{new_var}} := as.character(gsub("[^A-z.-]+", "", as.character({{var}}))))
    if(pull == TRUE){
      results <- pull(results)
    }
  }
  return(results)
}
#' @name utils_num_str
#' @export
remove_strings <- function(.data, ...){
  if(missing(...)){
    vars <- vars(everything())
  } else{
    vars <- vars(...)
  }
  results <-
    mutate_at(.data,
              .vars = vars,
              .funs = gsub,
              pattern = "[A-z]",
              replacement = "") %>%
    mutate_at(.vars = vars,
              .funs = as.numeric)
  return(results)
}
#' @name utils_num_str
#' @export
replace_number <- function(.data,
                           var,
                           new_var = new_var,
                           pattern = NULL,
                           replacement = "",
                           drop = FALSE,
                           pull = FALSE,
                           .before = NULL,
                           .after  = NULL){
  if(missing(pattern)){
    pattern <- "[0-9]"
  } else{
    pattern <- pattern
  }
  if (drop == FALSE){
    results <- .data %>%
      mutate({{new_var}} := gsub(pattern, replacement, as.character({{var}})))
    if(pull == TRUE){
      results <- pull(results)
    }
    if (!is.null(.before) | !is.null(.after)){
      results <- reorder_cols(results, {{new_var}}, .before = .before, .after = .after)
    }
  } else{
    results <- .data %>%
      transmute({{new_var}} := gsub(pattern, replacement, as.character({{var}})))
    if(pull == TRUE){
      results <- pull(results)
    }
  }
  return(results)
}
#' @name utils_num_str
#' @export
replace_string <- function(.data,
                           var,
                           new_var = new_var,
                           pattern = NULL,
                           replacement = "",
                           drop = FALSE,
                           pull = FALSE,
                           .before = NULL,
                           .after  = NULL){
  if(missing(pattern)){
    pattern <- "[A-z]"
  } else {
    pattern <- pattern
  }
  if (drop == FALSE){
    results <- .data %>%
      mutate({{new_var}} := gsub(pattern, replacement, as.character({{var}})))
    if(pull == TRUE){
      results <- pull(results)
    }
    if (!is.null(.before) | !is.null(.after)){
      results <- reorder_cols(results, {{new_var}}, .before = .before, .after = .after)
    }
  } else{
    results <- .data %>%
      transmute({{new_var}} := gsub(pattern, replacement, as.character({{var}})))
    if(pull == TRUE){
      results <- pull(results)
    }
  }
  return(results)
}
#' @name utils_num_str
#' @export
#' @importFrom dplyr mutate_if transmute
round_cols <- function(.data, ...,  digits = 2){
  rn_test <- has_rownames(.data)
  if(rn_test == TRUE){
    rnames <- rownames(.data)
  }
  if (missing(...)){
    .data %<>% mutate_if(is.numeric, round, digits = digits)
  } else{
    .data %<>% mutate_at(vars(...), round, digits = digits)
  }
  if(rn_test == TRUE){
    rownames(.data) <- rnames
  }
  return(.data)
}
#' @title Utilities for handling with rows and columns
#' @name utils_rows_cols
#' @description
#' * \code{add_cols()}: Add one or more columns to an existing data frame. If
#' specified \code{.before} or \code{.after} columns does not exist, columns are
#' appended at the end of the data. Return a data frame with all the original
#' columns in \code{.data} plus the columns declared in \code{...}. In
#' \code{add_cols()} columns in \code{.data} are available for the expressions.
#' So, it is possible to add a column based on existing data.
#' * \code{add_rows()}: Add one or more rows to an existing data frame. If
#' specified \code{.before} or \code{.after} rows does not exist, rows are
#' appended at the end of the data. Return a data frame with all the original
#' rows in \code{.data} plus the rows declared in \code{...}.
#' * \code{all_pairs()}: Get all the possible pairs between the levels of a
#' factor.
#' * \code{column_exists()}: Checks if a column exists in a data frame. Return a
#' logical value.
#' * \code{columns_to_first()}: Move columns to first positions in \code{.data}.
#' * \code{columns_to_last()}: Move columns to last positions in \code{.data}.
#' * \code{concatenate()}: Concatenate columns of a data frame. If \code{drop =
#' TRUE} then the existing variables are dropped.
#' * \code{get_levels()}: Get the levels of a factor variable.
#' * \code{get_level_size()}: Get the size of each level of a factor variable.
#' * \code{remove_cols()}: Remove one or more columns from a data frame.
#' * \code{remove_rows()}: Remove one or more rows from a data frame.
#' * \code{reorder_cols()}: Reorder columns in a data frame.
#' * \code{select_cols()}: Select one or more columns from a data frame.
#' * \code{select_first_col()}: Select first variable, possibly with an offset.
#' * \code{select_last_col()}: Select last variable, possibly with an offset.
#' * \code{select_numeric_cols()}: Select all the numeric columns of a data
#' frame.
#' * \code{select_non_numeric_cols()}: Select all the non-numeric columns of a
#' data frame.
#' * \code{select_rows()}: Select one or more rows from a data frame.
#'
#' @param .data A data frame
#' @param ... The argument depends on the function used.
#' * For \code{add_cols()} and \code{add_rows()} is name-value pairs. All values
#' must have one element for each row in \code{.data} when using
#' \code{add_cols()} or one element for each column in \code{.data} when using
#' \code{add_rows()}. Values of length 1 will be recycled when using
#' \code{add_cols()}.
#'
#' * For \code{remove_cols()} and \code{select_cols()},  \code{...} is the
#' column name or column index of the variable(s) to be dropped.
#'
#' * For \code{columns_to_first()} and \code{columns_to_last()},  \code{...} is
#' the column name or column index of the variable(s) to be moved to first or
#' last in \code{.data}.
#'
#' * For \code{remove_rows()} and \code{select_rows()}, \code{...} is an integer
#' row value.
#'
#' * For \code{concatenate()}, \code{...} is the unquoted variable names to be
#' concatenated.
#' @param .before,.after For \code{add_cols()}, \code{concatenate()}, and
#'   \code{reorder_cols()}, one-based column index or column name where to add
#'   the new columns, default: .after last column. For \code{add_rows()},
#'   one-based row index where to add the new rows, default: .after last row.
#' @param new_var The name of the new variable containing the concatenated
#'   values. Defaults to \code{new_var}.
#' @param sep The separator to appear between concatenated variables. Defaults
#'   to "_".
#' @param drop Logical argument. If \code{TRUE} keeps the new variable
#'   \code{new_var} and drops the existing ones. Defaults to \code{FALSE}.
#' @param pull Logical argument. If \code{TRUE}, returns the last column (on the
#'   assumption that's the column you've created most recently), as a vector.
#' @param cols A quoted variable name to check if it exists in \code{.data}.
#' @param group A factor variable to get the levels.
#' @param levels The levels of a factor or a numeric vector.
#' @param offset Set it to \emph{n} to select the \emph{n}th variable from the
#'   end (for \code{select_last_col()}) of from the begin (for
#'   \code{select_first_col()})
#' @md
#' @importFrom  tibble add_column add_row
#' @export
#' @examples
#' \donttest{
#' library(metan)
#'
#' ################# Adding columns #################
#' # Variables x and y .after last column
#' data_ge %>%
#'   add_cols(x = 10,
#'            y = 30)
#' # Variables x and y .before the variable GEN
#' data_ge %>%
#'   add_cols(x = 10,
#'            y = 30,
#'            .before = "GEN")
#'
#' # Creating a new variable based on the existing ones.
#' data_ge %>%
#'   add_cols(GY2 = GY^2,
#'            GY2_HM = GY2 + HM,
#'            .after = "GY")
#'
#' ############### Reordering columns ###############
#' reorder_cols(data_ge2, NKR, .before = "ENV")
#' reorder_cols(data_ge2, ENV, GEN, .after = "ED")
#'
#' ######## Selecting and removing columns ##########
#' select_cols(data_ge2, GEN, REP)
#' remove_cols(data_ge2, GEN, REP)
#'
#' ########## Selecting and removing rows ###########
#' select_rows(data_ge2, 2:3)
#' remove_rows(data_ge2, 2:3)
#'
#' ########### Concatenating columns ################
#' concatenate(data_ge, ENV, GEN, REP)
#' concatenate(data_ge, ENV, GEN, REP, drop = TRUE)
#'
#' # Combine with add_cols() and replace_string()
#'data_ge2 %>%
#'  add_cols(ENV_GEN = concatenate(., ENV, GEN, pull = TRUE),
#'           .after = "GEN") %>%
#'  replace_string(ENV_GEN,
#'                 pattern = "H",
#'                 replacement = "HYB_",
#'                 .after = "ENV_GEN")
#'
#'
#' ################### Adding rows ##################
#' data_ge %>%
#'   add_rows(ENV = "E_TEST",
#'            GEN = "G_TEST",
#'            REP = 3,
#'            GY = 10.3,
#'            HM = 100.11,
#'            .after = 1)
#' ########## checking if a column exists ###########
#' column_exists(data_g, "GEN")
#'
#' ####### get the levels and size of levels ########
#' get_levels(data_g, GEN)
#' get_level_size(data_g, GEN)
#'
#' ############## all possible pairs ################
#' all_pairs(data_g, GEN)
#'
#' ########## select numeric variables only #########
#' select_numeric_cols(data_g)
#' select_non_numeric_cols(data_g)
#' }
add_cols <- function(.data, ..., .before = NULL, .after = NULL){
  results <- mutate(.data, ...)
  pos_dots <- (ncol(results)) - ((ncol(results) - ncol(.data)) - 1)
  if (!is.null(.before) | !is.null(.after)){
    results <- reorder_cols(results,
                            pos_dots:ncol(results),
                            .before = .before,
                            .after = .after)
  }
  return(as_tibble(results, rownames = NA))
}
#' @name utils_rows_cols
#' @export
add_rows <- function(.data, ..., .before = NULL, .after = NULL){
  if(is.character(.before)){
    if(!(.before %in% rownames(.data))){
      .before <- NULL
    }
  }
  if(is.character(.after)){
    if(!(.after %in% rownames(.data))){
      .after <- NULL
    }
  }
  add_row(.data, ..., .before = .before, .after = .after)
}
#' @name utils_rows_cols
#' @export
all_pairs <- function(.data, levels){
  levels <-
    get_levels(.data, {{levels}})
  combn(levels, 2) %>%
    t() %>%
    as.data.frame()
}
#' @name utils_rows_cols
#' @export
column_to_first <-function(.data, ...){
  select_cols(.data, ..., everything())
}
#' @name utils_rows_cols
#' @export
column_to_last <-function(.data, ...){
  select_cols(.data, -c(!!!quos(...)), everything())
}
#' @name utils_rows_cols
#' @export
column_exists <-function(.data, cols){
  if(length(setdiff(cols, colnames(.data))) != 0){
    FALSE
  } else{
    TRUE
  }
}
#' @name utils_rows_cols
#' @export
concatenate <- function(.data,
                        ...,
                        new_var = new_var,
                        sep = "_",
                        drop = FALSE,
                        pull = FALSE,
                        .before = NULL,
                        .after  = NULL){
  if (drop == FALSE){
    conc <- select(.data, ...)
    results <- mutate(.data,
                      {{new_var}} := apply(conc, 1, paste, collapse = sep))
    if (pull == TRUE){
      results <- pull(results)
    }
    if (!is.null(.before) | !is.null(.after)){
      results <- reorder_cols(results, {{new_var}}, .before = .before, .after = .after)
    }
  } else{
    conc <- select(.data, ...)
    results <- transmute(.data,
                         {{new_var}} := apply(conc, 1, paste, collapse = sep))
    if (pull == TRUE){
      results <- pull(results)
    }
  }
  return(results)
}
#' @name utils_rows_cols
#' @export
get_levels <- function(.data, group){
  .data %>%
    mutate_if(~!is.numeric(.x), as.factor) %>%
    pull({{group}}) %>%
    levels()
}
#' @name utils_rows_cols
#' @importFrom dplyr count
#' @export
get_level_size <- function(.data, group){
  result <- .data %>%
    group_by({{group}}) %>%
    count()
  n <- result$n
  names(n) <- result %>% pull(1)
  return(n)
}
#' @name utils_rows_cols
#' @importFrom rlang quos
#' @export
reorder_cols <- function(.data, ..., .before = NULL, .after = NULL){
  args <- match.call()
  if (is.null(.after) && is.null(.before)){
    stop("At least '.before' or '.after' must be informed", call. = FALSE)
  }
  if (!is.null(.after) && !is.null(.before)){
    stop("'.before' and '.after' cannot be evaluated together.", call. = FALSE)
  }
  if (!is.null(.before)){
    if(is.character(.before)){
      if(!(.before %in% colnames(.data))){
        stop(paste("Column", args[[".before"]], "not present in .data"), call. = FALSE)
      }
    } else{
      .before <- colnames(.data[.before])
    }
    sel <- select(.data, ...)
    bfr <- .data[,1:which(colnames(.data) ==  .before)-1]
    if(any(colnames(sel) %in% colnames(bfr))){
      bfr <- bfr[, -which(names(bfr)  %in% names(sel))]
    }
    aft <- select(.data, -!!colnames(bfr), -!!colnames(sel))
    results <- cbind(bfr, sel, aft)
  }
  if(!is.null(.after)) {
    if(is.character(.after)){
      if(!(.after %in% colnames(.data))){
        stop(paste("Column", args[[".after"]], "not present in .data"), call. = FALSE)
      }
    } else{
      .after <- colnames(.data[.after])
    }
    if(which(colnames(.data) ==  .after)+1 > ncol(.data)){
      results <- select(.data, -c(!!!quos(...)), everything())
    } else{
      sel <- select(.data, ...)
      aft <- .data[,(which(colnames(.data) ==  .after)+1):ncol(.data)]
      if(any(colnames(sel) %in% colnames(aft))){
        aft <- aft[, -which(names(aft)  %in% names(sel))]
      }
      bfr <- select(.data, -!!colnames(aft), -!!colnames(sel))
      results <- cbind(bfr, sel, aft)
    }
  }
  return(as_tibble(results))
}
#' @name utils_rows_cols
#' @export
remove_cols <- function(.data, ...){
  select(.data, -c(!!!quos(...)))
}
#' @name utils_rows_cols
#' @export
remove_rows <- function(.data, ...){
  slice(.data, -c(!!!quos(...)))
}
#' @name utils_rows_cols
#' @export
select_first_col <- function(.data, offset = NULL){
  if(!missing(offset) && offset == 0){
    stop("`offset` must be greater than the 0", call. = FALSE)
  }
  offset <- ifelse(missing(offset), ncol(.data) - 1, ncol(.data) - offset)
  select(.data, last_col(offset = offset))
}
#' @name utils_rows_cols
#' @export
select_last_col <- function(.data, offset = NULL){
  offset <- ifelse(missing(offset), 0, offset)
  select(.data, last_col(offset = offset))
}
#' @name utils_rows_cols
#' @export
select_numeric_cols <- function(.data){
  if(is_grouped_df(.data)){
    .data <- ungroup(.data)
  }
  select_if(.data, is.numeric)
}
#' @name utils_rows_cols
#' @export
select_non_numeric_cols <- function(.data){
  if(is_grouped_df(.data)){
    .data <- ungroup(.data)
  }
  select_if(.data, ~!is.numeric(.x))
}
#' @name utils_rows_cols
#' @export
select_cols <- function(.data, ...){
  select(.data, ...)
}
#' @name utils_rows_cols
#' @export
select_rows <- function(.data, ...){
  slice(.data, ...)
}


#' @title Means by one or more factors
#' @description Computes the mean for all numeric variables of a data frame,
#'   grouping by one or more factors.
#' @param .data A data frame
#' @param ... One or more categorical variables for grouping the data.
#' @return An object of class tbl_df with the computed means by each level of
#'   the factor(s) declared in \code{...}.
#' @export
#' @examples
#' \donttest{
#' library(metan)
#' means_by(data_ge2, ENV)
#' means_by(data_ge2, GEN, ENV)
#'}
means_by <- function(.data, ...){
  group_by(.data, ...) %>%
    summarise_if(is.numeric, mean) %>%
    ungroup()
}
```
