## Supplementary material for "metan: an R package for multi-environment trial analysis": available at: waas.R

```
#' Weighted Average of Absolute Scores
#'
#' Compute the Weighted Average of Absolute Scores for AMMI analysis (Olivoto et
#' al., 2019).
#'
#' This function compute the weighted average of absolute scores, estimated as
#' follows:
#'
#' \deqn{ WAAS_i = \sum_{k = 1}^{p} |IPCA_{ik} \times EP_k|/ \sum_{k =
#' 1}^{p}EP_k}
#'
#' where \eqn{WAAS_i} is the weighted average of absolute scores of the
#' \emph{i}th genotype; \eqn{PCA_{ik}} is the score of the \emph{i}th genotype
#' in the \emph{k}th IPCA; and \eqn{EP_k} is the explained variance of the *k*th
#' IPCA for \emph{k = 1,2,..,p}, considering \emph{p} the number of significant
#' PCAs, or a declared number of PCAs. For example if \code{prob = 0.05}, all
#' axis that are significant considering this probability level are used. The
#' number of axis can be also informed by declaring \code{naxis = x}. This will
#' override the number of significant axes according to the argument code{prob}.
#'
#' @param .data The dataset containing the columns related to Environments,
#'   Genotypes, replication/block and response variable(s).
#' @param env The name of the column that contains the levels of the
#'   environments.
#' @param gen The name of the column that contains the levels of the genotypes.
#' @param rep The name of the column that contains the levels of the
#'   replications/blocks.
#' @param resp The response variable(s). To analyze multiple variables in a
#'   single procedure a vector of variables may be used. For example \code{resp
#'   = c(var1, var2, var3)}.
#' @param mresp A numeric vector of the same length of \code{resp}. The
#'   \code{mresp} will be the new maximum value after rescaling. By default, all
#'   variables in \code{resp} are rescaled so that de maximum value is 100 and
#'   the minimum value is 0.
#' @param wresp The weight for the response variable(s) for computing the WAASBY
#'   index. Must be a numeric vector of the same length of \code{resp}. Defaults
#'   to 50, i.e., equal weights for stability and mean performance.
#' @param prob The p-value for considering an interaction principal component
#'   axis significant.
#' @param naxis The number of IPCAs to be used for computing the WAAS index.
#'   Default is \code{NULL} (Significant IPCAs are used). If values are
#'   informed, the number of IPCAS will be used independently on its
#'   significance. Note that if two or more variables are included in
#'   \code{resp}, then \code{naxis} must be a vector.
#' @param ind_anova Logical argument set to \code{TRUE}. If \code{FALSE} the
#'   within-environment ANOVA is not performed.
#' @param verbose Logical argument. If \code{verbose = FALSE} the code is run
#'   silently.
#' @references Olivoto, T., A.D.C. L{\'{u}}cio, J.A.G. da silva, V.S. Marchioro,
#'   V.Q. de Souza, and E. Jost. 2019a. Mean performance and stability in
#'   multi-environment trials I: Combining features of AMMI and BLUP techniques.
#'   Agron. J. 111:2949-2960.
#'   \href{https://dl.sciencesocieties.org/publications/aj/abstracts/0/0/agronj2019.03.0220?access=0&view=pdf}{doi:10.2134/agronj2019.03.0220}
#'
#' @return An object of class \code{waas} with the following items for each
#'   variable:
#'
#' * \strong{individual} A within-environments ANOVA considering a fixed-effect
#' model.
#' * \strong{model} A data frame with the response variable, the scores of all
#' Principal Components, the estimates of Weighted Average of Absolute Scores,
#' and WAASY (the index that consider the weights for stability and productivity
#' in the genotype ranking.
#'
#' * \strong{MeansGxE} The means of genotypes in the environments
#'
#' * \strong{PCA} Principal Component Analysis.
#'
#' * \strong{anova} Joint analysis of variance for the main effects and
#' Principal Component analysis of the interaction effect.
#'
#' * \strong{Details} A list summarizing the results. The following information
#' are showed. \code{WgtResponse}, the weight for the response variable in
#' estimating WAASB, \code{WgtWAAS} the weight for stability, \code{Ngen} the
#' number of genotypes, \code{Nenv} the number of environments, \code{OVmean}
#' the overall mean, \code{Min} the minimum observed (returning the genotype and
#' environment), \code{Max} the maximum observed, \code{Max} the maximum
#' observed, \code{MinENV} the environment with the lower mean, \code{MaxENV}
#' the environment with the larger mean observed, \code{MinGEN} the genotype
#' with the lower mean, \code{MaxGEN} the genotype with the larger.
#' * \strong{residuals} The residuals of the model.
#' * \strong{probint} The p-value for the genotype-vs-environment interaction.
#' @md
#' @author Tiago Olivoto \email{tiagoolivoto@@gmail.com}
#' @seealso \code{\link{waasb}}
#' @export
#' @examples
#'\donttest{
#' library(metan)
#'
#' # Considering p-value <= 0.05 to compute the WAAS
#'
#'model <- waas(data_ge,
#'              env = ENV,
#'              gen = GEN,
#'              rep = REP,
#'              resp = GY)
#'
#'
#' # Declaring the number of axis to be used for computing WAAS
#' # and assigning a larger weight for the response variable when
#' # computing the WAASBY index.
#'
#' model2 <- waas(data_ge,
#'                env = ENV,
#'                gen = GEN,
#'                rep = REP,
#'                resp = GY,
#'                naxis = 3,
#'                wresp = 60)
#'
#' # Analyzing multiple variables (GY and HM) at the same time
#' # considering that smaller values of HM are better and higher
#' # values of GY are better, assigning a larger weight for the GY
#' # and a smaller weight for HM when computing WAASBY index.
#'
#' model3 <- waas(data_ge,
#'                env = ENV,
#'                gen = GEN,
#'                rep = REP,
#'                resp = c(GY, HM),
#'                mresp = c(100, 0),
#'                wresp = c(60, 40))
#'}
#'
waas <- function(.data, env, gen, rep, resp, mresp = NULL, wresp = NULL, prob = 0.05,
                 naxis = NULL, ind_anova = TRUE, verbose = TRUE) {
    factors  <- .data %>%
        select(ENV = {{env}},
               GEN = {{gen}},
               REP = {{rep}}) %>%
        mutate_all(as.factor)
    vars <- .data %>%
        select({{resp}}) %>%
        select_numeric_cols()
    nvar <- ncol(vars)
    if (!is.null(naxis)) {
        if (length(naxis) != nvar) {
            stop("The argument 'naxis' must have length ", nvar, ", the same number of variables in 'resp'.")
        }
    }
    if (is.null(mresp)) {
        mresp <- replicate(nvar, 100)
        minresp <- 100 - mresp
    } else {
        if (length(mresp) != nvar) {
            stop("The length of the numeric vector 'mresp' must be equal the number of variables in argument 'resp'")
        }
        if (sum(mresp == 100) + sum(mresp == 0) != nvar) {
            stop("The values of the numeric vector 'mresp' must be 0 or 100.")
        }
        mresp <- mresp
        minresp <- 100 - mresp
    }
    if (is.null(wresp)) {
        PesoResp <- replicate(nvar, 50)
        PesoWAASB <- 100 - PesoResp
    } else {
        if (length(wresp) != nvar) {
            stop("The length of the numeric vector 'wresp' must be equal the number of variables in argument 'resp'")
        }
        if (min(wresp) < 0 | max(wresp) > 100) {
            stop("The range of the numeric vector 'wresp' must be equal between 0 and 100.")
        }
        PesoResp <- wresp
        PesoWAASB <- 100 - PesoResp
    }
    listres <- list()
    vin <- 0
    for (var in 1:nvar) {
        data <- factors %>%
            mutate(Y = vars[[var]])
        Nenv <- length(unique(data$ENV))
        Ngen <- length(unique(data$GEN))
        minimo <- min(Nenv, Ngen) - 1
        vin <- vin + 1
        if(ind_anova == TRUE){
            individual <- data %>% anova_ind(ENV, GEN, REP, Y)
        } else{
            individual = NULL
        }
        model <- performs_ammi(data, ENV, GEN, REP, Y, verbose = FALSE)[[1]]
        PC <- model$PCA
        Escores <- model$model
        MeansGxE <- model$MeansGxE
        if (is.null(naxis)) {
            SigPC1 <- nrow(PC[which(PC[, 6] < prob), ])
        } else {
            if (nvar == 1) {
                SigPC1 <- naxis
            } else {
                SigPC1 <- naxis[vin]
            }
        }
        if (SigPC1 > minimo) {
            stop("The number of axis to be used must be lesser than or equal to ",
                 minimo, " [min(GEN-1;ENV-1)]")
        } else {
            Pesos <- slice(model$PCA[7], 1:SigPC1)
            WAAS <- Escores %>%
                select(contains("PC")) %>%
                abs() %>%
                t() %>%
                as.data.frame() %>%
                slice(1:SigPC1) %>%
                mutate(Percent = Pesos$Percent)
            WAASAbs <- mutate(Escores, WAAS = sapply(WAAS[, -ncol(WAAS)], weighted.mean, w = WAAS$Percent))
            if (nvar > 1) {
                WAASAbs %<>%
                    group_by(type) %>%
                    mutate(PctResp = (mresp[vin] - minresp[vin])/(max(Y) - min(Y)) * (Y - max(Y)) + mresp[vin],
                           PctWAAS = (0 - 100)/(max(WAAS) - min(WAAS)) * (WAAS - max(WAAS)) + 0,
                           wRes = PesoResp[vin],
                           wWAAS = PesoWAASB[vin],
                           OrResp = rank(-Y),
                           OrWAAS = rank(WAAS),
                           OrPC1 = rank(abs(PC1)),
                           WAASY = ((PctResp * wRes) + (PctWAAS * wWAAS))/(wRes + wWAAS),
                           OrWAASY = rank(-WAASY)) %>%
                    ungroup()
            } else {
                WAASAbs %<>%
                    group_by(type) %>%
                    mutate(PctResp = (mresp - minresp)/(max(Y) - min(Y)) * (Y - max(Y)) + mresp,
                           PctWAAS = (0 - 100)/(max(WAAS) - min(WAAS)) * (WAAS - max(WAAS)) + 0,
                           wRes = PesoResp,
                           wWAAS = PesoWAASB,
                           OrResp = rank(-Y),
                           OrWAAS = rank(WAAS),
                           OrPC1 = rank(abs(PC1)),
                           WAASY = ((PctResp * wRes) + (PctWAAS * wWAAS))/(wRes + wWAAS),
                           OrWAASY = rank(-WAASY)) %>%
                    ungroup()
            }
            min_group <- Escores %>% group_by(type) %>% top_n(1, -Y) %>% select(type, Code, Y) %>% slice(1) %>% as.data.frame()
            max_group <- Escores %>% group_by(type) %>% top_n(1, Y) %>% select(type, Code, Y) %>% slice(1) %>% as.data.frame()
            min <- MeansGxE %>% top_n(1, -Y) %>% select(ENV, GEN, Y) %>% slice(1)
            max <- MeansGxE %>% top_n(1, Y) %>% select(ENV, GEN, Y) %>% slice(1)
            Details <- tibble(Parameters = c("Ngen", "Nenv", "OVmean","Min", "Max", "MinENV", "MaxENV", "MinGEN", "MaxGEN", "SigPC"),
                              Values = c(Ngen, Nenv, round(mean(MeansGxE$Y), 4),
                                         paste0(round(min[3], 4), " (", min$GEN, " in ", min$ENV,")"),
                                         paste0(round(max$Y, 4), " (", max$GEN, " in ", max$ENV,")"),
                                         paste0(min_group[1,2], " (", round(min_group[1,3], 3),")"),
                                         paste0(max_group[1,2], " (", round(max_group[1,3], 3),")"),
                                         paste0(min_group[2,2], " (", round(min_group[2,3], 3), ") "),
                                         paste0(max_group[2,2], " (", round(max_group[2,3], 3), ") "),
                                         SigPC1))
            listres[[paste(names(vars[var]))]] <-
                structure(list(individual = individual[[1]],
                               model = WAASAbs,
                               MeansGxE = MeansGxE,
                               PCA = as_tibble(PC, rownames = NA),
                               anova = model$ANOVA,
                               Details = Details,
                               residuals = as_tibble(model$residuals, rownames = NA),
                               probint = model$probint),
                          class = "waas")

            if (verbose == TRUE) {
                cat("variable", paste(names(vars[var])),"\n")
                cat("---------------------------------------------------------------------------\n")
                cat("AMMI analysis table\n")
                cat("---------------------------------------------------------------------------\n")
                print(as.data.frame(model$ANOVA), digits = 3, row.names = FALSE)
                cat("---------------------------------------------------------------------------\n\n")
            }
        }
    }
    if (verbose == TRUE) {
        if (length(which(unlist(lapply(listres, function(x) {
            x[["probint"]]
        })) > prob)) > 0) {
            cat("------------------------------------------------------------\n")
            cat("Variables with nonsignificant GxE interaction\n")
            cat(names(which(unlist(lapply(listres, function(x) {
                x[["probint"]]
            })) > prob)), "\n")
            cat("------------------------------------------------------------\n")
        } else {
            cat("All variables with significant (p < 0.05) genotype-vs-environment interaction\n")
        }
        cat("Done!\n")
    }
    invisible(structure(listres, class = "waas"))
}
```
