## Supplementary material for "metan: an R package for multi-environment trial analysis": available at: wsmp.R

```
#' Weighting between stability and mean performance
#'
#' This function computes the WAASY or WAASBY indexes (Olivoto et al., 2019) considering different
#' scenarios of weights for stability and mean performance.
#'
#' After fitting a model with the functions \code{\link{waas}} or \code{\link{waasb}}
#' it is possible to compute the superiority indexes WAASY or WAASBY in different scenarios of
#' weights for stability and mean performance. The number of scenarios is defined by the arguments
#' \code{increment}. By default, twenty-one different scenarios are computed. In this case, the
#' the superiority index is computed considering the following weights: stability (waasb or waas) = 100;
#' mean performance = 0. In other words, only stability is considered for genotype ranking. In the
#' next iteration, the weights becomes 95/5 (since increment = 5). In the third scenario, the weights become
#' 90/10, and so on up to these weights become 0/100. In the last iteration, the genotype
#' ranking for WAASY or WAASBY matches perfectly with the ranks of the response variable.
#'
#' @param model Should be an object of class \code{waas} or \code{waasb}.
#' @param mresp A numeric value that will be the new maximum value after rescaling.
#' By default, the variable in \code{resp} is rescaled so that the original maximum
#' and minimum values are 100 and 0, respectively. Let us consider that for a specific
#' trait, say, lodging incidence, lower values are better. In this case, you should use
#' \code{mresp = 0} to rescale the response variable so that the lowest values will become 100
#' and the highest values 0.
#' @param increment The increment in the weight ratio for stability and mean performance.
#' Se the \bold{Details} section for more information.
#' @param saveWAASY Automatically save the WAASY values when the weight for
#' stability is \code{saveWAASY}. Default is 50. Please, note that \code{saveWAASY}
#' @param prob The p-value for considering an interaction principal component axis significant.
#' must be multiple of \code{increment}. If this assumption is not valid, an error will be occur.
#' @param progbar A logical argument to define if a progress bar is shown.
#' Default is \code{TRUE}.
#' @references Olivoto, T., A.D.C. L{\'{u}}cio, J.A.G. da silva, V.S. Marchioro,
#'  V.Q. de Souza, and E. Jost. 2019. Mean performance and stability in multi-environment
#'   trials I: Combining features of AMMI and BLUP techniques. Agron. J.
#'   \href{https://dl.sciencesocieties.org/publications/aj/abstracts/0/0/agronj2019.03.0220?access=0&view=pdf}{doi:10.2134/agronj2019.03.0220}
#' @return An object of class \code{wsmp} with the following items for each variable:
#' * \strong{scenarios} A list with the model for all computed scenarios.
#' * \strong{WAASY} The values of the WAASY estimated when the weight for the
#' stability in the loop match with argument \code{saveWAASY}.
#' * \strong{hetdata, hetcomb} The data used to produce the heatmaps.
#' * \strong{Ranks} All the values of WAASY estimated in the different
#' scenarios of WAAS/GY weighting ratio.
#' @md
#' @author Tiago Olivoto \email{tiagoolivoto@@gmail.com}
#' @seealso \code{\link{resca}}
#' @importFrom progress progress_bar
#' @export
#' @examples
#' \donttest{
#' library(metan)
#' model <- waas(data_ge2,
#'              env = ENV,
#'              gen = GEN,
#'              rep = REP,
#'              resp = PH)
#' scenarios <- wsmp(model)
#'}
wsmp <- function(model, mresp = 100, increment = 5, saveWAASY = 50,
                 prob = 0.05, progbar = TRUE) {
  if(!class(model) %in% c("waas", "waasb")){
    stop("The model must be an object of class 'waas' or 'waasb'")
  }
  test <- 100%%increment == 0
  test2 <- saveWAASY%%increment == 0
  if (!mresp %in% c(100, 0)) {
    stop("The value 'mresp' must be 0 or 100.")
  }
  if (test == FALSE) {
    stop("The argument 'increment = ", increment, "' is invalid. Please, note that this value must result in an integer in the expression '100 / increment'. Please, consider changing the values.")
  }
  if (test2 == FALSE) {
    stop("The argument 'saveWAASY = ", saveWAASY, "' must be divisible by 'increment' (",
         increment, "). Please, consider changing the values.")
  }
  datain <- model
  if (class(model) == "waasb") {
    dfs <- list()
    for (k in 1:length(model)) {
      PesoWAAS <- 100
      PesoResp <- 0
      minresp <- 100 - mresp
      data <- datain[[k]][["residuals"]] %>% select(ENV, GEN, REP, Y)
      nam <- names(datain[k])
      Nenv <- length(unique(data$ENV))
      Ngen <- length(unique(data$GEN))
      minimo <- min(Nenv, Ngen) - 1
      ncomb <- (100/increment) + 1
      totalcomb <- ncomb * minimo
      CombWAASY <- data.frame(type = matrix(".", (Ngen +
                                                    Nenv), 1))
      ovmean <- mean(data$Y)
      WAASY.Values <- list()
      initial <- 0
      model <- suppressWarnings(suppressMessages(lme4::lmer(Y ~
                                                              REP %in% ENV + ENV + (1 | GEN) + (1 | GEN:ENV),
                                                            data = data)))
      summ <- summary(model)
      bups <- lme4::ranef(model)
      blups <- data.frame(Names = rownames(bups$`GEN:ENV`))
      blups <- blups %>% data.frame(do.call("rbind", strsplit(as.character(blups$Names),
                                                              ":", fixed = TRUE))) %>% dplyr::select(-Names) %>%
        dplyr::select(-X1, everything()) %>% dplyr::mutate(BLUPge = bups[[1]]$`(Intercept)`) %>%
        dplyr::rename(Code = X2, GEN = X1) %>% dplyr::arrange(Code)
      intmatrix <- by(blups[, 3], blups[, c(2, 1)], function(x) sum(x,
                                                                    na.rm = TRUE))
      s <- svd(intmatrix)
      U <- s$u[, 1:minimo]
      LL <- diag(s$d[1:minimo])
      V <- s$v[, 1:minimo]
      Eigenvalue <- data.frame(Eigenvalue = s$d[1:minimo]^2) %>%
        dplyr::mutate(Proportion = s$d[1:minimo]^2/sum(s$d[1:minimo]^2) *
                        100, Accumulated = cumsum(Proportion), PC = paste("PC",
                                                                          1:minimo, sep = "")) %>% dplyr::select(PC,
                                                                                                                 everything())
      SCOREG <- U %*% LL^0.5
      SCOREE <- V %*% LL^0.5
      colnames(SCOREG) <- colnames(SCOREE) <- paste("PC",
                                                    1:minimo, sep = "")
      MEDIAS <- data %>% select(ENV, GEN, Y) %>% dplyr::group_by(ENV,
                                                                 GEN) %>% dplyr::summarise(Y = mean(Y)) %>% ungroup()
      MGEN <- MEDIAS %>% group_by(GEN) %>% summarise(Y = mean(Y)) %>%
        mutate(type = "GEN")
      MGEN <- cbind(MGEN, SCOREG) %>% rename(Code = GEN)
      MENV <- MEDIAS %>% group_by(ENV) %>% summarise(Y = mean(Y)) %>%
        mutate(type = "ENV")
      MENV <- cbind(MENV, SCOREE) %>% rename(Code = ENV)
      Escores <- rbind(MGEN, MENV) %>% select(type, everything())
      Pesos <- data.frame(Percent = Eigenvalue$Proportion)
      if (progbar == TRUE) {
        pb <- progress_bar$new(
          format = ":what [:bar]:percent (:eta left)",
          clear = F, total = totalcomb, width = 90)
      }
      for (k in 1:ncomb) {
        WAASB <- Escores %>% select(contains("PC")) %>%
          abs() %>% t() %>% as.data.frame() %>% mutate(Percent = Pesos$Percent)
        WAASAbsInicial <- Escores %>% mutate(WAASB = sapply(WAASB[,
                                                                  -ncol(WAASB)], weighted.mean, w = WAASB$Percent)) %>%
          group_by(type) %>% mutate(PctResp = (mresp -
                                                 minresp)/(max(Y) - min(Y)) * (Y - max(Y)) +
                                      mresp, PctWAASB = (minresp - mresp)/(max(WAASB) -
                                                                             min(WAASB)) * (WAASB - max(WAASB)) + minresp,
                                    wRes = PesoResp, wWAASB = PesoWAAS, OrResp = rank(-Y),
                                    OrWAASB = rank(WAASB), OrPC1 = rank(abs(PC1)),
                                    WAASBY = ((PctResp * wRes) + (PctWAASB * wWAASB))/(wRes +
                                                                                         wWAASB), OrWAASBY = rank(-WAASBY)) %>% dplyr::ungroup()
        inicial <- as.data.frame(WAASAbsInicial$OrWAASB)
        colnames(inicial) <- paste0(minimo, "PC")
        SigPC2 <- 1
        for (j in 1:minimo) {
          WAASB <- Escores %>% select(contains("PC")) %>%
            abs() %>% t() %>% as.data.frame() %>% slice(1:SigPC2) %>%
            mutate(Percent = Pesos$Percent[1:SigPC2])
          WAASAbs <- Escores %>% mutate(WAASB = sapply(WAASB[,
                                                             -ncol(WAASB)], weighted.mean, w = WAASB$Percent)) %>%
            group_by(type) %>% mutate(PctResp = (mresp -
                                                   minresp)/(max(Y) - min(Y)) * (Y - max(Y)) +
                                        mresp, PctWAASB = (minresp - mresp)/(max(WAASB) -
                                                                               min(WAASB)) * (WAASB - max(WAASB)) + minresp,
                                      wRes = PesoResp, wWAASB = PesoWAAS, OrResp = rank(-Y),
                                      OrWAASB = rank(WAASB), OrPC1 = rank(abs(PC1)),
                                      WAASBY = ((PctResp * wRes) + (PctWAASB *
                                                                      wWAASB))/(wRes + wWAASB), OrWAASBY = rank(-WAASBY)) %>%
            dplyr::ungroup()
          results <- as.data.frame(WAASAbs$OrWAASB)
          names(results) <- paste0(SigPC2, "PCA")
          final <- cbind(results, inicial)
          inicial <- final
          SigPC2 <- SigPC2 + 1
          ProcdAtua <- j
          initial <- initial + 1
          if (progbar == TRUE) {
            pb$tick(tokens = list(what = paste("Ranks considering ", PesoResp, "% for GY and ", PesoWAAS, "% for WAASB", sep = "")))
          }
        }
        initial <- initial
        WAAS <- WAASAbsInicial
        WAAS$type <- ifelse(WAAS$type == "GEN", "Genotype",
                            "Environment")
        CombWAASY[[sprintf("%.0f/%.0f", PesoWAAS, PesoResp)]] <- WAASAbsInicial$OrWAASBY
        WAASY.Values[[paste("WAAS/GY", PesoWAAS, "/",
                            PesoResp)]] <- data.frame(WAAS)
        PesoResp <- PesoResp + increment
        PesoWAAS <- PesoWAAS - increment
        if (PesoWAAS + increment == saveWAASY) {
          genotypes <- WAAS %>% dplyr::filter(type ==
                                                "Genotype") %>% dplyr::select(Code, wRes,
                                                                              wWAASB, WAASBY) %>% dplyr::arrange(WAASBY) %>%
            mutate(Mean = ifelse(WAASBY < mean(WAASBY),
                                 "below", "above"))
        }
      }
      Rank <- final[, -(SigPC2)]
      Names <- WAASAbsInicial %>% select(type, Code, OrResp,
                                         OrPC1, OrWAASB)
      Rank <- cbind(Names, Rank) %>% as_tibble(rownames = NA)
      hetdata <- Rank %>% dplyr::filter(type == "GEN") %>%
        column_to_rownames("Code") %>% select(contains("PCA")) %>%
        as_tibble(rownames = NA)
      CombWAASY %<>% select(-type) %>% mutate(type = Names$type,
                                              Code = Names$Code) %>% select(type, Code, everything()) %>%
        dplyr::filter(type == "GEN") %>% column_to_rownames("Code") %>%
        select(contains("/")) %>% as_tibble(rownames = NA)
      PC1 <- Pesos[1, 1]
      PC2 <- Pesos[2, 1]
      mean <- mean(WAAS$Y)
      dfs[[paste(nam)]] <- structure(list(scenarios = WAASY.Values,
                                          WAASY = genotypes, hetcomb = CombWAASY, hetdata = hetdata,
                                          Ranks = Rank), class = "wsmp")
    }
  }
  if (class(model) == "waas") {
    dfs <- list()
    for (k in 1:length(model)) {
      PesoWAAS <- 100
      PesoResp <- 0
      minresp <- 100 - mresp
      data <- datain[[k]][["residuals"]] %>% select(ENV,
                                                    GEN, REP, mean)
      nam <- names(datain[k])
      Nenv <- length(unique(data$ENV))
      Ngen <- length(unique(data$GEN))
      minimo <- min(Nenv, Ngen) - 1
      ncomb <- (100/increment) + 1
      CombWAASY <- data.frame(type = matrix(".", (Ngen +
                                                    Nenv), 1))
      WAASY.Values <- list()
      model <- performs_ammi(data, ENV, GEN, REP, mean, verbose = FALSE)[[1]]
      anova <- model$ANOVA
      PC <- model$PCA
      MeansGxE <- model$MeansGxE
      totalcomb <- ncomb * nrow(PC)
      initial <- 0
      if (progbar == TRUE) {
        pb <- progress_bar$new(
          format = ":what [:bar]:percent (:eta left)",
          clear = F, total = totalcomb, width = 90)
      }
      for (k in 1:ncomb) {
        Escores <- model$model
        SigPC1 <- nrow(PC[which(PC[, 6] < prob), ])
        Pesos <- as.data.frame(model$PCA[7][c(1:SigPC1),
                                                 ])
        colnames(Pesos) <- "Percent"
        WAAS <- Escores %>% select(contains("PC")) %>%
          abs() %>% t() %>% as.data.frame() %>% slice(1:SigPC1) %>%
          mutate(Percent = Pesos$Percent)
        WAASAbsInicial <- Escores %>% mutate(WAAS = sapply(WAAS[,
                                                                -ncol(WAAS)], weighted.mean, w = WAAS$Percent)) %>%
          group_by(type) %>% mutate(PctResp = (mresp -
                                                 minresp)/(max(Y) - min(Y)) * (Y - max(Y)) +
                                      mresp, PctWAAS = (minresp - mresp)/(max(WAAS) -
                                                                            min(WAAS)) * (WAAS - max(WAAS)) + minresp,
                                    wRes = PesoResp, wWAAS = PesoWAAS, OrResp = rank(-Y),
                                    OrWAAS = rank(WAAS), OrPC1 = rank(abs(PC1)),
                                    WAASY = ((PctResp * wRes) + (PctWAAS * wWAAS))/(wRes +
                                                                                      wWAAS), OrWAASY = rank(-WAASY)) %>% dplyr::ungroup()
        inicial <- as.data.frame(WAASAbsInicial$OrWAAS)
        colnames(inicial) <- paste0(SigPC1, "PCA")
        SigPC2 <- 1
        for (j in 1:nrow(PC)) {
          WAAS <- Escores %>% select(contains("PC")) %>%
            abs() %>% t() %>% as.data.frame() %>% slice(1:SigPC2) %>%
            mutate(Percent = Pesos$Percent[1:SigPC2])
          WAASAbs <- Escores %>% mutate(WAAS = sapply(WAAS[,
                                                           -ncol(WAAS)], weighted.mean, w = WAAS$Percent)) %>%
            group_by(type) %>% mutate(PctResp = (mresp -
                                                   minresp)/(max(Y) - min(Y)) * (Y - max(Y)) +
                                        mresp, PctWAAS = (minresp - mresp)/(max(WAAS) -
                                                                              min(WAAS)) * (WAAS - max(WAAS)) + minresp,
                                      wRes = PesoResp, wWAAS = PesoWAAS, OrResp = rank(-Y),
                                      OrWAAS = rank(WAAS), OrPC1 = rank(abs(PC1)),
                                      WAASY = ((PctResp * wRes) + (PctWAAS * wWAAS))/(wRes +
                                                                                        wWAAS), OrWAASY = rank(-WAASY)) %>% dplyr::ungroup()
          results <- as.data.frame(WAASAbs$OrWAAS)
          names(results) <- paste0(SigPC2, "PCA")
          final <- cbind(results, inicial)
          inicial <- final
          SigPC2 %<>% +1
          ProcdAtua <- j
          initial <- initial + 1
          if (progbar == TRUE) {
            pb$tick(tokens = list(what = paste("Ranks considering ", PesoResp, "% for GY and ", PesoWAAS, "% for WAAS", sep = "")))
          }
        }
        initial <- initial
        WAAS <- WAASAbsInicial
        WAAS$type <- ifelse(WAAS$type == "GEN", "Genotype",
                            "Environment")
        CombWAASY[[sprintf("%.0f/%.0f", PesoWAAS, PesoResp)]] <- WAASAbsInicial$OrWAASY
        WAASY.Values[[paste("WAAS/GY", PesoWAAS, "/",
                            PesoResp)]] <- as_tibble(WAAS)
        PesoResp <- PesoResp + increment
        PesoWAAS <- PesoWAAS - increment
        if (PesoWAAS + increment == saveWAASY) {
          genotypes <- WAAS %>% dplyr::filter(type ==
                                                "Genotype") %>% dplyr::select(Code, wRes,
                                                                              wWAAS, WAASY) %>% dplyr::arrange(WAASY) %>%
            mutate(Mean = ifelse(WAASY < mean(WAASY),
                                 "below", "above"))
        }
      }
      Rank <- final[, -(SigPC2)]
      Names <- WAASAbsInicial %>% select(type, Code, OrResp,
                                         OrPC1, OrWAAS)
      Rank <- cbind(Names, Rank) %>% as_tibble(rownames = NA)
      hetdata <- Rank %>% dplyr::filter(type == "GEN") %>%
        column_to_rownames("Code") %>% select(contains("PCA")) %>%
        as_tibble(rownames = NA)
      CombWAASY %<>% select(-type) %>% mutate(type = Names$type,
                                              Code = Names$Code) %>% select(type, Code, everything()) %>%
        dplyr::filter(type == "GEN") %>% column_to_rownames("Code") %>%
        select(contains("/")) %>% as_tibble(rownames = NA)
      PC1 <- Pesos[1, 1]
      PC2 <- Pesos[2, 1]
      mean <- mean(WAAS$Y)
      dfs[[paste(nam)]] <- structure(list(scenarios = WAASY.Values,
                                          WAASY = genotypes, hetcomb = CombWAASY, hetdata = hetdata,
                                          Ranks = Rank), class = "wsmp")
    }
  }
  return(structure(dfs, class = "wsmp"))
}
```
