## Supplementary material for "metan: an R package for multi-environment trial analysis": available at: zzz.R

```
#' @import ggplot2
#' @import stats
#' @importFrom dplyr  select  everything  mutate  group_by  group_by_if  group_keys
#'             group_split  left_join  sample_n  anti_join pull  summarise_all
#'             select_if arrange slice contains top_n summarise ungroup rename
#'             is_grouped_df mutate_at desc n summarise_if funs
#' @importFrom tibble rownames_to_column column_to_rownames tibble as_tibble
#' @importFrom grid grobTree textGrob
#' @importFrom ggrepel  geom_text_repel geom_label_repel
#' @importFrom magrittr  %>% %<>%
#' @importFrom GGally  ggally_text  eval_data_col  ggpairs
#' @importFrom ade4  mantel.randtest
#' @importFrom dendextend  set  rotate_DendSer
#' @importFrom lme4  ranef  VarCorr  fortify.merMod
#' @importFrom lmerTest  ranova  lmer
#' @importFrom gplots  heatmap.2
#' @importFrom FWDselect  selection
#' @importFrom grDevices  colorRampPalette  dev.off  pdf
#'             chull  tiff  boxplot.stats
#' @importFrom methods  is  as
#' @importFrom graphics  plot  boxplot  hist  par  mtext abline
#' @importFrom utils  head combn  stack
#' @importFrom methods setClass setGeneric setMethod setRefClass
#'
NULL

#' @title Data for examples
#'
#' @description This dataset contains the means for grain yield of 10 genotypes cultivated
#' in 5 environments. The interaction effects for this data is found in
#' \code{\link{int.effects}}
#'
#' @name meansGxE
#' @docType data
#' @author Tiago Olivoto \email{tiagoolivoto@@gmail.com}
#' @keywords data
NULL

#' Data for examples
#'
#' @name int.effects
#' @docType data
#' @author Tiago Olivoto \email{tiagoolivoto@@gmail.com}
#' @keywords data
NULL


#' @title Multi-environment trial of oat
#' @description This dataset contain data on two variables assessed in 10 genotypes growing
#' in in 11 environments. The experimental design was a RCBD with 3
#' replicates(blocks). This data provide examples for several functions of
#' \pkg{metan} package.
#' @format A tibble with 420 observations on the following 5 variables.
#' * \strong{ENV} A factor with 14 levels; each level represents one cultivation
#' environment.
#' * \strong{GEN} A factor with 10 levels; each level represents one genotype.
#' * \strong{REP} A factor with 3 levels; each level represents one
#' replication/block.
#' * \strong{GY} A continuous variable (grain yield) observed in each plot.
#' * \strong{HM} A continuous variable (hectoliter mass) observed in each plot.
#' @md
#' @source Personal data
#' @name data_ge
#' @docType data
#' @author Tiago Olivoto \email{tiagoolivoto@@gmail.com}
#' @keywords data
NULL

#' @title Multi-environment trial of maize
#' @description This dataset contain data on 15 traits assessed in 13 maize hybrids growing
#' in 4 environments. The experimental design was a RCBD with 3 blocks and 1
#' replications per block. It may be used as example in several functions of
#' \pkg{metan} package.
#' @format A tibble with 156 observations on the following 18 variables.
#' * \strong{ENV} A factor with 4 levels; each level represents one cultivation
#' environment.
#' * \strong{GEN} A factor with 13 levels; each level represents one maize hybrid.
#' * \strong{REP} A factor with 3 levels; each level represents one
#' replication/block.
#' * \strong{PH} Plant height, in cm.
#' * \strong{EH} Ear height, in cm.
#' * \strong{EP} Ear position, i.e., the ratio EH/PH.
#' * \strong{EL} Ear length, in cm.
#' * \strong{ED} Ear diameter, in mm.
#' * \strong{CL} Cob length, in cm.
#' * \strong{CD} Cob diameter, in mm.
#' * \strong{CW} Cob weight, in g.
#' * \strong{KW} Kernel weight, in cm.
#' * \strong{NR} Number of rows.
#' * \strong{NKR} Number of kernels per row.
#' * \strong{CDED} Cob diameter / Ear diameter ratio.
#' * \strong{PERK} Percentage of kernels.
#' * \strong{TKW} Thousand-kernel weight
#' * \strong{NKE} Number of kernels per row.
#' @md
#' @name data_ge2
#' @source Personal data
#' @docType data
#' @author Tiago Olivoto \email{tiagoolivoto@@gmail.com}
#' @keywords data
NULL


#' @title Single maize trial
#' @description This dataset contain data on 15 traits assessed in 13 maize hybrids.
#' The experimental design was a RCBD with 3 blocks and 1
#' replications per block. It is used as an example in the function \code{\link{gamem}}
#'  of the \pkg{metan} package.
#' @format A tibble with 39 observations on the following 17 variables.
#' * \strong{GEN} A factor with 13 levels; each level represents one maize hybrid.
#' * \strong{REP} A factor with 3 levels; each level represents one replication/block.
#' * \strong{PH} Plant height, in cm.
#' * \strong{EH} Ear height, in cm.
#' * \strong{EP} Ear position, i.e., the ratio EH/PH.
#' * \strong{EL} Ear length, in cm.
#' * \strong{ED} Ear diameter, in mm.
#' * \strong{CL} Cob length, in cm.
#' * \strong{CD} Cob diameter, in mm.
#' * \strong{CW} Cob weight, in g.
#' * \strong{KW} Kernel weight, in cm.
#' * \strong{NR} Number of rows.
#' * \strong{NKR} Number of kernels per row.
#' * \strong{CDED} Cob diameter / Ear diameter ratio.
#' * \strong{PERK} Percentage of kernels.
#' * \strong{TKW} Thousand-kernel weight
#' * \strong{NKE} Number of kernels per row.
#' @md
#' @name data_g
#' @source Personal data
#' @docType data
#' @author Tiago Olivoto \email{tiagoolivoto@@gmail.com}
#' @keywords data
NULL


#' @title Data from an alpha lattice design
#' @description Alpha lattice design of spring oats
#' @format A tibble with 72 observations on the following 5 variables.
#' * \strong{PLOT} Plot number
#' * \strong{REP} Replicate code
#' * \strong{BLOCK} Incomplete block code
#' * \strong{GEN} Genotype code
#' * \strong{YIELD} Observed dry matter yield (tonnes/ha)
#'
#' @details A spring oats trial grown in Craibstone. There were 24 varieties
#'  in 3 replicates, each consisting of 6 incomplete blocks of 4 plots.
#'  Planted in a resolvable alpha design. The plots were laid out in a single line.
#' @source J. A. John & E. R. Williams (1995). Cyclic and computer generated designs,
#'  Chapman and Hall, London. Page 146.
#' @name data_alpha
#' @md
#' @docType data
#' @author Tiago Olivoto \email{tiagoolivoto@@gmail.com}
#' @keywords data
NULL

.onAttach <- function(libname, pkgname) {
  packageStartupMessage("\n******************************************************")
  packageStartupMessage("metan has been successfully loaded in R ", paste0(R.Version()[c("major","minor")], collapse = "."))
  packageStartupMessage("Please, see the complete vignette at:\nhttps://tiagoolivoto.github.io/metan/")
  packageStartupMessage("******************************************************\n")
}

if (getRversion() >= "2.15.1") {
  utils::globalVariables(
    c("..density..", ".fitted", ".resid", ".scresid", "Accumul",
      "Accumulated", "BLUPe", "BLUPg", "BLUPge", "CI", "Code", "ENV", "Eigenvalue",
      "FAI", "GEN", "GY_HMRPGV", "GY_RPGV", "Genotype", "MODEL", "Mean", "PC", "PC1",
      "PC2", "Parameters", "PctResp", "PctWAAS", "PctWAASB", "Percent", "PesRes",
      "PesWAAS", "PesWAASB", "Predicted", "Proportion", "RMSPD", "Rank", "ResAMMI",
      "WAAS", "WAASBY", "WAASY", "X1", "X2", "Y", "Ypred", "YpredAMMI", "envPC1",
      "factors", "gen", "genPC1", "ggee", "id", "label", "nominal", "pred", "resOLS",
      "sel", "stdres", "type", "y", "z", "Corr", "combn", "cov2cor", "partial",
      "pt", ".", "linear", "my_custom_cor", "my_custom_smooth", "K", "direct", "VAR",
      "eq", "IndAmb", "REP", "gge", "ind", "cophenetic", "remaining", "index", "ge",
      "FA1", "FA2", "Gen", "wRes", "wWAASB", "OrResp", "OrPC1", "OrWAASB", "wWAAS",
      "OrWAAS", ".stdresid", "WAASB", "grp", "Names", "ID", "MTSI", "Pair", "LL", "UL",
      "ci", "mean_var", "se", "x", "d1", "d2", "radio", "x0", "x1_x", "x1_y", "y0",
      "val", "Statistic", "Cluster", "CCP", "U1", "U2", "V1", "V2", "Var", "LEVEL",
      "GROUP", "Group", "model", "Variance (%)", "Xs", "Xo", "Pr(>Chisq)", "dat",
      "BLOCK", "rowid", "grank", "rMean", "rShukaVar", "value", "DF", "without", "RPGV_Y",
      "HMRPGV_Y", "Pairs", "name", "v1", "v2", "slope", "new_var", "TYPE", "CODE"))
}
```
