## Supplementary material for "metan: an R package for multi-environment trial analysis": available at: metanref.bib

@article{Fox1990,
abstract = {Gains in yield and stability were analysed using grain yield data from the 8th to the 14th International Triticale Yield Nursery (ITYN) and differences in adaptation between complete and substituted triticales (X Triticosecale Wittmack) were examined. A trend of improving yield and adaptation was observed. As highly significant genotype x site interaction each year was not adequately described using regression analysis, yield stability was examined by ranking genotypes within sites and summarizing scores for individual genotypes across sites. Cluster analysis was used to study the relationships among sites and among genotypes. There was little evidence of sites grouping on a geographical basis. Grouping of genotypes showed a tendency for complete and substituted triticales to perform differently, suggesting that diversity for adaptation is maximized by utilizing both types in breeding. Progress for grain yield is compared with more spectacular gains in test weight and reasons for the different rates of improvement are postulated. {\textcopyright} 1990 Kluwer Academic Publishers.},
author = {Fox, P. N. and Skovmand, B. and Thompson, B. K. and Braun, H. J. and Cormier, R.},
doi = {10.1007/BF00040364},
file = {:D$\backslash$:/Documents/Mendeley Desktop/Fox et al/Yield and adaptation of hexaploid spring triticale - 1990 - Fox et al.pdf:pdf},
issn = {00142336},
journal = {Euphytica},
keywords = {X Triticosecale,cluster analysis,complete and substituted types,genotype x site interaction,ranking,triticale},
mendeley-groups = {Doutorado,Doutorado/METAN,Doutorado/Tese},
number = {1},
pages = {57--64},
publisher = {Kluwer Academic Publishers},
title = {{Yield and adaptation of hexaploid spring triticale}},
url = {https://link.springer.com/article/10.1007/BF00040364},
volume = {47},
year = {1990}
}
@article{Huehn1979,
author = {Huehn, M.},
journal = {EDV in Medizin und Biologie},
mendeley-groups = {Doutorado/METAN,Doutorado/Tese},
pages = {112--117},
title = {{Beitrage zur erfassung der phanotypischen stabilitat}},
volume = {10},
year = {1979}
}
@article{Shukla1972,
abstract = {Some statistical aspects of partitioning genotype-environmental components of variability},
author = {Shukla, G. K.},
doi = {10.1038/hdy.1972.87},
file = {:D$\backslash$:/Documents/Mendeley Desktop/Shukla/Some statistical aspects of partitioning genotype-environmental components of variability - 1972 - Shukla.pdf:pdf},
issn = {0018-067X},
journal = {Heredity},
mendeley-groups = {Doutorado/Methods of adaptability,Doutorado/METAN,Doutorado/Tese},
number = {2},
pages = {238--245},
pmid = {4507945},
title = {{Some statistical aspects of partitioning genotype-environmental components of variability}},
url = {http://www.nature.com/articles/hdy197287},
volume = {29},
year = {1972}
}
@article{Schmildt2011,
abstract = {The aim of this study was to compare three methodologies which analyze adaptability and stability of corn cultivars. It evaluated the productivity of corn cultivars of the national experiment of Embrapa, center, regarding the crops 2004/2005, 2005/2006 and 2006/2007, in order to determine the adaptability and stability, by the methods of Lin and Binns with decomposition (CRUZ; CARNEIRO, 2006), Annicchiarico with decomposition (SCHMILDT; CRUZ, 2005) and by another method proposed based on the Annicchiarico method with decomposition. The use of the average standard error instead of the standard deviation in the calculation of the variability of environments for each cultivar, as well as for the increasing of the confidence level of 75 to 95{\%} was proposed. It was noticed that the results of indication of cultivars by the three methods showed Spearman's high correlation (p {\textless} 0.01). The new method presents advantages: it values the deviations between the environments by taking into account the number of environments and presenting the lowest level of significance (allowing lowest type I error). The proposed methodology presents the numerical results of the confidence index equal to the results of the Annicchiarico method with decomposition when the compared number of environments is equal to six.},
author = {Schmildt, Edilson Romais and Nascimento, Adriel Lima and Cruz, Cosme Dami{\~{a}}o and Oliveira, Jaqueline Aparecida Raminelli},
doi = {10.4025/actasciagron.v33i1.5817},
file = {::},
issn = {16799275},
journal = {Acta Scientiarum - Agronomy},
keywords = {Genotype x environment interaction,Genotypic performance,Zea mays},
number = {1},
pages = {51--58},
title = {{Avalia{\c{c}}{\~{a}}o de metodologias de adaptabilidade e estabilidade de cultivares milho}},
volume = {33},
year = {2011}
}

@book{Robertson1959,
address = {New York, US},
author = {Robertson, A.},
booktitle = {Biometrical genetics},
mendeley-groups = {Doutorado},
pages = {186},
publisher = {Pergamon},
title = {{Experimental design on the measurement of heritabilities and genetic correlations: biometrical genetics}},
year = {1959}
}
@article{Cruz1991a,
author = {Cruz, C. D. and Castoldi, F. L.},
file = {::},
journal = {ceres.ufv.br},
mendeley-groups = {Doutorado/Tese},
pages = {422--430},
title = {{Decomposi{\c{c}}{\~{a}}o da intera{\c{c}}{\~{a}}o gen{\'{o}}tipos x ambientes em partes simples e complexa}},
url = {http://www.ceres.ufv.br/ojs/index.php/ceres/article/view/2165},
volume = {38},
year = {1991}
}
@article{Fox1982,
abstract = {In plant breeding yield trials the environmental range often exceeds the genotypic range. In such instances environmental main-effects (mean yields) may confound characterization of selection environments, as the general productivity of an environment may be unrelated to tendencies in the relative performance (ranking) of genetic material grown in that environment.},
author = {Fox, P. N. and Rosielle, A. A.},
doi = {10.1007/BF00039203},
file = {:C$\backslash$:/Users/tiago/AppData/Local/Mendeley Ltd./Mendeley Desktop/Downloaded/Fox, Rosielle - 1982 - Reducing the influence of environmental main-effects on pattern analysis of plant breeding environments.pdf:pdf;:D$\backslash$:/Documents/Mendeley Desktop/Fox, Rosielle/Reducing the influence of environmental main-effects on pattern analysis of plant breeding environments - 1982 - Fox, Rosielle(2).pdf:pdf},
issn = {15735060},
journal = {Euphytica},
keywords = {G x E interaction,environmental main-effects,pattern analysis,ranking,selection},
mendeley-groups = {Doutorado/WAASB},
number = {3},
pages = {645--656},
publisher = {Kluwer Academic Publishers},
title = {{Reducing the influence of environmental main-effects on pattern analysis of plant breeding environments}},
volume = {31},
year = {1982}
}

@article{Olivoto2019,
author = {Olivoto, T. and L{\'{u}}cio, A. D. C and Da silva, J. A. G. and Marchioro, V.S. and de Souza, V.Q. and Jost, E.},
doi = {10.2134/agronj2019.03.0220},
file = {:D$\backslash$:/Documents/Mendeley Desktop/Olivoto et al/Mean performance and stability in multi-environment trials I Combining features of AMMI and BLUP techniques - 2019 - Olivoto et al.pdf:pdf},
journal = {Agronomy Journal},
mendeley-groups = {Doutorado/METAN},
number = {6},
pages = {2949--2960},
title = {{Mean performance and stability in multi-environment trials I: Combining features of AMMI and BLUP techniques}},
volume = {111},
year = {2019}
}
@article{Olivoto2019a,
author = {Olivoto, T. and L{\'{u}}cio, A. D. C and Da silva, J. A. G. and Sari, B. G. and Diel, M. I.},
doi = {10.2134/agronj2019.03.0221},
file = {:D$\backslash$:/Documents/Mendeley Desktop/Olivoto et al/Mean performance and stability in multi-environment trials II Selection based on multiple traits - 2019 - Olivoto et al.pdf:pdf},
journal = {Agronomy Journal},
mendeley-groups = {Doutorado/METAN},
number = {6},
pages = {2961--2969},
title = {{Mean performance and stability in multi-environment trials II: Selection based on multiple traits}},
volume = {111},
year = {2019}
}
@article{murakami2004,
author = {Murakami, D.M. and Cruz, C.D.},
file = {:D$\backslash$:/Documents/Mendeley Desktop/Murakami, Cruz/Proposal of methodologies for environment stratification and analysis of genotype adaptability - 2004 - Murakami, Cruz.pdf:pdf},
journal = {Crop Breeding and Applied Biotechnology},
mendeley-groups = {Mestrado/Capitulo soja},
number = {1},
pages = {7--11},
title = {{Proposal of methodologies for environment stratification and analysis of genotype adaptability}},
url = {http://www.sbmp.org.br/cbab/siscbab/uploads/c8128f42-aefe-cdf5.pdf},
volume = {4},
year = {2004}
}
@book{Yan2003,
abstract = {Sect. I. Genotype-by-Environment Interaction and Stability Analysis. Ch. 1. Genotype-by-Environment Interaction. 1.1. Heredity and Environment. 1.2. Genotype-by-Environment Interaction. 1.3. Implications of GEI in Crop Breeding. 1.4. Causes of Genotype-by-Environment Interaction. Ch. 2. Stability Analyses in Plant Breeding and Performance Trials. 2.1. Stability Concepts and Statistics. 2.2. Dealing with Genotype-by-Environment Interaction. 2.3. GGE Biplot: Genotype + GE Interaction -- Sect. II. GGE Biplot and Multi-Environment Trial Data Analysis. Ch. 3. Theory of Biplot. 3.1. The Concept of Biplot. 3.2. The Inner-Product Property of a Biplot. 3.3. Visualizing the Biplot. 3.4. Relationships among Columns and among Rows.},
author = {Yan, Weikai. and Kang, Manjit S.},
file = {:C$\backslash$:/Users/tiago/AppData/Local/Mendeley Ltd./Mendeley Desktop/Downloaded/Yan, Kang - 2003 - GGE biplot analysis a graphical tool for breeders, geneticists, and agronomists.pdf:pdf},
isbn = {0849313384},
mendeley-groups = {Doutorado/Books},
pages = {271},
publisher = {CRC Press},
title = {{GGE biplot analysis : a graphical tool for breeders, geneticists, and agronomists}},
year = {2003}
}


@article{Yan2007,
author = {Yan, W. and Kang, M. S. and Ma, B. and Woods, S. and Cornelius, P. L.},
doi = {10.2135/cropsci2006.06.0374},
file = {:D$\backslash$:/Documents/Mendeley Desktop/Yan et al/GGE Biplot vs. AMMI Analysis of Genotype-by-Environment Data - 2007 - Yan et al.pdf:pdf},
issn = {1435-0653},
journal = {Crop Science},
mendeley-groups = {Doutorado/Methods of adaptability,Doutorado/Research project},
number = {2},
pages = {641--653},
title = {{GGE Biplot vs. AMMI Analysis of Genotype-by-Environment Data}},
url = {https://www.crops.org/publications/cs/abstracts/47/2/643},
volume = {47},
year = {2007}
}


@article{Wricke1965,
author = {Wricke, G},
journal = {Z. Pflanzenzuchtg},
mendeley-groups = {Mestrado/Capitulo soja},
pages = {127--138},
title = {{Zur berechnung der okovalenz bei sommerweizen und hafer}},
volume = {52},
year = {1965}
}


@article{Alves2018,
abstract = {Despite its numerous important traits for biodiesel production, little research exists on the Jatropha curcas crop, mainly with respect to repeated measures over time (crop years). The present study was thus developed to evaluate repeated measures over time in Jatropha curcas and examine the applicability and efficiency of the harmonic mean of the relative performance of genetic values (HMRPGV) method in the selection of families for yield, stability, and adaptability, simultaneously. The work involved data from the evaluation of 730 individuals of 73 half-sib families, in a complete randomized block design, during six crop years, for the trait grain yield. Restricted maximum likelihood/best linear unbiased prediction (REML/BLUP) procedure was performed to estimate the variance components and predict the genetic values. Ten crop years must be evaluated to obtain 80{\%} of the maximum coefficient of determination and 90{\%} accuracy in terms of genetic gain from selection. The efficiency of performing 10 measurements compared with the situation in which only one measurement is used was 67{\%}. The HMRPGV method has great potential in the selection of Jatropha curcas families for yield, stability, and adaptability, simultaneously.},
author = {Alves, Rodrigo Silva and {de Azevedo Peixoto}, Leonardo and Teodoro, Paulo Eduardo and Silva, Lidiane Aparecida and Rodrigues, Erina Vit{\'{o}}rio and de Resende, Marcos Deon Vilela and Laviola, Bruno Galveas and Bhering, Leonardo Lopes},
doi = {10.1016/J.INDCROP.2018.04.029},
issn = {0926-6690},
journal = {Industrial Crops and Products},
mendeley-groups = {Doutorado/WAASB},
month = {sep},
pages = {290--293},
publisher = {Elsevier},
title = {{Selection of Jatropha curcas families based on temporal stability and adaptability of genetic values}},
url = {https://www.sciencedirect.com/science/article/pii/S0926669018303406},
volume = {119},
year = {2018}
}


@article{Mojena1977,
abstract = {Hierarchical clustering procedures have received a great deal of emphasis in recent years, yet research has lagged in their empirical evaluation and in objective means to aid the user in selecting good partitions (rather than good hierarchies). The present paper aims to correct both of these deficiencies, first by empirically testing selected methods which have become popular and, second by proposing and evaluating statistical stopping rules. Results indicate: 1. that methods vary widely in their performance and 2. that the proposed stopping rules can aid the user in selecting partitions.},
author = {Mojena, R},
doi = {10.1093/comjnl/20.4.359},
file = {:D$\backslash$:/Documents/Mendeley Desktop/Mojena/Hierarchical grouping methods and stopping rules an evaluation - 1977 - Mojena.pdf:pdf},
issn = {0010-4620, 1460-2067},
journal = {The Computer Journal},
number = {4},
pages = {359--363},
shorttitle = {Hierarchical grouping methods and stopping rules},
title = {{Hierarchical grouping methods and stopping rules: an evaluation}},
url = {http://comjnl.oxfordjournals.org/content/20/4/359.short?rss=1{\&}ssource=mfc},
volume = {20},
year = {1977}
}


@article{Eberhart1966,
author = {Eberhart, S. A. and Russell, W. A.},
doi = {10.2135/cropsci1966.0011183X000600010011x},
issn = {0011-183X},
journal = {Crop Science},
mendeley-groups = {Mestrado/Capitulo soja,Doutorado/WAASB},
number = {1},
pages = {36--40},
title = {{Stability parameters for comparing Varieties}},
url = {https://www.crops.org/publications/cs/abstracts/6/1/CS0060010036},
volume = {6},
year = {1966}
}


@article{murakami2004,
author = {Murakami, D.M. and Cruz, C.D.},
file = {:D$\backslash$:/Documents/Mendeley Desktop/Murakami, Cruz/Proposal of methodologies for environment stratification and analysis of genotype adaptability - 2004 - Murakami, Cruz.pdf:pdf},
journal = {Crop Breeding and Applied Biotechnology},
number = {1},
pages = {7--11},
title = {{Proposal of methodologies for environment stratification and analysis of genotype adaptability}},
url = {http://www.sbmp.org.br/cbab/siscbab/uploads/c8128f42-aefe-cdf5.pdf},
volume = {4},
year = {2004}
}


@article{Lin1988,
author = {Lin, C. S. and Binns, M. R.},
doi = {10.4141/cjps88-018},
journal = {Canadian Journal of Plant Science},
mendeley-groups = {Mestrado/Capitulo soja},
number = {1},
pages = {193--198},
title = {{A superiority measure of cultivar performance for cultivar x location data}},
url = {http://pubs.aic.ca/doi/abs/10.4141/cjps88-018},
volume = {68},
year = {1988}
}


@article{Annicchiarico1992,
author = {Annicchiarico, P},
journal = {Journal of Genetics and Breeding},
mendeley-groups = {Mestrado/Capitulo soja},
pages = {269--278},
title = {{Cultivar adaptation and recommendation from alfalfa trials in Northern Italy}},
volume = {46},
year = {1992}
}


  @article{Olivoto:2018,
	title={WAASBdata},
	author={Olivoto, T.},
	journal={Mendeley Data},
	volume = {V1},
	year={2018},
	doi={10.17632/2sjz32k3s3.1}
}

@article{ColombariFilho2013,
author = {{Colombari Filho}, J. M. and Resende, M. D. V. and Morais, O. P. and Castro, A. P. and Guimar{\~{a}}es, {\'{E}}. P. and Pereira, J. A. and Utumi, M. M. and Breseghello, F.},
doi = {10.1007/s10681-013-0922-2},
file = {:D$\backslash$:/Documents/Mendeley Desktop/Colombari Filho et al/Upland rice breeding in Brazil a simultaneous genotypic evaluation of stability, adaptability and grain yield - 2013 - Colombari Filho e.pdf:pdf},
issn = {0014-2336},
journal = {Euphytica},
month = {jul},
number = {1},
pages = {117--129},
title = {{Upland rice breeding in Brazil: a simultaneous genotypic evaluation of stability, adaptability and grain yield}},
url = {http://link.springer.com/10.1007/s10681-013-0922-2},
volume = {192},
year = {2013}
}

@article{Piepho:1994,
	title={Best Linear Unbiased Prediction (BLUP) for regional yield trials: a comparison to additive main effects and multiplicative interaction (AMMI) analysis},
	author={Piepho, H.P.},
	journal={Theor. Appl. Genet.},
	volume = {89},
	number = {5},
	pages={647-654},
	year={1994},
	doi={10.1007/BF00222462}
}

@article{Gauch:1988,
	title={Predictive and postdictive success of statistical analyses of yield trials},
	author={Gauch, H.G, and Zobel, R.W.,},
	journal={Theor. Appl. Genet.},
	volume = {76},
	number = {1},
	pages={1-10},
	year={1988},
	doi={10.1007/BF00288824}
}

@article{Legates:1999,
	title={Evaluating the use of "goodness-of-fit" Measures in hydrologic and hydroclimatic model validation},
	author={Legates, D. R., and McCabe, Gregory J.},
	journal={Water Resources Research},
	volume = {35},
	number = {1},
	pages={233-241},
	year={1999},
	doi={10.1007/BF00288824}
}

@article{Gollob:1968,
author = {Gollob, H. F.},
doi = {10.1007/BF02289676},
file = {:D$\backslash$:/Documents/Mendeley Desktop/Gollob/A statistical model which combines features of factor analytic and analysis of variance techniques. - 1968 - Gollob.pdf:pdf},
journal = {Psychometrika},
number = {1},
pages = {73--115},
pmid = {5239571},
title = {{A statistical model which combines features of factor analytic and analysis of variance techniques.}},
url = {https://link.springer.com/article/10.1007{\%}2FBF02289676},
volume = {33},
year = {1968}
}

@article{Purchase2000,
abstract = {Thirteen winter and intermediate type bread wheat cultivars were evaluated for yield stability under dryland conditions over a four year period from 1991 to 1994 and over a total of 120 environments in the Western, Central and Eastern Free State wheat producing regions of South Africa. The following statistical analyses were conducted and procedures followed to determine yield stability: (i) Shukla's procedure of stability variance(*2j); (ii) Lin and Binns cultivar performance measure (Pi); (iii) Finlay and Wilkenson's regression analysis and coefficient (b); (iv) Eberhart and Russel's deviation from regression (S2d); (v) Wricke's ecovalence (W1); (vi) AMMI model. Since the AMMI model does not make provision for a quantitative stability measure, such a measure was developed to rank genotypes. Total correspondence for significance of Spearman's rank correlation coefficients for the different analysis procedures was noted over the three production regions. No significant rank correlation coefficients were f...},
author = {Purchase, J. L. and Hatting, Hesta and van Deventer, C. S.},
doi = {10.1080/02571862.2000.10634878},
file = {:D$\backslash$:/Documents/Mendeley Desktop/Purchase, Hatting, van Deventer/Genotype × environment interaction of winter wheat ( Triticum aestivum L.) in South Africa II. Stability analysis of yield performance -.pdf:pdf},
journal = {South African Journal of Plant and Soil},
month = {jan},
number = {3},
pages = {101--107},
publisher = {Taylor {\&} Francis Group},
title = {{Genotype × environment interaction of winter wheat ( Triticum aestivum L.) in South Africa: II. Stability analysis of yield performance}},
url = {http://www.tandfonline.com/doi/abs/10.1080/02571862.2000.10634878},
volume = {17},
year = {2000}
}

@article{Sneller1997,
author = {Sneller, C. H. and Kilgore-Norquest, L. and Dombek, D.},
doi = {10.2135/cropsci1997.0011183X003700020013x},
file = {:D$\backslash$:/Documents/Mendeley Desktop/Sneller, Kilgore-Norquest, Dombek/Repeatability of Yield Stability Statistics in Soybean - 1997 - Sneller, Kilgore-Norquest, Dombek(2).pdf:pdf},
journal = {Crop Science},
number = {2},
pages = {383--390},
publisher = {Crop Science Society of America},
title = {{Repeatability of Yield Stability Statistics in Soybean}},
url = {https://www.crops.org/publications/cs/abstracts/37/2/CS0370020383},
volume = {37},
year = {1997}
}

@article{Zali2012,
abstract = {This research was carried out to study stability analysis and comparison between some measures of stability from AMMI model in 17 chickpea genotypes, in RCBD with four replications in five different research stations of Iran across two years (2004-2005). Spearman's rank correlation was used to measure the relationship between measures of stability from AMMI model. To understand better relationships among these methods, principal component analysis, was performed. The first two PCs explained 87.65{\%} of the variance of original variables. The relationships among the different stability statistics from AMMI model are graphically displayed in a biplot of PC1 and PC2. The results also revealed that stability measures can be classified into three groups. The first group included, SIPC4, EV4, MASV, Dz4, AV (AMGE), Wi(AMMI) and FAi which were correlated with mean yield. The second group consists of ASV, Da4, Bi and Za4 which were correlated with mean yield too. Group 3 consist of FPi that was not associated with the more stability methods and mean yield. This measure may not be as good as the other methods. The results of the correlation matrix and the PC analysis between different measures of stability from AMMI model showed that all measures (except for FPi) can be used for evaluating the responses of chickpea genotypes to changing environments. Eventually, according to all measures of stability from AMMI model (except for FPi) and regarding the high mean yield, genotype G13 (FLIP 97-114) was recommended for national release in Iran, as it adequately demonstrated wide adaptation across environments.},
author = {Zali, H. and Farshadfar, E. and Sabaghpour, S.H. and Karimizadeh, R.},
file = {:D$\backslash$:/Documents/Mendeley Desktop/Zali et al/Evaluation of genotype × environment interaction in chickpea using measures of stability from AMMI model - 2012 - Zali et al.pdf:pdf},
journal = {Annals of Biological Research},
number = {7},
pages = {3126--3136},
title = {{Evaluation of genotype × environment interaction in chickpea using measures of stability from AMMI model}},
url = {http://eprints.icrisat.ac.in/7173/},
volume = {3},
year = {2012}
}

@article{Farshadfar2008,
abstract = {In order to determine stable bread wheat genotypes with high grain yield via a single parameter, field experiments were conducted with 20 genotypes for 4 consecutive years in two different conditions (irrigated and rainfed) in a completely randomized block design with three replications in each environment. Combined analysis of variance showed highly significant differences for the GE interaction indicating the possibility of selection for stable entries. The results of additive main effect and multiplicative interaction (AMMI) analysis revealed 10{\%} of total variability was justified by the GE interaction which was 2.5 times more than that of genotypes. Ordination techniques displayed high differences for the interaction principal components (IPC1, IPC2 and IPC3), exhibiting that 83{\%} of the GE sum of squares was justified by AMMI1, AMMI2 and AMMI3, i.e., 3.77 times more than that explained by the linear regression model displaying the relative efficiency of AMMI1 model in comparison with regression model. AMMI stability value (ASV) discriminated genotypes G6, G9, G14, G15, G17 and G18 as the stable entries. As stability per se is not a desirable selection criterion, because the most stable genotypes would not necessarily give the best yield performance, hence, simultaneous consideration of grain yield and ASV in a single non-parametric index entitled: Genotype Selection Index (GSI), discriminated G6 and G17 with general adaptability and high grain yield for rainfed and irrigated conditions which was in agreement with the results of biplot analysis.},
author = {Farshadfar, E},
file = {:D$\backslash$:/Documents/Mendeley Desktop/Farshadfar/Incorporation of AMMI stability value and grain yield in a single non-parametric index (GSI) in bread wheat. - 2008 - Farshadfar.pdf:pdf},
journal = {Pakistan journal of biological sciences},
month = {jul},
number = {14},
pages = {1791--1796},
pmid = {18817218},
title = {{Incorporation of AMMI stability value and grain yield in a single non-parametric index (GSI) in bread wheat.}},
url = {http://www.ncbi.nlm.nih.gov/pubmed/18817218},
volume = {11},
year = {2008}
}


@article{Rocha2018,
author = {Rocha, Jo{\~{a}}o Romero do Amaral Santos de Car and Machado, Juarez Campolina and Carneiro, Pedro Cresc{\^{e}}ncio Souza},
doi = {10.1111/gcbb.12443},
file = {:C$\backslash$:/Users/tiago/AppData/Local/Mendeley Ltd./Mendeley Desktop/Downloaded/Rocha, Machado, Carneiro - 2018 - Multitrait index based on factor analysis and ideotype-design proposal and application on elephant gra.pdf:pdf},
issn = {17571693},
journal = {GCB Bioenergy},
keywords = {BLUP,Cenchrus purpureus,Napier grass,REML,SEM,biomass,energy co-generation,multicollinearity,selection index},
mendeley-groups = {Doutorado/MTSI,Doutorado/WAASB},
month = {jan},
number = {1},
pages = {52--60},
publisher = {Wiley/Blackwell (10.1111)},
title = {{Multitrait index based on factor analysis and ideotype-design: proposal and application on elephant grass breeding for bioenergy}},
url = {http://doi.wiley.com/10.1111/gcbb.12443},
volume = {10},
year = {2018}
}


@article{Olivoto2018,
author = {Olivoto, T. and L{\'{u}}cio, A. D. C and Souza, V. Q. and Nardino, M. and Diel, M. I. and Sari, B. G. and Krysczun, D .K. and Meira, D. and Meier, C.},
doi = {10.2134/agronj2017.09.0566},
file = {:D$\backslash$:/Documents/Mendeley Desktop/Olivoto et al/Confidence interval width for Pearson's correlation coefficient a Gaussian-independent estimator based on sample size and strength of as.pdf:pdf},
journal = {Agronomy Journal},
mendeley-groups = {Doutorado},
month = {nov},
number = {1},
pages = {1--8},
title = {{Confidence interval width for Pearson's correlation coefficient: a Gaussian-independent estimator based on sample size and strength of association}},
url = {https://dl.sciencesocieties.org/publications/aj/abstracts/0/0/agronj2017.09.0566},
volume = {110},
year = {2018}
}


@article{Henderson1975,
abstract = {Mixed linear models are assumed in most animal breeding applications. Convenient methods for computing BLUE of the estimable linear functions of the fixed elements of the model and for computing best linear unbiased predictions of the random elements of the model have been available. Most data available to animal breeders, however, do not meet the usual requirements of random sampling, the problem being that the data arise either from selection experiments or from breeders' herds which are undergoing selection. Consequently, the usual methods are likely to yield biased estimates and predictions. Methods for dealing with such data are presented in this paper.},
author = {Henderson, C. R.},
doi = {10.2307/2529430},
file = {::},
journal = {Biometrics},
mendeley-groups = {Doutorado/Methods of adaptability},
month = {jun},
number = {2},
pages = {423--447},
title = {{Best Linear Unbiased Estimation and Prediction under a Selection Model}},
url = {https://www.jstor.org/stable/2529430?origin=crossref},
volume = {31},
year = {1975}
}


@article{Olivoto2017f,
author = {Olivoto, T. and Souza, V. Q. and Nardino, M. and Carvalho, I. R. and Ferrari, M. and Pelegrin, A. J. and Szareski, V. J. and Schmidt, D.},
doi = {10.2134/agronj2016.04.0196},
file = {:D$\backslash$:/Documents/Mendeley Desktop/Olivoto et al/Multicollinearity in path analysis a simple method to reduce its effects - 2017 - Olivoto et al.pdf:pdf},
issn = {0002-1962},
journal = {Agronomy Journal},
number = {1},
pages = {131--142},
title = {{Multicollinearity in path analysis: a simple method to reduce its effects}},
url = {https://dl.sciencesocieties.org/publications/aj/abstracts/109/1/131},
volume = {109},
year = {2017}
}


@article{Rocha2018,
author = {Rocha, Jo{\~{a}}o Romero do Amaral Santos de Car and Machado, Juarez Campolina and Carneiro, Pedro Cresc{\^{e}}ncio Souza},
doi = {10.1111/gcbb.12443},
file = {:D$\backslash$:/Documents/Mendeley Desktop/Rocha, Machado, Carneiro/Multitrait index based on factor analysis and ideotype-design proposal and application on elephant grass breeding for bioenergy - 2018 -.pdf:pdf},
issn = {17571693},
journal = {GCB Bioenergy},
keywords = {BLUP,Cenchrus purpureus,Napier grass,REML,SEM,biomass,energy co-generation,multicollinearity,selection index},
mendeley-groups = {Doutorado/MTSI,Doutorado/WAASB},
month = {jan},
number = {1},
pages = {52--60},
publisher = {Wiley/Blackwell (10.1111)},
title = {{Multitrait index based on factor analysis and ideotype-design: proposal and application on elephant grass breeding for bioenergy}},
url = {http://doi.wiley.com/10.1111/gcbb.12443},
volume = {10},
year = {2018}
}


@article{Mantel1967,
abstract = {The problem of identifying subtle time-space clustering of disease, as may be occurring in leukemia, is described and reviewed. Published approaches, generally associated with studies of leukemia, not dependent on knowledge of the underlying population for their validity, are directed towards identifying clustering by establishing a relationship between the temporal and the spatial separations for the n ( n - 1)/2 possible pairs which can be formed from the n observed cases of disease. Here it is proposed that statistical power can be improved by applying a reciprocal transform to these separations. While a permutational approach can give valid probability levels for any observed association, for reasons of practicability, it is suggested that the observed association be tested relative to its permutational variance. Formulas and computational procedures for doing so are given.

While the distance measures between points represent symmetric relationships subject to mathematical and geometric regularities, the variance formula developed is appropriate for arbitrary relationships. Simplified procedures are given for the case of symmetric and skew-symmetric relationships. The general procedure is indicated as being potentially useful in other situations as, for example, the study of interpersonal relationships. Viewing the procedure as a regression approach, the possibility for extending it to nonlinear and multivariate situations is suggested.

Other aspects of the problem and of the procedure developed are discussed.

Similarly, pure temporal clustering can be identified by a study of incidence rates in periods of widespread epidemics. In point of fact, many epidemics of communicable diseases are somewhat local in nature and so these do actually constitute temporal-spatial clusters. For leukemia and similar diseases in which cases seem to arise substantially at random rather than as clear-cut epidemics, it is necessary to devise sensitive and efficient procedures for detecting any nonrandom component of disease occurrence.

Various ingenious procedures which statisticians have developed for the detection of disease clustering are reviewed here. These procedures can be generalized so as to increase their statistical validity and efficiency. The technic to be given below for imparting statistical validity to the procedures already in vogue can be viewed as a generalized form of regression with possible useful application to problems arising in quite different contexts.},
author = {Mantel, N},
file = {::},
issn = {0008-5472},
journal = {Cancer research},
mendeley-groups = {Doutorado/MTSI},
month = {feb},
number = {2},
pages = {209--20},
pmid = {6018555},
publisher = {American Association for Cancer Research},
title = {{The detection of disease clustering and a generalized regression approach.}},
url = {http://www.ncbi.nlm.nih.gov/pubmed/6018555},
volume = {27},
year = {1967}
}


@article{Olivoto2017c,
abstract = {Information about data arrangement methodologies and optimal sample size in estimating the Pearson correlation coefficient (r) among maize traits are still limited. Furthermore, some data arrangement methodologies currently used may be increasing multicollinearity in multiple regression analysis. This study aimed to investigate the statistical behavior of the r and the multicollinearity of correlation matrices among maize traits in different data arrangement scenarios and different sample sizes. Data from 45 treatments [15 simple maize hybrids (Zea mays L.) conducted in three locations] were used. Eleven traits were accessed and three datasets (scenarios) were formed: (1) Coming from all the sampled observations (plants), n = 900; (2) Coming from the average of five plants per plot, n = 180; and (3) Coming from the average of treatments, n = 45. A thousand estimates of r were held in each scenario to 60 sample sizes by bootstrap simulations with replacement. Confidence intervals (CI) were estimated. One hundred eighty correlation matrices were estimated and the condition number (CN) calculated. Data coming from average values of plots and average values of treatments overestimates the r up to 24 and 34{\%}, resulting in an increase of 24 and 131{\%} in the matrices' CN. Trait pairs with high r require a smaller number of plants, being the CI inversely proportional to the magnitude of the r. Two hundred and ten plants are sufficient to estimate the r in the CI of 95{\%} {\textless} 0.30. Key words: Average values, bootstrap, confidence intervals, sample tracking, Zea mays L.},
author = {Olivoto, T., and Nardino, M., and Carvalho, I. R., and Follmann, D. N., and Ferrari, M., and de Pelegrin, A. J., and Szareski, V. J. and de Oliveira, A. C., and Caron, B. O., and Souza, V. Q.},
doi = {10.5897/AJAR2016.11799},
file = {:D$\backslash$:/Documents/Mendeley Desktop/Olivoto et al/Optimal sample size and data arrangement method in estimating correlation matrices with lesser collinearity A statistical focus in maize.pdf:pdf},
issn = {1991-637X},
journal = {African Journal of Agricultural Research},
month = {jan},
number = {2},
pages = {93--103},
publisher = {Academic Journals},
title = {{Optimal sample size and data arrangement method in estimating correlation matrices with lesser collinearity: A statistical focus in maize breeding}},
url = {http://academicjournals.org/journal/AJAR/article-abstract/5DB554B62359},
volume = {12},
year = {2017}
}


@article{Yates1938,
author = {Yates, F. and Cochran, W. G.},
doi = {10.1017/S0021859600050978},
file = {:D$\backslash$:/Documents/Mendeley Desktop/Yates, Cochran/The analysis of groups of experiments - 1938 - Yates, Cochran.pdf:pdf},
issn = {0021-8596},
journal = {The Journal of Agricultural Science},
mendeley-groups = {Mestrado/Capitulo soja},
month = {oct},
number = {04},
pages = {556--580},
publisher = {Cambridge University Press},
title = {{The analysis of groups of experiments}},
url = {http://www.journals.cambridge.org/abstract{\_}S0021859600050978},
volume = {28},
year = {1938}
}
