## Supplementary material for "metan: an R package for multi-environment trial analysis": available at: vignettes_metan.Rmd

````
---
title: "Multi-environment Trial Analysis"
author: "Tiago Olivoto"
date: "`r Sys.Date()`"
output: rmarkdown::html_vignette
fig_caption: yes
link-citations: true
bibliography: metanref.bib    
vignette: >
  %\VignetteIndexEntry{Multi-environment Trial Analysis}
  %\VignetteEngine{knitr::rmarkdown}
  %\VignetteEncoding{UTF-8}
---

```{r global_options, include = FALSE}
knitr::opts_chunk$set(comment = "")
```


Provides tools for computing several world-known stability statistics. In the latest version (v 1.2.1), the following stability methods are implemented in their respective functions.

* Parametric methods
   * AMMI analysis with [`performs_ammi()`](https://tiagoolivoto.github.io/metan/reference/performs_ammi.html).
   * GGE biplot analysis with [`gge()`](https://tiagoolivoto.github.io/metan/reference/gge.html).
   * BLUP prediction for multi-environment trials with [`waasb()`](https://tiagoolivoto.github.io/metan/reference/waasb.html).
   * AMMI-based stability statistics with [`AMMI_indexes()`](https://tiagoolivoto.github.io/metan/reference/AMMI_indexes.html).
   * BLUP-based stability indexes with [`Resende_indexes()`](https://tiagoolivoto.github.io/metan/reference/Resende_indexes.html).
   * Joint regression analysis [@Eberhart1966] using [`ge_reg()`](https://tiagoolivoto.github.io/metan/reference/ge_reg.html).
   * Genotypic confidence index [@Annicchiarico1992] using [`Annicchiarico()`](https://tiagoolivoto.github.io/metan/reference/Annicchiarico.html).
   * Modified genotypic confidence index [@Schmildt2011] using [`Schmildt()`](https://tiagoolivoto.github.io/metan/reference/Schmildt.html).
   * Ecovalence [@Wricke1965] using [`ecovalence()`](https://tiagoolivoto.github.io/metan/reference/ecovalence.html).
   * Stability analysis and environmental stratification [@murakami2004] using [`ge_factanal()`](https://tiagoolivoto.github.io/metan/reference/ge_factanal.html).
   * Shukla's stability variance [@Shukla1972] using [`Shukla()`](https://tiagoolivoto.github.io/metan/reference/Shukla.html).
   * Weighted Average of Absolute Scores [@Olivoto2019] using [`waas()`](https://tiagoolivoto.github.io/metan/reference/waas.html) (fixed-effect model) and [`waasb()`](https://tiagoolivoto.github.io/metan/reference/waasb.html) (mixed-effect model).

* Non-parametric methods
   * Superiority index [@Lin1988], using the function [`superiority()`](https://tiagoolivoto.github.io/metan/reference/superiority.html).
   * Fox's stability statistic [@Fox1990] with [`Fox()`](https://tiagoolivoto.github.io/metan/reference/Fox.html).
   * Geometric adaptability index [@Shahbazi2019] with [`gai()`](https://tiagoolivoto.github.io/metan/reference/gai.html).
   * Huehn's stability statistic [@Huehn1979] with [`Huehn()`](https://tiagoolivoto.github.io/metan/reference/Huehn.html).
   * Thennarasu's stability statistics [@Thennarasu1995] with [`Thennarasu()`](https://tiagoolivoto.github.io/metan/reference/Thennarasu.html).
   

The easiest way to compute the above-mentioned stability indexes is by using the function [`ge_stats()`](https://tiagoolivoto.github.io/metan/reference/ge_stats.html). This is a wrapper that basically returns a summary of each method. If you are looking for more details from each method like `plot()` and `print()`, I suggest computing that methods using their own function.

The complete functionality of the package is described at https://tiagoolivoto.github.io/metan/index.html. You're welcome to check it out!

# Brief examples

Brief examples will be shown using the dataset `data_ge` that contains data on two variables assessed in 10 genotypes growing in 14 environments. For more details type `?data_ge`.

# Checking data
First of all, we will check the data for possible problems with the function `inspect()`.
```{r, message=FALSE, warning=FALSE}
library(metan)
inspect(data_ge)
```


Then, the details of the multi-environment trial can be obtained with the function `ge_details()`. Note that to apply the function to all numeric variables quickly, we can use the [select helper](https://tiagoolivoto.github.io/metan/articles/vignettes_helper.html#select-helpers) `everything()` in the argument `resp`.

```{r}
ge_details(data_ge,
           env = ENV,
           gen = GEN,
           resp = everything())
```

We can create a plot to show the performance of the genotypes across the environments with `ge_plot()`.

```{r fig.height=4, fig.width=5}
ge_plot(data_ge, GEN, ENV, GY)
```

Or obtain the means for genotypes, environments or genotype-environment interaction with `ge_means()`. Note that the function `round_cols()` provided by `metan` round all numeric columns of a data frame to two (default) significant figures.
```{r}
mge <- ge_means(data_ge,
                env = ENV,
                gen = GEN,
                resp = everything())
# Genotype-environment means
get_model_data(mge) %>% round_cols()
# Environment means
get_model_data(mge, what = "env_means") %>% round_cols()
# Genotype means
get_model_data(mge, what = "gen_means") %>% round_cols()
```


# AMMI model
## Fitting the model
The AMMI model may be fitted with with both functions `performs_ammi()` and `waas()`, which is the acronym for the weighted average of absolute scores [@Olivoto2019].

```{r}
ammi_model <- performs_ammi(data_ge, ENV, GEN, REP, resp = c(GY, HM))
waas_index <- waas(data_ge, ENV, GEN, REP, GY, verbose = FALSE)
```

## Cross-validation procedures
The cross-validation procedures implemented in the `metan` are based on the splitting of the original data into a training set and a validation set. The model is fitted using the training set and the predicted value is compared with the validation set. This process is iterated many times, say, 1000 times. The lesser the difference between predicted and validation data, the higher the predictive accuracy of the model. More information may be found [here.](https://tiagoolivoto.github.io/metan/articles/vignettes_cross-validation.html)


## Biplots

The well-known AMMI2 biplot may be obtained using the function `plot_scores()`. ggplot2-based graphics are obtained. Please, note that since `performs_ammi()` and , `waas()` functions allow analyzing multiple variables at the same time, e.g., `resp = c(v1, v2, ...)`, the output `ammi_model` is a list that in this case has two elements, (GY and HM).

```{r, fig.height=8, fig.width=5,  message=FALSE, warning=FALSE}
a <- plot_scores(ammi_model, axis.expand = 1.2)
b <- plot_scores(ammi_model,
                 type = 2,
                 polygon = TRUE,
                 col.gen = "black",
                 col.env = "gray70",
                 col.segm.env = "gray70",
                 axis.expand = 1.5)
arrange_ggplot(a, b, labels = letters[1:2], ncol = 1)
```


## Predict the response variable
The S3 method `predict()` is implemented for objects of class `performs_ammi` and may be used to estimate the response of each genotype in each environment considering different number of Interaction Principal Component Axis (IPCA). As a example, to predict the variables GY and HM we will use four and six IPCA (number of significant IPCAs, respectively). In addition, we will create a two way table with `make_mat()` to show the predicted values for the variable GY.

```{r }
predicted <- predict(ammi_model, naxis = c(4, 6))
make_mat(predicted$GY, GEN, ENV, YpredAMMI) %>% 
  round_cols()
```

# BLUP model
The implementation of linear-mixed effect models to predict the response variable in MET is made with the function `waasb()`. By default, genotype and genotype-vs-environment interaction are assumed to have random effects. Use the argument `random` to change this default. In the following example the model is fitted to all numeric variables in `data_ge2`.

```{r warning=FALSE, message=FALSE}
model2 <- waasb(data_ge, ENV, GEN, REP, everything())
```

## Residual plots
Several residual plots may be obtained applying the S3 generic function `plot()` to an object of class `waasb` (or `waas`). 

```{r,fig.height=12, fig.width=4, message=FALSE, warning=FALSE}
plot(model2, which = c(1, 2, 7), ncol = 1)
```

## Distribution of random effects
The distribution of the random effects may be obtained using the argument
`type = "re"`.

```{r,fig.height=8, fig.width=4}
plot(model2, type = "re", which = c(1, 2), ncol = 1)
```

## Genetic parameters and variance components
We can get easily the model results such as the Likelihood Ration Test for random effects, the variance components, and the BLUPs for genotypes with `get_model_data()`. By default for an object of class `waasb` the function returns the genetic parameters.

```{r}
get_model_data(model2) %>% round_cols(digits = 3)

```


## Plotting the BLUPs for genotypes

```{r, fig.height=8, fig.width=4}
library(ggplot2)
c <- plot_blup(model2)
d <- plot_blup(model2,
               prob = 0.1,
               col.shape  =  c("gray20", "gray80")) +
      coord_flip()
arrange_ggplot(c, d,labels = letters[3:4], ncol = 1)
```


## BLUPS for genotype-vs-environment interaction

```{r }
get_model_data(model2, what = "blupge") %>% 
  round_cols()
```


## BLUP-based stability index
The WAASB index [@Olivoto2019] is a quantitative stability measure based on the weighted average of the absolute scores from the singular value decomposition of the BLUPs for genotype-vs-interaction effects. We can use the argument `what = "WAASB"` to get this statistic.

```{r }
get_model_data(model2, what = "WAASB") %>% 
  round_cols()
```


The function `Resende_indexes()` may be used to compute the harmonic mean of genotypic values (HMGV), the relative performance of the genotypic values (RPGV) and the harmonic mean of the relative performance of genotypic values (HMRPGV). See @Alves2018 for more details. We use the function `get_model_data()` to get the HMRPGV (default) for all analyzed variables.

```{r }
index <- Resende_indexes(model2)
get_model_data(index, "HMRPGV") %>% 
  round_cols()
```


# GGE model
## Fitting the model
The GGE model is fitted with the function `gge()`. This function produces a GGE model based on both a two-way table (in our case the object `table`) with genotypes in the rows and environments in columns, or a data.frame containing at least the columns for genotypes, environments and the response variable(s).

```{r echo = TRUE}
gge_model <- gge(data_ge, ENV, GEN, GY)

```

## Visualizing the Biplot
The generic function `plot()` is used to generate a biplot using as input a fitted model of class `gge`.  The type of biplot is chosen by the argument `type` in the function.  Ten biplots type are available according to {@Yan2003}.

* `type = 1` A basic biplot.
* `type =  2` Mean performance vs. stability.
* `type =  3` Which-won-where.
* `type =  4` Discriminativeness vs. representativeness.
* `type =  5` Examine an environment.
* `type =  6` Ranking environments.
* `type =  7` Examine a genotype.
* `type =  8` Ranking gentoypes.
* `type =  9` Compare two genotypes.
* `type =  10` Relationship among environments.

```{r echo = TRUE, fig.width = 4, fig.height=8, message=F, warning=F}
e <- plot(gge_model)
f <- plot(gge_model, type = 2)
arrange_ggplot(e, f, labels = letters[5:6], ncol = 1)
```


# Wrapper function `ge_stats()`
To compute all the stability statistics at once, we can use the function `ge_stats()`. Again we get the results with `get_model_data()`.

```{r}
stat_ge <- ge_stats(data_ge, ENV, GEN, REP, GY)
get_model_data(stat_ge) %>% 
  round_cols()
```


# Selection based on multiple traits

The multi-trait stability index (MTSI) was proposed by @Olivoto2019a and is used for simultaneous selection considering mean performance and stability (of several traits) in the analysis of METs using both fixed and mixed-effect models. For more details see the [complete vignette](https://tiagoolivoto.github.io/metan/articles/vignettes_indexes.html).

# Getting help

* If you encounter a clear bug, please file a minimal reproducible example on [github](https://github.com/TiagoOlivoto/metan/issues)

# References
````
