## Supplementary figures and images for "metan: an R package for multi-environment trial analysis"

### logo.png

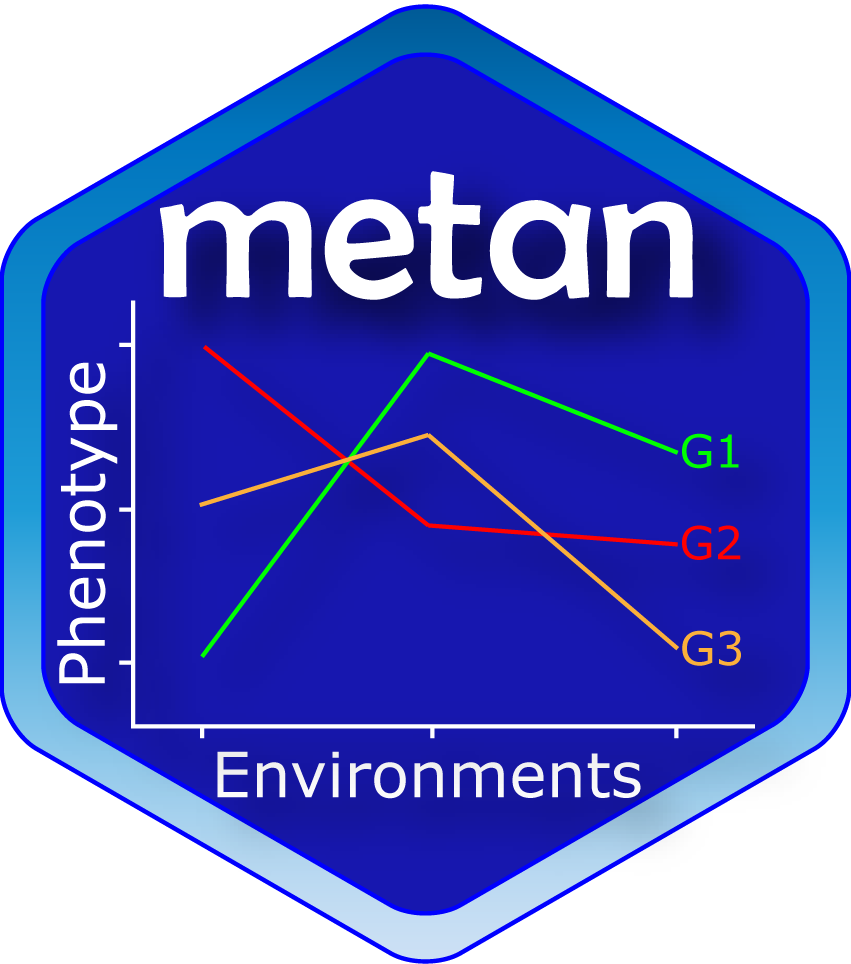

### README-AMMI-1.png

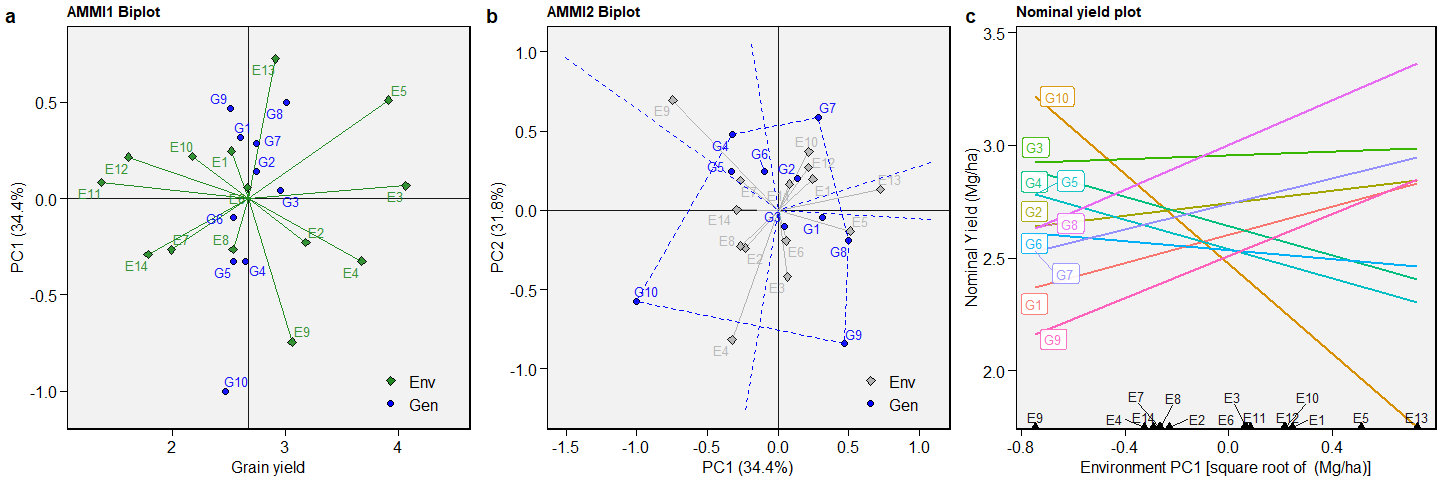

### README-BLUP-1.png

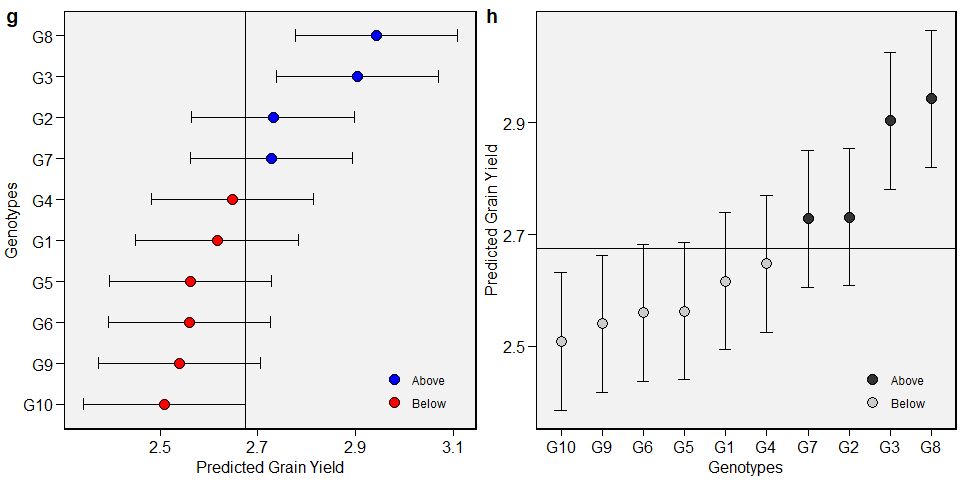

### README-GGE-1.png

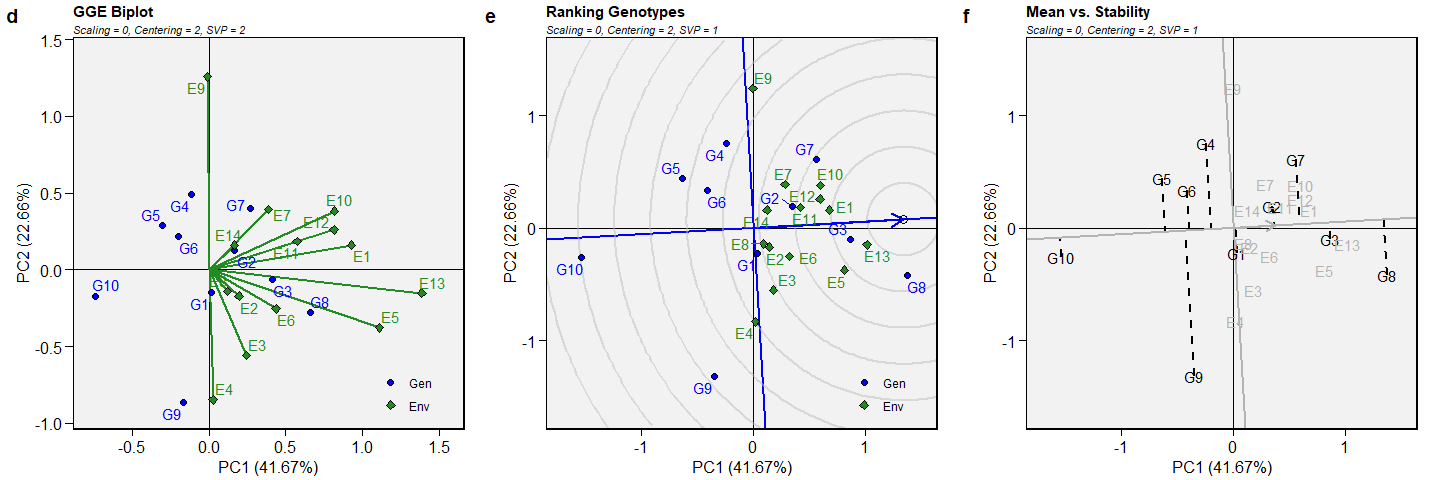

### README-unnamed-chunk-4-1.png

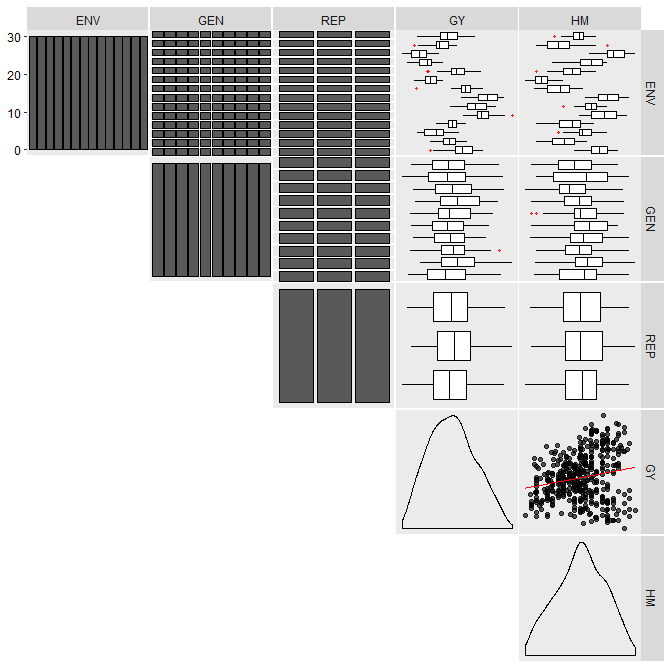

### README-unnamed-chunk-4-2.png

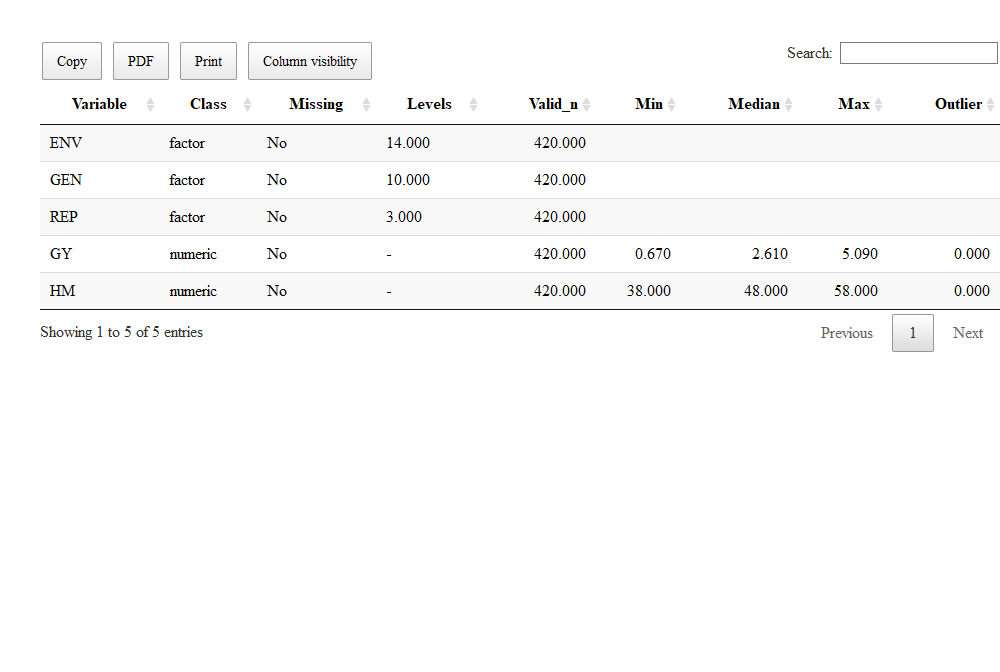

### README-unnamed-chunk-5-1.png

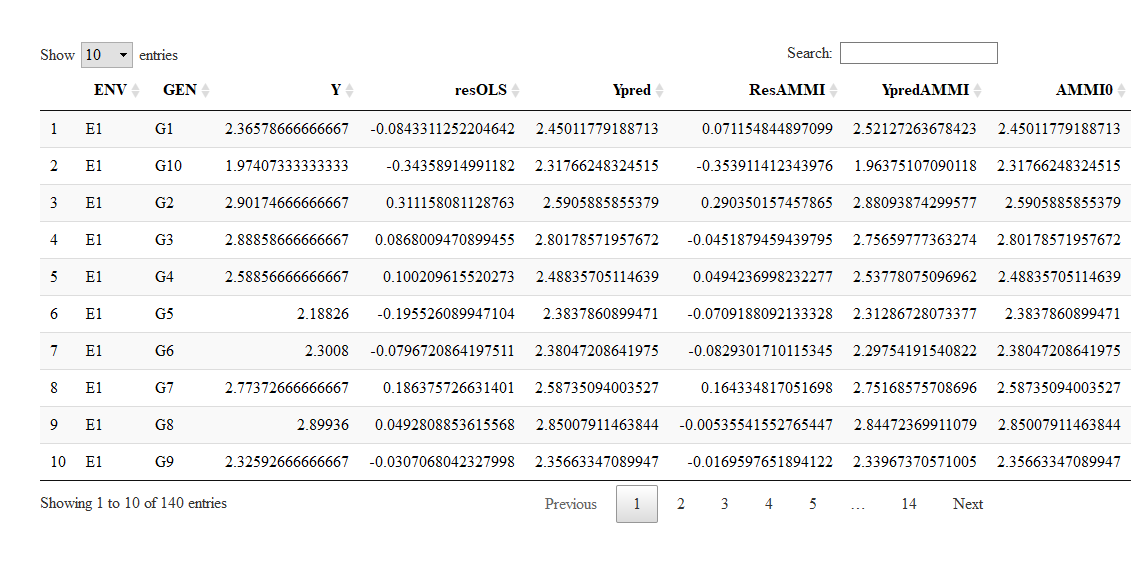

### README-unnamed-chunk-6-1.png

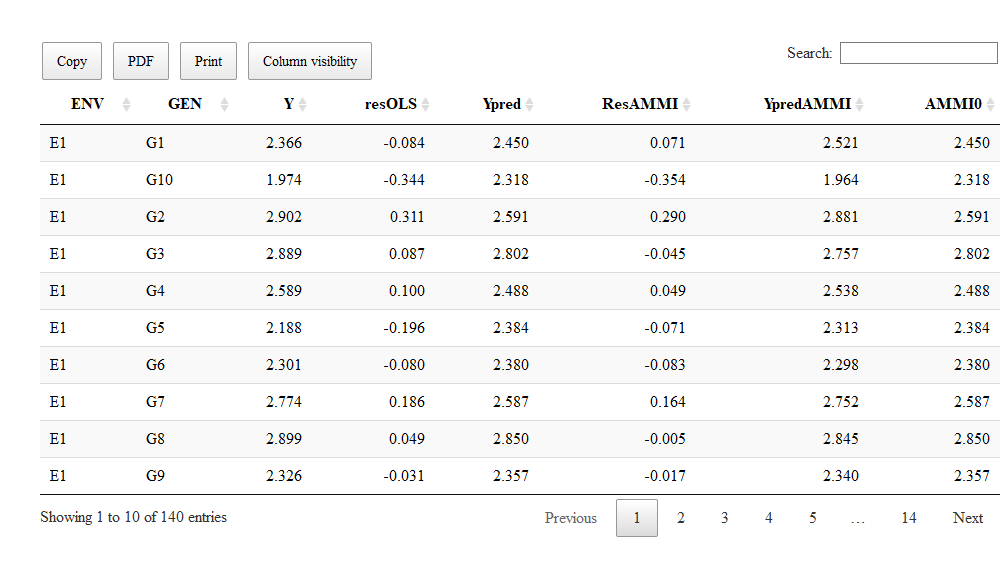

### README-unnamed-chunk-7-1.png

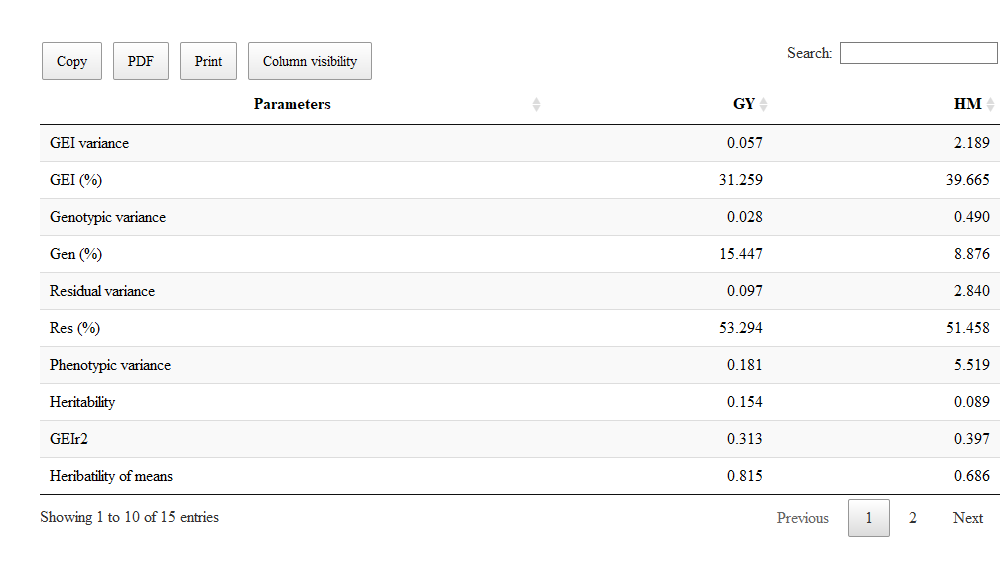

### README-unnamed-chunk-7-2.png

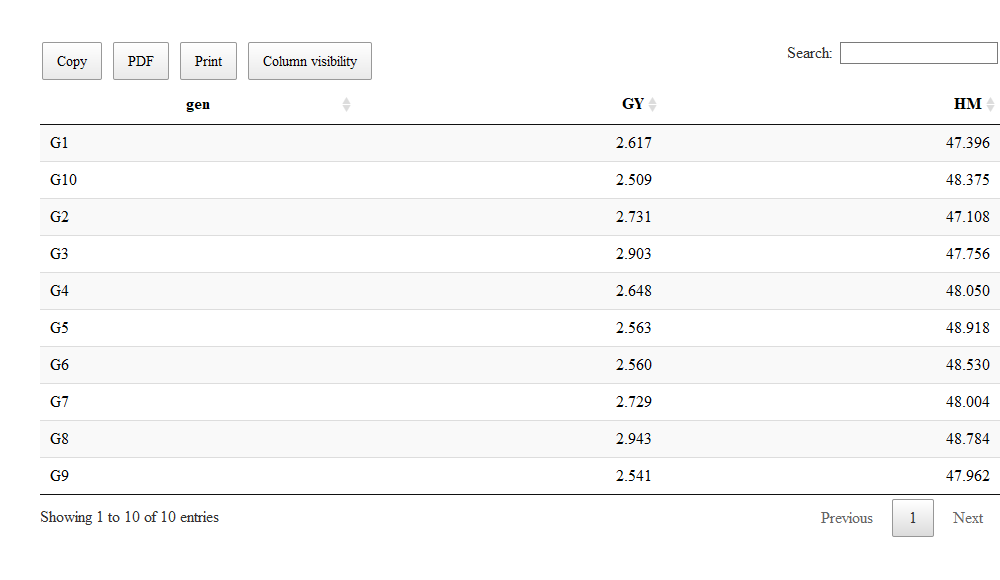

### README-unnamed-chunk-8-1.png

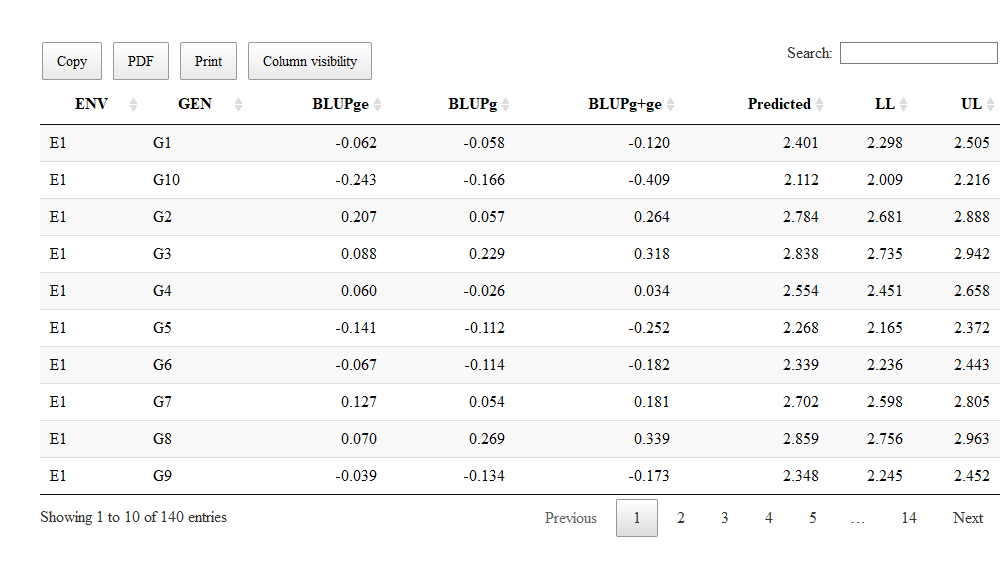

### README-unnamed-chunk-9-1.png

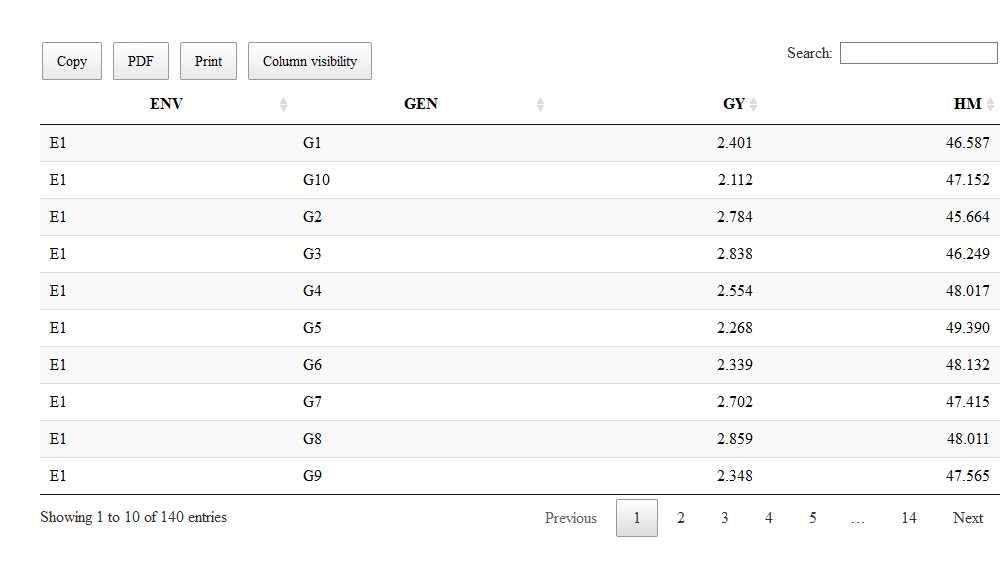

### README-unnamed-chunk-10-1.png

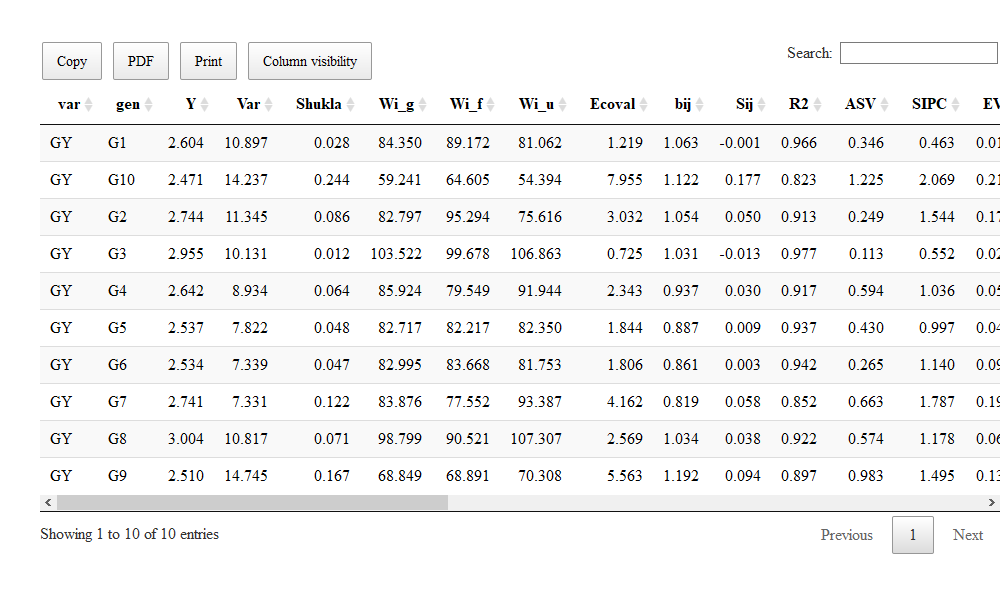

### README-unnamed-chunk-10-2.png

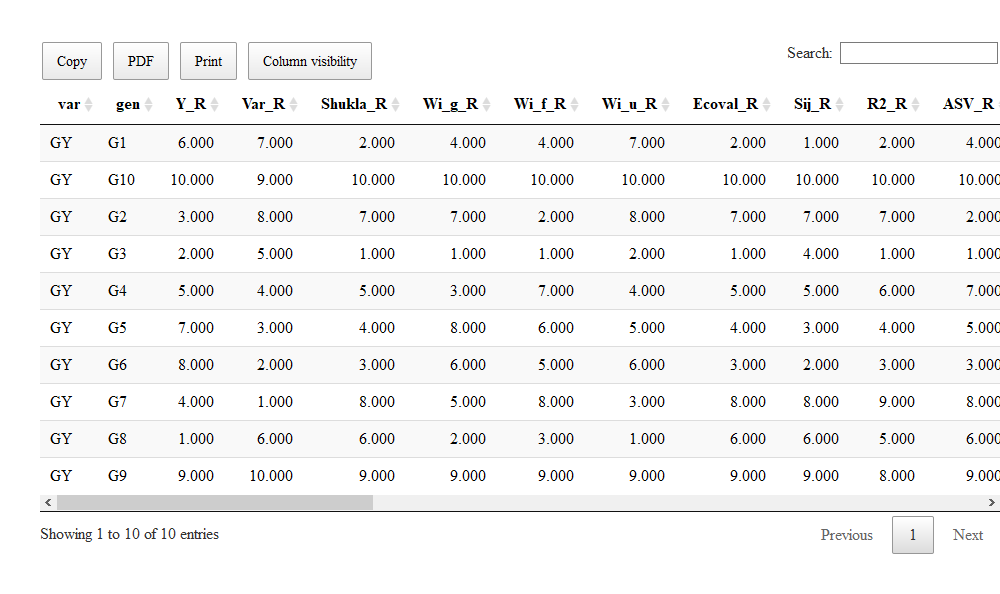

### validation.png

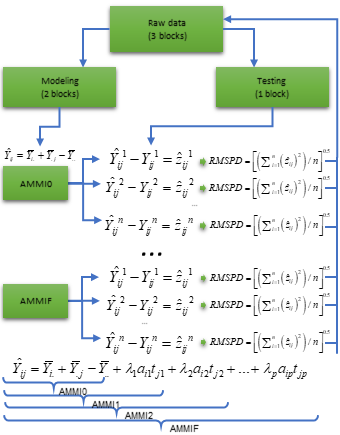
