## Appendix S1 for "metan: an R package for multi-environment trial analysis"

#### Contents

**1 An overview on scientific production related to multi-environment trials** **4**

**2 About `metan`** **5**

**3 Installing `metan`** **5**

**4 Example data** **6**

**5 Helper functions** **6**

**11 References** **110**

### 1 An overview on scientific production related to multi-environment trials

Aiming at understanding the dynamics of publications related to multi-environment trials analysis in the last half-century we carried out a bibliometric survey in the SCOPUS database using the R package `bibliometrix` (Aria & Cuccurullo, 2017).

We found 6590 documents published between 1969–2019 in 902 sources by 19.351 authors. The maximum anual scientific production was observed in 2017 (515 documents). The observed growth rate of scientific production was 11.22%, while the average total citations per doccument showed a clear and linear decrease after 2004.

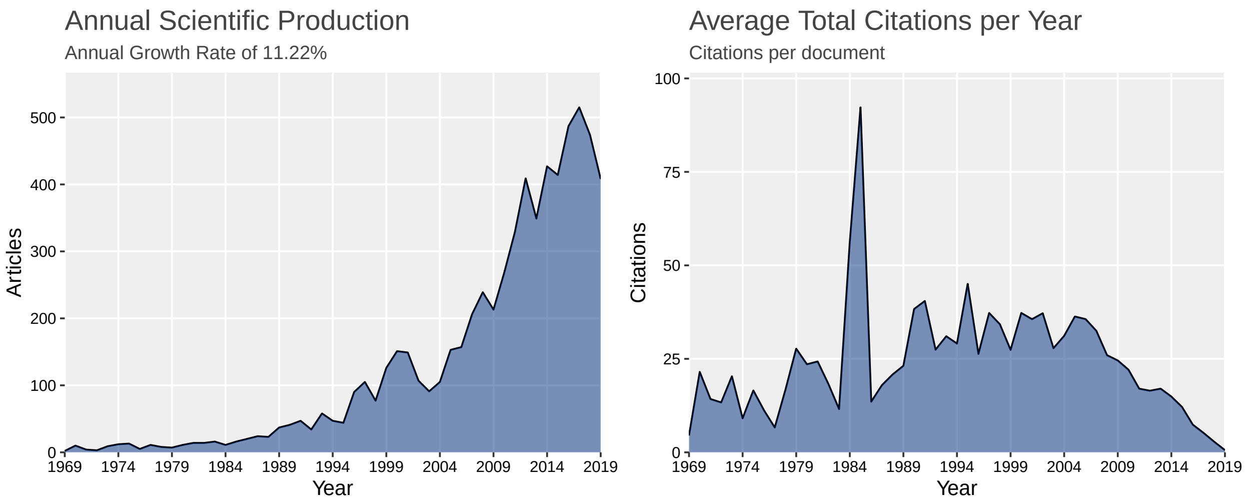

United States of America (USA) ranked the top as the most productive country with 780 documents published, followed by Brazil (371) and India (338). The USA was also the country with the higher number of citations, while Philippines was the country with the higher number of citations per document.

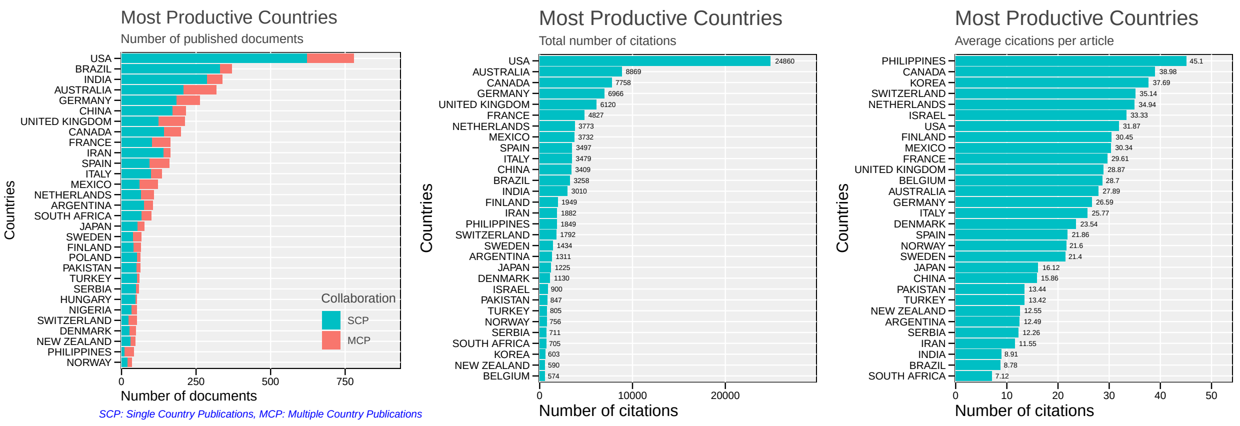

The top 50 key-words are shown in the Figure below. *"GENOTYPE BY ENVIRONMENT INTERACTION"*, *"GENOTYPE ENVIRONMENT INTERACTION"* and *"STABILITY"* was the terms most frequently used as key-words in the published articles, with more than 400 occurrences.

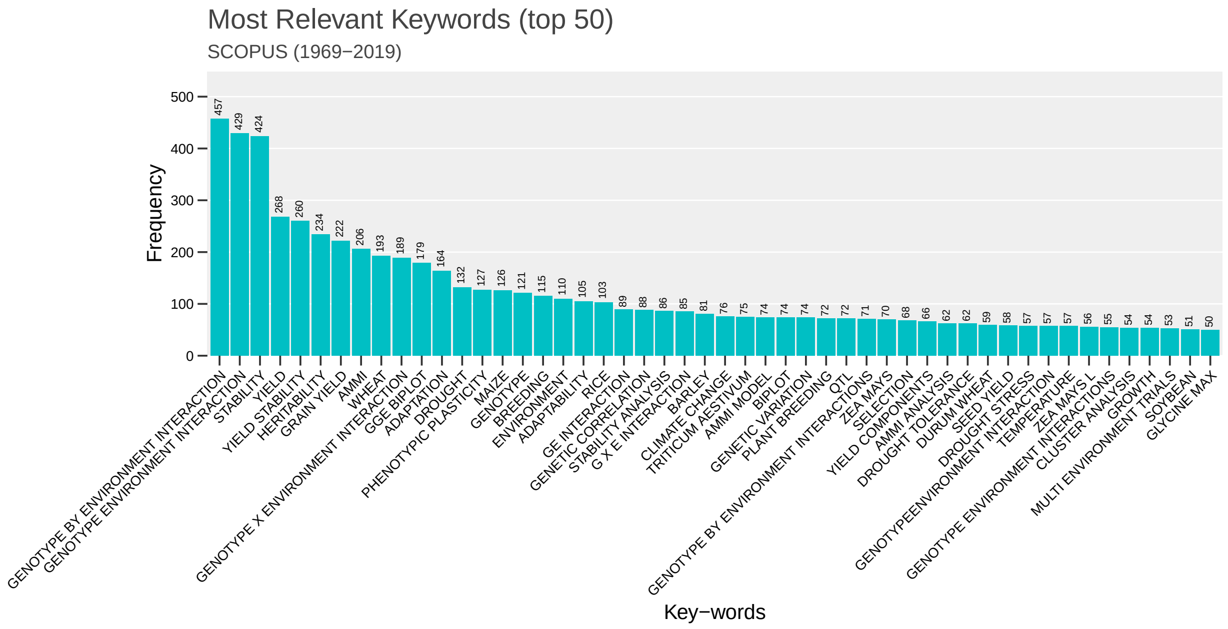

#### 2 About `metan`

`metan` is an R package that provides a collection of functions to analyze data from multi-environment trials, with a special focus on plant breeding. This document contains code to reproduce the figures used in our manuscript: **"metan: an R package for multi-environment trial analysis"**. Please note that a lot of the information presented below is also on our online vignette, accessible at https://tiagoolivoto.github.io/metan/. The online documents are updated regularly and may contain information not shown here.

#### 3 Installing `metan`

To install the released version of `metan` from CRAN type:

```
install.packages("metan")
```

The latest development version of `metan` can be installed from the GitHub repository. The installation process requires the `devtools` package, which needs to be installed first. If you are a Windows user, you should also first download and install the latest version of Rtools.

```
if(!require(devtools)) install.packages("devtools")
```

After `devtools` properly installed, you can install `metan` by running the following code. Please, note that the installation will also download the dependencies required to run the package.

```
devtools::install_github("TiagoOlivoto/metan")
```

Them, load `metan` by running

```
library(metan)
```

### 4 Example data

To explore `metan` capabilities we have provided example data that can be used immediately once the package is loaded. These data are: `data_alpha`, `data_g`, `data_ge`, `data_ge2`, `int.effects`, and `meansGxE`. We can obtain more details by clicking in the link for each data or typing, for example, `?data_g` in the R console.

### 5 Helper functions

#### 5.1 Utilities for rows and columns

##### 5.1.1 Selecting or removing columns and rows

The functions `select_cols()` and `select_rows()` can be used to select columns and rows, respectively from a data frame. Note that we use `head()` to limit the output to six rows only. Let's check it out.

```
select_cols(data_ge2, ENV, GEN) %>% head()
```

```
# A tibble: 6 x 2
  ENV   GEN
  <fct> <fct>
1 A1    H1
2 A1    H1
3 A1    H1
4 A1    H10
5 A1    H10
6 A1    H10
```

Numeric columns can be selected quickly by using the function `select_numeric_cols()`. Non-numeric columns are selected with `select_non_numeric_cols()`.

```
select_numeric_cols(data_ge2) %>% head()
```

```
# A tibble: 6 x 15
     PH    EH    EP    EL    ED    CL    CD    CW    KW    NR   NKR  CDED  PERK
  <dbl> <dbl> <dbl> <dbl> <dbl> <dbl> <dbl> <dbl> <dbl> <dbl> <dbl> <dbl> <dbl>
1  2.61  1.71 0.658  16.1  52.2  28.1  16.3  25.1  217.  15.6  36.6 0.538  89.6
2  2.87  1.76 0.628  14.2  50.3  27.6  14.5  21.4  184.  16    31.4 0.551  89.5
```

```
3  2.68  1.58 0.591  16.0  50.7  28.4  16.4  24.0  208.  17.2  31.8 0.561  89.7
4  2.83  1.64 0.581  16.7  54.1  31.7  17.4  26.2  194.  15.6  32.8 0.586  87.9
5  2.79  1.71 0.616  14.9  52.7  32.0  15.5  20.7  176.  17.6  28   0.607  89.7
6  2.72  1.51 0.554  16.7  52.7  30.4  17.5  26.8  207.  16.8  32.8 0.577  88.5
# ... with 2 more variables: TKW <dbl>, NKE <dbl>
```

```
select_non_numeric_cols(data_ge2) %>% head()
```

```
# A tibble: 6 x 3
  ENV   GEN   REP
  <fct> <fct> <fct>
1 A1    H1    1
2 A1    H1    2
3 A1    H1    3
4 A1    H10   1
5 A1    H10   2
6 A1    H10   3
```

To remove columns or rows, use `remove_cols()` and `remove_rows()`.

```
remove_cols(data_ge2, ENV, GEN) %>% head()
```

```
# A tibble: 6 x 16
  REP      PH    EH    EP    EL    ED    CL    CD    CW    KW    NR   NKR  CDED
  <fct> <dbl> <dbl> <dbl> <dbl> <dbl> <dbl> <dbl> <dbl> <dbl> <dbl> <dbl> <dbl>
1 1      2.61  1.71 0.658  16.1  52.2  28.1  16.3  25.1  217.  15.6  36.6 0.538
2 2      2.87  1.76 0.628  14.2  50.3  27.6  14.5  21.4  184.  16    31.4 0.551
3 3      2.68  1.58 0.591  16.0  50.7  28.4  16.4  24.0  208.  17.2  31.8 0.561
4 1      2.83  1.64 0.581  16.7  54.1  31.7  17.4  26.2  194.  15.6  32.8 0.586
5 2      2.79  1.71 0.616  14.9  52.7  32.0  15.5  20.7  176.  17.6  28   0.607
6 3      2.72  1.51 0.554  16.7  52.7  30.4  17.5  26.8  207.  16.8  32.8 0.577
# ... with 3 more variables: PERK <dbl>, TKW <dbl>, NKE <dbl>
```

Since `metan` allows –in most of its functions– analyzing multiple variables at the same time, select helpers provided by non-standard evaluation can be used for selecting variables that match an expression. This means that we can use a function to select variables instead of typing its own names. `metan` reexports the tidy select helpers and implements three own select helpers based on operations with prefixes and suffixes: `difference_var()` for selecting variables that startwith a prefix **AND NOT** end with a suffix. `intersect_var()`, for selecting variables that starts with a prefix **AND** end with a suffix, and `union_var()` for selecting variables that starts with a prefix **OR** ends with a suffix.

Select helpers can be used into functions in the argument `resp` or as argument of the function `select_cols()`.

- **Selecting variables that start with a prefix.**

If we want to select the variables that start with "N", we can use:

```
select_cols(data_ge2, starts_with("C")) %>% head()
```

```
# A tibble: 6 x 4
     CL    CD    CW  CDED
  <dbl> <dbl> <dbl> <dbl>
1  28.1  16.3  25.1 0.538
2  27.6  14.5  21.4 0.551
3  28.4  16.4  24.0 0.561
4  31.7  17.4  26.2 0.586
5  32.0  15.5  20.7 0.607
6  30.4  17.5  26.8 0.577
```

but if we want to select the ones that don't start with "C" we simply add "-" just before `starts_with()`.

```
select_cols(data_ge2, -starts_with("C")) %>% head()
```

```
# A tibble: 6 x 14
  ENV   GEN   REP      PH    EH    EP    EL    ED    KW    NR   NKR  PERK   TKW
  <fct> <fct> <fct> <dbl> <dbl> <dbl> <dbl> <dbl> <dbl> <dbl> <dbl> <dbl> <dbl>
1 A1    H1    1      2.61  1.71 0.658  16.1  52.2  217.  15.6  36.6  89.6  418.
2 A1    H1    2      2.87  1.76 0.628  14.2  50.3  184.  16    31.4  89.5  361.
3 A1    H1    3      2.68  1.58 0.591  16.0  50.7  208.  17.2  31.8  89.7  367.
4 A1    H10   1      2.83  1.64 0.581  16.7  54.1  194.  15.6  32.8  87.9  374.
5 A1    H10   2      2.79  1.71 0.616  14.9  52.7  176.  17.6  28    89.7  347.
6 A1    H10   3      2.72  1.51 0.554  16.7  52.7  207.  16.8  32.8  88.5  394.
# ... with 1 more variable: NKE <dbl>
```

- **Selecting variables that end with a suffix.**

Similarly, if we want to select the variables that end with "D", we can use:

```
select_cols(data_ge2, ends_with("D")) %>% head()
```

```
# A tibble: 6 x 3
     ED    CD  CDED
  <dbl> <dbl> <dbl>
1  52.2  16.3 0.538
2  50.3  14.5 0.551
3  50.7  16.4 0.561
4  54.1  17.4 0.586
5  52.7  15.5 0.607
6  52.7  17.5 0.577
```

- **Selecting variables that start with a prefix *AND* end with a suffix.**

Now, if we want to select variables that start with "C" and end with "D", i.e., the intersection between start letter "C" and end letter "D" we can:

```
select_cols(data_ge2, intersect_var("C", "D")) %>% head()
```

```
# A tibble: 6 x 2
     CD  CDED
  <dbl> <dbl>
1  16.3 0.538
2  14.5 0.551
3  16.4 0.561
4  17.4 0.586
5  15.5 0.607
6  17.5 0.577
```

- **Selecting variables that start with a prefix *OR* end with a suffix.**

We can also get the union between start letter "C" and end letter "D", i.e., variables that start with "C" or end with "D".

```
select_cols(data_ge2, union_var("C", "D")) %>% head()
```

```
# A tibble: 6 x 5
     CL    CD    CW  CDED    ED
  <dbl> <dbl> <dbl> <dbl> <dbl>
1  28.1  16.3  25.1 0.538  52.2
2  27.6  14.5  21.4 0.551  50.3
3  28.4  16.4  24.0 0.561  50.7
4  31.7  17.4  26.2 0.586  54.1
5  32.0  15.5  20.7 0.607  52.7
6  30.4  17.5  26.8 0.577  52.7
```

- **Selecting variables that start with a prefix *AND NOT* end with a suffix.**

We can also get the difference between start letter "C" and end letter "D", i.e., variables that start with "C" and not end with "D".

```
select_cols(data_ge2, difference_var("C", "D")) %>% head()
```

```
# A tibble: 6 x 2
     CL    CW
  <dbl> <dbl>
1  28.1  25.1
```

```
2   27.6  21.4
3   28.4  24.0
4   31.7  26.2
5   32.0  20.7
6   30.4  26.8
```

- **Selecting variables that contains a literal string.**

If variables in the data set have a pattern that differences between a group of variables, we can use the following code to select variables with a pattern. First we will mutate the names of the variables "PH", "EH", "EP", and "EL by including"_PLANT" to indicate that their are plant–related variables. Then we will select these variables with the function `contains()`.

```
data_vars <- data_ge2 %>%
  rename(PH_PLANT = PH,
         EH_PLANT = EH,
         EP_PLANT = EP,
         EL_PLANT = EL)
names(data_vars)
```

```
 [1] "ENV"      "GEN"      "REP"      "PH_PLANT" "EH_PLANT" "EP_PLANT"
 [7] "EL_PLANT" "ED"       "CL"       "CD"       "CW"       "KW"
[13] "NR"       "NKR"      "CDED"     "PERK"     "TKW"      "NKE"
```

```
select_cols(data_vars, contains("PLANT")) %>% head()
```

```
# A tibble: 6 x 4
  PH_PLANT EH_PLANT EP_PLANT EL_PLANT
     <dbl>    <dbl>    <dbl>    <dbl>
1     2.61     1.71    0.658     16.1
2     2.87     1.76    0.628     14.2
3     2.68     1.58    0.591     16.0
4     2.83     1.64    0.581     16.7
5     2.79     1.71    0.616     14.9
6     2.72     1.51    0.554     16.7
```

- **Selecting variables that matches a regular expression.**

More sophisticated selections can be made by using `matches()`. Assuming that we would like to select the variables that start with "E" has the second letter between "A" and "L" and end with "T", we would use something like:

```
select_cols(data_vars, matches("^E[A-L].*T$")) %>% head()
```

```
# A tibble: 6 x 2
  EH_PLANT EL_PLANT
     <dbl>    <dbl>
1     1.71     16.1
2     1.76     14.2
3     1.58     16.0
4     1.64     16.7
5     1.71     14.9
6     1.51     16.7
```

- **Selecting the last or first variables, possibly with an offset.**

We can select the first $n$th first or last column with `select_last_col()` or `select_first_col()`. We can set the argument `offset` to $n$ to select the $n$th var from the end or from the begin.

```
select_first_col(data_vars) %>% head()
```

```
# A tibble: 6 x 1
  ENV
  <fct>
1 A1
2 A1
3 A1
4 A1
5 A1
6 A1
```

```
select_last_col(data_vars) %>% head()
```

```
# A tibble: 6 x 1
    NKE
  <dbl>
1  521.
2  494.
3  565.
4  519.
5  502.
6  525.
```

##### 5.1.2 Add columns and rows

The functions `add_cols()` and `add_rows()` can be used to add columns and rows, respectively to a data frame.

```
add_cols(data_ge,
         ROW_ID = 1:420) %>%
  head()
```

```
# A tibble: 6 x 6
  ENV   GEN   REP      GY    HM ROW_ID
  <fct> <fct> <fct> <dbl> <dbl>  <int>
1 E1    G1    1      2.17  44.9      1
2 E1    G1    2      2.50  46.9      2
3 E1    G1    3      2.43  47.8      3
4 E1    G2    1      3.21  45.2      4
5 E1    G2    2      2.93  45.3      5
6 E1    G2    3      2.56  45.5      6
```

It is also possible to add a column based on existing data. Note that the arguments `.after` and `.before` are used to select the position of the new column(s). This is particularly useful to put variables of the same category together.

```
add_cols(data_ge,
         GY2 = GY^2,
         .after = "GY") %>%
  head()
```

```
# A tibble: 6 x 6
  ENV   GEN   REP      GY   GY2    HM
  <fct> <fct> <fct> <dbl> <dbl> <dbl>
1 E1    G1    1      2.17  4.70  44.9
2 E1    G1    2      2.50  6.27  46.9
3 E1    G1    3      2.43  5.89  47.8
4 E1    G2    1      3.21 10.3   45.2
5 E1    G2    2      2.93  8.60  45.3
6 E1    G2    3      2.56  6.58  45.5
```

##### 5.1.3 Concatenating columns

The function `concatetate()` can be used to concatenate multiple columns of a data frame. It return a data frame with all the original columns in `.data` plus the concatenated variable, after the last column. To chose the position of the new variable, use the argument `.after` or `.before`, as follows.

```
concatenate(data_ge, ENV, GEN, REP, .after = "REP") %>% head()
```

```
# A tibble: 6 x 6
  ENV   GEN   REP   new_var     GY    HM
  <fct> <fct> <fct> <chr>    <dbl> <dbl>
1 E1    G1    1     E1_G1_1   2.17  44.9
```

```
2 E1    G1    2     E1_G1_2  2.50  46.9
3 E1    G1    3     E1_G1_3  2.43  47.8
4 E1    G2    1     E1_G2_1  3.21  45.2
5 E1    G2    2     E1_G2_2  2.93  45.3
6 E1    G2    3     E1_G2_3  2.56  45.5
```

To drop the existing variables and keep only the concatenated column, use the argument `drop = TRUE`. To use `concatetate()` within a given function like `add_cols()` use the argument `pull = TRUE` to pull out the results to a vector.

```
concatenate(data_ge, ENV, GEN, REP, drop = TRUE) %>% head()
```

```
# A tibble: 6 x 1
  new_var
  <chr>
1 E1_G1_1
2 E1_G1_2
3 E1_G1_3
4 E1_G2_1
5 E1_G2_2
6 E1_G2_3
```

```
concatenate(data_ge, ENV, GEN, REP, pull = TRUE) %>% head()
```

```
[1] "E1_G1_1" "E1_G1_2" "E1_G1_3" "E1_G2_1" "E1_G2_2" "E1_G2_3"
```

##### 5.1.4 Reordering columns

The function `reorder_cols()` can be used to reorder the columns of a data frame.

```
reorder_cols(data_vars, contains("PLANT"), .before = "ENV") %>% head()
```

```
# A tibble: 6 x 18
  PH_PLANT EH_PLANT EP_PLANT EL_PLANT ENV   GEN   REP      ED    CL    CD    CW
     <dbl>    <dbl>    <dbl>    <dbl> <fct> <fct> <fct> <dbl> <dbl> <dbl> <dbl>
1     2.61     1.71    0.658     16.1 A1    H1    1      52.2  28.1  16.3  25.1
2     2.87     1.76    0.628     14.2 A1    H1    2      50.3  27.6  14.5  21.4
3     2.68     1.58    0.591     16.0 A1    H1    3      50.7  28.4  16.4  24.0
4     2.83     1.64    0.581     16.7 A1    H10   1      54.1  31.7  17.4  26.2
5     2.79     1.71    0.616     14.9 A1    H10   2      52.7  32.0  15.5  20.7
6     2.72     1.51    0.554     16.7 A1    H10   3      52.7  30.4  17.5  26.8
# ... with 7 more variables: KW <dbl>, NR <dbl>, NKR <dbl>, CDED <dbl>,
#   PERK <dbl>, TKW <dbl>, NKE <dbl>
```

```r
reorder_cols(data_vars, ENV, GEN, .after = "ED") %>% head()
```

```
# A tibble: 6 x 18
  REP   PH_PLANT EH_PLANT EP_PLANT EL_PLANT    ED ENV   GEN      CL    CD    CW
  <fct>    <dbl>    <dbl>    <dbl>    <dbl> <dbl> <fct> <fct> <dbl> <dbl> <dbl>
1 1         2.61     1.71    0.658     16.1  52.2 A1    H1     28.1  16.3  25.1
2 2         2.87     1.76    0.628     14.2  50.3 A1    H1     27.6  14.5  21.4
3 3         2.68     1.58    0.591     16.0  50.7 A1    H1     28.4  16.4  24.0
4 1         2.83     1.64    0.581     16.7  54.1 A1    H10    31.7  17.4  26.2
5 2         2.79     1.71    0.616     14.9  52.7 A1    H10    32.0  15.5  20.7
6 3         2.72     1.51    0.554     16.7  52.7 A1    H10    30.4  17.5  26.8
# ... with 7 more variables: KW <dbl>, NR <dbl>, NKR <dbl>, CDED <dbl>,
#   PERK <dbl>, TKW <dbl>, NKE <dbl>
```

It is possible to put columns at first and last places quicly with `columns_to_first()` and `columns_to_last()`, respectively.

```r
column_to_first(data_ge2, NKE, NR) %>% head()
```

```
# A tibble: 6 x 18
    NKE    NR ENV   GEN   REP      PH    EH    EP    EL    ED    CL    CD    CW
  <dbl> <dbl> <fct> <fct> <fct> <dbl> <dbl> <dbl> <dbl> <dbl> <dbl> <dbl> <dbl>
1  521.  15.6 A1    H1    1      2.61  1.71 0.658  16.1  52.2  28.1  16.3  25.1
2  494.  16   A1    H1    2      2.87  1.76 0.628  14.2  50.3  27.6  14.5  21.4
3  565.  17.2 A1    H1    3      2.68  1.58 0.591  16.0  50.7  28.4  16.4  24.0
4  519.  15.6 A1    H10   1      2.83  1.64 0.581  16.7  54.1  31.7  17.4  26.2
5  502.  17.6 A1    H10   2      2.79  1.71 0.616  14.9  52.7  32.0  15.5  20.7
6  525.  16.8 A1    H10   3      2.72  1.51 0.554  16.7  52.7  30.4  17.5  26.8
# ... with 5 more variables: KW <dbl>, NKR <dbl>, CDED <dbl>, PERK <dbl>,
#   TKW <dbl>
```

##### 5.1.5 Getting levels

To get the levels and the size of the levels of a factor, the functions `get_levels()` and `get_level_size()` can be used.

```r
get_levels(data_ge, ENV)
```

```
 [1] "E1"  "E10" "E11" "E12" "E13" "E14" "E2"  "E3"  "E4"  "E5"  "E6"  "E7" 
[13] "E8"  "E9" 
```

```r
get_level_size(data_ge, ENV)
```

```
 E1 E10 E11 E12 E13 E14  E2  E3  E4  E5  E6  E7  E8  E9 
 30  30  30  30  30  30  30  30  30  30  30  30  30  30 
```

#### 5.2 Utilities for numbers and strings

##### 5.2.1 Rounding whole data frames

The function `round_cols()` round a selected column or a whole data frame to the specified number of decimal places (default 0). If no variables are informed, then all numeric variables are rounded.

```
head(data_ge2)
```

```
# A tibble: 6 x 18
  ENV   GEN   REP      PH    EH    EP    EL    ED    CL    CD    CW    KW    NR
  <fct> <fct> <fct> <dbl> <dbl> <dbl> <dbl> <dbl> <dbl> <dbl> <dbl> <dbl> <dbl>
1 A1    H1    1      2.61  1.71 0.658  16.1  52.2  28.1  16.3  25.1  217.  15.6
2 A1    H1    2      2.87  1.76 0.628  14.2  50.3  27.6  14.5  21.4  184.  16
3 A1    H1    3      2.68  1.58 0.591  16.0  50.7  28.4  16.4  24.0  208.  17.2
4 A1    H10   1      2.83  1.64 0.581  16.7  54.1  31.7  17.4  26.2  194.  15.6
5 A1    H10   2      2.79  1.71 0.616  14.9  52.7  32.0  15.5  20.7  176.  17.6
6 A1    H10   3      2.72  1.51 0.554  16.7  52.7  30.4  17.5  26.8  207.  16.8
# ... with 5 more variables: NKR <dbl>, CDED <dbl>, PERK <dbl>, TKW <dbl>,
#   NKE <dbl>
```

```
round_cols(data_ge2) %>% head()
```

```
# A tibble: 6 x 18
  ENV   GEN   REP      PH    EH    EP    EL    ED    CL    CD    CW    KW    NR
  <fct> <fct> <fct> <dbl> <dbl> <dbl> <dbl> <dbl> <dbl> <dbl> <dbl> <dbl> <dbl>
1 A1    H1    1      2.61  1.71 0.66   16.1  52.2  28.1  16.3  25.1  217.  15.6
2 A1    H1    2      2.87  1.76 0.63   14.2  50.3  27.6  14.5  21.4  184.  16
3 A1    H1    3      2.68  1.58 0.59   16.0  50.7  28.4  16.4  24.0  208.  17.2
4 A1    H10   1      2.83  1.64 0.580  16.7  54.0  31.7  17.4  26.2  194.  15.6
5 A1    H10   2      2.79  1.71 0.62   14.9  52.7  32.0  15.5  20.7  176.  17.6
6 A1    H10   3      2.72  1.51 0.55   16.7  52.7  30.4  17.5  26.8  207.  16.8
# ... with 5 more variables: NKR <dbl>, CDED <dbl>, PERK <dbl>, TKW <dbl>,
#   NKE <dbl>
```

Alternatively, select variables to round.

```
round_cols(data_ge2, PH, EP, digits = 1) %>% head()
```

```
# A tibble: 6 x 18
  ENV   GEN   REP      PH    EH    EP    EL    ED    CL    CD    CW    KW    NR
  <fct> <fct> <fct> <dbl> <dbl> <dbl> <dbl> <dbl> <dbl> <dbl> <dbl> <dbl> <dbl>
1 A1    H1    1       2.6  1.71   0.7  16.1  52.2  28.1  16.3  25.1  217.  15.6
2 A1    H1    2       2.9  1.76   0.6  14.2  50.3  27.6  14.5  21.4  184.  16
3 A1    H1    3       2.7  1.58   0.6  16.0  50.7  28.4  16.4  24.0  208.  17.2
4 A1    H10   1       2.8  1.64   0.6  16.7  54.1  31.7  17.4  26.2  194.  15.6
```

```
5 A1    H10   2       2.8  1.71   0.6  14.9  52.7  32.0  15.5  20.7  176.  17.6
6 A1    H10   3       2.7  1.51   0.6  16.7  52.7  30.4  17.5  26.8  207.  16.8
# ... with 5 more variables: NKR <dbl>, CDED <dbl>, PERK <dbl>, TKW <dbl>,
#   NKE <dbl>
```

##### 5.2.2 Extracting and replacing numbers

The functions `extract_number()`, and `replace_number()` can be used to extract or replace numbers. As an example, we will extract the number of each genotype in `data_g`. By default, the extracted numbers are put as a new variable called `new_var` after the last column of the data.

```
extract_number(data_ge, GEN, .after = "GEN") %>% head()
```

```
# A tibble: 6 x 6
  ENV   GEN   new_var REP      GY    HM
  <fct> <fct>   <dbl> <fct> <dbl> <dbl>
1 E1    G1          1 1      2.17  44.9
2 E1    G1          1 2      2.50  46.9
3 E1    G1          1 3      2.43  47.8
4 E1    G2          2 1      3.21  45.2
5 E1    G2          2 2      2.93  45.3
6 E1    G2          2 3      2.56  45.5
```

If the argument `drop` is set to `TRUE`, only the new variable is kept and all others are dropped.

```
extract_number(data_ge, GEN, drop = TRUE) %>% head()
```

```
# A tibble: 6 x 1
  new_var
    <dbl>
1       1
2       1
3       1
4       2
5       2
6       2
```

To pull out the results into a vector, use the argument `pull = TRUE`. This is particularly useful when `extract_*` or `replace_*` are used within a function like `add_cols()` .

```
extract_number(data_ge, GEN, pull = TRUE) %>% head()
```

```
[1] 1 1 1 2 2 2
```

To replace numbers of a given column with a specified replacement, use `replace_number()`. By default, numbers are replaced with `""`. The argument `drop` and `pull` can also be used, as shown above.

```
replace_number(data_ge, GEN) %>% head()
```

```
# A tibble: 6 x 6
  ENV   GEN   REP      GY    HM new_var
  <fct> <fct> <fct> <dbl> <dbl> <chr>
1 E1    G1    1      2.17  44.9 G
2 E1    G1    2      2.50  46.9 G
3 E1    G1    3      2.43  47.8 G
4 E1    G2    1      3.21  45.2 G
5 E1    G2    2      2.93  45.3 G
6 E1    G2    3      2.56  45.5 G
```

```
replace_number(data_ge,
               var = REP,
               pattern = "1",
               replacement = "Rep_1",
               new_var = R_ONE,
               .after = "REP") %>%
  head()
```

```
# A tibble: 6 x 6
  ENV   GEN   REP   R_ONE    GY    HM
  <fct> <fct> <fct> <chr> <dbl> <dbl>
1 E1    G1    1     Rep_1  2.17  44.9
2 E1    G1    2     2      2.50  46.9
3 E1    G1    3     3      2.43  47.8
4 E1    G2    1     Rep_1  3.21  45.2
5 E1    G2    2     2      2.93  45.3
6 E1    G2    3     3      2.56  45.5
```

##### 5.2.3 Extracting, replacing and removing strings

The functions `extract_string()`, and `replace_string()` are used in the same context of `extract_number()`, and `replace_number()`, but for handling with strings.

```
extract_string(data_ge, GEN, .after = "GEN") %>% head()
```

```
# A tibble: 6 x 6
  ENV   GEN   new_var REP      GY    HM
  <fct> <fct> <chr>   <fct> <dbl> <dbl>
1 E1    G1    G       1      2.17  44.9
2 E1    G1    G       2      2.50  46.9
3 E1    G1    G       3      2.43  47.8
4 E1    G2    G       1      3.21  45.2
5 E1    G2    G       2      2.93  45.3
6 E1    G2    G       3      2.56  45.5
```

To replace strings, we can use the functionreplace_strings().

```
replace_string(data_ge,
               var = GEN,
               new_var = GENOTYPE,
               replacement = "GENOTYPE_",
               .after = "GEN") %>%
  head()
```

```
# A tibble: 6 x 6
  ENV   GEN   GENOTYPE   REP      GY    HM
  <fct> <fct> <chr>      <fct> <dbl> <dbl>
1 E1    G1    GENOTYPE_1 1      2.17  44.9
2 E1    G1    GENOTYPE_1 2      2.50  46.9
3 E1    G1    GENOTYPE_1 3      2.43  47.8
4 E1    G2    GENOTYPE_2 1      3.21  45.2
5 E1    G2    GENOTYPE_2 2      2.93  45.3
6 E1    G2    GENOTYPE_2 3      2.56  45.5
```

To remove all strings of a data frame, use remove_strings().

```
remove_strings(data_ge)
```

```
# A tibble: 420 x 5
     ENV   GEN   REP    GY    HM
   <dbl> <dbl> <dbl> <dbl> <dbl>
 1     1     1     1  2.17  44.9
 2     1     1     2  2.50  46.9
 3     1     1     3  2.43  47.8
 4     1     2     1  3.21  45.2
 5     1     2     2  2.93  45.3
 6     1     2     3  2.56  45.5
 7     1     3     1  2.77  46.7
 8     1     3     2  3.62  43.2
 9     1     3     3  2.28  47.8
10     1     4     1  2.36  47.9
# ... with 410 more rows
```

#### 5.3 Splitting data frames

Helper functions in `metan` are used internally to make the code cleaner. One of the most used is `split_factors()`. This function splits a data frame into a list where each object is a level of a factor or combination of factors. This is particularly useful when one statistic needs to be computed for each level of a factor. The following code splits the data `data_ge()` considering each level of the factor environment (ENV). If users need to split a data frame into a list considering all combinations of factors the easiest way is by using `as.split_factors()`.

```r
g1 <- split_factors(data_ge, ENV)
```

```
Warning: Use 'keep_factors = TRUE' to keep the columns ENV GEN REP in the
grouped data.
```

```r
names(g1$dfs)
```

```
 [1] "E1"  "E10" "E11" "E12" "E13" "E14" "E2"  "E3"  "E4"  "E5"  "E6"  "E7" 
[13] "E8"  "E9" 
```

```r
g2 <- as.split_factors(data_ge)
names(g2$dfs)[1:6]
```

```
[1] "E1 / G1 / 1"  "E1 / G1 / 2"  "E1 / G1 / 3"  "E1 / G10 / 1" "E1 / G10 / 2"
[6] "E1 / G10 / 3"
```

#### 5.4 Making two-way tables

The function `make_mat()` can be used to make a two-way table using a *"long"* format data.

```r
head(data_ge)
```

```
# A tibble: 6 x 5
  ENV   GEN   REP      GY    HM
  <fct> <fct> <fct> <dbl> <dbl>
1 E1    G1    1      2.17  44.9
2 E1    G1    2      2.50  46.9
3 E1    G1    3      2.43  47.8
4 E1    G2    1      3.21  45.2
5 E1    G2    2      2.93  45.3
6 E1    G2    3      2.56  45.5
```

```r
make_mat(data_ge, row = GEN, col = ENV, val = GY) %>% round(2)
```

```
      E1  E10  E11  E12  E13  E14   E2   E3   E4   E5   E6   E7   E8   E9
G1  2.37 2.31 1.36 1.34 3.00 1.53 3.04 4.08 3.49 4.17 2.81 1.90 2.27 2.78
G10 1.97 1.54 0.90 1.02 1.83 1.86 3.15 4.11 4.27 3.37 2.48 2.24 2.70 3.15
G2  2.90 2.30 1.49 1.99 3.03 1.43 3.23 4.57 3.72 3.83 2.54 1.99 2.05 3.36
G3  2.89 2.34 1.57 1.76 3.47 2.06 3.61 4.13 4.13 4.13 2.98 2.16 2.85 3.29
G4  2.59 2.17 1.37 1.53 2.64 1.86 3.19 3.85 3.30 3.78 2.70 1.98 2.30 3.72
G5  2.19 2.14 1.33 1.69 2.57 1.78 3.14 3.74 3.38 3.47 2.43 1.66 2.71 3.30
G6  2.30 2.21 1.50 1.39 2.91 1.80 3.29 3.43 3.40 3.57 2.34 1.76 2.54 3.04
G7  2.77 2.44 1.36 1.95 3.18 1.94 2.61 4.10 3.02 4.05 2.67 2.55 2.58 3.14
G8  2.90 2.57 1.68 2.00 3.52 1.99 3.44 4.11 4.14 4.81 2.91 2.26 2.88 2.83
G9  2.33 1.74 1.13 1.41 2.95 1.57 3.09 4.51 3.90 3.93 2.77 1.39 2.49 1.94
```

#### 5.5 Dealing with matrices

The functions `make_upper_tri()` and `make_lower_tri()` can be used to produce upper or lower triangular matrices, while `make_sym` produces a symmetric matrix.

```
cor_mat <- cor(data_ge2[, 4:8])

# Upper triangular
upp_tri <- make_upper_tri(cor_mat)
upp_tri
```

```
   PH        EH        EP        EL        ED
PH NA 0.9318282 0.6384123 0.3801960 0.6613148
EH NA        NA 0.8695460 0.3626537 0.6302561
EP NA        NA        NA 0.2634237 0.4580196
EL NA        NA        NA        NA 0.3851451
ED NA        NA        NA        NA        NA
```

```
# Lower triangular
low_tri <- make_lower_tri(cor_mat)
low_tri
```

```
          PH        EH        EP        EL ED
PH        NA        NA        NA        NA NA
EH 0.9318282        NA        NA        NA NA
EP 0.6384123 0.8695460        NA        NA NA
EL 0.3801960 0.3626537 0.2634237        NA NA
ED 0.6613148 0.6302561 0.4580196 0.3851451 NA
```

```
# Symmetric matrix
make_sym(low_tri)
```

```
          PH        EH        EP        EL        ED
PH        NA 0.9318282 0.6384123 0.3801960 0.6613148
EH 0.9318282        NA 0.8695460 0.3626537 0.6302561
EP 0.6384123 0.8695460        NA 0.2634237 0.4580196
EL 0.3801960 0.3626537 0.2634237        NA 0.3851451
ED 0.6613148 0.6302561 0.4580196 0.3851451        NA
```

The function `reorder_cormat()` can be used to reorder a correlation matrix according to the correlation coefficient by using hclust for hierarchical clustering order. This is particularly useful to identify the hidden pattern of correlations in a heat map.

```
cor_mat
```

```
          PH        EH        EP        EL        ED
PH 1.0000000 0.9318282 0.6384123 0.3801960 0.6613148
EH 0.9318282 1.0000000 0.8695460 0.3626537 0.6302561
EP 0.6384123 0.8695460 1.0000000 0.2634237 0.4580196
EL 0.3801960 0.3626537 0.2634237 1.0000000 0.3851451
ED 0.6613148 0.6302561 0.4580196 0.3851451 1.0000000
```

```
reorder_cormat(cor_mat)
```

```
          EL        ED        EP        PH        EH
EL 1.0000000 0.3851451 0.2634237 0.3801960 0.3626537
ED 0.3851451 1.0000000 0.4580196 0.6613148 0.6302561
EP 0.2634237 0.4580196 1.0000000 0.6384123 0.8695460
PH 0.3801960 0.6613148 0.6384123 1.0000000 0.9318282
EH 0.3626537 0.6302561 0.8695460 0.9318282 1.0000000
```

#### 5.6 Harmonic and geometric means

The functions `hm_mean()` and `gm_mean()` are useful for computing harmonic and geometric means, respectively. Both work with data frames by computing the means for each numeric variable. It is also possible to select variables using a comma-separated vector of unquoted variable names.

```
num <- c(1:20, 30, 50)
mean(num)
```

```
[1] 13.18182
```

```
gm_mean(num)
```

```
[1] 9.552141
```

```
hm_mean(num)
```

```
[1] 6.025626
```

```
hm_mean(data_ge2) %>% round(2)
```

```
    PH     EH     EP     EL     ED     CL     CD     CW     KW     NR    NKR
  2.44   1.28   0.53  15.06  49.38  28.83  15.88  23.05 166.32  15.96  31.88
  CDED   PERK    TKW    NKE
  0.58  87.39 331.66 501.45
```

```
gm_mean(data_ge2, EP, EL, CL) %>% round(2)
```

```
   EP    EL    CL
 0.53 15.11 28.92
```

#### 5.7 Pairwise combinations of variables

The function `comb_vars()` generates pairwise combinations of variables that will be the result of a function (defaults to sum) applied to each combination.

```
data <- data.frame(A = runif(n = 3, min = 3, max = 30),
                   B = runif(n = 3, min = 1, max = 10),
                   C = runif(n = 3, min = 9, max = 90),
                   D = runif(n = 3, min = 1, max = 90),
                   E = runif(n = 3, min = 5, max = 10)) %>%
  round(0)
data
```

```
   A B  C  D E
1 18 2 42 31 6
2 13 6 83 30 7
3  8 7 68 29 9
```

```
comb_vars(data)
```

```
# A tibble: 3 x 10
    AxB   AxC   AxD   AxE   BxC   BxD   BxE   CxD   CxE   DxE
  <dbl> <dbl> <dbl> <dbl> <dbl> <dbl> <dbl> <dbl> <dbl> <dbl>
1    20    60    49    24    44    33     8    73    48    37
2    19    96    43    20    89    36    13   113    90    37
3    15    76    37    17    75    36    16    97    77    38
```

```
comb_vars(data, FUN = "*")
```

```
# A tibble: 3 x 10
    AxB   AxC   AxD   AxE   BxC   BxD   BxE   CxD   CxE   DxE
  <dbl> <dbl> <dbl> <dbl> <dbl> <dbl> <dbl> <dbl> <dbl> <dbl>
1    36   756   558   108    84    62    12  1302   252   186
2    78  1079   390    91   498   180    42  2490   581   210
3    56   544   232    72   476   203    63  1972   612   261
```

#### 5.8 Binding with missing values

The function `rbind_fill()` combines data frames by row and fills with "." (default) missing values.

```
df1 <- data.frame(v1 = c(1, 2), v2 = c(2, 3))
df2 <- data.frame(v3 = c(4, 5))
# rbind(df1, df2) return a error
rbind_fill(df1, df2)
```

```
  v1 v2 v3
1  1  2  .
2  2  3  .
3  .  .  4
4  .  .  5
```

```
rbind_fill(df1, df2, fill = NA)
```

```
  v1 v2 v3
1  1  2 NA
2  2  3 NA
3 NA NA  4
4 NA NA  5
```

#### 5.9 Rescale a continuous vector

The function `resca()` is used to rescale a variable to have specified minimum and maximum values. Users can rescale numeric vectors, variables in data frames or rescale within levels of a factor. By default, variables are rescaled to assume a range of 0-100.

```
# Numeric vector
resca(values = c(1:5))
```

```
[1]   0  25  50  75 100
```

```
data_ge %>%
resca(GY, HM, new_min = 0, new_max = 1) %>%
head()
```

```
# A tibble: 6 x 7
  ENV   GEN   REP      GY    HM GY_res HM_res
  <fct> <fct> <fct> <dbl> <dbl>  <dbl>  <dbl>
1 E1    G1    1      2.17  44.9  0.338  0.346
2 E1    G1    2      2.50  46.9  0.414  0.445
3 E1    G1    3      2.43  47.8  0.397  0.487
4 E1    G2    1      3.21  45.2  0.574  0.36
5 E1    G2    2      2.93  45.3  0.512  0.365
6 E1    G2    3      2.56  45.5  0.428  0.375
```

### 6 Check, manipulate and summarise data

#### 6.1 Inspecting data

`metan` was designed to work best with data frame objects, including objects of class `data.frame`, `data.table` and `tibble`. Once data is imported into R, we use `inspect()` to quickly scan it for common issues. This includes the number of factor variables, unbalanced data, missing values, and possible outliers. First, we will create an object `out_data` to simulate an outlier in the data `data_ge`.

```
out_data <- data_ge
out_data[34, 4] <- out_data[34, 4] * 3
inspect(out_data, plot = TRUE)
```

```
# A tibble: 5 x 9
  Variable Class   Missing Levels Valid_n   Min Median   Max Outlier
  <chr>    <fct>   <fct>   <fct>    <int> <dbl>  <dbl> <dbl>   <dbl>
1 ENV      factor  No      14         420 NA     NA    NA         NA
2 GEN      factor  No      10         420 NA     NA    NA         NA
3 REP      factor  No      3          420 NA     NA    NA         NA
4 GY       numeric No      -          420  0.67   2.61  8.35       1
5 HM       numeric No      -          420 38     48    58          0

Warning: Possible outliers in variable(s) GY. Use 'find_outliers()' for more
details.
```

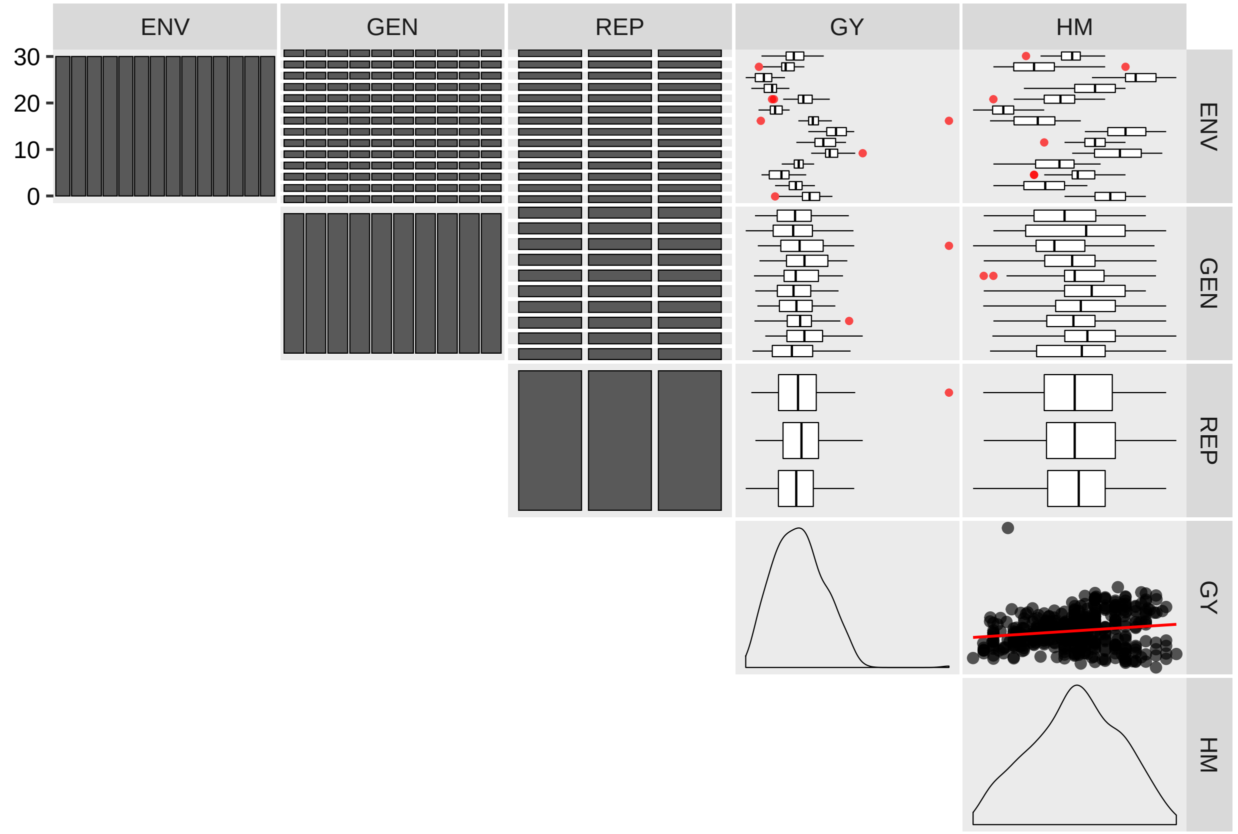

#### 6.2 Finding outliers

The results above show that possible outliers are present in the variable GY. A more in-depth check should be performed. To do that, we will use the function `find_outliers()` with the argument `plot = TRUE` that will generate the Figure 1a of the manuscript.

```
find_outliers(out_data, GY, plots = TRUE)
```

```
Warning: The factors ENV GEN REP were ignored. Use 'split_factors()' to perform
an analysis for each level of a factor.

Number of possible outliers: 1
Lines: 34
Proportion: 0.2%
Mean of the outliers: 8.352
Maximum of the outliers: 8.352  | Line 34
Minimum of the outliers: 8.352  | Line 34
With outliers:    mean = 2.687 | CV = 35.897%
Without outliers: mean = 2.674 | CV = 34.6%
```

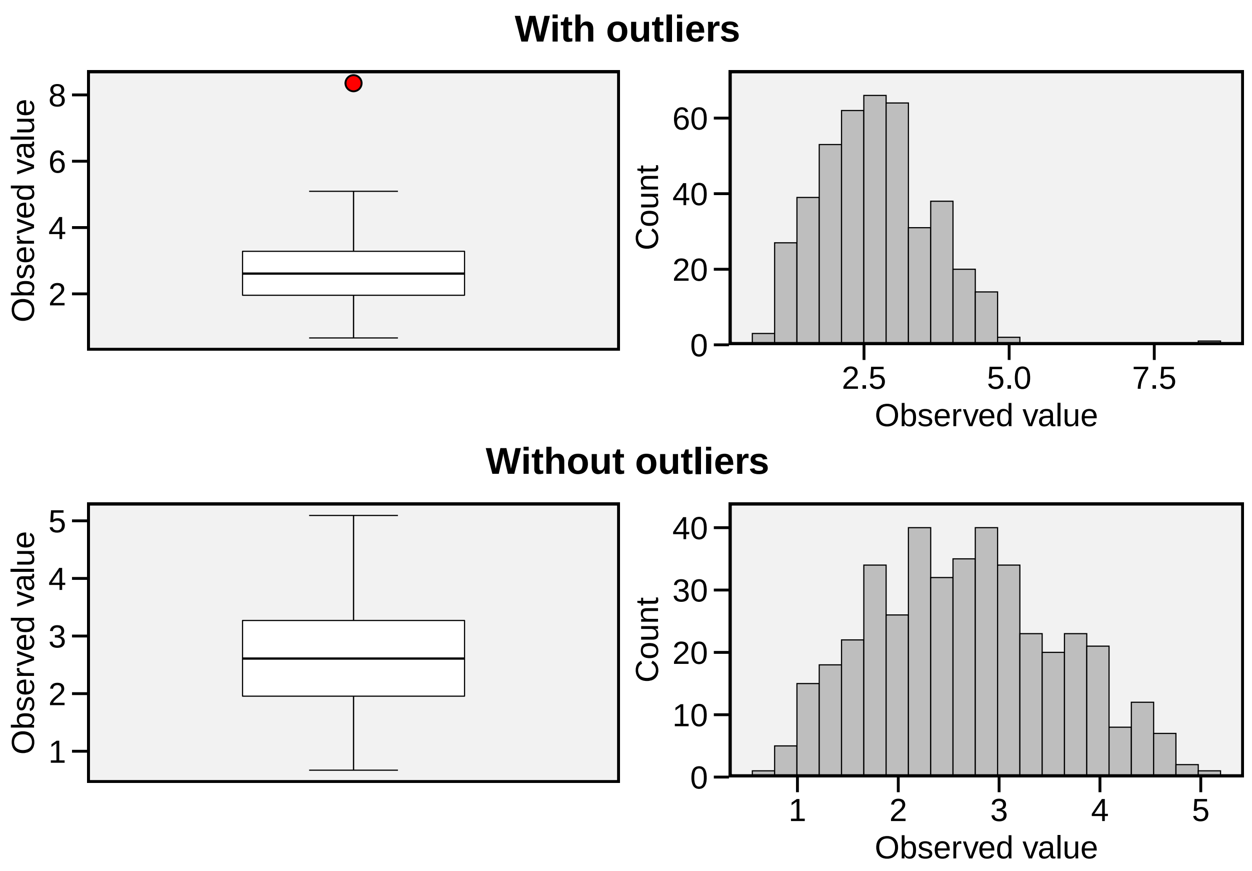

The results show that the possible outlier is in the line 34 and has a value of 8.352 (above 1.5 times the interquartile range of GY). The researcher should then check if this value is assumed to be an outlier and take some action or simply follow with the analysis without a guilty conscience.

**Quick tip**

To compute an outlier check for each level of a factor we can use the argument `by` that calls `split_factors()` internally. For example, to check for outliers in the variable GY in each environment of `data_ge` we should use: `find_outliers(data_ge, GY, by = ENV)`. To check for more than one grouping variable we should pass the subsetted data from `split_factors()`, e.g., `data_ge %>% split_factors(ENV, GEN) %>% find_outliers(GY)`

#### 6.3 Descriptive statistics

Here –using the data example `data_ge2`– we will compute, for each environment, seven descriptive statistics (coefficient of variation, maximum, mean, median, minimum, standard error, and standard deviation) using the function `desc_stat()`. These are the statistics that are computed by default. To compute all the statistics, use the argument `stats = "all"`. In addition, we can use a comma-separated vector with the statistic names, e.g., `stats = c("mean, cv")` for computing only chosen statistics. To facilitate the typing, the comma-separated vector of names has `"` only at the beginning and end of the vector. Note that the statistic names ARE NOTE case sensitive, i.e., both `"mean"` and `"MEAN"` are recognized.

- This will work:
  - `stats = c("mean, SE.mean, CV, max, min")`
  - `stats = "mean"`
- This will NOT work:
  - `stats = c("mean", "SE.mean", "CV", "max", "min")`
  - `stats = c("mean,SE.mean, CV, max,min")`

```
data_ge2 %>%
  desc_stat(PH, CW, NKR,
            hist = TRUE) %>%
  print()
```

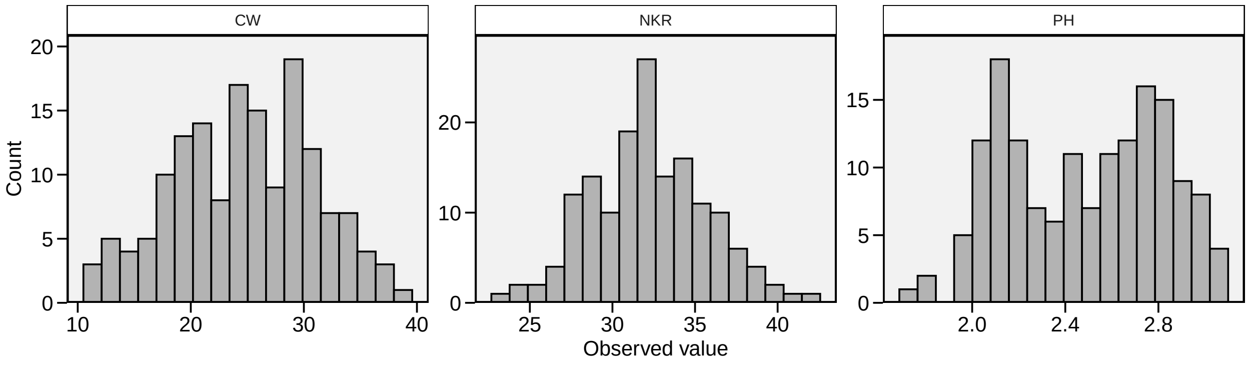

```
# A tibble: 7 x 4
  Statistic      CW    NKR      PH
  <chr>       <dbl>  <dbl>   <dbl>
1 cv         25.2   10.7   13.4
2 max        38.5   42      3.04
3 mean       24.8   32.2    2.48
4 median     24.5   32      2.52
5 min        11.1   23.2    1.71
6 se.mean     0.501  0.277  0.0267
7 var.amo    39.2   12.0    0.112
```

By default, when `split_factors()` is called internally with the argument `by`, the results are shown in a *"long"* format, with grouping levels and statistics in the rows and variables in columns. The results for each variable may be converted to a *"wide"* format using, the function `desc_wider()`.

```
stats <-
  desc_stat(data_ge2,
            PH, CW, NKR,
            by = ENV,
            verbose = F)
desc_wider(stats, PH)
```

```
# A tibble: 4 x 8
  ENV      cv   max  mean median   min se.mean var.amo
  <chr> <dbl> <dbl> <dbl>  <dbl> <dbl>   <dbl>   <dbl>
1 A1     4.42  3.04  2.79   2.78  2.55  0.0198  0.0153
2 A2    16.2   3.04  2.46   2.27  1.81  0.0639  0.159
3 A3    10.3   2.77  2.17   2.10  1.71  0.0358  0.0499
4 A4     6.61  2.81  2.52   2.47  2.11  0.0266  0.0277
```

The means of all numeric variables for each level of a factor can be computed quickly with the function `means_by()`.

```
means_by(data_ge2, ENV) %>% round_cols()
```

```
# A tibble: 4 x 16
  ENV      PH    EH    EP    EL    ED    CL    CD    CW    KW    NR   NKR  CDED
  <fct> <dbl> <dbl> <dbl> <dbl> <dbl> <dbl> <dbl> <dbl> <dbl> <dbl> <dbl> <dbl>
1 A1     2.79  1.58 0.570  15.6  51.6  29.7  16.4  28.3  199.  16.9  33.9 0.580
2 A2     2.46  1.31 0.53   15.2  48.7  28.5  15.9  23.8  168.  15.8  32.3 0.580
3 A3     2.17  1.08 0.5    14.7  47.9  28.4  15.8  20.8  147.  15.8  30.4 0.59
4 A4     2.52  1.41 0.56   15.1  49.9  29.4  15.8  26.4  177.  16.0  32.4 0.59
# ... with 3 more variables: PERK <dbl>, TKW <dbl>, NKE <dbl>
```

```
means_by(data_ge2, ENV, GEN) %>% round_cols()
```

```
# A tibble: 52 x 17
   ENV   GEN      PH    EH    EP    EL    ED    CL    CD    CW    KW    NR   NKR
   <fct> <fct> <dbl> <dbl> <dbl> <dbl> <dbl> <dbl> <dbl> <dbl> <dbl> <dbl> <dbl>
 1 A1    H1     2.72  1.68 0.63   15.4  51.1  28.0  15.7  23.5  203.  16.3  33.3
 2 A1    H10    2.78  1.62 0.580  16.1  53.2  31.4  16.8  24.6  192.  16.7  31.2
 3 A1    H11    2.75  1.58 0.570  16.6  48.9  29.0  17.2  23.6  188.  15.2  34.6
 4 A1    H12    2.69  1.54 0.580  15.2  50.0  29.8  15.6  25.6  180.  17.3  32.7
 5 A1    H13    2.77  1.58 0.570  14.8  53.5  31.2  15.4  31.3  219.  18.7  32.9
 6 A1    H2     2.79  1.34 0.48   15.0  51.9  26.6  15.5  20.2  204.  19.2  33.5
 7 A1    H3     2.94  1.59 0.54   15.4  51.8  28    16.4  24.4  198.  18.5  34.1
 8 A1    H4     2.87  1.68 0.580  16.0  50.7  27.4  16.7  27.7  202.  16.4  38.3
 9 A1    H5     2.83  1.59 0.56   15.8  49.8  28.3  16.7  32.1  193.  14.5  37.4
10 A1    H6     2.77  1.57 0.570  16.7  54.1  31.7  17.6  35.8  232.  16.8  35.5
# ... with 42 more rows, and 4 more variables: CDED <dbl>, PERK <dbl>,
#   TKW <dbl>, NKE <dbl>
```

#### 6.4 Manipulating data

In MET analysis sometimes we need to convert a *"long"* data to a typical two-way table with genotypes in rows and environments in columns. If you want to do that quickly, then `make_mat()` is what you're looking for. Let's check it out.

```
twm <- make_mat(data_ge2, GEN, ENV, PH)
twm
```

```
          A1       A2       A3       A4
H1  2.722667 2.930000 2.197333 2.635333
H10 2.783333 2.049333 2.038000 2.388667
H11 2.748667 2.146667 2.101333 2.563333
H12 2.692667 2.093333 2.430000 2.534000
H13 2.772000 2.227333 2.598667 2.556667
H2  2.792000 2.946667 2.154000 2.524000
H3  2.935333 2.940667 2.043333 2.460133
H4  2.868667 2.846000 2.049333 2.556000
H5  2.834667 2.704667 2.094667 2.638000
H6  2.768667 2.824667 2.153333 2.485333
H7  2.773867 2.136667 2.177333 2.529333
H8  2.692000 1.959333 2.100000 2.554933
H9  2.925333 2.199333 2.028000 2.304667
```

To convert a two-way table to a *"long"* format use the function `make_long()`.

```
make_long(twm)
```

```
# A tibble: 52 x 3
   GEN   ENV       Y
   <chr> <chr> <dbl>
 1 H1    A1     2.72
 2 H1    A2     2.93
 3 H1    A3     2.20
 4 H1    A4     2.64
 5 H10   A1     2.78
 6 H10   A2     2.05
 7 H10   A3     2.04
 8 H10   A4     2.39
 9 H11   A1     2.75
10 H11   A2     2.15
# ... with 42 more rows
```

> **Quick tip**
>
> If a typical MET data (with replicates) is used, then the argument `fun` in `make_mat()` can be used to compute any statistic based on the replicate's values. For example, to show the maximum value among replicates of PH for each cell of the two-way table we should use:
> `max_twm <- make_mat(data_ge2, GEN, ENV, PH, fun = max)`

### 7 Analyzing individual environments

A within-environment ANOVA considering a fixed-effect model is computed with the function `anova_ind()`. For each environment the Mean Squares for block, genotypes and error are shown. Estimated F-value and the probability error are also shown for block and genotype effects. Some measures of experimental precision are calculated, namelly, coefficient of variation, $CV = (\sqrt{MS_{res}}/Mean) \times 100$; the heritability, $h2 = (MS_{gen} - MS_{res})/MS_{gen}$, and the accuracy of selection, $As = \sqrt{h2}$.

```
indiv <- anova_ind(data_ge, ENV, GEN, REP, GY, verbose = FALSE)
print(indiv$GY$individual)
```

```
# A tibble: 14 x 12
   ENV    MEAN    MSB   MSG    MSR    FCB    PRFB   FCG    PRFG    CV    h2
   <chr> <dbl>  <dbl> <dbl>  <dbl>  <dbl>   <dbl> <dbl>   <dbl> <dbl> <dbl>
 1 E1     2.52 0.0652 0.337 0.144   0.453 6.43e-1  2.34 5.94e-2 15.1  0.573
 2 E10    2.18 0.654  0.296 0.0267 24.5   7.28e-6 11.1  1.10e-5  7.51 0.910
 3 E11    1.37 0.377  0.151 0.105   3.59  4.86e-2  1.44 2.44e-1 23.7  0.304
 4 E12    1.61 0.0919 0.320 0.0535  1.72  2.08e-1  5.98 6.47e-4 14.4  0.833
 5 E13    2.91 0.0767 0.713 0.0994  0.772 4.77e-1  7.18 2.10e-4 10.8  0.861
 6 E14    1.78 0.104  0.131 0.0753  1.37  2.78e-1  1.73 1.53e-1 15.4  0.423
 7 E2     3.18 0.698  0.207 0.179   3.91  3.88e-2  1.16 3.76e-1 13.3  0.136
 8 E3     4.06 0.489  0.335 0.179   2.73  9.21e-2  1.87 1.23e-1 10.4  0.466
 9 E4     3.68 0.116  0.531 0.138   0.846 4.46e-1  3.86 7.12e-3 10.1  0.741
10 E5     3.91 0.219  0.526 0.0664  3.30  6.02e-2  7.93 1.10e-4  6.59 0.874
11 E6     2.66 0.160  0.135 0.0586  2.73  9.22e-2  2.30 6.35e-2  9.09 0.565
12 E7     1.99 0.381  0.337 0.0910  4.19  3.22e-2  3.70 8.73e-3 15.2  0.730
13 E8     2.54 0.817  0.215 0.0278 29.4   2.15e-6  7.72 1.31e-4  6.57 0.870
14 E9     3.06 0.583  0.679 0.111   5.25  1.60e-2  6.12 5.62e-4 10.9  0.837
# ... with 1 more variable: AS <dbl>
```

The function `gamem()` can be used to analyze single experiments (one-way experiments) using a mixed-effect model according to the following model:

$$y_{ij} = \mu + \alpha_i + \tau_j + \varepsilon_{ij}$$

where $y_{ij}$ is the value observed for the $i$th genotype in the $j$th replicate ($i = 1, 2, \ldots$ $g$; $j = 1, 2,$ .., $r$); being $g$ and $r$ the number of genotypes and replicates, respectively; $\alpha_i$ is the random effect of the $i$th genotype; $\tau_j$ is the fixed effect of the $j$th replicate; and $\varepsilon_{ij}$ is the random error associated to $y_{ij}$. In this example, we will use the example data data_g from metan package.

```
gen_mod <- gamem(data_g, GEN, REP,
                 resp = c(ED, CL, CD, KW, TKW, NKR))
```

```
Evaluating variable ED 0 %
Evaluating variable CL 20 %
Evaluating variable CD 40 %
Evaluating variable KW 60 %
Evaluating variable TKW 80 %
Evaluating variable NKR 100 %
------------------------------------------------------------
Variables with nonsignificant genotype effect
CD NKR
------------------------------------------------------------
Done!
```

The easiest way of obtaining the results of the model above is by using the function get_model_data(). Let's do it.

- **Likelihood ratio test for genotype effect**

```
get_model_data(gen_mod, "pval_lrt")
```

```
# A tibble: 2 x 7
  model             ED           CL      CD       KW       TKW    NKR
  <chr>          <dbl>        <dbl>   <dbl>    <dbl>     <dbl>  <dbl>
1 Complete NA          NA           NA      NA       NA        NA
2 Genotype   0.0000273  0.00000225  0.118  0.0253  0.00955  0.216
```

- **Variance components and genetic parameters**

```
get_model_data(gen_mod, "genpar")
```

```
# A tibble: 11 x 7
   Parameters     ED     CL     CD       KW       TKW    NKR
   <chr>       <dbl>  <dbl>  <dbl>    <dbl>     <dbl>  <dbl>
 1 Gen_var      5.37   4.27  0.240  181.      841.      2.15
 2 Gen (%)     68.8   75.1   27.4     39.2      45.2   21.6
 3 Res_var      2.43   1.41  0.634  280.     1018.      7.80
 4 Res (%)     31.2   24.9   72.6     60.8      54.8   78.4
 5 Phen_var     7.80   5.68  0.873  461.     1859.      9.94
```

```
 6 H2          0.688  0.751  0.274   0.392    0.452  0.216
 7 H2mg        0.869  0.901  0.532   0.659    0.712  0.452
 8 Accuracy    0.932  0.949  0.729   0.812    0.844  0.673
 9 CVg         4.84   7.26   3.10    9.16     9.13   4.82
10 CVr         3.26   4.18   5.05   11.4     10.0    9.19
11 CV ratio    1.49   1.74   0.615   0.803    0.909  0.525
```

- **Predicted means**

```
get_model_data(gen_mod, "blupg")
```

```
# A tibble: 13 x 7
   gen      ED    CL    CD    KW   TKW   NKR
   <chr> <dbl> <dbl> <dbl> <dbl> <dbl> <dbl>
 1 H1     50.2  30.7  15.8  153.  354.  29.5
 2 H10    44.4  25.1  15.5  129.  268.  31.7
 3 H11    47.2  26.6  15.6  143.  297.  31.3
 4 H12    47.8  26.1  15.2  148.  293.  30.0
 5 H13    50.3  27.4  15.9  170.  319.  31.2
 6 H2     50.3  30.0  16.3  156.  338.  29.6
 7 H3     47.2  28.6  16.1  142.  331.  30.2
 8 H4     46.1  27.8  16.2  145.  310.  31.8
 9 H5     49.8  30.1  16.2  156.  309.  31.3
10 H6     49.7  31.6  15.2  140.  325.  28.6
11 H7     48.7  30.0  15.5  153.  346.  30.0
12 H8     46.3  29.0  15.8  143.  339.  29.5
13 H9     44.4  27.0  15.8  131.  301.  30.4
```

In the above example, the experimental design was a complete randomized block. It is also possible to an experiment conducted in an alpha-lattice design with the function `gamem()`. In this case, the following model is fitted:

$$y_{ijk} = \mu + \alpha_i + \gamma_j + (\gamma\tau)_{jk} + \varepsilon_{ijk}$$

where $y_{ijk}$ is the observed value of the $i$th genotype in the $k$th block of the $j$th replicate ($i = 1$, 2, … $g$; $j = 1$, 2, .., $r$; $k = 1$, 2, .., $b$); respectively; $\alpha_i$ is the random effect of the $i$th genotype; $\gamma_j$ is the fixed effect of the $j$th complete replicate; $(\gamma\tau)_{jk}$ is the random effect of the $k$th incomplete block nested within the $j$ replicate; and $\varepsilon_{ijk}$ is the random error associated to $y_{ijk}$. In this example, we will use the example data `data_alpha` from metan package.

```
gen_alpha <- gamem(data_alpha, GEN, REP, YIELD, block = BLOCK)
```

```
Done!
```

```r
get_model_data(gen_alpha, "pval_lrt")
```

```
# A tibble: 3 x 2
  model            YIELD
  <chr>            <dbl>
1 Complete   NA
2 Genotype    0.00000118
3 rep:block  0.00335
```

```r
get_model_data(gen_alpha, "details")
```

```
# A tibble: 6 x 2
  Parameters YIELD
  <chr>      <chr>
1 Ngen       24
2 OVmean     4.4795
3 Min        2.8873 (G03 in B6 of R3)
4 Max        5.8757 (G05 in B1 of R1)
5 MinGEN     3.3431 (G03)
6 MaxGEN     5.1625 (G01)
```

```r
get_model_data(gen_alpha, "genpar")
```

```
# A tibble: 13 x 2
   Parameters      YIELD
   <chr>           <dbl>
 1 Gen_var        0.143
 2 Gen (%)       48.5
 3 rep:block_var  0.0702
 4 rep:block (%) 23.8
 5 Res_var        0.0816
 6 Res (%)       27.7
 7 Phen_var       0.295
 8 H2             0.485
 9 H2mg           0.798
10 Accuracy       0.893
11 CVg            8.44
12 CVr            6.38
13 CV ratio       1.32
```

### 8 Stability analysis

The function `ge_plot()` can be used to visualize the genotype's performance across the environments. Two types of plots can be produced by setting the argument `type`. `type = 1` (default) produces a heat map while `type = 2` produces a line plot.

```r
a <- ge_plot(data_ge, ENV, GEN, GY)
b <- ge_plot(data_ge, ENV, GEN, GY, type = 2)
arrange_ggplot(a, b, labels = letters[1:2])
```

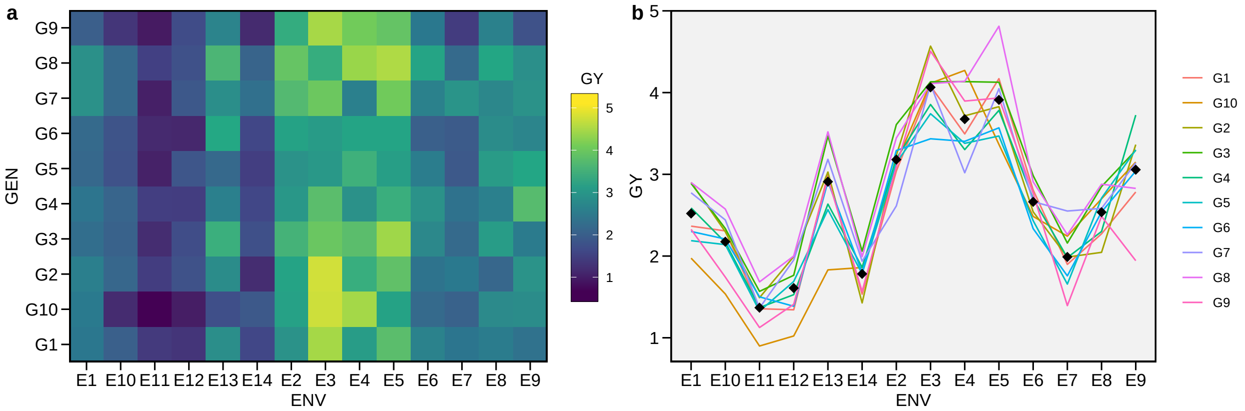

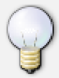

**Quick tip**

Iterative plots can be obtained using the `ggplotly()` function of the plotly package. This function converts a `ggplot2` object into an iterative plot that is shown in the *viewer* tab of the RStudio. To make `p1` iterative, we should use `plotly::ggplotly(p1)`

To identify the winner genotype within each environment, we can use the function `ge_winners()`.

```r
ge_winners(data_ge2, ENV, GEN, resp = everything())
```

```
# A tibble: 4 x 16
  ENV   PH    EH    EP    EL    ED    CL    CD    CW    KW    NR    NKR   CDED
  <fct> <chr> <chr> <chr> <chr> <chr> <chr> <chr> <chr> <chr> <chr> <chr> <chr>
1 A1    H3    H1    H1    H6    H6    H8    H6    H6    H6    H2    H4    H8
2 A2    H2    H1    H1    H6    H2    H2    H6    H2    H2    H2    H6    H13
3 A3    H13   H13   H6    H4    H13   H6    H2    H7    H13   H13   H4    H6
4 A4    H5    H5    H10   H7    H11   H5    H7    H5    H7    H11   H9    H10
# ... with 3 more variables: PERK <chr>, TKW <chr>, NKE <chr>
```

To show the ranks of all genotypes within each environment, for each variable, we se the argument `type = "ranks"` in the function `ge_winners()`.

```r
ge_winners(data_ge2, ENV, GEN, resp = everything(), type = "ranks")
```

```
# A tibble: 52 x 16
   ENV   PH    EH    EP    EL    ED    CL    CD    CW    KW    NR    NKR   CDED
   <fct> <chr> <chr> <chr> <chr> <chr> <chr> <chr> <chr> <chr> <chr> <chr> <chr>
 1 A1    H3    H1    H1    H6    H6    H8    H6    H6    H6    H2    H4    H8
 2 A1    H9    H4    H10   H11   H13   H9    H11   H8    H13   H13   H5    H9
 3 A1    H4    H9    H4    H10   H10   H6    H9    H9    H9    H3    H6    H7
 4 A1    H5    H10   H7    H4    H9    H10   H10   H7    H2    H7    H11   H12
 5 A1    H2    H7    H12   H5    H8    H13   H5    H5    H1    H12   H3    H11
 6 A1    H10   H5    H11   H9    H2    H7    H4    H13   H4    H8    H2    H10
 7 A1    H7    H3    H6    H3    H3    H12   H8    H4    H3    H6    H1    H6
 8 A1    H13   H11   H9    H7    H1    H11   H3    H12   H8    H10   H13   H13
 9 A1    H6    H13   H13   H1    H7    H5    H7    H10   H5    H4    H8    H5
10 A1    H11   H6    H5    H12   H4    H1    H1    H3    H10   H1    H12   H1
# ... with 42 more rows, and 3 more variables: PERK <chr>, TKW <chr>, NKE <chr>
```

For more details about the trials, we can use `ge_details()`.

```
ge_details(data_ge, ENV, GEN, resp = everything())
```

```
# A tibble: 10 x 3
   Parameters GY                   HM
   <chr>      <chr>                <chr>
 1 Mean       "2.67"               "48.09"
 2 Std_err    "0.92"               "4.37"
 3 Desv_pad   "0.05"               "0.21"
 4 CV         "34.56"              "9.09"
 5 Min        "0.67 (G10 in E11)"  "38 (G2 in E14)"
 6 Max        "5.09 (G8 in E5)"    "58 (G8 in E11)"
 7 MinENV     "E11 (1.37)"         "E14 (41.03)"
 8 MaxENV     "E3 (4.06)"          "E11 (54.2)"
 9 MinGEN     "G10 (2.47) "        "G2 (46.66) "
10 MaxGEN     "G8 (3) "            "G5 (49.3) "
```

The function `ge_effects()` can be used to compute genotype-environment effects ($ge_{ij}$) or genotype plus genotyp-environment effects ($gge_{ij}$), with

$$ge_{ij} = y_{ij} - \mu - \alpha_i - \tau_j gge_{ij} = y_{ij} - \mu - \tau_j$$

where $\mu$ is the grand mean, $\alpha_i$ is the effect of genotype $i$; and $\tau_j$ is the effect of the environment $j$.

```
a <- ge_effects(data_ge, ENV, GEN, REP, GY)
b <- ge_effects(data_ge, ENV, GEN, REP, HM, type = "gge")
arrange_ggplot(plot(a), plot(b), labels = letters[1:2])
```

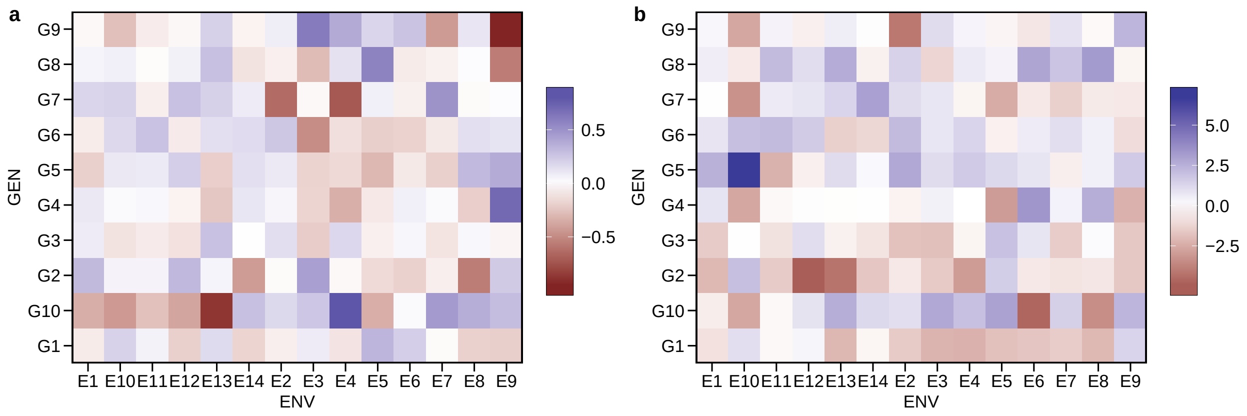

The plot shows a clear change in the rank order of genotypes across environments, which characterize a cross-over interaction. Statistically, this can be tested in a joint-ANOVA, performed with the function `anova_joint()`. This function implements the simplest and well–known linear model with interaction effect used to analyze data from multi-environment trials, namely:

$$y_{ijk} = \mu + \alpha_i + \tau_j + (\alpha\tau)_{ij} + \gamma_{jk} + \varepsilon_{ijk} \tag{8.1}$$

where $y_{ijk}$ is the response variable (e.g., grain yield) observed in the $k$th block of the $i$th genotype in the $j$th environment ($i = 1, 2, \ldots, g$; $j = 1, 2, \ldots, e$; $k = 1, 2, \ldots, b$); $\mu$ is the grand mean; $\alpha_i$ is the effect of the ith genotype; $\tau_j$ is the effect of the $j$th environment; $(\alpha\tau)_{ij}$ is the interaction effect of the $i$th genotype with the $j$th environment; $\gamma_{jk}$ is the effect of the $k$th block within the $j$th environment; and $\varepsilon_{ijk}$ is the random error.

```
joint <- anova_joint(data_ge, ENV, GEN, REP, GY, verbose = FALSE)
print(joint)
```

```
Variable GY
---------------------------------------------------------------------------
# A tibble: 8 x 6
  Source        Df `Sum Sq` `Mean Sq` `F value`   `Pr(>F)`
  <chr>      <dbl>    <dbl>     <dbl>     <dbl>      <dbl>
1 ENV        13       280.      21.5       222.    7.25e-130
2 GEN         9        13.0      1.44       14.9   2.19e- 19
3 REP(ENV)   28         9.66     0.345       3.57  3.59e-  8
4 ENV:GEN   117        31.2      0.267       2.76  1.01e- 11
5 Residuals 252        24.4      0.0967     NA    NA
6 CV(%)      11.6      NA       NA          NA    NA
7 MSR+/MSR-   6.71     NA       NA          NA    NA
8 OVmean      2.67     NA       NA          NA    NA
---------------------------------------------------------------------------
```

The genotype-vs-environment interaction was highly significant. So, it is suggested to proceed with some stability analysis to explore such interaction.

#### 8.1 ANOVA-based stability analysis

The function `Annicchiarico()` computes the known genotypic confidence index (Annicchiarico, 1992), which measures the superiority of the genotype in relation to the average of each environment, according to the following model:

$$Z_{ij} = \frac{Y_{ij}}{\bar{Y}_{.j}} \times 100$$

The genotypic confidence index of the genotype $i$ ($G_i$) is then estimated as follows:

$$G_i = Z_{i.}/e - \alpha \times sd(Z_{i.})$$

Where $e$ is the number of environments and $\alpha$ is the quantile of the standard normal distribution at a given probability error ($\alpha \approx |1.64|$ at 0.05).

```r
ann <- Annicchiarico(data_ge, ENV, GEN, REP, GY)
print(ann)
```

```
Variable GY
---------------------------------------------------------------------------
Environmental index
---------------------------------------------------------------------------
# A tibble: 14 x 4
   ENV    Mean   index class
   <fct> <dbl>   <dbl> <chr>
 1 E1     2.52 -0.154  unfavorable
 2 E10    2.18 -0.499  unfavorable
 3 E11    1.37 -1.31   unfavorable
 4 E12    1.61 -1.07   unfavorable
 5 E13    2.91  0.235  favorable
 6 E14    1.78 -0.892  unfavorable
 7 E2     3.18  0.506  favorable
 8 E3     4.06  1.39   favorable
 9 E4     3.68  1.00   favorable
10 E5     3.91  1.24   favorable
11 E6     2.66 -0.0110 unfavorable
12 E7     1.99 -0.685  unfavorable
13 E8     2.54 -0.138  unfavorable
14 E9     3.06  0.382  favorable
---------------------------------------------------------------------------
Analysis for all environments
---------------------------------------------------------------------------
# A tibble: 10 x 6
   GEN       Y Mean_rp Sd_rp    Wi  rank
   <chr> <dbl>   <dbl> <dbl> <dbl> <dbl>
 1 G1     2.60    96.5  7.38  91.5     6
 2 G10    2.47    90.3 18.9   77.5    10
 3 G2     2.74   103.  12.1   94.5     4
 4 G3     2.96   111.   4.59 108.      1
```

```
 5 G4     2.64    99.1  8.03  93.7     5
 6 G5     2.54    95.5  7.74  90.2     8
 7 G6     2.53    95.5  7.61  90.4     7
 8 G7     2.74   105.  12.5   96.0     3
 9 G8     3.00   113.   8.87 107.      2
10 G9     2.51    91.6 13.8   82.3     9
---------------------------------------------------------------------------
Analysis for favorable environments
---------------------------------------------------------------------------
# A tibble: 10 x 6
   GEN        Y Mean_rp Sd_rp    Wi  rank
   <chr> <dbl>   <dbl> <dbl> <dbl> <dbl>
 1 G1     3.43    98.6  5.76  94.8     4
 2 G10    3.31    94.7 18.3   82.4    10
 3 G2     3.62   105.   5.61 101.      3
 4 G3     3.79   110.   6.29 106.      1
 5 G4     3.41    99.0 11.8   91.0     5
 6 G5     3.27    94.6  7.54  89.5     7
 7 G6     3.27    95.2  7.03  90.5     6
 8 G7     3.35    96.8 11.7   88.9     8
 9 G8     3.81   110.  11.7  102.      2
10 G9     3.39    96.6 16.9   85.3     9
---------------------------------------------------------------------------
Analysis for unfavorable environments
---------------------------------------------------------------------------
# A tibble: 10 x 6
   GEN        Y Mean_rp Sd_rp    Wi  rank
   <chr> <dbl>   <dbl> <dbl> <dbl> <dbl>
 1 G1     1.99    94.9  8.40  89.2     8
 2 G10    1.84    86.9 19.8   73.6    10
 3 G2     2.09   101.  15.6   90.7     5
 4 G3     2.33   112.   3.04 110.      2
 5 G4     2.06    99.2  4.41  96.2     4
 6 G5     1.99    96.1  8.35  90.5     6
 7 G6     1.98    95.7  8.49  90.0     7
 8 G7     2.28   110.  10.3  103.      3
 9 G8     2.40   116.   5.36 113.      1
10 G9     1.85    87.8 10.6   80.6     9
```

The function `ecovalence()` computes the Wricke's stability parameter (Wricke, 1965), known as ecovalence ($\omega_i$), estimated by

$$\omega_i = \sum_j \left(Y_{ij} - \bar{Y}_{i.} - \bar{Y}_{.j} + \bar{Y}_{..}\right)$$

where $Y_{ij}$ is the average of genotype $i$ in environment $j$; $\bar{Y}i.$ is the average value of the genotype $i$; $\bar{Y}.j$ is the average value of the environment $j$, and $\bar{Y}..$ is the grand mean. In this method, the most stable genotypes were the ones with the lowest estimates of ($\omega_i$).

```r
eco <- ecovalence(data_ge, ENV, GEN, REP, GY)
print(eco)
```

```
Variable GY
---------------------------------------------------------------------------
Genotypic confidence index
---------------------------------------------------------------------------
# A tibble: 10 x 18
   GEN         E1     E10     E11     E12     E13      E14       E2      E3
   <chr>    <dbl>   <dbl>   <dbl>   <dbl>   <dbl>    <dbl>    <dbl>   <dbl>
 1 G1     -0.0843  0.203   0.0580 -0.196   0.159  -0.178   -0.0687  0.0866
 2 G10    -0.344  -0.436  -0.266  -0.383  -0.876   0.280    0.169   0.253
 3 G2      0.311   0.0519  0.0526  0.315   0.0488 -0.425   -0.0188  0.434
 4 G3      0.0868 -0.120  -0.0816 -0.125   0.278  -0.00338  0.147  -0.213
 5 G4      0.100   0.0245  0.0341 -0.0489 -0.242   0.112    0.0424 -0.178
 6 G5     -0.196   0.102   0.0943  0.217  -0.205   0.139    0.0971 -0.186
 7 G6     -0.0797  0.173   0.273  -0.0826  0.142   0.157    0.251  -0.489
 8 G7      0.186   0.200  -0.0706  0.276   0.208   0.0877  -0.634  -0.0297
 9 G8      0.0493  0.0703 -0.0142  0.0621  0.281  -0.119   -0.0646 -0.284
10 G9     -0.0307 -0.269  -0.0789 -0.0351  0.205  -0.0500   0.0790  0.606
# ... with 9 more variables: E4 <dbl>, E5 <dbl>, E6 <dbl>, E7 <dbl>, E8 <dbl>,
#   E9 <dbl>, Ecoval <dbl>, Ecov_perc <dbl>, rank <dbl>
```

The function `Shukla()` computes the Shukla's stability variance parameter (Shukla, 1972) and uses the Kang's nonparametric stability (Kang & Pham, 1991) to incorporate the mean performance and stability into a single selection criterion.

```r
Shu <- Shukla(data_ge, ENV, GEN, REP, GY)
print(Shu)
```

```
Variable GY
---------------------------------------------------------------------------
Shukla stability variance
---------------------------------------------------------------------------
# A tibble: 10 x 6
   GEN       Y ShuklaVar rMean rShukaVar ssiShukaVar
   <fct> <dbl>     <dbl> <dbl>     <dbl>       <dbl>
 1 G1     2.60    0.0280     6         2           8
 2 G10    2.47    0.244     10        10          20
 3 G2     2.74    0.0861     3         7          10
 4 G3     2.96    0.0121     2         1           3
 5 G4     2.64    0.0640     5         5          10
 6 G5     2.54    0.0480     7         4          11
 7 G6     2.53    0.0468     8         3          11
 8 G7     2.74    0.122      4         8          12
 9 G8     3.00    0.0712     1         6           7
10 G9     2.51    0.167      9         9          18
```

#### 8.2 Regression-based stability analysis

Eberhart & Russell (1966) popularized the regression-based stability analysis. In these procedures, the adaptability and stability analysis is performed by means of adjustments of regression equations where the dependent variable is predicted as a function of an environmental index, according to the following model:

$$Y_{ij} = \beta_{0i} + \beta_{1i}I_j + \delta_{ij} + \bar{\varepsilon}_{ij}$$

where $\beta_{0i}$ is the grand mean of the genotype $i$ ($i = 1, 2, \ldots$, I); $\beta_{1i}$ is the linear response (slope) of the genotype $i$ to the environmental index; $Ij$ is the environmental index ($j = 1, 2, \ldots, e$), where $I_j = [(y_{.j}/g) - (y_{..}/ge)]$, $\delta_{ij}$ is the deviation from the regression, and $\bar{\varepsilon}_{ij}$ is the experimental error. The model is fitted with the function `ge_reg()`.

```
reg_model <- ge_reg(data_ge, ENV, GEN, REP, GY)
print(reg_model)
```

```
Variable GY
---------------------------------------------------------------------------
Joint-regression Analysis of variance
---------------------------------------------------------------------------
# A tibble: 17 x 6
   SV                           Df `Sum Sq` `Mean Sq` `F value`  `Pr(>F)`
   <chr>                     <dbl>    <dbl>     <dbl>     <dbl>     <dbl>
 1 "Total"                     139  324.        2.33      NA     NA
 2 "GEN"                         9   13.0       1.44       6.28   3.05e- 7
 3 "ENV + (GEN x ENV)"         130  311.        2.39      NA     NA
 4 "ENV (linear)"                1  280.      280.        NA     NA
 5 " GEN x ENV (linear)"         9    3.61      0.402      1.75   8.58e- 2
 6 "Pooled deviation"          120   27.6       0.230     NA     NA
 7 "G1"                         12    1.11      0.0924     1.06   3.92e- 1
 8 "G10"                        12    7.54      0.629      7.22   1.66e-11
 9 "G2"                         12    2.95      0.246      2.82   1.14e- 3
10 "G3"                         12    0.699     0.0582     0.669  7.81e- 1
11 "G4"                         12    2.23      0.186      2.14   1.48e- 2
12 "G5"                         12    1.49      0.124      1.42   1.55e- 1
13 "G6"                         12    1.27      0.106      1.22   2.71e- 1
14 "G7"                         12    3.25      0.270      3.11   3.72e- 4
15 "G8"                         12    2.54      0.211      2.43   5.15e- 3
16 "G9"                         12    4.54      0.378      4.34   2.42e- 6
17 "Pooled error"              280   24.4       0.0870    NA     NA
---------------------------------------------------------------------------
Regression parameters
---------------------------------------------------------------------------
# A tibble: 10 x 6
   GEN       Y slope deviations  RMSE    R2
   <chr> <dbl> <dbl>      <dbl> <dbl> <dbl>
 1 G1     2.60 1.06    -0.00142 0.162 0.966
```

```
 2 G10     2.47 1.12      0.177   0.424 0.823
 3 G2      2.74 1.05      0.0497  0.265 0.913
 4 G3      2.96 1.03     -0.0128  0.129 0.977
 5 G4      2.64 0.937     0.0298  0.231 0.917
 6 G5      2.54 0.887     0.00902 0.188 0.937
 7 G6      2.53 0.861     0.00304 0.174 0.942
 8 G7      2.74 0.819     0.0579  0.278 0.852
 9 G8      3.00 1.03      0.0382  0.246 0.922
10 G9      2.51 1.19      0.0938  0.329 0.897
---------------------------------------------------------------------------
```

```
plot(reg_model)
```

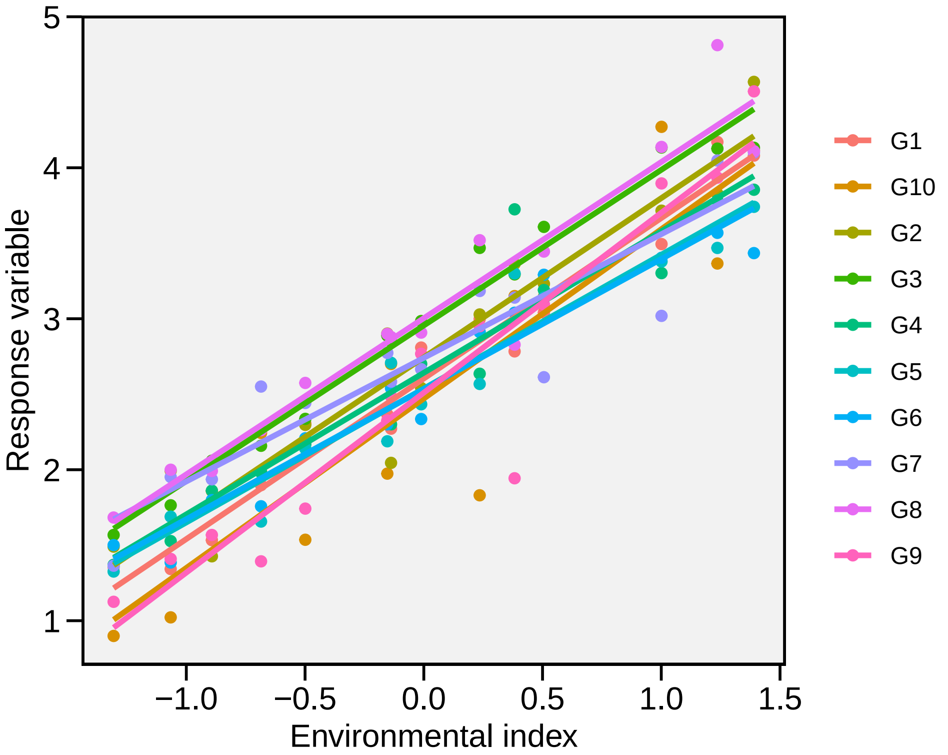

#### 8.3 Non-parametric models

The function superiority() implements the nonparametric method proposed by Lin & Binns (1988), which considers that a measure of cultivar general superiority for cultivar x location data is defined as the distance mean square between the cultivar's response and the maximum response averaged over all locations, according to the following model.

$$P_i = \sum_{j=1}^{n} (y_{ij} - y_{.j})^2/(2n)$$

where $n$ is the number of environments

```r
super <- superiority(data_ge, ENV, GEN, REP, GY)
print(super)
```

```
Variable GY
---------------------------------------------------------------------------
Superiority index considering all, favorable and unfavorable environments
---------------------------------------------------------------------------
# A tibble: 10 x 8
   GEN       Y   Pi_a   R_a   Pi_f   R_f    Pi_u   R_u
   <chr> <dbl>  <dbl> <dbl>  <dbl> <dbl>   <dbl> <dbl>
 1 G1     2.60 0.169      5 0.228      4 0.125       6
 2 G10    2.47 0.344     10 0.475     10 0.245      10
 3 G2     2.74 0.126      3 0.149      3 0.108       5
 4 G3     2.96 0.0410     1 0.0723     1 0.0175      2
 5 G4     2.64 0.173      6 0.289      5 0.0853      4
 6 G5     2.54 0.240      8 0.382      8 0.133       7
 7 G6     2.53 0.238      7 0.377      7 0.134       8
 8 G7     2.74 0.149      4 0.318      6 0.0214      3
 9 G8     3.00 0.0412     2 0.0882     2 0.00588     1
10 G9     2.51 0.291      9 0.390      9 0.217       9
---------------------------------------------------------------------------
```

The function Fox() performs a stability analysis based on the criteria of Fox, Skovmand, Thompson, Braun, & Cormier (1990), using the statistical “TOP third” only. A stratified ranking of the genotypes at each environment is done. The proportion of locations at which the genotype occurred in the top third is expressed in the output.

```r
fox <- Fox(data_ge, ENV, GEN, REP, GY)
print(fox)
```

```
Variable GY
---------------------------------------------------------------------------
Fox TOP third criteria
---------------------------------------------------------------------------
# A tibble: 10 x 3
   GEN       Y   TOP
   <fct> <dbl> <int>
 1 G1     2.60     2
 2 G10    2.47     3
 3 G2     2.74     5
 4 G3     2.96     9
 5 G4     2.64     3
 6 G5     2.54     1
 7 G6     2.53     1
 8 G7     2.74     5
 9 G8     3.00    12
10 G9     2.51     1
```

#### 8.4 Stability analysis based on factor analysis

A method that combines stability analysis and environmental stratification using factor analysis was proposed by Murakami & Cruz (2004). This model is implemented with the function `ge_factanal()`

```
fact <- ge_factanal(data_ge, ENV, GEN, REP, GY)
plot(fact)
```

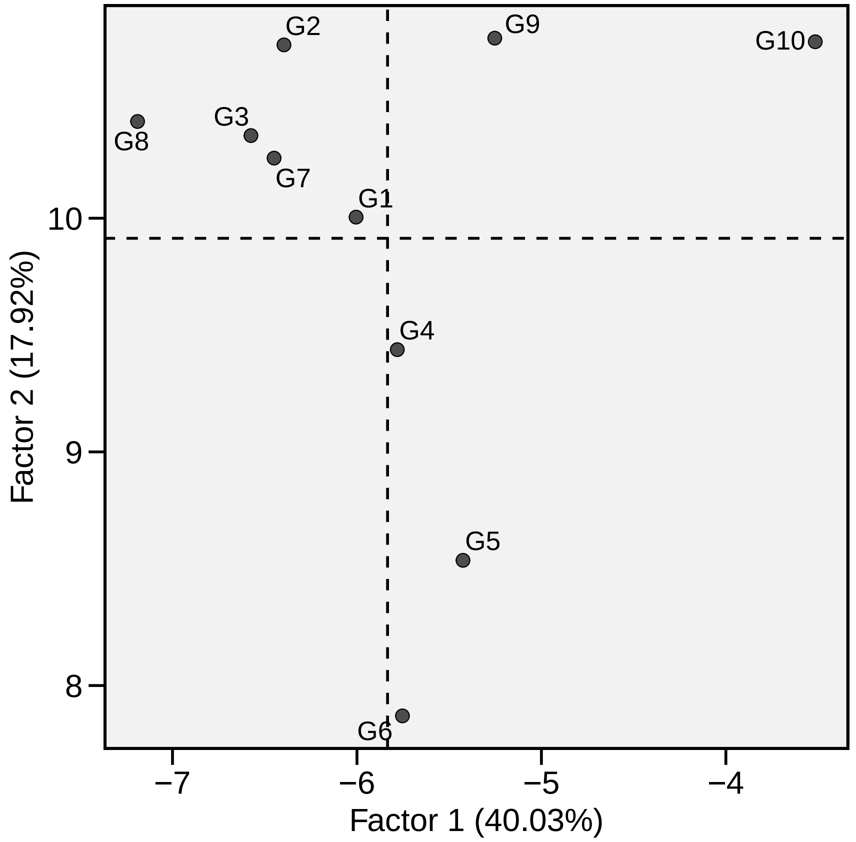

```
print(fact)
```

```
Variable GY
---------------------------------------------------------------------------------
Correlation matrix among environments
---------------------------------------------------------------------------------
# A tibble: 14 x 15
   ENV         E1       E10      E11      E12      E13      E14      E2       E3
   <chr>    <dbl>     <dbl>    <dbl>    <dbl>    <dbl>    <dbl>    <dbl>    <dbl>
 1 E1      1         0.7826   0.7818   0.8685   0.8249   0.1973  0.2273   0.3565
 2 E10     0.7826   1         0.9169   0.7974   0.8124   0.2322  0.1013  -0.1357
 3 E11     0.7818   0.9169   1         0.7491   0.8400   0.2342  0.4384  -0.1651
 4 E12     0.8685   0.7974   0.7491   1         0.7401   0.1552  0.05270  0.2494
```

```
 5 E13    0.8249   0.8124   0.8400   0.7401   1        0.1606  0.2464   0.1828
 6 E14    0.1973   0.2322   0.2342   0.1552   0.1606   1       0.2468  -0.4269
 7 E2     0.2273   0.1013   0.4384   0.05270  0.2464   0.2468  1       -0.06944
 8 E3     0.3565  -0.1357  -0.1651   0.2494   0.1828  -0.4269 -0.06944  1
 9 E4    -0.01148 -0.3422  -0.1297  -0.1902  -0.04504  0.1489  0.6611   0.4093
10 E5     0.7043   0.6755   0.6347   0.5672   0.8140   0.2178  0.2101   0.3104
11 E6     0.5799   0.3685   0.3489   0.3152   0.6375   0.2738  0.2893   0.4457
12 E7     0.4538   0.3943   0.1883   0.3496   0.1518   0.5661 -0.1665   0.04646
13 E8    -0.05268  0.02451  0.07734  0.03424  0.1216   0.8205  0.3168  -0.2927
14 E9     0.2193   0.2813   0.2406   0.2165  -0.1840   0.3229  0.1193  -0.3746
# ... with 6 more variables: E4 <dbl>, E5 <dbl>, E6 <dbl>, E7 <dbl>, E8 <dbl>,
#   E9 <dbl>
-------------------------------------------------------------------------------
Eigenvalues and explained variance
-------------------------------------------------------------------------------
# A tibble: 14 x 4
   PCA   Eigenvalues   Variance Cumul_var
   <fct>       <dbl>      <dbl>     <dbl>
 1 PC1     5.604e+ 0  4.003e+ 1     40.03
 2 PC2     2.509e+ 0  1.792e+ 1     57.95
 3 PC3     2.406e+ 0  1.719e+ 1     75.14
 4 PC4     1.371e+ 0  9.795e+ 0     84.94
 5 PC5     1.127e+ 0  8.049e+ 0     92.98
 6 PC6     4.873e- 1  3.481e+ 0     96.47
 7 PC7     3.029e- 1  2.164e+ 0     98.63
 8 PC8     1.295e- 1  9.249e- 1     99.55
 9 PC9     6.243e- 2  4.459e- 1    100
10 PC10    1.720e-16  1.229e-15    100
11 PC11    1.413e-16  1.009e-15    100
12 PC12   -1.744e-16 -1.246e-15    100
13 PC13   -2.667e-16 -1.905e-15    100
14 PC14   -3.008e-16 -2.149e-15    100
-------------------------------------------------------------------------------
Initial loadings
-------------------------------------------------------------------------------
# A tibble: 14 x 6
   Env        PC1      PC2       PC3      PC4      PC5
   <fct>    <dbl>    <dbl>     <dbl>    <dbl>    <dbl>
 1 E1    -0.9230  -0.06635  0.1988    0.1005   0.2390
 2 E10   -0.8778  -0.3987   0.08996  -0.1172  -0.1036
 3 E11   -0.8706  -0.2409   0.008640 -0.4145   0.03163
 4 E12   -0.8295  -0.2430   0.2138    0.06387  0.08795
 5 E13   -0.9082   0.08532  0.2121   -0.1810  -0.2260
 6 E14   -0.3737  -0.07328 -0.8644    0.1969  -0.1510
 7 E2    -0.3126   0.4096  -0.3996   -0.6131   0.4430
 8 E3    -0.1636   0.5788   0.5718    0.4218   0.3106
 9 E4    -0.04258  0.8411  -0.3329   -0.01830  0.3691
10 E5    -0.8513   0.3284   0.1117    0.08357 -0.2274
```

```
11 E6    -0.6748   0.5340   0.01988   0.1843  -0.07906
12 E7    -0.4507  -0.2416  -0.3579    0.7060   0.1730
13 E8    -0.2188   0.2887  -0.8149    0.05187 -0.3359
14 E9    -0.1171  -0.6865  -0.3437    0.06174  0.5888
---------------------------------------------------------------------------------
Loadings after varimax rotation and commonalities
---------------------------------------------------------------------------------
# A tibble: 14 x 8
   Env         FA1      FA2       FA3      FA4      FA5 Communality Uniquenesses
   <fct>     <dbl>    <dbl>     <dbl>    <dbl>    <dbl>       <dbl>        <dbl>
 1 E1    -0.8813    0.3273   0.009273 -0.06306  0.2743       0.9631     0.03694
 2 E10   -0.9419   -0.1585  -0.08198   0.1131   0.1736       0.9620     0.03799
 3 E11   -0.9288   -0.2335  -0.03364  -0.2420   0.1100       0.9889     0.01106
 4 E12   -0.8478    0.1346   0.02632   0.09415  0.2412       0.8047     0.1953
 5 E13   -0.9397    0.1085  -0.08419  -0.06373 -0.2345       0.9609     0.03907
 6 E14   -0.1501   -0.1229  -0.9157   -0.08719  0.2646       0.9537     0.04629
 7 E2    -0.1979   -0.05213 -0.1260   -0.9688   0.03278      0.9973     0.002659
 8 E3    -0.08057   0.9100   0.3407   -0.01732 -0.1102       0.9630     0.03697
 9 E4     0.2093    0.5430  -0.2721   -0.7277  -0.1202       0.9567     0.04331
10 E5    -0.7771    0.3920  -0.2692   -0.04704 -0.2675       0.9037     0.09629
11 E6    -0.5240    0.5692  -0.3092   -0.1740  -0.2380       0.7811     0.2189
12 E7    -0.2438    0.3424  -0.5197    0.2967   0.6190       0.9180     0.08197
13 E8     0.001609 -0.05886 -0.9143   -0.2255  -0.1434       0.9109     0.08911
14 E9    -0.07939  -0.2910  -0.01825  -0.05390  0.9271       0.9537     0.04633
---------------------------------------------------------------------------------
Environmental stratification based on factor analysis
---------------------------------------------------------------------------------
# A tibble: 14 x 6
   Env   Factor  Mean    Min   Max     CV
   <fct> <fct>  <dbl>  <dbl> <dbl>  <dbl>
 1 E1    FA1    2.521 1.974  2.902 13.30
 2 E10   FA1    2.175 1.536  2.575 14.44
 3 E11   FA1    1.368 0.8991 1.683 16.39
 4 E12   FA1    1.609 1.022  2     20.31
 5 E13   FA1    2.910 1.831  3.520 16.76
 6 E5    FA1    3.910 3.366  4.812 10.71
 7 E3    FA2    4.064 3.435  4.569  8.223
 8 E6    FA2    2.663 2.336  2.985  7.955
 9 E14   FA3    1.782 1.427  2.060 11.70
10 E8    FA3    2.536 2.045  2.879 10.55
11 E2    FA4    3.180 2.613  3.608  8.253
12 E4    FA4    3.675 3.019  4.271 11.45
13 E7    FA5    1.989 1.393  2.551 16.85
14 E9    FA5    3.057 1.943  3.725 15.57
---------------------------------------------------------------------------------
Mean = mean; Min = minimum; Max = maximum; CV = coefficient of variation (%)
The print statistics are based on the men values of  3 replicates
---------------------------------------------------------------------------------
```

#### 8.5 AMMI analysis

##### 8.5.1 The model

The response variable of the genotype $i$ in the environment $j$ using The Additive Main Effect and Multiplicative interaction (AMMI) model, is estimated by

$$y_{ij} = \mu + \alpha_i + \tau_j + \sum_{k=1}^{p} \lambda_k a_{ik} t_{jk} + \rho_{ij} + \varepsilon_{ij}$$

where $\lambda_k$ is the singular value for the $k$-th interaction principal component axis (IPCA); $a_{ik}$ is the $i$-th element of the $k$-th eigenvector; $t_{jk}$ is the $j$th element of the $k$th eigenvector. A residual $\rho_{ij}$ remains, if not all $p$ IPCA are used, where $p \leq min(g-1; e-1)$. The AMMI model is fitted with the function `performs_ammi()` or `waas()`. The last computes the Weighted Average of Absolute Scores (T. Olivoto, Lúcio, et al., 2019) and can also be used to fit an AMMI model.

```
AMMI_model <- performs_ammi(data_ge, ENV, GEN, REP, GY)
```

```
variable GY
---------------------------------------------------------------------------
AMMI analysis table
---------------------------------------------------------------------------
    Source  Df  Sum Sq Mean Sq F value   Pr(>F) Percent Accumul
       ENV  13 279.574 21.5057   62.33 0.00e+00       .       .
  REP(ENV)  28   9.662  0.3451    3.57 3.59e-08       .       .
  GEN        9  12.995  1.4439   14.93 2.19e-19       .       .
   ENV:GEN 117  31.220  0.2668    2.76 1.01e-11       .       .
       PC1  21  10.749  0.5119    5.29 0.00e+00    34.4    34.4
       PC2  19   9.924  0.5223    5.40 0.00e+00    31.8    66.2
       PC3  17   4.039  0.2376    2.46 1.40e-03    12.9    79.2
       PC4  15   3.074  0.2049    2.12 9.60e-03     9.8      89
       PC5  13   1.446  0.1113    1.15 3.18e-01     4.6    93.6
       PC6  11   0.932  0.0848    0.88 5.61e-01       3    96.6
       PC7   9   0.567  0.0630    0.65 7.53e-01     1.8    98.4
       PC8   7   0.362  0.0518    0.54 8.04e-01     1.2    99.6
       PC9   5   0.126  0.0252    0.26 9.34e-01     0.4     100
 Residuals 252  24.367  0.0967      NA       NA       .       .
     Total 419 357.816  0.8540      NA       NA       .       .
---------------------------------------------------------------------------

All variables with significant (p < 0.05) genotype-vs-environment interaction
Done!
```

```
AMMI_model2 <- waas(data_ge, ENV, GEN, REP, GY, verbose = FALSE)
print(AMMI_model, digits = 2)
```

```
Variable GY
```

```
---------------------------------------------------------------------------
AMMI analysis table
---------------------------------------------------------------------------
# A tibble: 15 x 8
   Source          Df `Sum Sq` `Mean Sq` `F value` `Pr(>F)` Percent Accumul
   <chr>        <dbl>    <dbl>     <dbl>     <dbl>    <dbl> <chr>   <chr>
 1 "ENV"           13   280.      22.         62.     0.       .       .
 2 "REP(ENV)"      28     9.7      0.35        3.6    3.6e- 8 .       .
 3 "GEN      "      9    13.       1.4        15.     2.2e-19 .       .
 4 "ENV:GEN"      117    31.       0.27        2.8    1.0e-11 .       .
 5 "PC1"           21    11.       0.51        5.3    0.      34.4    34.4
 6 "PC2"           19     9.9      0.52        5.4    0.      31.8    66.2
 7 "PC3"           17     4.0      0.24        2.5    1.4e- 3 12.9    79.2
 8 "PC4"           15     3.1      0.20        2.1    9.6e- 3 9.8     89
 9 "PC5"           13     1.4      0.11        1.2    3.2e- 1 4.6     93.6
10 "PC6"           11     0.93     0.085       0.88   5.6e- 1 3       96.6
11 "PC7"            9     0.57     0.063       0.65   7.5e- 1 1.8     98.4
12 "PC8"            7     0.36     0.052       0.54   8.0e- 1 1.2     99.6
13 "PC9"            5     0.13     0.025       0.26   9.3e- 1 0.4     100
14 "Residuals"   252    24.       0.097      NA     NA       .       .
15 "Total"       419   358.       0.85       NA     NA       .       .
---------------------------------------------------------------------------
Scores for genotypes and environments
---------------------------------------------------------------------------
# A tibble: 24 x 12
   type  Code      Y    PC1    PC2    PC3    PC4    PC5    PC6    PC7    PC8
   <chr> <fct> <dbl>  <dbl>  <dbl>  <dbl>  <dbl>  <dbl>  <dbl>  <dbl>  <dbl>
 1 GEN   G1      2.6  0.32  -0.044 -0.036 -0.066 -0.31   0.43  -0.15   0.25
 2 GEN   G10     2.5 -1.0   -0.57  -0.17  -0.33  -0.12  -0.11  -0.035  0.069
 3 GEN   G2      2.7  0.14   0.20  -0.73   0.47  -0.048 -0.28  -0.077  0.087
 4 GEN   G3      3.0  0.043 -0.10   0.23   0.18  -0.13  -0.14   0.46  -0.097
 5 GEN   G4      2.6 -0.33   0.48  -0.091  0.14  -0.19   0.35   0.089 -0.29
 6 GEN   G5      2.5 -0.33   0.25   0.25   0.18   0.47   0.033 -0.29  -0.12
 7 GEN   G6      2.5 -0.098  0.24   0.56   0.24   0.051 -0.10   0.059  0.33
 8 GEN   G7      2.7  0.28   0.59  -0.21  -0.71   0.23  -0.084  0.13   0.047
 9 GEN   G8      3.0  0.50  -0.19   0.32  -0.17  -0.33  -0.29  -0.27  -0.22
10 GEN   G9      2.5  0.47  -0.84  -0.12   0.064  0.38   0.19   0.092 -0.052
# ... with 14 more rows, and 1 more variable: PC9 <dbl>
```

To inspect the fitted model above we can use the S3 generic function `plot()`.

```
plot(AMMI_model,
     which = c(1, 2, 5, 7),
     ncol = 4,
     labels = TRUE,
     size.lab.out = 4)
```

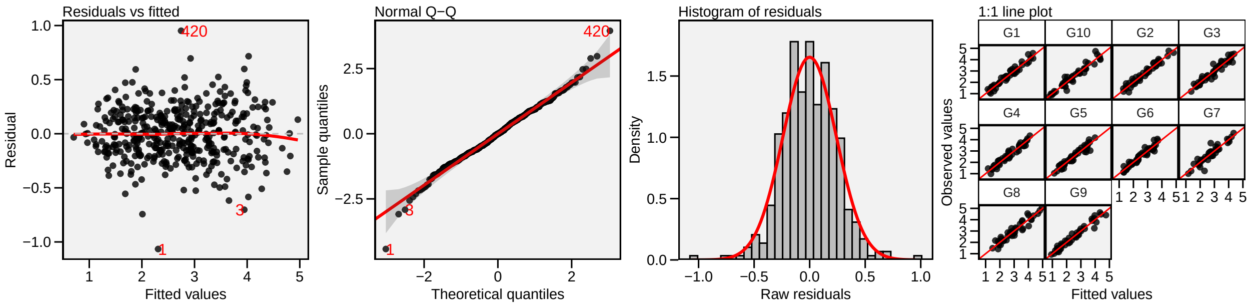

**Quick tip**

1:1 line plots are useful for identifying the predictive ability of models. Because it is a scatter plot with a 1:1 reference line, the x and y axes must have the same scale. It is assumed that the diagonal line has an intercept equal to zero and *slope* equal to one. A hypothesis testing for these parameters may be useful.

##### 8.5.2 Prediction

The response variable of a two-way table (for example, the yield of $g$ genotypes in $e$ environments) can be estimated using the S3 method `predict()`. This estimation is based on the number of multiplicative terms declared in the function. A summary of all possible AMMI models is presented below.

| Member of AMMI family | Espected response of the $i$-th genotype in the $j$th environment |
|---|---|
| AMMI0 | $\hat{y}_{ij} = \bar{y}_{i.} + \bar{y}_{.j} - \bar{y}_{..}$ |
| AMMI1 | $\hat{y}_{ij} = \bar{y}_{i.} + \bar{y}_{.j} - \bar{y}_{..} + \lambda_1 a_{i1} t_{j1}$ |
| AMMI2 | $\hat{y}_{ij} = \bar{y}_{i.} + \bar{y}_{.j} - \bar{y}_{..} + \lambda_1 a_{i1} t_{j1} + \lambda_2 a_{i2} t_{j2}$ |
| ... | |
| AMMIF | $\hat{y}_{ij} = \bar{y}_{i.} + \bar{y}_{.j} - \bar{y}_{..} + \lambda_1 a_{i1} t_{j1} + \lambda_2 a_{i2} t_{j2} + ... + \lambda_p a_{ip} t_{jp}$ |

Procedures based on postdictive success, such as Gollobs's test (Gollob, 1968) or predictive success, such as cross-validation (T. Olivoto, Lúcio, et al., 2019) should be used to define the number of IPCA used for estimating the response variable in AMMI analysis. In our example, four IPCAs were significant. So we will predict the GY using this number of IPCAs and use the function `make_mat()` to create a two-way table with the predicted values that -for better printing- will show the environments in rows and genotypes in columns.

```
predicted <- predict(AMMI_model, naxis = 4)
pred_mat <- make_mat(predicted$GY, ENV, GEN, YpredAMMI)
print(pred_mat, digits = 4)
```

```
      G1   G10    G2    G3    G4    G5    G6    G7    G8    G9
E1 2.521 1.964 2.881 2.757 2.538 2.313 2.298 2.752 2.845 2.340
```

```
E10 2.151 1.516 2.277 2.459 2.242 2.091 2.174 2.479 2.574 1.789
E11 1.296 0.888 1.478 1.711 1.407 1.323 1.400 1.364 1.718 1.099
E12 1.596 1.042 1.928 1.842 1.662 1.436 1.428 1.884 1.924 1.343
E13 3.047 1.828 3.017 3.279 2.704 2.643 2.877 3.124 3.620 2.957
E14 1.625 1.903 1.485 2.072 1.785 1.767 1.773 1.883 2.098 1.429
E2  3.005 3.134 3.232 3.621 3.158 3.210 3.262 2.631 3.434 3.113
E3  4.060 4.156 4.617 4.221 3.880 3.620 3.396 4.051 4.265 4.377
E4  3.527 4.210 3.617 4.055 3.382 3.472 3.407 3.054 3.971 4.058
E5  4.020 3.336 3.843 4.206 3.607 3.555 3.683 4.196 4.590 4.069
E6  2.626 2.544 2.692 2.944 2.512 2.437 2.431 2.698 3.060 2.687
E7  1.873 2.179 1.904 2.087 2.063 1.828 1.669 2.567 2.193 1.525
E8  2.394 2.746 2.052 2.884 2.399 2.492 2.557 2.502 2.972 2.366
E9  2.710 3.151 3.394 3.237 3.648 3.335 3.121 3.188 2.788 1.993
```

##### 8.5.3 Biplots

Provided that an object of class `performs_ammi` is available in the global environment, the graphics may be obtained using the function `plot_scores()`.

- **biplot type 1: GY x PC1**

```
a <- plot_scores(AMMI_model)
b <- plot_scores(AMMI_model,
                 col.gen = "black",
                 col.env = "gray70",
                 col.segm.env = "gray70",
                 plot_theme = theme_metan_minimal())
arrange_ggplot(a, b, labels = letters[1:2])
```

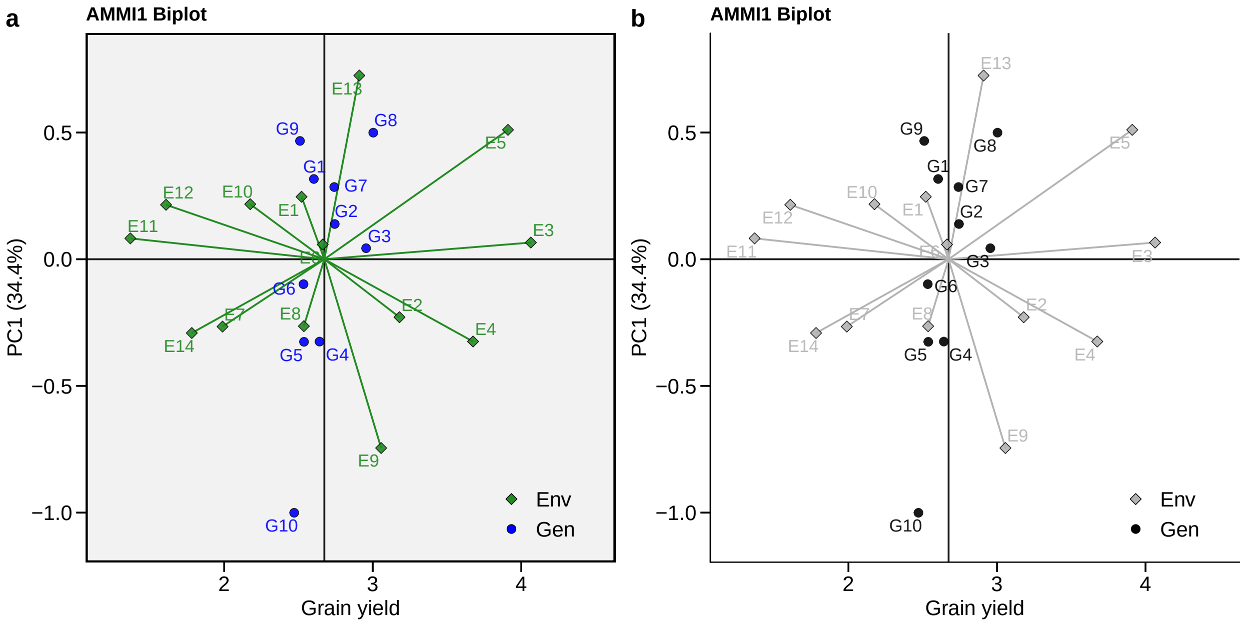

- **biplot type 2: PC1 x PC2**

```
c <- plot_scores(AMMI_model, type = 2)
d <- plot_scores(AMMI_model,
                 type = 2,
                 polygon = T,
                 col.segm.env = "#FFFFFF00", # Transparent
                 axis.expand = 1.5,
                 plot_theme = theme_metan(grid = "both"))

arrange_ggplot(c, d, labels = letters[3:4])
```

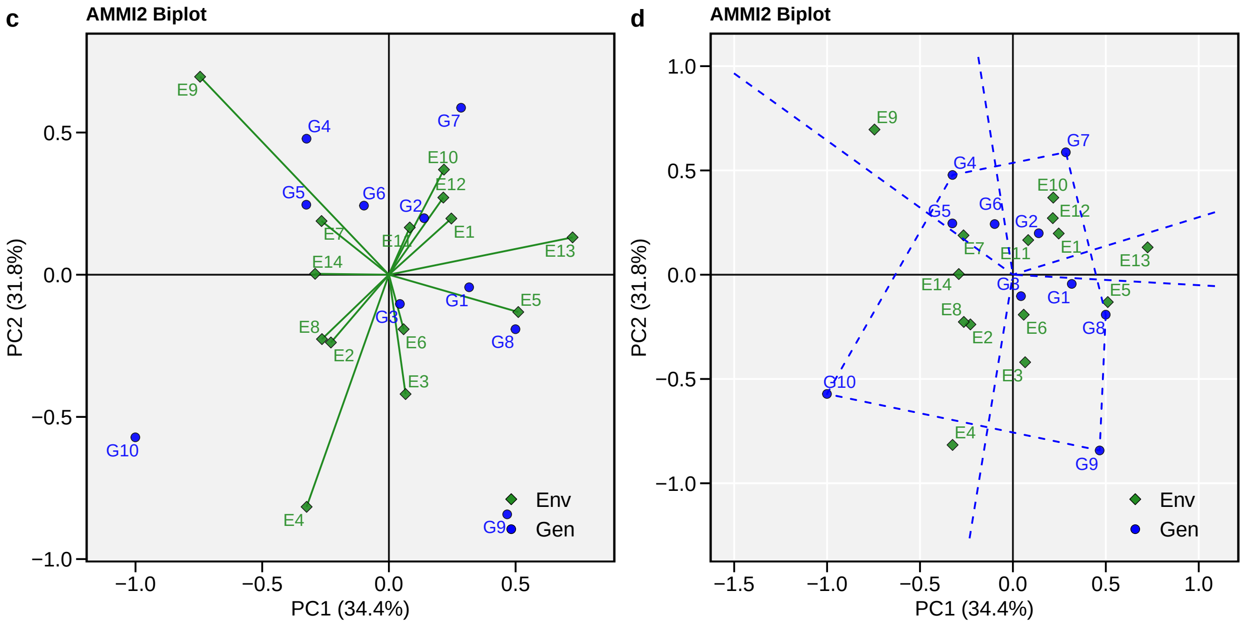

- **biplot type 3: GY x WAAS**

The quadrants proposed by T. Olivoto, Lúcio, et al. (2019) in the following biplot represent four classifications regarding the joint interpretation of mean performance and stability. The genotypes or environments included in quadrant I can be considered unstable genotypes or environments with high discrimination ability, and with productivity below the grand mean. In the quadrant II are included unstable genotypes, although with productivity above the grand mean. The environments included in this quadrant deserve special attention since, in addition to providing high magnitudes of the response variable, they present a good discrimination ability. Genotypes within quadrant III have low productivity, but can be considered stable due to the lower values of WAASB. The lower this value, the more stable the genotype can be considered. The environments included in this quadrant can be considered as poorly productive and with low discrimination ability. The genotypes within the quadrant IV are highly productive and broadly adapted due to the high magnitude of the response variable and high stability performance (lower values of WAASB). Only objects of class `waas` can be used to produce such biplot (in our example, `waas_index2`).

```
e <- plot_scores(AMMI_model2, type = 3)
f <- plot_scores(AMMI_model2,
                 type = 3,
                 x.lab = "My custom x label",
                 size.shape = 4, # Size of the shape point
                 col.gen = "gray50", # Color for genotypes
                 size.tex.pa = 4, # Size of the text
                 col.alpha.env = 0, # Transparency of environment's point
                 x.lim = c(2.4, 3.1), # Limits of x axis
                 x.breaks = seq(2.4, 3.1, by = 0.1), # Markers of x axis
                 y.lim = c(0, 0.7))+
    theme(legend.position = "none") +
    theme_metan_minimal() +
    ggtitle("WAASB vs GY plot", subtitle = "Zoom in genotypes' scores")
arrange_ggplot(e, f, labels = letters[5:6])
```

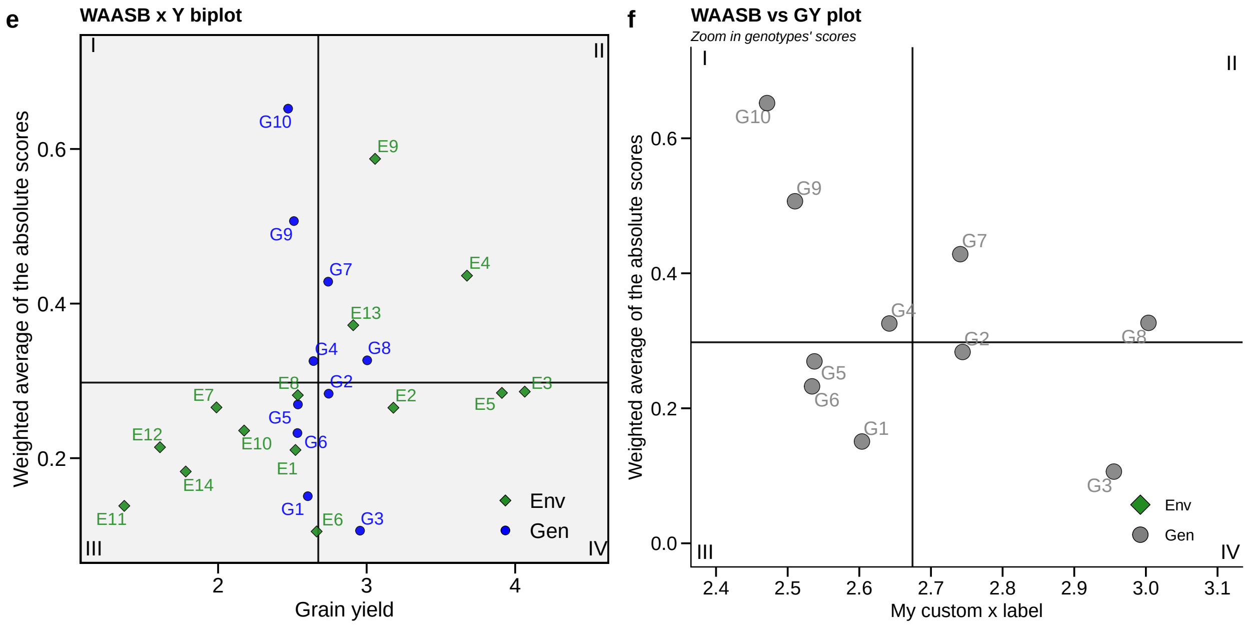

- **nominal yield and environment IPCA1**

A graphic with the nominal yield ($\hat{y}_{ij}^*$ ) as a function of the environment IPCA1 scores can be used for identifying possible mega-environments as well as visualizing the "which-won-where" pattern was produced. In this graphic, each genotype is depicted by a straight line with the equation $\hat{y}_{ij}^* = \mu_i + \lambda_1^{0.5} a_{i1} \times \lambda_1^{0.5} t_{j1}$ , where $\mu_i$ is the nominal yield for the ith genotype in $i$ the environment $j$; $\mu_i$ is the grand mean of the genotype $i$; $\lambda_1^{0.5} a_{i1}$ is the IPCA1 score of the genotype $i$; and $\lambda_1^{0.5} t_{j1}$ is the IPCA1 score of the environment $j$. The winner genotype in a given environment has the highest nominal yield in that environment (Gauch & Zobel, 1997).

```
g <- plot_scores(AMMI_model, type = 4)
h <- plot_scores(AMMI_model,
                 type = 4,
                 color = FALSE,
                 col.alpha.gen = 0,
                 col.alpha.env = 0,
                 plot_theme = theme_metan(grid = "y"))
arrange_ggplot(g, h, labels = letters[7:8])
```

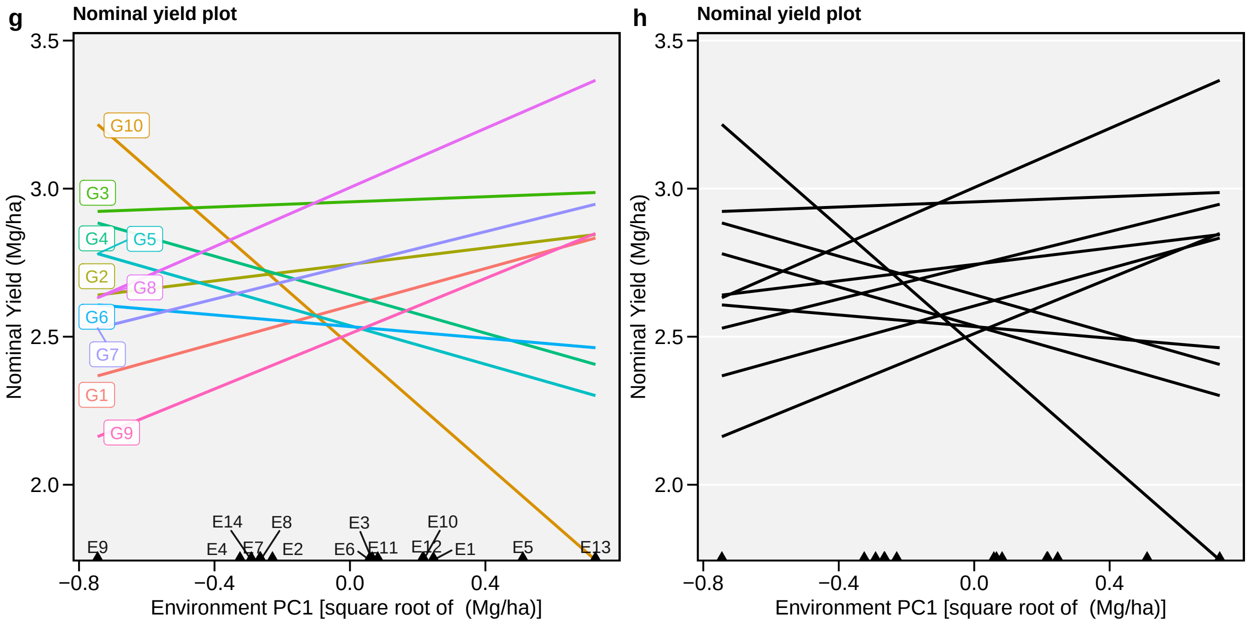

##### 8.5.4 AMMI-based stability statistics

The function `AMMI_indexes()` can be used to compute the following AMMI-based stability statistics

- **AMMI stability value, ASV, (Purchase, Hatting, & Deventer, 2000).**

$$ASV_i = \sqrt{\left[\frac{\lambda_1^2}{\lambda_2^2} \times (\lambda_1^{0.5} a_{i1})\right]^2 + (\lambda_2^{0.5} a_{i2})^2}$$

- **Sums of the absolute value of the IPCA scores**

$$SIPC_i = \sum_{k=1}^{P} \left|\lambda_k^{0.5} a_{ik}\right|$$

- **Averages of the squared eigenvector values**

$$EV_i = \sum_{k=1}^{P} a_{ik}^2/P$$

described by Sneller, Kilgore-Norquest, & Dombek (1997), where $P$ is the number of IPCA retained via F-tests.

- **Absolute value of the relative contribution of IPCAs to the interaction, Za, (Zali, Farshadfar, Sabaghpour, & Karimizadeh, 2012).**

$$Za_i = \sum_{k=1}^{P} \theta_k a_{ik}$$

where $\theta_k$ is the percentage sum of squares explained by the $k$-th IPCA.

- **Weighted Average of Absolute Scores (T. Olivoto, Lúcio, et al., 2019)**

$$WAAS_i = \sum_{k=1}^{P} |\lambda_k^{0.5} a_{ik} \times \theta_k| / \sum_{k=1}^{P} \theta_k$$

Simultaneous selection indexes (ssi), are computed by summation of the ranks of the ASV, SIPC, EV, Za, and WAAS indexes and the ranks of the mean yields (Farshadfar, 2008), which results in ssiASV, ssiSIPC, ssiEV, ssiZa, and ssiWAAS, respectively.

The `AMMI_indexes()` function has two arguments. The first (`x`) is the model, which must be an object of the class `waas` or `performs_ammi`. The second, (`order.y`) is the order for ranking the response variable. By default, it is set to NULL, which means that the response variable is ordered in descending order. If `x` is a list with more than one variable, `order.y` must be a vector of the same length of `x`. Each element of the vector must be one of the `order.y = "h"` or `order.y = "l"`. If `order.y = "h"` is used, the response variable will be ordered from maximum to minimum. If `order.y = "l"` is used then the response variable will be ordered from minimum to maximum.

To show how functions of `metan` interact easily with `%>%`, in the next example, we will fit an AMMI model for all numeric variables in `data_ge2`, except the ones that contains `"E"` (with `-contains("E")`), compute the stability statistics and extract the data for the Weigthed Average of Absolute Scores (WAAS) and their respective ranks.

```
ammi_ind <- data_ge2 %>%
  performs_ammi(ENV, GEN, REP,
                resp = -contains("E"),
                verbose = FALSE) %>%
  AMMI_indexes()
get_model_data(ammi_ind, "WAAS")
```

```
# A tibble: 13 x 9
   gen      PH    CL    CD    CW    KW    NR   NKR   TKW
   <fct> <dbl> <dbl> <dbl> <dbl> <dbl> <dbl> <dbl> <dbl>
 1 H1    0.318 0.799 0.130 1.34  0.782 0.427 0.929 2.72
```

```
 2 H10   0.230 0.948 0.306 1.04  0.895 0.372 0.506 2.15
 3 H11   0.201 0.407 0.214 0.692 0.321 0.794 0.836 1.26
 4 H12   0.364 0.530 0.351 0.257 2.15  0.309 0.228 0.558
 5 H13   0.363 0.866 0.520 1.39  1.95  0.498 0.946 0.514
 6 H2    0.342 1.16  0.550 2.11  3.37  0.796 0.404 4.41
 7 H3    0.374 0.603 0.552 0.507 2.81  0.611 0.252 4.10
 8 H4    0.294 0.511 0.415 0.537 1.69  0.436 0.281 3.07
 9 H5    0.168 0.411 0.237 0.766 0.519 0.837 0.611 0.738
10 H6    0.270 0.733 0.569 0.935 3.22  0.369 1.60  1.64
11 H7    0.228 0.453 0.595 1.57  3.03  0.284 0.518 3.44
12 H8    0.315 1.03  0.770 1.81  3.89  0.352 0.941 4.91
13 H9    0.146 0.699 0.608 1.61  3.11  0.502 0.888 5.50
```

```r
get_model_data(ammi_ind, "WAAS_R")
```

```
# A tibble: 13 x 9
   gen      PH    CL    CD    CW    KW    NR   NKR   TKW
   <fct> <dbl> <dbl> <dbl> <dbl> <dbl> <dbl> <dbl> <dbl>
 1 H1        9     9     1     8     3     6    10     7
 2 H10       5    11     4     7     4     5     5     6
 3 H11       3     1     2     4     1    11     8     4
 4 H12      12     5     5     1     7     2     1     2
 5 H13      11    10     7     9     6     8    12     1
 6 H2       10    13     8    13    12    12     4    11
 7 H3       13     6     9     2     8    10     2    10
 8 H4        7     4     6     3     5     7     3     8
 9 H5        2     2     3     5     2    13     7     3
10 H6        6     8    10     6    11     4    13     5
11 H7        4     3    11    10     9     1     6     9
12 H8        8    12    13    12    13     3    11    12
13 H9        1     7    12    11    10     9     9    13
```

#### 8.6 BLUP prediction in MET analysis

##### 8.6.1 The model

Assuming $\alpha_i$ and $(\alpha\tau)_{ij}$ to be random effects, the model in Eq.(8.1) can be conviently rewriten in a standard linear mixed-effect model

$$\mathbf{y} = \mathbf{Xb} + \mathbf{Zu} + \mathbf{e}$$

where $\mathbf{y}$ is an $n[=\sum_{j=1}^{e}(gb)] \times 1$ vector of response variable $\mathbf{y} = [y_{111}, y_{112}, \dots, y_{geb}]'$; $\mathbf{b}$ is an $(eb) \times 1$ vector of unknown fixed effects $\mathbf{b} = [\gamma_{11}, \gamma_{12}, ..., \gamma_{eb}]'$; $\mathbf{u}$ is an $m[= g + ge] \times 1$ vector of random effects $\mathbf{u} = \left[\alpha_1, \alpha_2, ..., \alpha_g, (\alpha\tau)_{11}, (\alpha\tau)_{12}, ..., (\alpha\tau)_{ge}\right]'$; $\mathbf{X}$ is an $n \times (eb)$ design matrix relating $\mathbf{y}$ to $\mathbf{b}$; $\mathbf{Z}$ is an $n \times m$ design matrix relating $\mathbf{y}$ to $\mathbf{u}$ ; $\mathbf{e}$ is an $n \times 1$ vector of random errors $\mathbf{e} = [y_{111}, y_{112}, \dots, y_{geb}]'$;

The vectors **b** and **u** are estimated using the well-known mixed model equation

$$\begin{bmatrix} \hat{\mathbf{b}} \\ \hat{\mathbf{u}} \end{bmatrix} = \begin{bmatrix} \mathbf{X'R^{-1}X} & \mathbf{X'R^{-1}Z} \\ \mathbf{Z'R^{-1}X} & \mathbf{Z'R^{-1}Z+G^{-1}} \end{bmatrix}^{-} \begin{bmatrix} \mathbf{X'R^{-1}y} \\ \mathbf{Z'R^{-1}y} \end{bmatrix}$$

where **G** and **R** are the variance-covariance matrices for random-effect vector **u** and residual vector **e**, respectively.

The function `waasb()` is used to fit the linear mixed-effect model. By default, genotype and genotype-vs-environment interaction are assumed to be random effects. Other effects may be considered using the argument `random`. Here, we will analyze a vector of response variables (PH, ED, TKW, and NKR) from the data example `data_ge2`.

```
WAASB_model <- waasb(data_ge2, ENV, GEN, REP,
                     resp = c(PH, ED, TKW, NKR))
```

```
Evaluating variable PH 0 %
Evaluating variable ED 33.3 %
Evaluating variable TKW 66.7 %
Evaluating variable NKR 100 %
All variables with significant (p < 0.05) genotype-vs-environment interaction
Done!
```

Similarly than in the AMMI model the function `plot()` can be used to generate diagnostic plots of residuals of the model. The normality of the random effects may be also obtained by using `type = "re"`. Let's do it.

```
plot(WAASB_model, type = "re", ncol = 4)
```

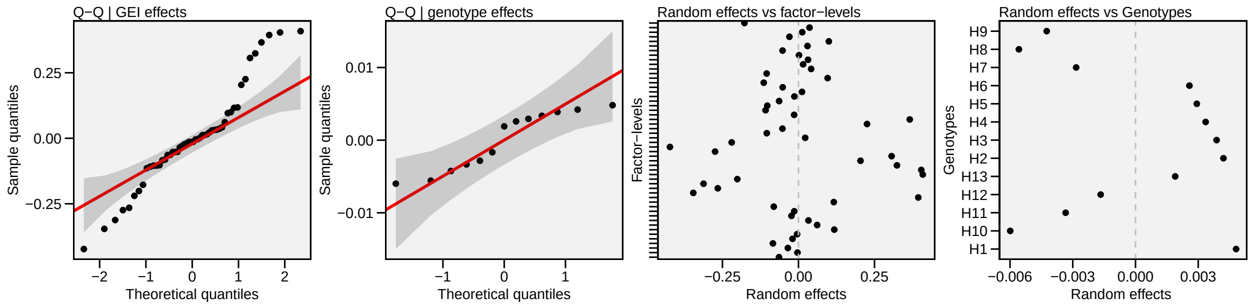

#### Curiosity

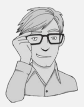

Residual plots are not widely used in MET analysis to check model assumptions. These plots have many advantages over statistical methods, especially when hypothesis testing is made in data sets with a large sample size. An interesting discussion comparing statistical methods and plots in verifying model assumptions is presented by Kozak & Piepho (2017).

##### 8.6.2 Likelihood Ratio Tests

The significance of random effects in the model is verified with a Likelihood Ratio Tests (LRT). Statistics and p-values for the LRT can be obtained with the function `get_model_data()`

```
get_model_data(WAASB_model, what = "pval_lrt")
```

```
# A tibble: 3 x 5
  model             PH            ED       TKW      NKR
  <chr>          <dbl>         <dbl>     <dbl>    <dbl>
1 Complete NA         NA            NA        NA
2 Genotype  9.39e- 1  0.299          1.00e+ 0  0.787
3 Gen:Env   1.09e-13  0.0000000169  4.21e-10  0.00404
```

##### 8.6.3 Variance components

The variance components for the random effects in the model can be obtained using

```
get_model_data(WAASB_model, what = "vcomp")
```

```
# A tibble: 3 x 5
  Group           PH    ED     TKW   NKR
  <chr>        <dbl> <dbl>   <dbl> <dbl>
1 GEN       0.000456 0.556 8.27e-7 0.187
2 GEN:ENV   0.0425   2.82  1.15e+3 2.96
3 Residual  0.0224   2.59  9.19e+2 7.85
```

##### 8.6.4 Genetic parameters

Beyond the variance components for the declared random effects, some important parameters are also calculated.

- **The broad-sense heritability, $h_g^2$, estimated by**

$$h_g^2 = \frac{\hat\sigma_g^2}{\hat\sigma_g^2 + \hat\sigma_i^2 + \hat\sigma_e^2}$$

where $\hat\sigma_g^2$ is the genotypic variance; $\hat\sigma_i^2$ is the genotype-by-environment interaction variance; and $\hat\sigma_e^2$ is the residual variance.

- **The coefficient of determination of the interaction effects, $r_i^2$, estimated by**

$$r_i^2 = \frac{\hat\sigma_i^2}{\hat\sigma_g^2 + \hat\sigma_i^2 + \hat\sigma_e^2}$$

- **The heritability on the mean basis, $h^2_{gm}$, estimated by**

$$h^2_{gm} = \frac{\hat{\sigma}^2_g}{[\hat{\sigma}^2_g + \hat{\sigma}^2_i / e + \hat{\sigma}^2_e / (eb)]}$$

where $e$ and $b$ are the number of environments and blocks, respectively;

- **The accuracy of selection, *Ac*, estimated by**

$$Ac = \sqrt{h^2_{gm}}$$

- **The genotype-environment correlation, $r_{ge}$, estimated by**

$$r_{ge} = \frac{\hat{\sigma}^2_g}{\hat{\sigma}^2_g + \hat{\sigma}^2_i}$$

- **The genotypic coefficient of variation and the residual coefficient of variation estimated, respectively, by**

$$CVg = \left(\sqrt{\hat{\sigma}^2_g}/\mu\right) \times 100$$

and

$$CVr = \left(\sqrt{\hat{\sigma}^2_e}/\mu\right) \times 100$$

where $\mu$ is the grand mean.

```
get_model_data(WAASB_model, what = "genpar")
```

```
# A tibble: 15 x 5
   Parameters                 PH      ED      TKW     NKR
   <chr>                   <dbl>   <dbl>    <dbl>   <dbl>
 1 GEI variance           0.0425    2.82   1.15e+ 3  2.96
 2 GEI (%)               65.0      47.2    5.55e+ 1 26.9
 3 Genotypic variance     0.000456  0.556  8.27e- 7  0.187
 4 Gen (%)                0.698     9.32   4.00e- 8  1.70
 5 Residual variance      0.0224    2.59   9.19e+ 2  7.85
 6 Res (%)               34.3      43.4    4.45e+ 1 71.4
 7 Phenotypic variance    0.0654    5.97   2.06e+ 3 11.0
 8 Heritability           0.00698   0.0932 4.00e-10  0.0170
 9 GEIr2                  0.650     0.472  5.55e- 1  0.269
10 Heribatility of means  0.0352    0.376  2.28e- 9  0.118
11 Accuracy               0.188     0.614  4.77e- 5  0.344
```

```
12 rge                    0.655    0.521  5.55e- 1  0.274
13 CVg                    0.860    1.51   2.68e- 4  1.34
14 CVr                    6.03     3.25   8.95e+ 0  8.69
15 CV ratio               0.143    0.463  3.00e- 5  0.154
```

##### 8.6.5 Predicted means

To obtain the predicted means for each genotypes, simply use the argument `what = 'blupg'` in the function `get_model_data()`.

```
get_model_data(WAASB_model, what = "blupg")
```

```
# A tibble: 13 x 5
   gen      PH    ED   TKW   NKR
   <fct> <dbl> <dbl> <dbl> <dbl>
 1 H1     2.49  50.2  339.  32.2
 2 H10    2.48  49.1  339.  32.3
 3 H11    2.48  49.2  339.  32.3
 4 H12    2.48  49.2  339.  32.0
 5 H13    2.49  49.9  339.  32.1
 6 H2     2.49  50.1  339.  32.2
 7 H3     2.49  49.5  339.  32.1
 8 H4     2.49  49.4  339.  32.6
 9 H5     2.49  49.7  339.  32.4
10 H6     2.49  50.3  339.  32.3
11 H7     2.48  49.5  339.  32.2
12 H8     2.48  49.1  339.  32.1
13 H9     2.48  48.8  339.  32.3
```

In the same way, use `what = 'blupge'` to obtain the predicted means for each genotype-environment combination.

```
WAASB_model %>%
  get_model_data(what = "blupge") %>%
  head() # First 6 rows
```

```
# A tibble: 6 x 6
  ENV   GEN      PH    ED   TKW   NKR
  <fct> <fct> <dbl> <dbl> <dbl> <dbl>
1 A1    H1     2.73  51.4  377.  33.5
2 A1    H10    2.78  52.7  369.  32.5
3 A1    H11    2.75  49.5  355.  34.3
4 A1    H12    2.71  50.3  339.  33.2
5 A1    H13    2.78  53.1  366.  33.3
6 A1    H2     2.79  52.0  335.  33.7
```

##### 8.6.6 BLUP-based stability statistics

Colombari Filho et al. (2013) have shown the use of three BLUP-based indexes for selecting genotypes with performance and stability. This method have been frequently used for analyzing MET data (Candido et al., 2018; Dias et al., 2018; Torres et al., 2018; Martins et al., 2019; Rosado et al., 2019; Vasconcelos, Echer, Kliemann, & Lang, 2019).

The first is the harmonic mean of genotypic values (HMGV) a stability index that considers the genotype with the highest harmonic mean of BLUPs across environments as the most stable, as follows:

$$HMGV_i = \frac{1}{e}\sum_{j=1}^{e}\frac{1}{BLUP_{ij}}$$

The second is the relative performance of genotypic values (RPGV), an adaptability index estimated as follows:

$$RPGV_i = \frac{1}{e}\sum_{j=1}^{e}BLUP_{ij}/\mu_j$$

The third and last is the harmonic mean of relative performance of genotypic values (HMRPGV), a simultaneous selection index for stability, adaptability and mean performance, estimated as follows:

$$HMRPGV_i = \frac{1}{e}\sum_{j=1}^{e}\frac{1}{BLUP_{ij}/\mu_j}$$

These BLUP-based stability indexes are computed with the function `Resende_indexes()`. We can also get the results using `get_model_data()`. For example, to get the HMRPGV in the same unit of the response variable we should use `what = "HMRPGV_Y"`.

```
stab_blup <- Resende_indexes(WAASB_model)
get_model_data(stab_blup, what = "HMRPGV_Y")
```

```
# A tibble: 13 x 5
   gen      PH    ED   TKW   NKR
   <chr> <dbl> <dbl> <dbl> <dbl>
 1 H1     2.59  50.9  358.  32.2
 2 H10    2.33  48.5  320.  32.3
 3 H11    2.40  48.9  333.  32.7
 4 H12    2.44  48.7  320.  31.2
 5 H13    2.53  50.4  340.  31.5
 6 H2     2.58  50.7  350.  32.0
 7 H3     2.56  49.4  342.  31.7
 8 H4     2.55  49.3  343.  33.8
 9 H5     2.55  49.8  340.  33.2
10 H6     2.54  51.2  357.  32.4
```

```
11 H7      2.41  49.5  341.   31.8
12 H8      2.33  48.5  320.   31.6
13 H9      2.37  47.8  310.   32.4
```

The WAASB index, an acronym for **W**eighted **A**verage of the **A**bsolute **S**cores from the Singular Value Decomposition of the **BLUP**s for genotype-vs-environment interaction effects obtained by a Linear Mixed-effect model was proposed by T. Olivoto, Lúcio, et al. (2019). The WAASB was computed with the function `waasb()`.

$$WAASB_i = \sum_{k=1}^{P} |\lambda_k^{0.5} a_{ik} \times \theta_k| / \sum_{k=1}^{P} \theta_k$$

where $WAASB_i$ is the weighted average of absolute scores of the genotype $i$; $\lambda_k^{0.5} a_{ik}$ is the scores of the genotype $i$ in the IPCA $k$; and $\theta_k$ is the explained variance of the IPCA $k$ for $k = 1, 2, .., p$, being $p = min(g - 1; e - 1)$. We can now get the values using the argument `what = "WAASB"` in the function `get_model_data()`.

```
get_model_data(WAASB_model, what = "WAASB")
```

```
# A tibble: 13 x 5
   gen      PH    ED   TKW   NKR
   <fct> <dbl> <dbl> <dbl> <dbl>
 1 H1    0.319 0.695 3.49  0.545
 2 H10   0.287 0.879 2.47  0.291
 3 H11   0.210 0.579 0.801 0.533
 4 H12   0.298 0.344 1.59  0.433
 5 H13   0.258 0.758 0.423 0.547
 6 H2    0.312 0.990 3.86  0.271
 7 H3    0.340 0.364 3.08  0.172
 8 H4    0.268 0.321 2.77  0.615
 9 H5    0.171 0.465 0.476 0.562
10 H6    0.232 0.528 0.591 1.07
11 H7    0.209 0.301 2.55  0.369
12 H8    0.334 0.640 4.50  0.612
13 H9    0.208 0.913 5.13  0.632
```

#### 8.7 Cross-validation for AMMI and BLUP models

##### 8.7.1 Theory

Gauch (2013) pointed out that predictive accuracy merits special attention for model diagnosis in MET analysis. Due to the great data processing power of the current computers, it is reasonable to affirm that the choice of the best method to predict yield (or other response variables) should be based on the predictive ability assessment in each situation. To our current knowledge, no one other R package performs cross-validation for AMMI and BLUP models in MET analysis. In `metan` the predictive accuracy of both AMMI and BLUP models may be obtained using a cross-validation procedure implemented by the functions `cv_ammi()`, `cv_ammif()` (AMMI model) and `cv_blup()`

(BLUP model). The function `cv_ammif()` provides a complete cross-validation procedure for all members of the AMMI model family (AMMI0-AMMIF) using replicate-based data. If the user, for some reason, needs to compute cross-validation for a specific member of the AMMI-family model, then the function `cv_ammi()` can be used. Automatically the first validation is carried out considering the AMMIF (all possible axis used). Considering this model, the original data set is split up into two sets: *"training"* set and *"validation"* sets. The *"training"* set has all combinations (genotype-*vs*-environment) with R-1 replicates (R). The *"validation"* set has one replicate that was not included in the *"training"* set.

The splitting of the data set into *"training"* and *"validation"* sets depends on the experimental design. For a Randomized Complete Block Design (default option), completely blocks are randomly selected within environments, as shown by T. Olivoto, Lúcio, et al. (2019). The remaining block serves as validation data. If `design = "CRD"` is informed, thus declaring that a completely randomized design was used, single observations are randomized for each treatment (genotype-by-environment combination). This is the same procedure suggested by Gauch (1988). The estimated values for each member of the AMMI model family in each re-sampling cycle are compared with the observed values in the validation data. Then, the Root Mean Square Prediction Difference is computed as follows:

$$RMSPD = \left[\left(\sum_{i=1}^{n} (\hat{y}_{ij} - y_{ij})^2\right)/n\right]^{0.5}$$

where $\widehat{y}_{ij}$ is the model predicted value; and $y_{ij}$ is the observed value in the validation set. The number of random selection of blocks/replicates ($n$) is defined in the argument `nboot`. At the end of the $n$ cycles for all models, a list with all estimated RMSPD and the average of RMSPD is returned and we can use `plot()` to create a boxplot with the results, as shown in Figure 1i of the paper.

##### 8.7.2 Computing the cross-validation procedures

As an example, we will compute a cross-validation test for AMMI0, AMMI2, and AMMIF (9 axes) models, as well as for the full AMMI-family model and BLUP model. By default, 200 re-samples are performed for each cross-validation procedure.

**Warning**

The cross-validation procedure may take several minutes to run. All the examples below were run in ~7 min using an 8-GB-RAN machine with Intel(R) Core(TM) i-5-3210M CPU at 2.5 GHz

```
AMMI0 <- cv_ammi(data_ge, ENV, GEN, REP, GY, naxis = 0) # AMMI0
AMMI2 <- cv_ammi(data_ge, ENV, GEN, REP, GY, naxis = 2) # AMMI2
AMMI9 <- cv_ammi(data_ge, ENV, GEN, REP, GY, naxis = 9) # AMMI9
AMMIF <- cv_ammif(data_ge, ENV, GEN, REP, GY) #AMMI0-AMMIF
```

The function `cv_blup()` provides a cross-validation of replicate-based data using mixed models. By default, complete blocks are randomly selected for each environment. Using the argument `random` it is possible to choose the random effects of the model, as shown below.

- **Genotype and genotype-vs-environment as random effects**

```
BLUP_g <- cv_blup(data_ge, ENV, GEN, REP, GY, random = "gen")
```

- **Environment, replication-within-environment and interaction as random effects**

```
BLUP_e <- cv_blup(data_ge, ENV, GEN, REP, GY, random = "env")
```

- **A random model (all terms as random effects)**

```
BLUP_ge <- cv_blup(data_ge, ENV, GEN, REP, GY, random = "all")
```

##### 8.7.3 Visualizing the results

We can use the function `bind_cv()` to bind the cross-validation objects aiming at showing the mean results in a table. Let's do it.

```
bind_means <- bind_cv(AMMIF, BLUP_g, BLUP_e, BLUP_ge, bind = "means")
print(bind_means$RMSPD)
```

```
# A tibble: 13 x 6
   MODEL         mean     sd      se  Q2.5 Q97.5
   <fct>        <dbl>  <dbl>   <dbl> <dbl> <dbl>
 1 BLUP_e_RCBD  0.401 0.0251 0.00178 0.354 0.446
 2 BLUP_g_RCBD  0.406 0.0276 0.00195 0.349 0.456
 3 BLUP_ge_RCBD 0.406 0.0240 0.00170 0.362 0.451
 4 AMMI2        0.413 0.0257 0.00182 0.366 0.457
 5 AMMI3        0.414 0.0233 0.00165 0.367 0.455
 6 AMMI4        0.414 0.0231 0.00163 0.368 0.458
 7 AMMI5        0.423 0.0229 0.00162 0.380 0.464
 8 AMMI6        0.424 0.0224 0.00159 0.378 0.467
 9 AMMIF        0.425 0.0209 0.00148 0.387 0.462
10 AMMI7        0.427 0.0211 0.00149 0.381 0.465
11 AMMI8        0.427 0.0212 0.00150 0.387 0.466
12 AMMI1        0.429 0.0264 0.00187 0.372 0.474
13 AMMI0        0.431 0.0310 0.00219 0.375 0.491
```

We can also combine all results to create an object of class `cvalidation` suitable for creation of a boxplot with `plot()`

```
bind_ammi <- bind_cv(AMMI0, AMMI2, AMMI9)
bind_ammi_blup <- bind_cv(AMMIF, BLUP_g, BLUP_e, BLUP_ge)
a <- plot(bind_ammi)
b <- plot(bind_ammi_blup)
c <- plot(bind_ammi_blup, order_box = TRUE)

arrange_ggplot(a, b, c, nrow = 1, labels = letters[1:3])
```

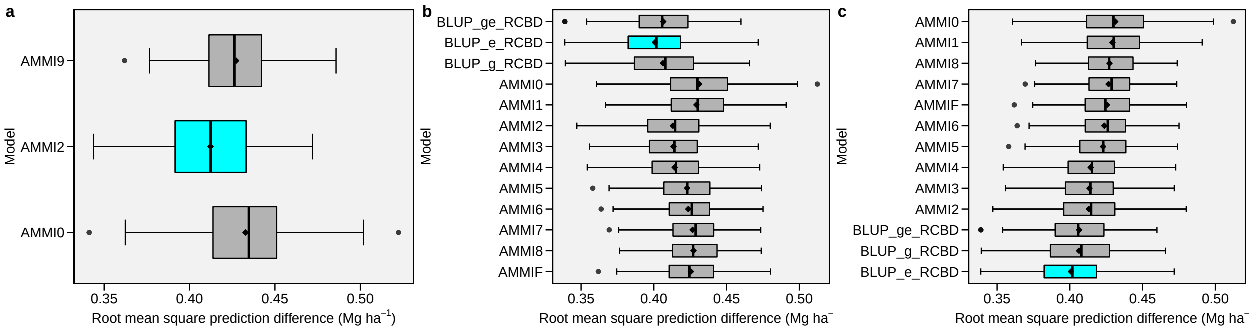

#### 8.8 GGE analysis

##### 8.8.1 The model

Genotype plus Genotype-vs-Environment interaction (GGE) model has been widely used to genotype evaluation and mega-environment identification in multi-environment trials (MET). This model considers a GGE (i.e., G + GE) biplot, which is constructed by the first two symmetrically scaled principal components (PC1 and PC2) derived from singular value decomposition of environment-centered MET data. The GGE biplot graphically displays G plus GE of a MET in a way that facilitates visual genotype evaluation and mega-environment identification (Yan, Kang, Ma, Woods, & Cornelius, 2007).

The mean yield of genotype $i$ in environment $j$ is commonly described by a general linear model

$$\hat{y}_{ij} + \mu + \alpha_i + \beta_j + \phi_{ij}$$

where $\hat{y}_{ij}$ is the mean yield of genotype $i$ in environment $j$, $i = 1, ...g; j = 1, ...e$ being $g$ and $e$ the numbers of genotypes and environments, respectively; $\mu$ is the grand mean; $\alpha_i$ is the main effect of the genotype $i$; $\beta_j$ is the main effect of the environment $j$, and $\phi_{ij}$ is the interaction effect between genotype $i$ and environment $j$. Subjecting the $\phi_{ij}$ to Singular Value Decomposition (SVD) results in the AMMI model. The deletion of $\alpha_i$ allows the variation explained by this term to be absorbed into the $\phi_{ij}$ term. In the Genotype plus Genotype-vs-Environment interaction (GGE) model the $\alpha_i$ term is deleted from the above model and then the environment-centered data matrix, $\phi_{ij}$, is subjected to SVD (Yan & Kang, 2003; Yan et al., 2007). Explicitly, we have

$$\phi_{ij} = \hat{y}_{ij} - \mu - \beta_j = \sum_{k=1}^{p} \xi_{ik}^{*} \eta_{jk}^{*}$$

where $\xi_{ik}^{*} = \lambda_k^{\alpha} \xi_{ik}$; $\eta_{jk}^{*} = \lambda_k^{1-\alpha} \eta_{jk}$ being $\lambda_k$ the $k$th eigenvalue from the SVD ($k = 1, ...p$), with $p \le min(e, g)$; $\alpha$ is the the singular value partition factor for the Principal Component (PC) $k$; $\xi_{ik}^{*}$ and $\eta_{jk}^{*}$ are the PC scores for genotype $i$ and environment $j$, respectively.

##### 8.8.2 Model options

The function `gge()` is used to produce a GGE model. According to Yan & Kang (2003), the function supports four methods of data centering, two methods of data scaling and three options for singular value partitioning:

- **Centering methods available**
  - `0` or `"none"` for no centering;
  - `1` or `"global"` for global centered (E+G+GE);
  - `2` or `"environment"` (default), for environment-centered (G+GE);
  - `3` or `"double"` for double centred (GE). A biplot cannot be produced with models produced without centering.
- **Scaling methods available**
  - `0` or `"none"` (default) for no scaling;
  - `1` or `"sd"` where each value is divided by the standard deviation of its corresponding environment (column). This will put all testers roughly the same range of values.
- **Singular Value Partitioning methods available**
  - `1` or `"genotype"` The singular value is entirely partitioned into the genotype eigenvectors, also called row metric preserving;
  - `2` or `"environment"` (default) the singular value is entirely partitioned into the environment eigenvectors, also called column metric preserving;
  - `3` or `"symmetrical"` The singular value is symmetrically partitioned into the genotype and the environment eigenvectors This SVP is most often used in AMMI analysis and other biplot analysis, but it is not ideal for visualizing either the relationship among entries or that among the testers.

The function `gge()` is used to fit the model. This function produces a GGE model based on both a two-way table with genotypes in the rows and environments in columns or a data frame containing at least the columns for genotypes, environments and the response variable(s).

```
# Using a data frame
gge_model <- gge(data_ge, ENV, GEN, GY)
```

The model above was fitted considering (i) column metric preserving (where the singular value is entirely partitioned into the environment eigenvectors); (ii) environment centered (the biplot will contain mixed information of G + GEI); and no scaling method. To change these default settings, use the arguments `svp`, `centering`, and `scaling`, respectively. Please, note that in the second example the argument `table` was set to `TRUE` to indicate that the input data is a two-way table.

##### 8.8.3 Predict a GGE model

The S3 generic function `predict()` is used to predict the response variable of a two-way table based on a GGE model. This prediction is based on the number of principal components used (Yan et al., 2007).

```
predict(gge_model)
```

```
$GY
# A tibble: 10 x 14
     E1   E10   E11   E12   E13   E14    E2    E3    E4    E5    E6    E7    E8
```

```
 * <dbl> <dbl> <dbl> <dbl> <dbl> <dbl> <dbl> <dbl> <dbl> <dbl> <dbl> <dbl> <dbl>
 1  2.51  2.14 1.35   1.59  2.95  1.76  3.20  4.14  3.78  3.97  2.70  1.95  2.56
 2  1.93  1.62 0.988  1.07  2.07  1.66  3.08  3.99  3.78  3.28  2.43  1.69  2.48
 3  2.67  2.33 1.47   1.75  3.09  1.82  3.19  4.04  3.59  4.03  2.70  2.08  2.54
 4  2.83  2.44 1.56   1.88  3.40  1.83  3.26  4.18  3.73  4.32  2.83  2.10  2.59
 5  2.50  2.25 1.39   1.64  2.71  1.83  3.09  3.81  3.32  3.65  2.52  2.11  2.47
 6  2.32  2.06 1.27   1.46  2.52  1.78  3.09  3.87  3.46  3.54  2.49  1.99  2.47
 7  2.40  2.11 1.31   1.52  2.65  1.78  3.12  3.92  3.52  3.66  2.54  2.00  2.49
 8  2.78  2.49 1.56   1.88  3.17  1.87  3.17  3.93  3.40  4.04  2.68  2.21  2.52
 9  2.99  2.54 1.64   2.00  3.71  1.84  3.33  4.33  3.89  4.61  2.96  2.11  2.64
10  2.27  1.78 1.15   1.30  2.82  1.64  3.28  4.43  4.28  4.03  2.79  1.65  2.62
# ... with 1 more variable: E9 <dbl>
```

##### 8.8.4 GGE Biplot

The generic function `plot()` is used to generate a biplot using as input a fitted model of class `gge`. The type of biplot is chosen by the argument `type` in the function. Ten biplots types are available according to Yan & Kang (2003).

- `type = 1` A basic biplot.
- `type = 2` Mean performance vs. stability.
- `type = 3` Which-won-where.
- `type = 4` Discriminativeness vs. representativeness.
- `type = 5` Examine an environment.
- `type = 6` Ranking environments.
- `type = 7` Examine a genotype.
- `type = 8` Ranking gentoypes.
- `type = 9` Compare two genotypes.
- `type = 10` Relationship among environments.

In this material, for each biplot type, two graphics are produced. One with the default settings and the other to show some graphical options of the function.

- **Biplot type 1: A basic biplot** This is the default setting in the function plot, thus, this biplot is produced by just calling `plot(model)`, as shown below.

```
a <- plot(gge_model)
b <- plot(gge_model,
          col.gen = "orange2",
          size.text.env = 2)
arrange_ggplot(a, b, labels = letters[1:2])
```

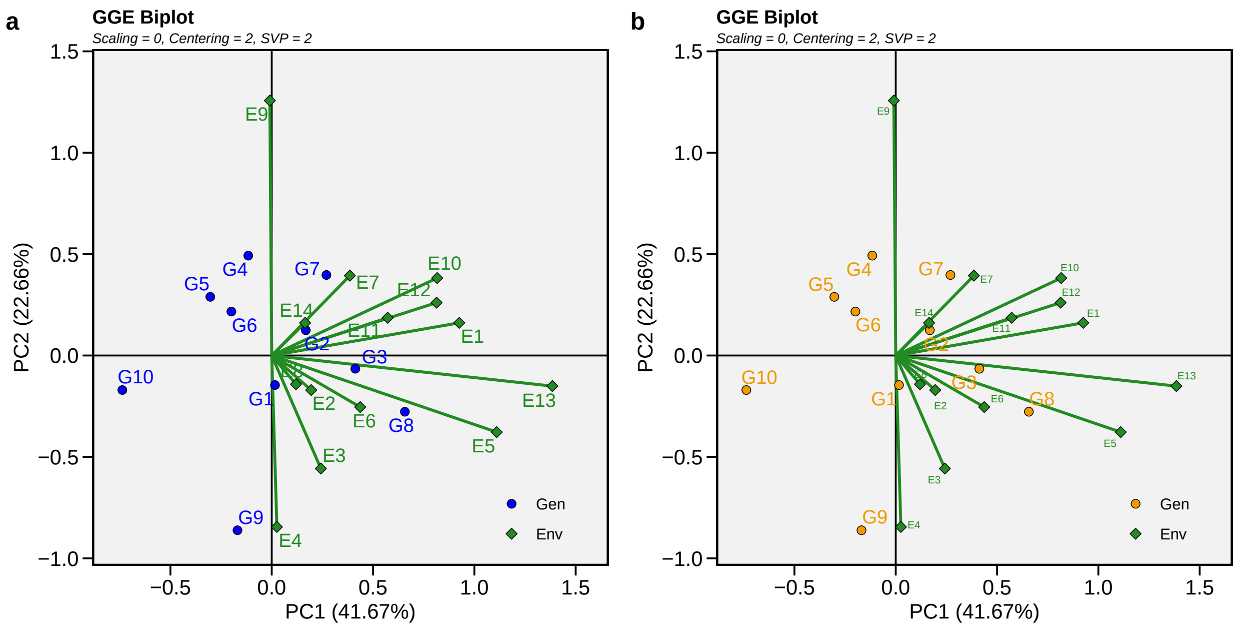

- **Biplot type 2: Mean performance vs. stability**

In this biplot, the visualization of the mean and stability of genotypes is achieved by drawing an average environment coordinate (AEC) on the genotype-focused biplot. First, an average environment, represented by the small circle, is defined by the mean PC1 and PC2 scores of the environments. The line that passes through the biplot origin and the AEC may be called the average. The projections of genotype markers onto this axis should, therefore, approximate the mean yield of the genotypes. Thus, the G8 was clearly the highest-yielding genotype, on average.

The AEC ordinate is the line that passes through the biplot origin and is perpendicular to the AEC abscissa. Therefore, if the AEC abscissa represents the G, the AEC ordinate must approximate the GEI associated with each genotype, which is a measure of variability or instability of the genotypes (Yan et al., 2007). A greater projection onto the AEC ordinate, regardless of the direction, means greater instability. In our example, G3 was found to be the most stable and the second most productive genotype, while G9 had great instability.

```
gge_model <- gge(data_ge, ENV, GEN, GY, svp = "genotype")
a <- plot(gge_model, type = 2)
b <- plot(gge_model,
          type = 2,
          col.gen = "black",
          col.env = "gray",
          axis_expand = 1.5)
arrange_ggplot(a, b, labels = letters[1:2])
```

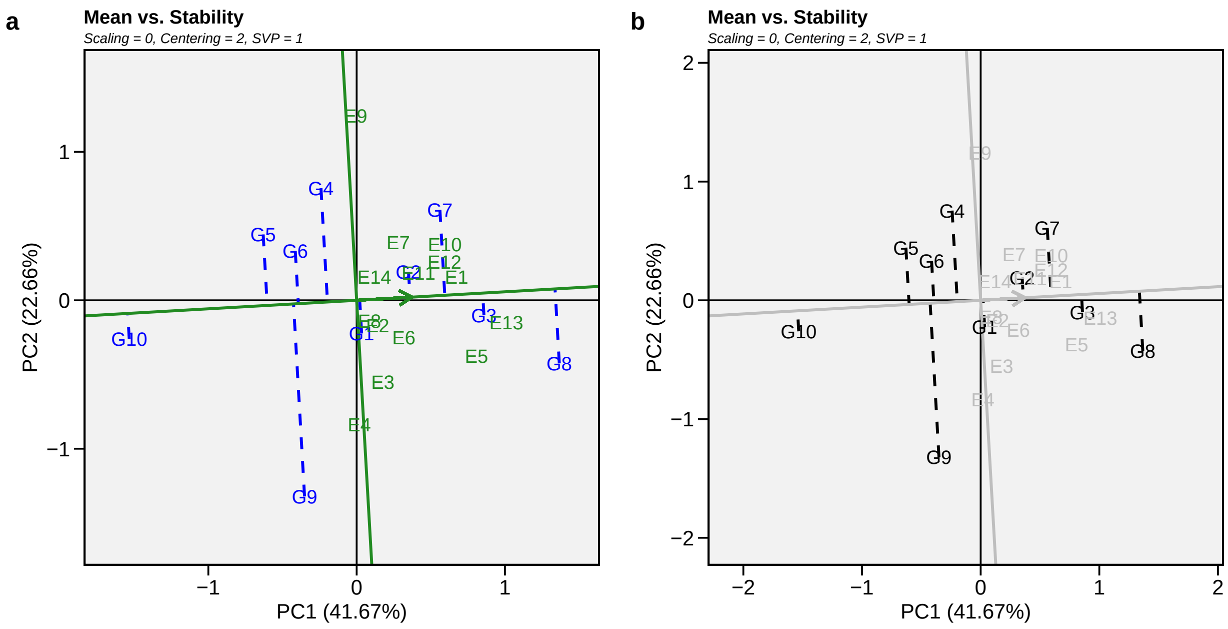

- **Biplot type 3: Which-won-where**

In this biplot, a polygon is drawn joining the genotypes (G7, G8, G9, G10, and G4) that are located farthest from the biplot origin so that all other genotypes are contained in the polygon. The vertex genotypes have the longest vectors, in their respective directions, which is a measure of responsiveness to environments. The vertex genotypes are, therefore, among the most responsive genotypes; all other genotypes are less responsive in their respective directions. A genotype located at the origin would rank the same in all environments and is not at all responsive to the environments.

The perpendicular lines to the sides of the polygon divide the biplot into sectors. Each sector has a vertex genotype. For example, the sector with the vertex genotype G4 may be referred to as the G4 sector; and one environment (E9), fell in this sector. As a rule, the vertex genotype is the highest-yielding genotype in all environments that share the sector with it (Yan et al., 2007). In this case, G4 was the highest-yielding in E9.

```
gge_model <- gge(data_ge, ENV, GEN, GY, svp = "symmetrical")
a <- plot(gge_model, type = 3)
b <- plot(gge_model,
          type = 3,
          size.shape.win = 5,
          large_label = 6,
          col.gen = "black",
          col.env = "gray",
          annotation = FALSE,
          title = FALSE)
arrange_ggplot(a, b, labels = letters[1:2])
```

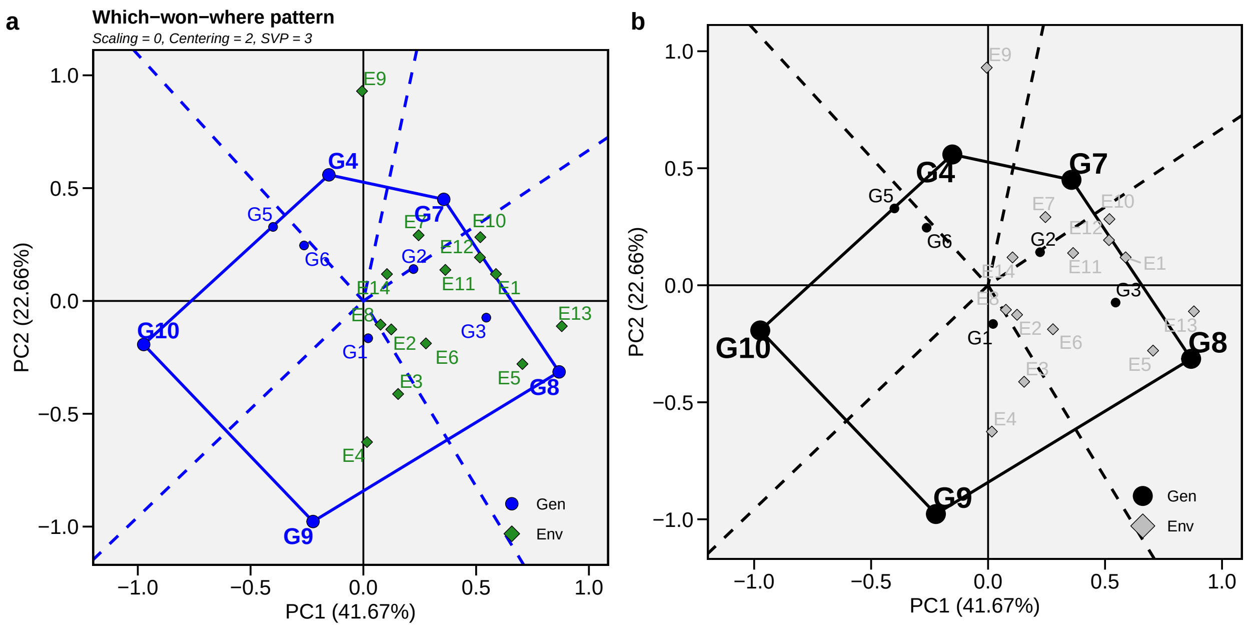

- **Biplot type 4: Discriminativeness vs. representativeness**

```
a <- plot(gge_model, type = 4)
b <- plot(gge_model,
          type = 4,
          plot_theme = theme_gray())+
     theme(legend.position = "bottom")
arrange_ggplot(a, b, labels = letters[1:2])
```

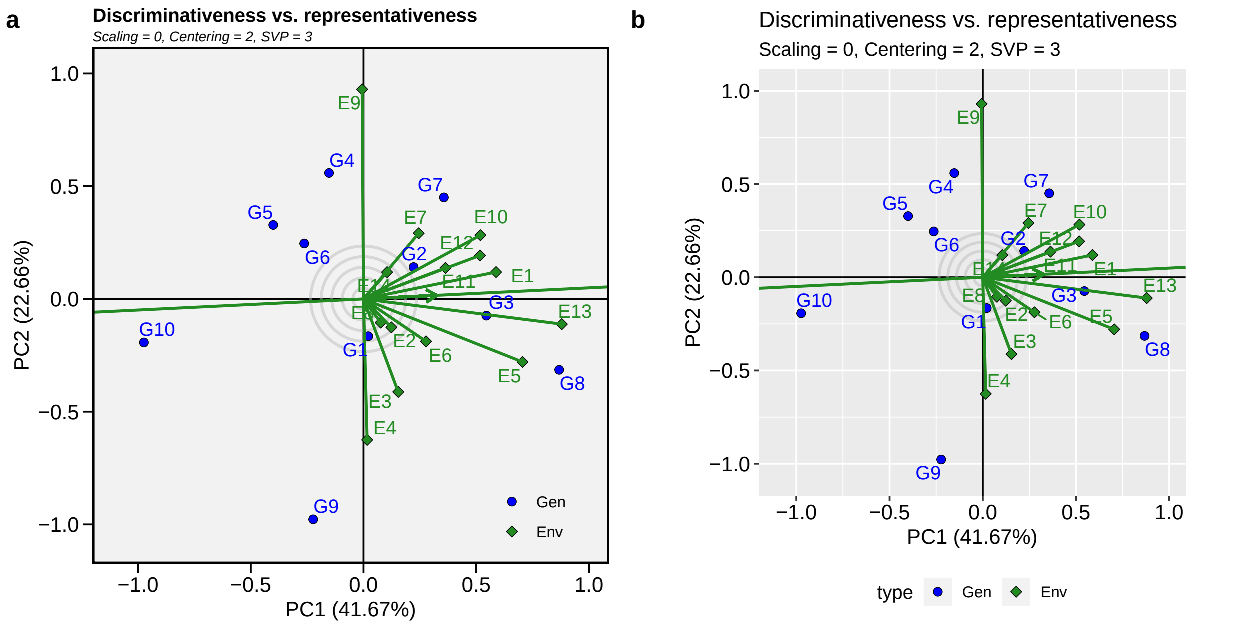

- **Biplot type 5: Examine an environment**

Identifying genotypes most adapted to an environment can be easily achieved via a GGE biplot. For example, to visualize the performance of different genotypes in a given environment, e.g., E10, simply draw a line that passes through the biplot origin and the marker of E10. The genotypes can be ranked according to their projections onto the E10 axis based on their performance in E10, in the direction pointed by the arrow. In our example, at E10, the highest-yielding genotype was G8, and the lowest-yielding genotype was G10, and the order of the genotypes were G8 > G7 > G3 > G2 > G4 > G1 > G6 > G5 > G9 > G10.

```
gge_model <- gge(data_ge, ENV, GEN, GY, svp = "symmetrical")
a <- plot(gge_model, type = 5, sel_env = "E10")
b <- plot(gge_model,
          type = 5,
          sel_env = "E10",
          col.gen = "black",
          col.env = "black",
          size.text.env = 10,
          axis_expand = 1.5)
arrange_ggplot(a, b, labels = letters[1:2])
```

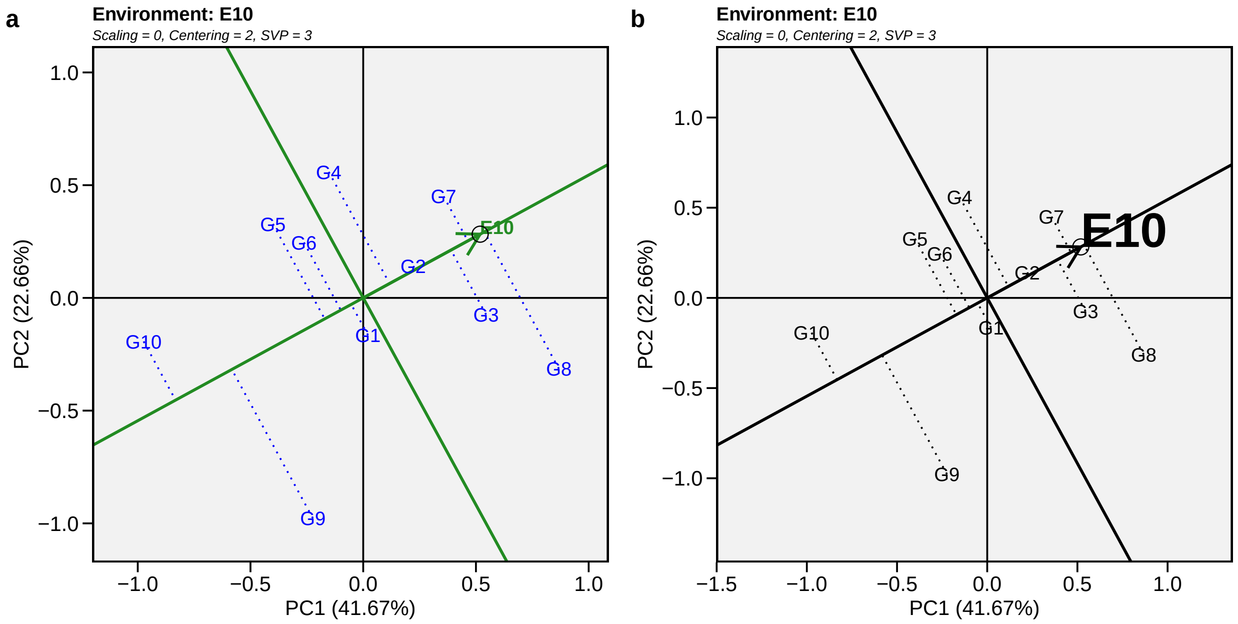

- **Biplot type 6: Ranking environments**

In this biplot, the *"ideal"* environment is used as the center of a set of concentric lines that serve as a ruler to measure the distance between an environment and the ideal environment. since the main focus in this biplot is environments, then, the singular value partition used is `"environment"` (default). It can be seen that E13 is the closest to the ideal environment, and, therefore, is most desirable of all 14 environments. E4 and E9 were the least desirable test environments.

```
gge_model <- gge_model <- gge(data_ge, ENV, GEN, GY)
a <- plot(gge_model, type = 6)
b <- plot(gge_model,
          type = 6,
          col.gen = "black",
          col.env = "black",
          size.text.env = 10,
          axis_expand = 1.5)
arrange_ggplot(a, b, labels = letters[1:2])
```

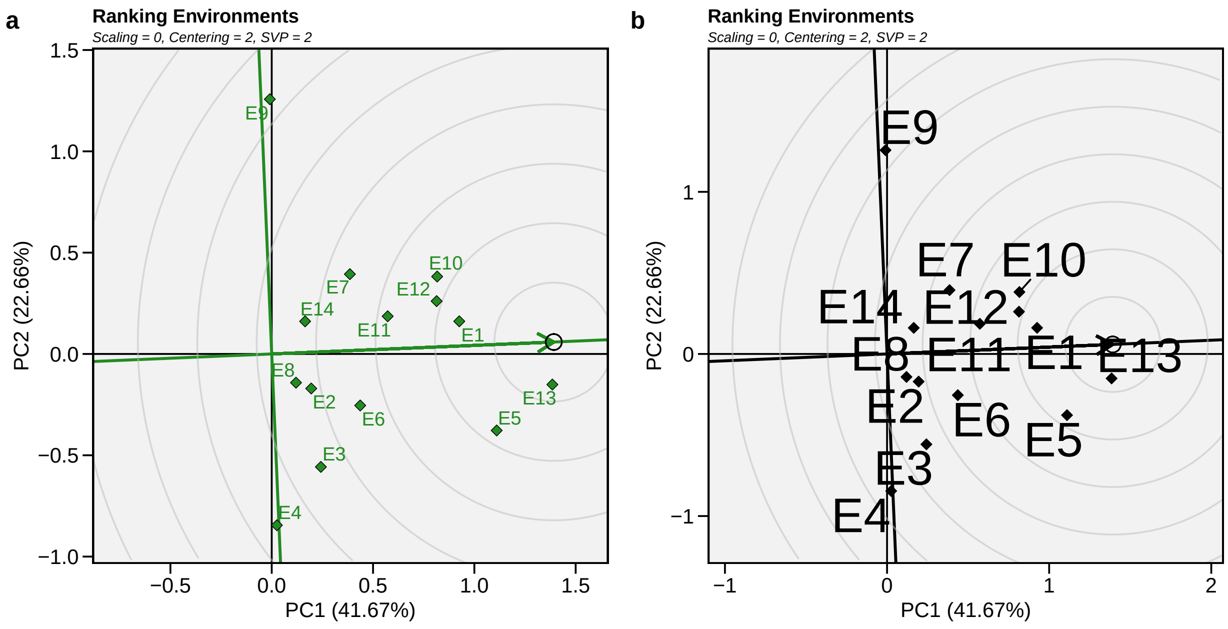

- **Biplot type 7: Examine a genotype**

Analogous to visualizing genotype performances in a given environment (biplot 5) visualization of the mean and stability of genotypes is achieved by drawing an average environment coordinate (AEC) on the genotype-focused biplot (Yan et al., 2007).

```
gge_model <- gge(data_ge, ENV, GEN, GY, svp = "genotype")
a <- plot(gge_model, type = 7, sel_gen = "G8")
b <- plot(gge_model,
          type = 7,
          sel_gen = "G8",
          col.gen = "black",
          col.env = "black",
          size.text.env = 10,
          axis_expand = 1.5)
arrange_ggplot(a, b, labels = letters[1:2])
```

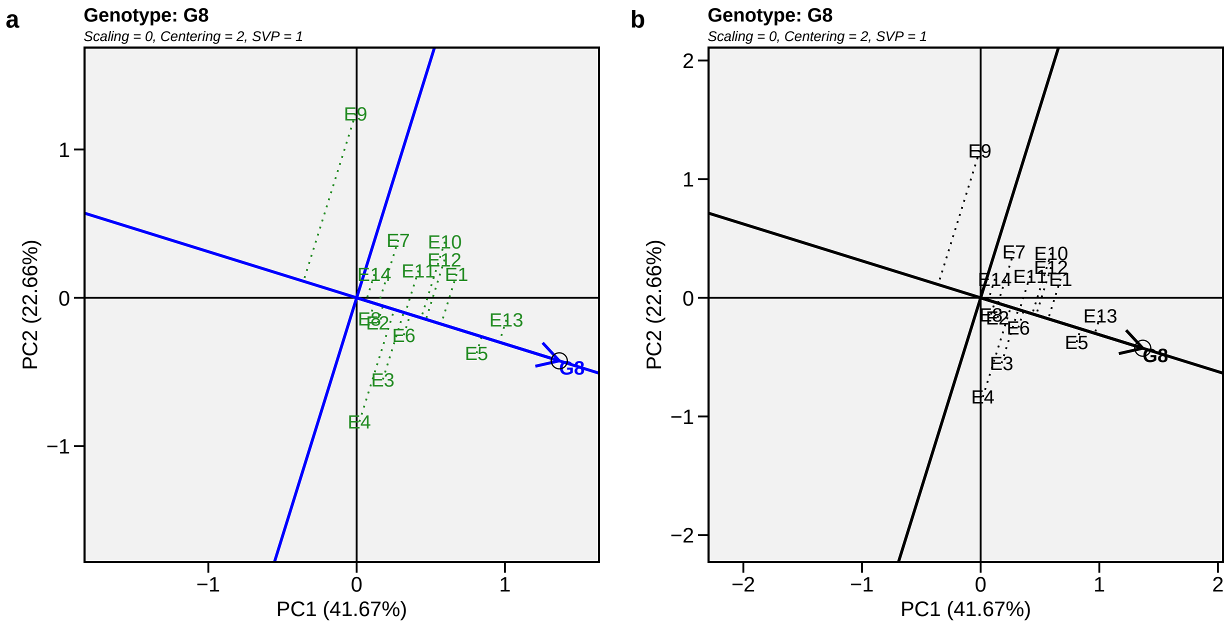

- **Biplot type 8: Ranking genotypes**

This biplot compares all genotypes with the *"ideal"* genotype. The ideal genotype, represented by the small circle with an arrow pointing to it, is defined as having the highest yield in all environments. That is, it has the highest mean yield and is absolutely stable. The genotypes are ranked based on their distance from the ideal genotype (Yan et al., 2007). In our example, G3 and G8 were found to outperform the other genotypes.

```
gge_model <- gge(data_ge, ENV, GEN, GY, svp = "genotype")
a <- plot(gge_model, type = 8)
b <- plot(gge_model,
          type = 8,
          col.gen = "black",
          col.env = "gray",
          size.text.gen = 6)
arrange_ggplot(a, b, labels = letters[1:2])
```

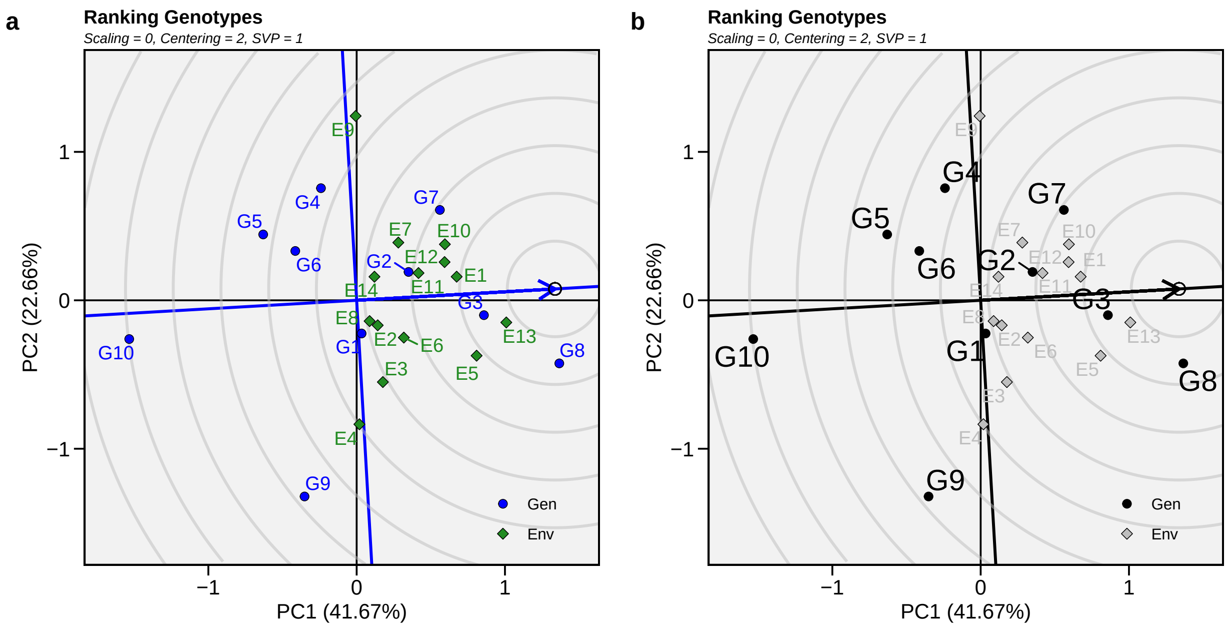

- **Biplot type 9: Compare two genotypes**

To compare two genotypes, for example, G10 and G8, draw a connector line to connect them and draw a perpendicular line that passes through the biplot origin and is perpendicular to the connector line. We see one environment –E9– is on the same side of the perpendicular line as G10, and the other 13 environments are on the other side of the perpendicular line, together with G8. This indicates that G10 yielded more than G8 in E9, but G8. yielded more than G10 in the other 13 environments (Yan et al., 2007).

```
gge_model <- gge(data_ge, ENV, GEN, GY, svp = "symmetrical")
a <- plot(gge_model, type = 9, sel_gen1 = "G8", sel_gen2 = "G10")
b <- plot(gge_model,
          type = 9,
          sel_gen1 = "G8",
          sel_gen2 = "G10",
          col.gen = "black",
          title = FALSE,
          annotation = FALSE)
arrange_ggplot(a, b, labels = letters[1:2])
```

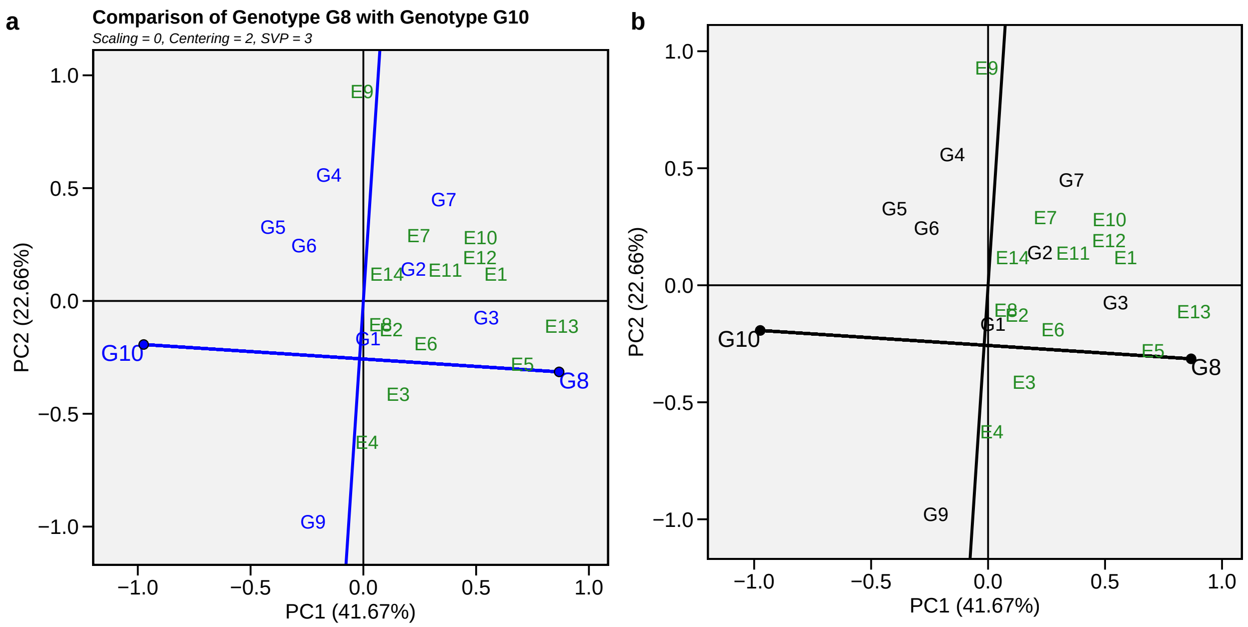

- **Biplot type 10: Relationship among environments**

```
gge_model <- gge(data_ge, ENV, GEN, GY)
a <- plot(gge_model, type = 10)
b <- plot(gge_model,
          type = 10,
          col.gen = "black",
          title = FALSE,
          annotation = FALSE)
arrange_ggplot(a, b, labels = letters[1:2])
```

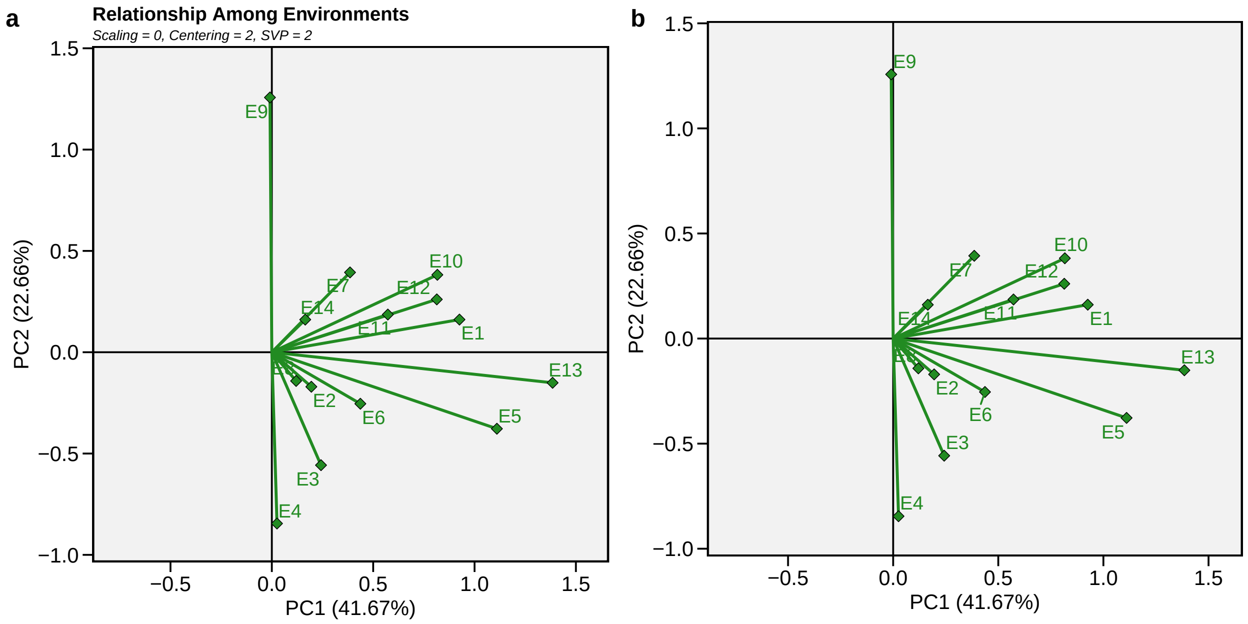

#### 8.9 Multi-trait stability index

The multi-trait stability index (*MTSI*) proposed by (T. Olivoto et al., 2019) is computed for evaluating simultaneous selection for stability and mean performance across many traits. The MTSI is computed considering the WAASBY index (T. Olivoto, Lúcio, et al., 2019), a superiority index based on mixed-effect models, as follows

$$WAASBY_i = \frac{(rG_i \times \theta_Y) + (rW_i \times \theta_S)}{\theta_Y + \theta_S}$$

where $WAASBY_i$ is the simultaneous selection index for the $i$-th genotype that weights between performance and stability; $rY_i$ and $rW_i$ are the rescaled values (0-100) for dependent variable and $WAASB$, respectively; $\theta_Y$ and $\theta_S$ are the weights for dependent variable and $WAASB$, respectively. Rescaled values are used to make $WAASB$ and $Y$ directly comparable. Assuming that the highest value for the dependent variable is better, say, for grain yield, the genotype with the highest mean will have $rY_i = 100$ after rescaling. On the other hand, if the lowest value is better, say, for lodging, the genotype with the lowest mean will have $rY_i = 100$ after rescaling. The genotype with the lowest $WAASB$ will then have $rW_i = 100$. The code below computes the WAASBY index with a higher weight for the mean performance (65 in argument `wresp`) and considering that high values are better for all variables (100 in argument `mresp`).

```
model <- waasb(data_ge2, ENV, GEN, REP,
               resp = c(KW, NKE, PH, EH, TKW),
               random = "gen", # Default
               wresp = rep(65, 5), # Defaults to 50
               mresp = rep(100, 5)) # Default
```

```
Evaluating variable KW 0 %
Evaluating variable NKE 25 %
Evaluating variable PH 50 %
Evaluating variable EH 75 %
Evaluating variable TKW 100 %
All variables with significant (p < 0.05) genotype-vs-environment interaction
Done!
```

To obtain the WAASBY indes for all variables in the model, use `get_model_data()` with the argument `what = 'WAASBY'`.

```
get_model_data(model, what = "WAASBY")
```

```
# A tibble: 13 x 6
   gen          KW   NKE    PH    EH   TKW
   <fct>     <dbl> <dbl> <dbl> <dbl> <dbl>
 1 H1     8.33e+ 1  35.1 69.3  65     77.2
 2 H10    4.66e+ 1  46.1 11.0  26.9   31.0
 3 H11    6.17e+ 1  41.7 42.7  30.7   59.1
 4 H12    2.32e+ 1  34.2 34.7  18.6   32.6
```

```
 5 H13    6.91e+ 1  59.8 64.4  50.9   70.7
 6 H2     6.88e+ 1  63.2 67.1  39.6   64.3
 7 H3     4.81e+ 1  19.2 59.4  47.9   57.5
 8 H4     7.85e+ 1  91.8 71.1  59.9   60.3
 9 H5     8.27e+ 1  83.6 88.7  67.2   71.1
10 H6     6.77e+ 1  36.7 74.0  68.0   96.9
11 H7     4.55e+ 1  36.5 46.0  49.4   60.6
12 H8     1.71e+ 1  22.0  3.75  1.63  17.9
13 H9    -9.24e-15  14.9 37.9  49.3    0
```

After computing the WAASBY index, the MTSI index can be computed with the function `mtsi()`

```
index <- mtsi(model,
              index = "waasby",
              mineval = 0.7,
              verbose = FALSE)
print(index)
```

```
-------------------- Correlation matrix used used in factor analysis -----------------
           KW       NKE        PH        EH       TKW
KW  1.0000000 0.7142147 0.7612267 0.6367073 0.8785741
NKE 0.7142147 1.0000000 0.5599209 0.3897729 0.4408535
PH  0.7612267 0.5599209 1.0000000 0.8744239 0.7799807
EH  0.6367073 0.3897729 0.8744239 1.0000000 0.6653946
TKW 0.8785741 0.4408535 0.7799807 0.6653946 1.0000000

---------------------------- Principal component analysis -----------------------------
# A tibble: 5 x 4
  PC    Eigenvalues `Variance (%)` `Cum. variance (%)`
  <chr>       <dbl>          <dbl>               <dbl>
1 PC1        3.713          74.26               74.26
2 PC2        0.7118         14.24               88.49
3 PC3        0.4244          8.487              96.98
4 PC4        0.1043          2.087              99.07
5 PC5        0.04671         0.9342            100

--------------------------------- Initial loadings ----------------------------------
# A tibble: 5 x 3
  VAR       PC1     PC2
  <chr>   <dbl>   <dbl>
1 KW    -0.9300  0.1948
2 NKE   -0.7015  0.6699
3 PH    -0.9309 -0.1987
4 EH    -0.8370 -0.4069
5 TKW   -0.8881 -0.1415
```

```
-------------------------- Loadings after varimax rotation ---------------------------
# A tibble: 5 x 3
  VAR       FA1    FA2
  <chr>   <dbl>  <dbl>
1 KW    -0.6590 0.6844
2 NKE   -0.2028 0.9486
3 PH    -0.8813 0.3597
4 EH    -0.9208 0.1348
5 TKW   -0.8137 0.3829

--------------------------- Scores for genotypes-ideotype -----------------------------
# A tibble: 14 x 3
   GEN         FA1     FA2
   <chr>     <dbl>   <dbl>
 1 H1    -3.164     1.022
 2 H10   -0.5663    1.839
 3 H11   -1.553     1.671
 4 H12   -0.8617    1.121
 5 H13   -2.242     2.014
 6 H2    -1.866     2.331
 7 H3    -2.539     0.2883
 8 H4    -1.958     3.177
 9 H5    -2.663     2.727
10 H6    -3.482     0.7752
11 H7    -2.138     0.9181
12 H8     0.008421  0.9992
13 H9    -1.561    -0.4268
14 ID1   -3.768     2.946

---------------------------- Multitrait stability index ------------------------------
      H5      H13       H4       H2       H1       H6      H11       H7
1.126808 1.787992 1.824705 1.999554 2.017441 2.190083 2.556014 2.602273
      H3      H10      H12       H9       H8
2.928670 3.387928 3.432086 4.031264 4.248973

--------------------------- Selection differential (index) ----------------------------
# A tibble: 5 x 6
  VAR   Factor    Xo    Xs    SD SDperc
  <chr> <chr>  <dbl> <dbl> <dbl>  <dbl>
1 PH    FA 1   51.53 76.53 25.00  48.53
2 EH    FA 1   44.24 59.08 14.84  33.54
3 TKW   FA 1   53.78 70.89 17.11  31.82
4 KW    FA 2   53.25 75.92 22.67  42.56
5 NKE   FA 2   44.98 71.68 26.70  59.36

-------------------------- Mean of Selection differential -----------------------------
      Xo       Xs       SD   SDperc
49.55600 70.81921 21.26321 43.16009
```

```
------------------------- Selection differential (variables) --------------------------
# A tibble: 5 x 6
  VAR   Factor      xo      Xs        SD SDperc
  <chr> <chr>    <dbl>   <dbl>     <dbl>  <dbl>
1 PH    FA 1     2.485   2.553   0.06852 2.758
2 EH    FA 1     1.343   1.362   0.01901 1.415
3 TKW   FA 1   338.7   340.6     1.912   0.5647
4 KW    FA 2   172.9   181.7     8.798   5.087
5 NKE   FA 2   511.6   539.7    28.03    5.479

-------------------------------- Selected genotypes -----------------------------------
H5 H13
```

```
plot(index)
```

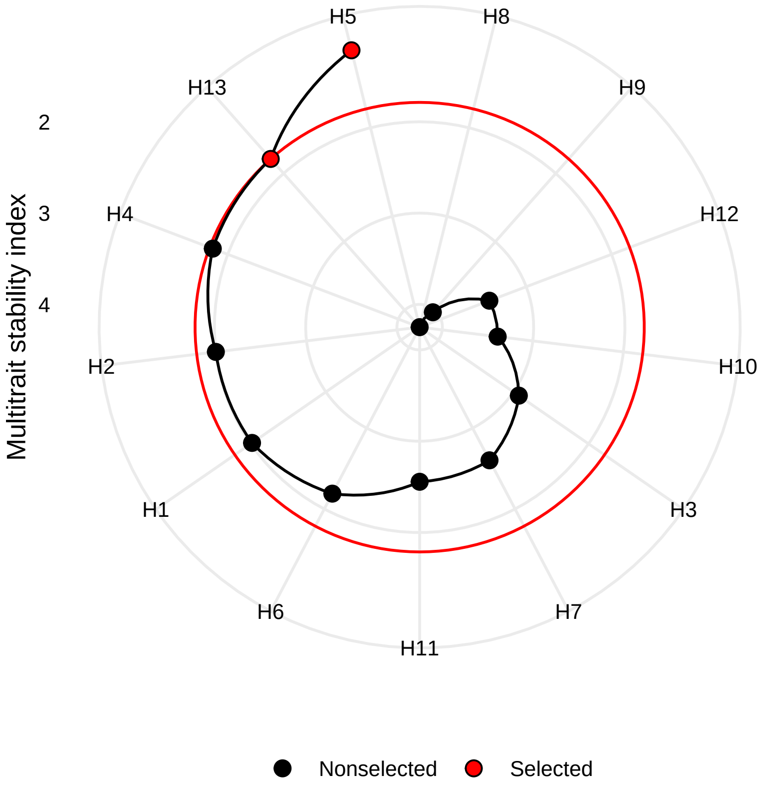

#### 8.10 Wrapper function for stability indexes

The easiest way to compute the stability indexes in `metan` is by using the function `ge_stats()`. It is a wrapper function that computes several stability indexes at once. We can then use `get_model_data()` to get the stability statistics or the genotype's ranks for each statistic.

```r
stab_ind <- ge_stats(data_ge2, ENV, GEN, REP, resp = c(EH, EP, EL, NKE))
```

```
Evaluating variable EH 0 %
Evaluating variable EP 33.3 %
Evaluating variable EL 66.7 %
Evaluating variable NKE 100 %
```

```r
get_model_data(stab_ind, "stats")
```

```
# A tibble: 52 x 33
   var   gen       Y    CV   Var  Shukla  Wi_g  Wi_f  Wi_u Ecoval   bij     Sij
   <chr> <chr> <dbl> <dbl> <dbl>   <dbl> <dbl> <dbl> <dbl>  <dbl> <dbl>   <dbl>
 1 EH    H1     1.50  20.6 0.288 0.0522   84.6 100.   78.3  0.425 1.06  0.0637
 2 EH    H10    1.26  24.8 0.294 0.0303   72.4 101.   62.8  0.258 1.30  0.0303
 3 EH    H11    1.27  20.1 0.195 0.0220   75.6  95.0  61.2  0.195 1.00  0.0256
 4 EH    H12    1.28  21.6 0.231 0.0783   61.3  96.4  33.8  0.624 0.590 0.0861
 5 EH    H13    1.35  14.5 0.116 0.0345   76.5  99.8  59.6  0.290 0.575 0.0298
 6 EH    H2     1.38  20.8 0.246 0.0906   69.4  76.2  73.4  0.718 0.525 0.0981
 7 EH    H3     1.41  24.5 0.359 0.0715   72.2  93.4  58.7  0.572 1.15  0.0870
 8 EH    H4     1.43  23.5 0.337 0.0366   80.7 102.   63.3  0.306 1.41  0.0335
 9 EH    H5     1.37  20.7 0.241 0.00986  86.8  96.6  76.3  0.103 1.30  0.00449
10 EH    H6     1.41  13.8 0.114 0.0126   91.1  97.5  99.4  0.124 0.780 0.0106
# ... with 42 more rows, and 21 more variables: R2 <dbl>, ASV <dbl>,
#   SIPC <dbl>, EV <dbl>, ZA <dbl>, WAAS <dbl>, HMGV <dbl>, RPGV <dbl>,
#   HMRPGV <dbl>, Pi_a <dbl>, Pi_f <dbl>, Pi_u <dbl>, Gai <dbl>, S1 <dbl>,
#   S2 <dbl>, S3 <dbl>, S6 <dbl>, N1 <dbl>, N2 <dbl>, N3 <dbl>, N4 <dbl>
```

```r
get_model_data(stab_ind, "ranks")
```

```
# A tibble: 52 x 32
   var   gen     Y_R  CV_R Var_R Shukla_R Wi_g_R Wi_f_R Wi_u_R Ecoval_R Sij_R
   <chr> <chr> <dbl> <dbl> <dbl>    <dbl>  <dbl>  <dbl>  <dbl>    <dbl> <dbl>
 1 EH    H1        1     5     9        9      3      4      2        9     9
 2 EH    H10      12    13    10        6      9      2      8        6     7
 3 EH    H11      11     4     4        5      8      9      9        5     5
 4 EH    H12       9     8     5       12     12      8     13       12    11
 5 EH    H13       7     2     2        7      7      5     10        7     6
 6 EH    H2        5     7     7       13     11     13      5       13    13
 7 EH    H3        4    11    13       11     10     10     11       11    12
 8 EH    H4        2     9    12        8      5      1      7        8     8
 9 EH    H5        6     6     6        1      2      7      4        1     2
10 EH    H6        3     1     1        3      1      6      1        3     3
# ... with 42 more rows, and 21 more variables: R2_R <dbl>, ASV_R <dbl>,
#   SIPC_R <dbl>, EV_R <dbl>, ZA_R <dbl>, WAAS_R <dbl>, HMGV_R <dbl>,
#   RPGV_R <dbl>, HMRPGV_R <dbl>, Pi_a_R <dbl>, Pi_f_R <dbl>, Pi_u_R <dbl>,
```

```
#   Gai_R <dbl>, S1_R <dbl>, S2_R <dbl>, S3_R <dbl>, S6_R <dbl>, N1_R <dbl>,
#   N2_R <dbl>, N3_R <dbl>, N4_R <dbl>
```

In addition, by using the function `corr_stab_ind()` it is possible to compute a Spearman's rank correlation matrix between the computed stability indexes. By default, all statistics are included. It is possible to include only parametric statistics, using `stats = "par"`, nonparametric statistics using `stats = "nonpar"`, AMMI-based stability indexes using `stats = "ammi"`. To include specific statistics, use a character vector with the statis=tcs names.

```r
a <- corr_stab_ind(stab_ind,
                   plot = FALSE,
                   stats = c("HMRPGV, RPGV, ASV, Wi_g, Sij, Ecoval, N2"))
b <- corr_stab_ind(stab_ind,
                   plot = FALSE,
                   stats = "nonpar")
arrange_ggplot(a$plot, b$plot, labels = letters[1:2])
```

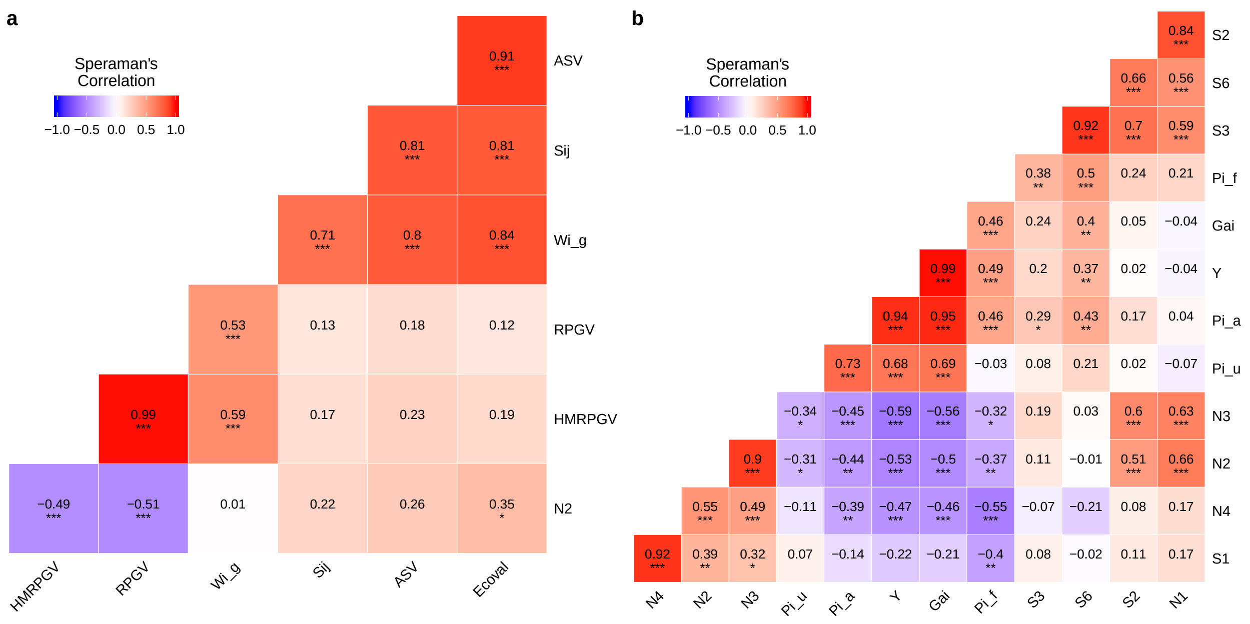

### 9 Biometrical models

#### 9.1 Correlation coefficient with p-values

Pearson's correlation coefficient can be easily computed with the function `corr_coef()`. Users can use a data frame that may contain factor variables; only numeric variables will be used. Indeed, users can print the results with `print()` or create correlation heat map can be created using `plot()`.

```r
coef_all <- corr_coef(data_ge2)
a <- plot(coef_all)
```

```r
coef_sel <- corr_coef(data_ge2, PH, EH, CD, CW, PERK)
print(coef_sel)
```

```
---------------------------------------------------------------------------
Pearson's correlation coefficient
---------------------------------------------------------------------------
         PH      EH      CD     CW    PERK
PH   1.0000  0.9318  0.3154  0.505  0.0408
EH   0.9318  1.0000  0.2805  0.519 -0.0213
CD   0.3154  0.2805  1.0000  0.484 -0.0482
CW   0.5047  0.5193  0.4840  1.000 -0.6811
PERK 0.0408 -0.0213 -0.0482 -0.681  1.0000
---------------------------------------------------------------------------
p-values for the correlation coefficients
---------------------------------------------------------------------------
           PH       EH       CD       CW     PERK
PH   0.00e+00 1.11e-69 6.06e-05 1.83e-11 6.13e-01
EH   1.11e-69 0.00e+00 3.90e-04 3.76e-12 7.91e-01
CD   6.06e-05 3.90e-04 0.00e+00 1.54e-10 5.50e-01
CW   1.83e-11 3.76e-12 1.54e-10 0.00e+00 1.34e-22
PERK 6.13e-01 7.91e-01 5.50e-01 1.34e-22 0.00e+00
```

```r
b <- plot(coef_sel)
c <- plot(coef_sel,
          diag = TRUE,
          type = "upper",
          col.high = "darkgreen",
          col.low = "darkblue")
arrange_ggplot(a, b, c, ncol = 3, labels = letters[1:3])
```

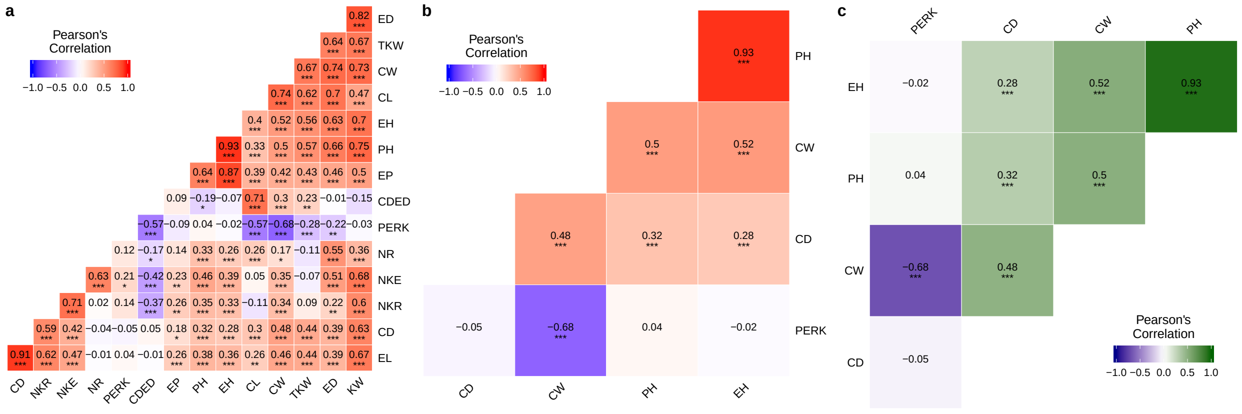

#### 9.2 Graphical and numerical visualization of a correlation matrix

The function `corr_plot()` can be used to visualize (both graphically and numerically) a correlation matrix. Pairwise of scatterplots are produced and may be shown in the upper or lower diagonal, which may be seen as a nicer and customizable ggplot2-based version of the `pairs()` base R function.

By calling `corr_plot(data)` where `data` is the data set, all numeric variables in `data` are plotted. The selection of variables to plot can be made by only using a comma-separated list of unquoted variable names, as follows.

```
a <- corr_plot(data_ge2, CD, EL, PERK)
b <- corr_plot(data_ge2, CD, EL, PERK,
               lower = NULL,
               upper = "corr")
c <- corr_plot(data_ge2, CD, EL, PERK,
               shape.point = 19,
               size.point = 2,
               size.axis.label = 7,
               alpha.point = 0.5,
               alpha.diag = 0,
               pan.spacing = 0,
               diag.type = "boxplot",
               col.sign = "gray",
               alpha.sign = 0.3,
               axis.labels = TRUE)
d <- corr_plot(data_ge2, CD, EL, PERK,
               lower = "corr", # Define lower pannel
               upper = "scatter", # Define upper pannel
               prob = 0.01, # Significance value
               shape.point = 21, # Shape of the point
               col.point = "black", # Color of the point
               fill.point = "orange", # Color to fill points
               size.point = 2, # Size of the point
               alpha.point = 0.6, # Transparency of the color
               maxsize = 4, # Size of the maximum correlation
               minsize = 1, # Size of the minimum correlation
               smooth = TRUE, # Linear smooth line
               size.smooth = 1, # Size of the smooth line
               col.smooth = "red", # Color of the smooth line
               col.sign = "cyan", # Color for significant pairs
               col.up.panel = "black", # Color for the upper pannel
               col.lw.panel = "black", # Color for the lower pannel
               col.dia.panel = "black", # Color for the diagonal pannel
               diag.type = "density", # What diagonal shows?
               pan.spacing = 0, # Space between pannels
               lab.position = "tl") # Position of the labels
arrange_ggplot(a, b, c, d, ncol = 4, labels = letters[1:4])
```

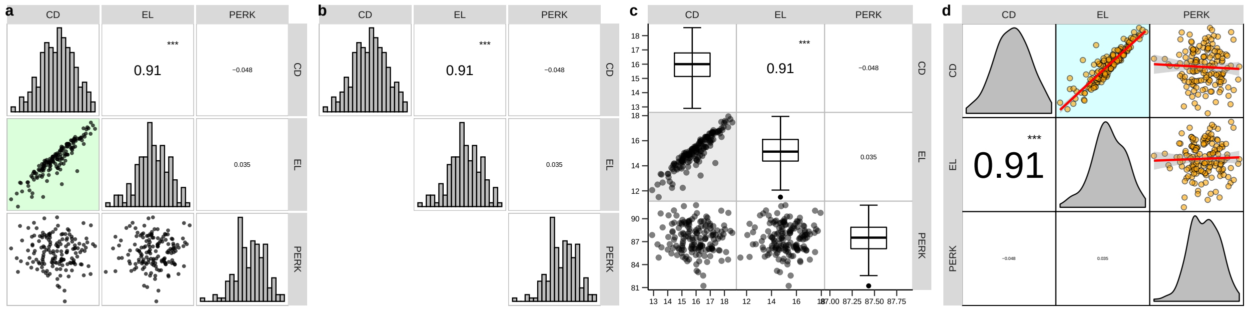

#### 9.3 Nonparametric confidence interval for Pearson's correlation

The function `corr_ci()` can be used to estimate the confidence interval for Pearson's correlation coefficient using a Gaussian-independent estimator (Olivoto et al., 2018). It is possible to estimate the confidence interval by declaring the sample size (n) and the correlation coefficient (r) or using a data frame. The following code computes the confidence interval and makes a plot to show the results.

```
library(ggplot2)
a <- data_ge2 %>%
  corr_ci(verbose = FALSE) %>%
  plot_ci() +
  theme(axis.text.y = element_blank())

ci <- data_ge2 %>%
  corr_ci(EH, PERK, EP, EL, ED, verbose = FALSE)

b <- plot_ci(ci)
c <- plot_ci(ci,
             fill.shape = "gray",
             size.shape = 4,
             main = FALSE,
             width.errbar = 0.2)
arrange_ggplot(a, b, c, ncol = 3, labels = letters[1:3])
```

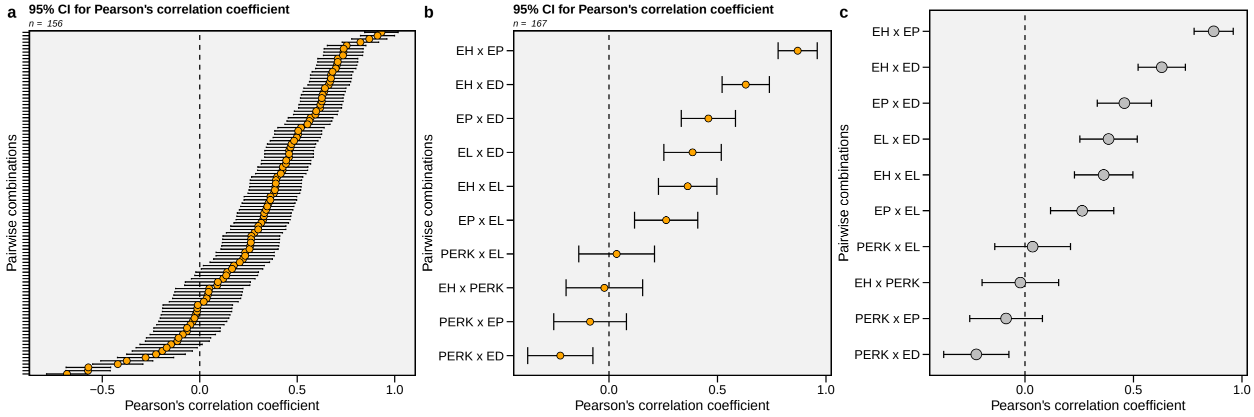

In the following examples, the confidence interval is calculated by declaring the sample size (n) and the correlation coefficient (r). If `by` is used, then the confidence interval will be calculated within each level of the grouping variable, in this case, environment (ENV).

```
# Inform n and r
corr_ci(n = 145, r = 0.34)
```

```
--------------------------------------------------
Nonparametric 95% half-width confidence interval
--------------------------------------------------
Level of significance: 5%
Correlation coefficient: 0.34
Sample size: 145
Confidence interval: 0.1422
True parameter range from: 0.1978 to 0.4822
--------------------------------------------------
```

```
# Compute the confidence for each level of ENV
corr_ci(data_ge2,
        EH, PERK, EL,
        by = ENV)
```

```
# A tibble: 12 x 6
   LEVEL Pair         Corr    CI      LL    UL
   <chr> <chr>       <dbl> <dbl>   <dbl> <dbl>
 1 A1    EH x EL    0.148  0.320 -0.172  0.467
 2 A1    EH x PERK -0.205  0.306 -0.510  0.101
 3 A1    PERK x EL -0.0270 0.352 -0.379  0.325
 4 A2    EH x EL    0.498  0.242  0.255  0.740
 5 A2    EH x PERK  0.0187 0.354 -0.335  0.373
 6 A2    PERK x EL  0.0498 0.345 -0.296  0.395
 7 A3    EH x EL   -0.0977 0.333 -0.430  0.235
 8 A3    EH x PERK  0.126  0.325 -0.199  0.451
 9 A3    PERK x EL  0.0391 0.348 -0.309  0.388
10 A4    EH x EL    0.250  0.295 -0.0446 0.545
11 A4    EH x PERK  0.250  0.295 -0.0447 0.545
12 A4    PERK x EL  0.182  0.311 -0.129  0.493
```

#### 9.4 Sample size planning

The function `corr_ss()` can be used to plan the required sample size to obtain a pre-established (if the correlation coefficient is previously known). Olivoto et al. (2018) suggests computing the required sample size considering a null correlation ($r = 0$) since any correlation greater than 0 will have a small confidence interval. Here, we will plan the sample size for a correlation study to achieve a half-width confidence interval of 0.15.

```r
corr_ss(r = 0, CI = 0.15)
```

```
-------------------------------------------------
Sample size planning for correlation coefficient
-------------------------------------------------
Level of significance: 5%
Correlation coefficient: 0
95% half-width CI: 0.15
Required sample size: 223
-------------------------------------------------
```

#### 9.5 Partial correlation coefficient

Pearson's linear correlation does not consider the influence of a set of traits on the relationship between two traits. For example, the hypothetical correlation of $r = 0.9$ between $x$ and $y$ may be due to the influence of a third trait or group of traits acting together. To identify this linear effect between $x$ and $y$ controlling statistically the effect of others traits, the partial correlation is used. From Pearson's simple correlation matrix, the partial correlation is calculated as follows

$$r_{xy.m} = \frac{-a_{xy}}{\sqrt{a_{xx}a_{yy}}}$$

Where $r_{xy.m}$ is the partial correlation coefficient between the traits $x$ and $y$, excluding the effects of the $m$ remaining traits of the set; $-a_{ij}$ is the inverse element of the correlation matrix corresponding to xy, $a_{ii}a_{jj}$ are the diagonal elements of the inverse matrix of correlation associated with trait $x$ and $y$ , respectively. Both the linear and partial correlation coefficients may be obtained using the function `lpcor()`.

```r
lpcor(data_ge2, NR, NKR, NKE)
```

```
        Pairs      linear      partial            t         prob
1     PH x EH  0.93182821  0.985278587  68.43576425 0.000000e+00
2     PH x EP  0.63841233 -0.950971899 -36.51145994 0.000000e+00
3     PH x EL  0.38019601 -0.058285302  -0.69327821 4.892754e-01
4     PH x ED  0.66131483  0.140319976   1.68285723 9.461577e-02
5     PH x CL  0.32516481 -0.156567454  -1.88235003 6.184913e-02
6     PH x CD  0.31539095 -0.007850816  -0.09322614 9.258562e-01
7     PH x CW  0.50473880 -0.047792236  -0.56815059 5.708364e-01
8     PH x KW  0.75344390  0.114119292   1.36400247 1.747402e-01
9     PH x NR  0.32860646  0.215504295   2.62054700 9.741135e-03
10   PH x NKR  0.35304952  0.175952721   2.12243573 3.554887e-02
11  PH x CDED -0.19201795  0.142114402   1.70481853 9.043063e-02
12  PH x PERK  0.04081442 -0.063254136  -0.75260838 4.529399e-01
13   PH x TKW  0.56853845 -0.007890020  -0.09369172 9.254871e-01
14   PH x NKE  0.45838005 -0.109640743  -1.30980816 1.923908e-01
15    EH x EP  0.86954597  0.981725615  61.25711425 0.000000e+00
16    EH x EL  0.36265373  0.048186955   0.57285385 5.676560e-01
```

```
17      EH x ED  0.63025605 -0.136239106  -1.63297564 1.047053e-01
18      EH x CL  0.39719352  0.149260358   1.79244768 7.520541e-02
19      EH x CD  0.28051181  0.006948814   0.08251459 9.343545e-01
20      EH x CW  0.51931358  0.041167817   0.48925551 6.254210e-01
21      EH x KW  0.70294690 -0.104088880  -1.24273752 2.160273e-01
22      EH x NR  0.26480509 -0.205183182  -2.48938037 1.395912e-02
23     EH x NKR  0.33105286 -0.197976463  -2.39831055 1.777916e-02
24    EH x CDED -0.06590990 -0.138009161  -1.65460094 1.002298e-01
25    EH x PERK -0.02134916  0.056543205   0.67248924 5.023728e-01
26     EH x TKW  0.56235676  0.038035919   0.45197858 6.519789e-01
27     EH x NKE  0.38812143  0.129778940   1.55418333 1.223824e-01
28      EP x EL  0.26342374 -0.016085023  -0.19102378 8.487817e-01
29      EP x ED  0.45801958  0.114258593   1.36568947 1.742110e-01
30      EP x CL  0.39082392 -0.120729960  -1.44415226 1.509145e-01
31      EP x CD  0.17504485 -0.054356204  -0.64639979 5.190712e-01
32      EP x CW  0.42480976 -0.044732469  -0.53170088 5.957699e-01
33      EP x KW  0.49741927  0.117760756   1.40812927 1.612938e-01
34      EP x NR  0.14043147  0.197748139   2.39543197 1.791372e-02
35     EP x NKR  0.25883185  0.242820641   2.97229250 3.476892e-03
36    EP x CDED  0.08966393  0.114678017   1.37076936 1.726250e-01
37    EP x PERK -0.08708974 -0.054178652  -0.64428213 5.204392e-01
38     EP x TKW  0.42631086 -0.064089374  -0.76258690 4.469837e-01
39     EP x NKE  0.23305230 -0.157477037  -1.89356286 6.033174e-02
40      EL x ED  0.38514512 -0.023996179  -0.28502091 7.760463e-01
41      EL x CL  0.25540676 -0.002205465  -0.02618852 9.791440e-01
42      EL x CD  0.91186526  0.795439402  15.58548377 0.000000e+00
43      EL x CW  0.45817278 -0.052188243  -0.62054669 5.358997e-01
44      EL x KW  0.66856012  0.098388161   1.17399075 2.423778e-01
45      EL x NR -0.01387378 -0.090008477  -1.07314736 2.850382e-01
46     EL x NKR  0.61715434 -0.091880285  -1.09565247 2.750988e-01
47    EL x CDED -0.01257977  0.005496759   0.06527139 9.480504e-01
48    EL x PERK  0.03526159 -0.021277238  -0.25271041 8.008600e-01
49     EL x TKW  0.44210108  0.039604034   0.47064108 6.386243e-01
50     EL x NKE  0.46569935  0.086961050   1.03653194 3.017284e-01
51      ED x CL  0.69746290  0.988412265  77.32059491 0.000000e+00
52      ED x CD  0.38971282  0.054405882   0.64699231 5.186888e-01
53      ED x CW  0.73713052 -0.526499799  -7.35358822 1.439671e-11
54      ED x KW  0.82414264  0.323910615   4.06540059 7.935336e-05
55      ED x NR  0.55253448  0.128630330   1.54019554 1.257539e-01
56     ED x NKR  0.22207274 -0.046635102  -0.55436431 5.802078e-01
57    ED x CDED -0.01003537 -0.988443687 -77.42749093 0.000000e+00
58    ED x PERK -0.22439531 -0.505305834  -6.95317408 1.224172e-10
59     ED x TKW  0.64198696  0.121324449   1.45136941 1.488983e-01
60     ED x NKE  0.50508084  0.102249238   1.22053950 2.242977e-01
61      CL x CD  0.30036364 -0.026905492  -0.31960072 7.497441e-01
62      CL x CW  0.73833793  0.539275577   7.60398938 3.681500e-12
63      CL x KW  0.47093101 -0.315422733  -3.94692287 1.244293e-04
64      CL x NR  0.26193592 -0.063667285  -0.75754405 4.499881e-01
```

```
65      CL x NKR -0.11494054  0.004778778   0.05674550 9.548283e-01
66     CL x CDED  0.70799925  0.996334355 138.30019683 0.000000e+00
67     CL x PERK -0.57313356  0.500461334   6.86409249 1.956586e-10
68      CL x TKW  0.61870013 -0.085603234  -1.02022702 3.093681e-01
69      CL x NKE  0.04894224 -0.084714288  -1.00955550 3.144376e-01
70       CD x CW  0.48402989  0.001424136   0.01691070 9.865318e-01
71       CD x KW  0.62598062 -0.058951531  -0.70123020 4.843151e-01
72       CD x NR -0.03584984 -0.036510033  -0.43382185 6.650809e-01
73      CD x NKR  0.59332061  0.232418570   2.83752067 5.217558e-03
74     CD x CDED  0.04530711  0.042012094   0.49930682 6.183414e-01
75     CD x PERK -0.04820386  0.006219481   0.07385367 9.412315e-01
76      CD x TKW  0.44332490  0.047540319   0.56514901 5.728705e-01
77      CD x NKE  0.41561684  0.006521650   0.07744195 9.383818e-01
78       CW x KW  0.73486220  0.591378621   8.70818957 7.549517e-15
79       CW x NR  0.16565752 -0.044476554  -0.52865296 5.978774e-01
80      CW x NKR  0.34031548  0.038954521   0.46291066 6.441421e-01
81     CW x CDED  0.29986077 -0.522308966  -7.27297493 2.224310e-11
82     CW x PERK -0.68106736 -0.963234041 -42.57281727 0.000000e+00
83      CW x TKW  0.67345853  0.095758548   1.14231918 2.552574e-01
84      CW x NKE  0.34627743  0.109231019   1.30485423 1.940681e-01
85       KW x NR  0.36214470 -0.064547929  -0.76806591 4.437324e-01
86      KW x NKR  0.59737013  0.035584983   0.42281605 6.730736e-01
87     KW x CDED -0.14702901  0.311050756   3.88631010 1.560615e-04
88     KW x PERK -0.02683251  0.612152054   9.19253133 4.440892e-16
89      KW x TKW  0.67303715  0.649507403  10.14324936 0.000000e+00
90      KW x NKE  0.68107556  0.599697532   8.89874468 2.442491e-15
91      NR x NKR  0.02055097 -0.467031188  -6.27169411 4.105635e-09
92     NR x CDED -0.16965700  0.085315683   1.01677480 3.110021e-01
93     NR x PERK  0.12054109  0.021530021   0.25571411 7.985443e-01
94      NR x TKW -0.10876267 -0.031483195  -0.37402764 7.089456e-01
95      NR x NKE  0.62608544  0.370813890   4.74118321 5.140262e-06
96    NKR x CDED -0.37442456 -0.023822635  -0.28295842 7.776237e-01
97    NKR x PERK  0.13554094 -0.006181769  -0.07340584 9.415872e-01
98     NKR x TKW  0.09286139  0.033169489   0.39408271 6.941153e-01
99     NKR x NKE  0.70783318  0.352573751   4.47387559 1.569665e-05
100 CDED x PERK -0.57138058 -0.496524946  -6.79234396 2.848701e-10
101  CDED x TKW  0.23282587  0.086622859   1.03247034 3.036195e-01
102  CDED x NKE -0.42051242  0.075770195   0.90231509 3.684283e-01
103  PERK x TKW -0.27789427  0.065871985   0.78388903 4.344201e-01
104  PERK x NKE  0.20528475  0.072594616   0.86429371 3.888945e-01
105   TKW x NKE -0.06515986 -0.869921187 -20.94471865 0.000000e+00
```

```r
# Compute the correlations for each level of the factor ENV
lpc2 <- lpcor(data_ge2,
              NR, NKR, NKE,
              by = ENV,
              verbose = FALSE)
print(lpc2$summary)
```

```
# A tibble: 12 x 6
   ENV   Pairs      linear partial     t     prob
   <chr> <chr>       <dbl>   <dbl> <dbl>    <dbl>
 1 A1    NR x NKR  -0.207   -0.856 -9.94 7.25e-12
 2 A1    NR x NKE   0.399    0.756  6.92 4.18e- 8
 3 A1    NKR x NKE  0.678    0.814  8.41 5.06e-10
 4 A2    NR x NKE   0.727    0.932 15.4  0.
 5 A2    NKR x NKE  0.675    0.850  9.67 1.49e-11
 6 A2    NR x NKR   0.0363  -0.822 -8.65 2.59e-10
 7 A3    NKR x NKE  0.442    0.881 11.2  3.12e-13
 8 A3    NR x NKR   0.127   -0.622 -4.76 3.09e- 5
 9 A3    NR x NKE   0.579    0.885 11.4  1.60e-13
10 A4    NR x NKR  -0.215   -0.716 -6.16 4.24e- 7
11 A4    NR x NKE   0.681    0.816  8.47 4.30e-10
12 A4    NKR x NKE  0.783    0.935 15.8  0.
```

#### 9.6 (co)variance and correlations for designed experiments

The function `covcor_design()` can be used to compute genetic, phenotypic and residual correlation/(co)variance matrices through Analysis of Variance (ANOVA) method using randomized complete block design (RCBD) or completely randomized design (CRD).

The phenotypic ($r^p_{xy}$), genotypic ($r^g_{xy}$) and residual ($r^r_{xy}$) correlations between $x$ and $y$ are computed as follows:

$$r^p_{xy} = \frac{cov^p_{xy}}{\sqrt{var^p_x var^p_y}} \qquad r^g_{xy} = \frac{cov^g_{xy}}{\sqrt{var^g_x var^g_y}} \qquad r^r_{xy} = \frac{cov^r_{xy}}{\sqrt{var^r_x var^r_y}}$$

Using Mean Squares from the ANOVA method, the variances ($var$) and covariances ($cov$) are computed as follows:

$$cov^p_{xy} = [(MST_{x+y} - MST_x - MST_y)/2]/r$$
$$var^p_x = MST_x/r$$
$$var^p_y = MST_y/r$$
$$cov^g_{xy} = [(cov^p_{xy} \times r) - cov^r_{xy}]/r$$
$$var^g_x = (MST_x - MSE_x)/r$$
$$var^g_y = (MST_x - MSE_y)/r$$
$$cov^r_{xy} = (MSR_{x+y} - MSR_x - MSR_y)/2$$
$$var^r_x = MSR_x$$
$$var^r_y = MSR_y$$

where $MST$ is the mean square for treatment, $MSR$ is the mean square for residuals, and $r$ is the number of replications.

##### 9.6.1 Genetic correlations

```
data = subset(data_ge2, ENV == "A1")
covcor_design(data, GEN, REP,
              resp = c(PH, EH, NKE, TKW),
              type = "gcor")
```

```
              PH           EH        NKE        TKW
PH   1.000000000 -0.006544623  0.2801806  0.2459377
EH  -0.006544623  1.000000000 -0.7752497  0.7247684
NKE  0.280180560 -0.775249741  1.0000000 -0.5117645
TKW  0.245937657  0.724768430 -0.5117645  1.0000000
```

##### 9.6.2 Phenotypic correlations

```
covcor_design(data, GEN, REP,
              resp = c(PH, EH, NKE, TKW),
              type = "pcor")
```

```
              PH         EH        NKE          TKW
PH  1.0000000000  0.3307336  0.1417114  0.0008856916
EH  0.3307336444  1.0000000 -0.4388300  0.3828624624
NKE 0.1417113819 -0.4388300  1.0000000 -0.5522625652
TKW 0.0008856916  0.3828625 -0.5522626  1.0000000000
```

##### 9.6.3 Residual correlations

```
covcor_design(data, GEN, REP,
              resp = c(PH, EH, NKE, TKW),
              type = "rcor")
```

```
             PH          EH         NKE         TKW
PH   1.00000000  0.53436776  0.09568529 -0.15533646
EH   0.53436776  1.00000000 -0.23177906 -0.04084134
NKE  0.09568529 -0.23177906  1.00000000 -0.63443188
TKW -0.15533646 -0.04084134 -0.63443188  1.00000000
```

##### 9.6.4 Residual (co)variance matrix

```
rcov <- covcor_design(data, GEN, REP,
                      resp = c(PH, EH, NKE, TKW),
                      type = "rcov")
rcov
```

```
              PH           EH          NKE           TKW
PH   0.014566384  0.007240156    0.6990162    -0.5971700
EH   0.007240156  0.012602700   -1.5749695    -0.1460429
NKE  0.699016154 -1.574969487  3663.8047009 -1223.2067115
TKW -0.597169989 -0.146042893 -1223.2067115  1014.6059358
```

#### 9.7 Path Analysis

##### 9.7.1 The model

Path analysis (Wright, 1923) is –within certain limitations– a method of assessing the logical consequences of a causal relationship hypothesis in a system of correlated traits. The statistical method is consolidated and used worldwide in several areas of science.

The decomposition of linear correlations into direct and indirect effects of a set of explanatory variables is based on the system of normal equations.

$$X'X\hat{\beta} = X'Y$$

whose resolution is

$$\hat{\beta} = X'X^{-1}X'Y$$

where $\hat{\beta}$ is the partial regression coefficient vector ($\hat{\beta}_1$, $\hat{\beta}_2$, $\hat{\beta}_3$, ..., $\hat{\beta}_p$) with $p + 1$; $X'X^{-1}$ is the inverse of the linear correlation matrix between the explanatory variables and $X'Y$ is the correlation matrix of each explanatory variable, with the dependent variable.

After estimating regression coefficients ($\hat{\beta}_p$), the direct and indirect effects of the set of explanatory p-variables can be estimated. Consider the following example, where a set of explanatory variables ($a$, $b$, $c$) are used to explain cause and effect relationships in the response of a dependent variable (say, $y$). After estimates of the partial regression coefficients ($\hat{\beta}_1$, $\hat{\beta}_2$ and $\hat{\beta}_3$), the direct and indirect effects of $a$ on $y$ are given by:

$$r_{a:y} = \hat{\beta}_1 + \hat{\beta}_{2_{ra:b}} + \hat{\beta}_{3_{ra:c}}$$

where $r_{a:y}$ is the linear correlation between $a$ and $y$, $\hat{\beta}_1$ is the direct effect of $a$ on $y$; $\hat{\beta}_{2_{ra:b}}$ is the indirect effect of $a$ on $y$ via $b$ and $\hat{\beta}_{3_{ra:c}}$ is the indirect effect of $a$ on $y$ via $c$. Similar regressions are used to estimate the effects of $b$ and $c$ as follows:

$$r_{b:y} = \hat{\beta}_{1_{rb:a}} + \hat{\beta}_2 + \hat{\beta}_{3_{rb:c}}$$
$$r_{c:y} = \hat{\beta}_{1_{rc:a}} + \hat{\beta}_{2_{rc:b}} + \hat{\beta}_3$$

Although path analysis reveals cause and effect associations, its estimation is based on multiple regression principles. Thus, parameter estimates may be biased due to the complex nature of the data, where the response of dependent traits is linked to a large number of explanatory variables, which are often correlated or multicollinear to each other (Graham, 2003). Thus, whenever two explanatory variables are highly associated, it is difficult to estimate the relationships of each explanatory variable individually, since several parameters solve the system of normal equations. This particularity is called multicollinearity (Blalock, 1963).

##### 9.7.2 Collinearity diagnosis

The main ways used to identify the degree of multicollinearity in an array of explanatory variables are as follows (T. Olivoto, Souza, et al., 2017).

- **Condition number, CN**, calculated by the ratio of the highest and lowest eigenvalues ($\lambda$) of the $X'X$ correlation matrix, according to the expression

$$\text{NC} = \frac{\lambda_{\text{Max}}}{\lambda_{\text{Min}}}$$

- **Matrix determinant, D** estimated by the product of eigenvalues of $X'X$, for $\lambda_j > 0$, according to the expression

$$D_{\boldsymbol{X'X}} = \prod_{j=1}^{p} \lambda j$$

- **Variance Inflation Factor, VIF**, that measure how much the variance of the estimated regression coefficients ($\hat{\beta}_p$) was inflated compared to when explanatory traits are not linearly associated. The estimation of VIF for the $k$th element of the $\hat{\beta}$ is given by the sum of the quotients of each squared component of the eigenvector divided by its respective associated eigenvalue:

$$\text{VIF}_{\beta_k} = \left( \frac{(\text{EV}_{k_{C1}})^2}{\lambda_1} + \frac{(\text{EV}_{k_{C2}})^2}{\lambda_2} + \ldots + \frac{(\text{EV}_{k_{Cp}})^2}{\lambda_p} \right)$$

where $VIF_{\beta_k}$ is the variance inflation factor the $k$th element of $\beta$ ($k = 1, 2, ..., p$); $EV_{k_{C1}}$ is the component of the $k$th eigenvector; and ($C = 1, 2, ..., p$); and $\lambda$ is the eigenvalue associated with the respective eigenvector ($\lambda = 1, 2, ..., p$). VIFs can also be considered as the diagonal elements of $X'X^{-1}$. The presence of VIFs greater than 10 is considered to be indicative of multicollinearity.

The function `colindiag()` computes a collinearity diagnostic of a correlation matrix of predictor traits. Several indicators, such as Variance Inflation Factor, Condition Number, and Matrix Determinant are used (Tiago Olivoto et al., 2017; T. Olivoto, Souza, et al., 2017). If only the data is informed in the function, all the numeric variables will be considered in the diagnostic. Here, we will check the colinearity in the correlation matrix between PH, EP, EH, and CD from `data_ge2`.

```
col_diag <- colindiag(data_ge2, PH, EP, EH, CD)
```

```
Severe multicollinearity in the matrix! Pay attention on the variables listed bellow
CN = 1020.931
Matrix determinant: 0.0023242
Largest correlation: PH x EH = 0.932
Smallest correlation: EP x CD = 0.175
```

```
Number of VIFs > 10: 3
Number of correlations with r >= |0.8|: 2
Variables with largest weight in the last eigenvalues:
EH > PH > EP > CD
```

```
# Diagnostic for each environment
# All numeric variables minus variables that contains 'E'
col_diag <- colindiag(data_ge2, -contains("E"), by = ENV)
```

```
-----------------------------------------------------------------------------
Level: A1
-----------------------------------------------------------------------------
Weak multicollinearity in the matrix
NC = 72.59
Matrix determinant: 0.0076727
Largest correlation: CL x CW = 0.662
Smallest correlation: CL x NR = -0.006
Number of VIFs > 10: 0
Number of correlations with r >= |0.8|: 0
Variables with largest weight in the last eigenvalues:
TKW > KW > NR > NKR > CW > CL > PH > CD

-----------------------------------------------------------------------------
Level: A2
-----------------------------------------------------------------------------
The multicollinearity in the matrix should be investigated.
NC = 188.535
Largest VIF = 23.8677009257904
Matrix determinant: 0.0001001
Largest correlation: KW x TKW = 0.83
Smallest correlation: CD x NR = 0.064
Number of VIFs > 10: 2
Number of correlations with r >= |0.8|: 3
Variables with largest weight in the last eigenvalues:
KW > TKW > NR > NKR > CL > CD > CW > PH

-----------------------------------------------------------------------------
Level: A3
-----------------------------------------------------------------------------
Weak multicollinearity in the matrix
NC = 52.096
Matrix determinant: 0.002208
Largest correlation: CL x CW = 0.771
Smallest correlation: PH x CL = -0.038
Number of VIFs > 10: 0
Number of correlations with r >= |0.8|: 0
Variables with largest weight in the last eigenvalues:
```

```
KW > TKW > NR > NKR > CL > CW > PH > CD

----------------------------------------------------------------------------
Level: A4
----------------------------------------------------------------------------
Weak multicollinearity in the matrix
NC = 88.695
Matrix determinant: 0.0022499
Largest correlation: CW x KW = 0.815
Smallest correlation: PH x NKR = 0.032
Number of VIFs > 10: 1
Number of correlations with r >= |0.8|: 1
Variables with largest weight in the last eigenvalues:
KW > NKR > TKW > NR > CW > CD > PH > CL
```

Although the problems related to multicollinearity, some measures can be taken to mitigate their undesirable effects when detected by the above methods. It is now known that excluding variables responsible for inflating the variance of a regression coefficient is one of the most appropriate methods for reducing multicollinearity in explanatory variable matrices (T. Olivoto, Souza, et al., 2017). Identifying these variables, however, can be a difficult task. Recently, T. Olivoto, Nardino, et al. (2017) proposed the use of stepwise procedures along with sequential path analysis to identify a set of variables with high explanatory power but not highly correlated. When the exclusion of variables causing multicollinearity is not a procedure considered by the researcher –for example, due to a small number of explanatory variables or the importance of knowing their effects– a third option is to perform a path analysis with all explanatory variables, but including a small value in the diagonal elements of **X'X**. This procedure is known as *ridge regression* (Hoerl & Kennard, 1976). This procedure, however, overestimates the direct effects, especially of those variables with high VIF (T. Olivoto, Souza, et al., 2017).

##### 9.7.3 Estimation of path coefficients

The function `path_coeff()` is used to compute the path coefficients. Here, we will ignore the warning regarding the multicollinearity observed above and estimate the path coefficients considering the variable CW as the dependent trait and all others as predictor traits.

```
path_all <- path_coeff(data_ge2, KW)
```

```
The factors ENV GEN REP where excluded to perform the analysis. If you want to perform an analy

Severe multicollinearity.
Condition Number = 7865.84
Please, consider using a correction factor, or use 'brutstep = TRUE'.
```

According to NC, VIF, and D, the multicollinearity in the explanatory variable matrix is severe. For example, ten VIFs> 10 were observed and the matrix determinant was $1.089 \times 10^{-11}$. Analysis of the eigenvalues-eigenvectors (`path_all$weightvar`) indicated that, in order of importance, the variables that most contribute to multicollinearity are: CL > ED > CDED > EH > CW > PH

> NKE > EP > TKW > PERK > NR > EL > NKR > CD. As discussed, we have basically two options for circumventing the problems in our data. Exclude the variables responsible for multicollinearity, or keep all variables and include a correction factor on the diagonal **X'X**. Let's start with the last option.

For teaching purposes, we will choose, for the moment, an arbitrary value of $k$ equal to 0.05, included in the analysis with the argument `correction`

```
path_all_k <- path_coeff(data_ge2, KW, correction = 0.01)
```

```
The factors ENV GEN REP where excluded to perform the analysis. If you want to perform an analy

Moderate multicollinearity!
Condition Number = 532.621
Please, cautiosely evaluate the VIF and matrix determinant.
```

By including the correction factor ($k = 0.01$) the multicollinearity was classified as moderate (NC = 532.621). Inevitably, we have two options for lower levels of multicollinearity. The first is to increase the value of $k$, say, to 0.05. This would further reduce the multicollinearity level in our matrix, however, the bias in estimating the coefficients would increase. The second (and most reasonable) option is the exclusion of the variables that cause the most multicollinearity problems. For example, we can consider the variables with the highest weight in the last eigenvalues, or those with the highest VIF. Variables CL and EL are highly correlated, so, we could keep only one of these variables. The same interpretation can be considered for CDED. This is a covariate (ratio between cob diameter and ear diameter, CDED = CD / ED). Let's consider then excluding these variables.

The adjustment of the new model excluding these variables is easily accomplished. For this we will use two arguments of the `path_coeff()` function not seen so far: `pred` and `exclude`. The variables entered in `pred` can be either the predictor variables (*default*) or the variables to be excluded if `exclude = TRUE`. Let's go to the example.

```
path_exclude <- data_ge2 %>%
  path_coeff(resp = KW,
             pred = c(PERK, EH, CDED),
             exclude = TRUE)
```

```
Moderate multicollinearity!
Condition Number = 125.742
Please, cautiosely evaluate the VIF and matrix determinant.
```

The levels of multicollinearity in excluding variables still worry. We have seen that both identifying the variables responsible for multicollinearity and adjusting the model by declaring specific predictors is a relatively simple procedure using the `path_coeff()`. But what if some statistical-computational procedure made this task even easier? Let us now consider this.

T. Olivoto, Nardino, et al. (2017) suggested the use of stepwise regressions to select a set of predictors with minimal multicollinearity in path analysis. This option is available in the function `path_coeff()`. Based on an iterative algorithm called with the argument `brutstep = TRUE`, a set of predictors with minimal multicollinearity is selected based on the values of VIF. Subsequently, a series of stepwise regressions are adjusted. The first stepwise regression is adjusted by considering $p - 1$ selected predictor variables, with $p$ being the number of variables selected in the iterative process. The second model adjusts a regression considering $p - 2$ selected variables, and so on to the last model, which considers only two selected variables. Let's go to the example.

```
path_step <- path_coeff(data_ge2, KW, brutstep = TRUE)
```

```
--------------------------------------------------------------------------
The algorithm has selected a set of 10 predictors with largest VIF = 7.16.
Selected predictors: PERK EP CDED NKR PH NR TKW EL CD ED
A forward stepwise-based selection procedure will fit 8 models.
--------------------------------------------------------------------------
Adjusting the model 1 with 9 predictors (12.5% concluded)
Adjusting the model 2 with 8 predictors (25% concluded)
Adjusting the model 3 with 7 predictors (37.5% concluded)
Adjusting the model 4 with 6 predictors (50% concluded)
Adjusting the model 5 with 5 predictors (62.5% concluded)
Adjusting the model 6 with 4 predictors (75% concluded)
Adjusting the model 7 with 3 predictors (87.5% concluded)
Adjusting the model 8 with 2 predictors (100% concluded)
Done!
--------------------------------------------------------------------------
Summary of the adjusted models
--------------------------------------------------------------------------
  Model  AIC Numpred    CN Determinant    R2 Residual maxVIF
 Model8 1232       2  1.57     0.95068 0.860   0.1402   1.05
 Model7 1148       3  1.34     0.97871 0.919   0.0808   1.02
 Model6 1129       4 21.07     0.17146 0.930   0.0705   5.71
 Model5 1116       5 26.39     0.08049 0.936   0.0642   5.71
 Model4 1103       6 35.70     0.03481 0.942   0.0582   5.71
 Model3 1097       7 37.46     0.02618 0.944   0.0555   6.42
 Model2 1098       8 45.48     0.00396 0.945   0.0550   6.80
 Model1 1099       9 50.99     0.00216 0.945   0.0545   6.96
--------------------------------------------------------------------------
```

Note that the algorithm has selected a set of 10 predictors (PERK, EP, CDED, NKR, PH, NR, TKW, EL, CD, and ED) that has multicollinearity at acceptable levels. Thus, any of these models could be used without major problems in this regard. The stepwise procedure performed with different numbers of selected variables also allows the selection of a more parsimonious model, a task that will be at the discretion of the researcher. Here, for teaching porpuses, we will show the path coefficients using as predictors the traits NR, NKR, and TKW (Model 7).

```
path_final <- path_coeff(data_ge2,
                         resp = KW,
                         pred = c(NR, NKR, TKW))
```

```
Weak multicollinearity.
Condition Number = 1.339
You will probably have path coefficients close to being unbiased.
```

```
print(path_final)
```

```
------------------------------------------------------------------------------------------
Correlation matrix between the predictor traits
------------------------------------------------------------------------------------------
# A tibble: 3 x 3
        NR     NKR      TKW
*    <dbl>   <dbl>    <dbl>
1  1       0.02055 -0.1088
2  0.02055 1        0.09286
3 -0.1088  0.09286  1
------------------------------------------------------------------------------------------
Vector of correlations between dependent and each predictor
------------------------------------------------------------------------------------------
          NR       NKR       TKW
KW 0.3621447 0.5973701 0.6730371
------------------------------------------------------------------------------------------
Multicollinearity diagnosis and goodness-of-fit
------------------------------------------------------------------------------------------
Condition number:  1.3389
Determinant:       0.97871
R-square:          0.9192
Residual:          0.0808
Response:          KW
Predictors:        NR NKR TKW
------------------------------------------------------------------------------------------
Variance inflation factors
------------------------------------------------------------------------------------------
# A tibble: 3 x 2
  VAR     VIF
  <chr> <dbl>
1 NR    1.013
2 NKR   1.010
3 TKW   1.021
------------------------------------------------------------------------------------------
Eigenvalues and eigenvectors
------------------------------------------------------------------------------------------
# A tibble: 3 x 4
  Eigenvalues     NR     NKR      TKW
        <dbl>  <dbl>   <dbl>    <dbl>
1      1.133  0.5310 -0.4278 -0.7315
2      1.020  0.6466  0.7625  0.02346
3      0.8464 0.5477 -0.4854  0.6815
```

```
--------------------------------------------------------------------------------
Variables with the largest weight in the eigenvalue of smallest magnitude
--------------------------------------------------------------------------------
TKW > NR > NKR
--------------------------------------------------------------------------------
Direct (diagonal) and indirect (off-diagonal) effects
--------------------------------------------------------------------------------
# A tibble: 3 x 3
       NR      NKR      TKW
*   <dbl>    <dbl>    <dbl>
1  0.4242  0.008718 -0.04614
2  0.01082 0.5264    0.04888
3 -0.07290 0.06224   0.6703
--------------------------------------------------------------------------------
```

#### 9.8 Canonical Correlations

Canonical correlations can be computed using the function `can_corr()`. The first argument of the function is the (optional) data set that must contain the numeric variables used for the estimation of canonical correlations. Variable groups are defined by the arguments `FG` (first / smallest group) and`SG` (second / largest group). By default, a multicollinearity diagnosis is performed on each variable group. In the example below, the coefficients were stored in the object **ccc**.

```
ccc = can_corr(data_ge2,
               FG = c(PH, EH, EP),
               SG = c(EL, ED, CL, CD, CW, KW, NR),
               verbose = FALSE)
print(ccc)
```

```
---------------------------------------------------------------------------
Matrix (correlation/covariance) between variables of first group (FG)
---------------------------------------------------------------------------
      PH    EH    EP
PH 1.000 0.932 0.638
EH 0.932 1.000 0.870
EP 0.638 0.870 1.000

---------------------------------------------------------------------------
Collinearity diagnostic between first group
---------------------------------------------------------------------------
The multicollinearity in the matrix should be investigated.
NC = 977.586
Largest VIF = 229.164618380199
Matrix determinant: 0.0025852
Largest correlation: PH x EH = 0.932
Smallest correlation: PH x EP = 0.638
Number of VIFs > 10: 3
Number of correlations with r >= |0.8|: 2
```

```
Variables with largest weight in the last eigenvalues:
EH > PH > EP

---------------------------------------------------------------------------
Matrix (correlation/covariance) between variables of second group (SG)
---------------------------------------------------------------------------
        EL    ED    CL      CD    CW    KW      NR
EL  1.0000 0.385 0.255  0.9119 0.458 0.669 -0.0139
ED  0.3851 1.000 0.697  0.3897 0.737 0.824  0.5525
CL  0.2554 0.697 1.000  0.3004 0.738 0.471  0.2619
CD  0.9119 0.390 0.300  1.0000 0.484 0.626 -0.0358
CW  0.4582 0.737 0.738  0.4840 1.000 0.735  0.1657
KW  0.6686 0.824 0.471  0.6260 0.735 1.000  0.3621
NR -0.0139 0.553 0.262 -0.0358 0.166 0.362  1.0000

---------------------------------------------------------------------------
Collinearity diagnostic between second group
---------------------------------------------------------------------------
Weak multicollinearity in the matrix
NC = 68.376
Matrix determinant: 0.0015322
Largest correlation: EL x CD = 0.912
Smallest correlation: EL x NR = -0.014
Number of VIFs > 10: 0
Number of correlations with r >= |0.8|: 2
Variables with largest weight in the last eigenvalues:
KW > ED > EL > CD > CL > CW > NR

---------------------------------------------------------------------------
Matrix (correlation/covariance) between FG and SG)
---------------------------------------------------------------------------
      EL    ED    CL    CD    CW    KW    NR
PH 0.380 0.661 0.325 0.315 0.505 0.753 0.329
EH 0.363 0.630 0.397 0.281 0.519 0.703 0.265
EP 0.263 0.458 0.391 0.175 0.425 0.497 0.140

---------------------------------------------------------------------------
Correlation of the canonical pairs and hypothesis testing
---------------------------------------------------------------------------
        Var Percent   Sum  Corr Lambda Chisq DF   p_val
U1V1 0.6315   76.19  76.2 0.795  0.296 181.8 21 0.00000
U2V2 0.1867   22.53  98.7 0.432  0.805  32.5 12 0.00116
U3V3 0.0106    1.28 100.0 0.103  0.989   1.6  5 0.90148

---------------------------------------------------------------------------
Canonical coefficients of the first group
---------------------------------------------------------------------------
      U1    U2     U3
```

```
PH  2.53  5.87   7.32
EH -2.44 -8.26 -12.45
EP  1.14  2.75   6.49

---------------------------------------------------------------------------
Canonical coefficients of the second group
---------------------------------------------------------------------------
         V1     V2     V3
EL -0.00893 -0.936  0.767
ED  0.19372  0.297 -1.824
CL -0.08385 -1.215  0.172
CD -0.30662  1.137 -1.423
CW -0.15226  0.191  0.478
KW  1.16752 -0.126  1.125
NR -0.05866  0.486  0.622
```

#### 9.9 Cluster Analysis

Cluster analysis is a very useful multivariate procedure in plant breeding. The basic principle is to group individuals (genotypes) according to their similarities (analyzed traits). Distance matrices and the implementation of hierarchical clustering algorithms for constructing dendrograms can be made with the function `clustering()`.

Assuming that the researcher wishes to compute the distances between each genotype (which is logical in a MET analysis) with a typical MET data the argument `means_by = GEN` argument. In this case, the average of each genotype is internally computed for each numeric variable and the distance is computed using these averages. In addition, the `plot()` function can be used to plot a dendrogram. A line is drawn at the suggested cutoff point according to Mojena (1977). Let's go to an example.

```
d2 = clustering(data_ge2,
                NKR, TKW, NKE,
                means_by = GEN)
plot(d2, horiz = FALSE, ylab = "Euclidean distance")
```

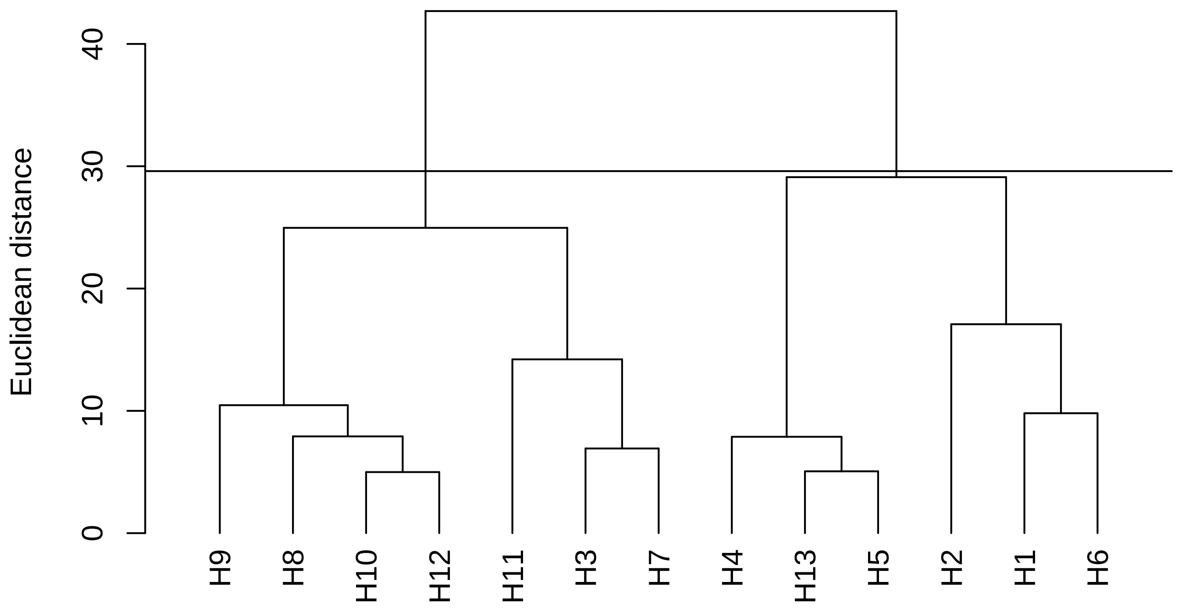

One way to measure how well the generated dendrogram reflects your data is to calculate the correlation between the cophenetic distances and the original distance matrix. The cophenetic correlation coefficient was already computed and can be assessed by typing

```
d2$cophenetic
```

```
[1] 0.8640355
```

The function `clustering()` counts with an algorithm for variable selection. The aim is to select a group of traits that most contribute to explain the variability of the original data. Let's say that if a few traits could be used to cluster genotypes without loss of information, human and financial resources could be spared. So instead of evaluating 15 traits (as in `data_ge2`), we could evaluate only those that really contribute to the genotype distinction.

The variable selection algorithm is invoked when the argument `selvar = TRUE` is included in the function. Variable selection is based on the eigenvalue/eigenvector solution. Assuming a dataset with $p$ traits, we need:

**1:** compute the distance matrix and the cophenetic correlation with the original traits (all numeric traits in the data set)

`for (i in 1:p - 2){`

**2:** compute the eigenvalues and eigenvectors of the correlation matrix between traits;

**3:** delete the variable with the highest weight (highest eigenvector at lowest eigenvalue);

**4:** compute the distance matrix and cophenetic correlation with the remaining traits;

**5:** compute the Mantel correlation between the obtained distance matrix and the original distance matrix;

}

At the end of the iterations, a summary of the models is returned. The distance is calculated with the traits that generated the model with the highest cophenetic correlation. We suggest a careful evaluation in order to choose a parsimonious model, that is, the one with the smallest number of traits, which presents acceptable cophenetic correlation and high similarity with the original distances.

```
sel_var =  clustering(data_ge2, means_by = GEN, selvar = TRUE)
```

```
Calculating model 1 with 15 variables. EH excluded in this step (7.1%).
Calculating model 2 with 14 variables. EP excluded in this step (14.3%).
Calculating model 3 with 13 variables. CDED excluded in this step (21.4%).
Calculating model 4 with 12 variables. PH excluded in this step (28.6%).
Calculating model 5 with 11 variables. CL excluded in this step (35.7%).
Calculating model 6 with 10 variables. NR excluded in this step (42.9%).
Calculating model 7 with 9 variables. PERK excluded in this step (50%).
Calculating model 8 with 8 variables. EL excluded in this step (57.1%).
Calculating model 9 with 7 variables. CD excluded in this step (64.3%).
Calculating model 10 with 6 variables. ED excluded in this step (71.4%).
Calculating model 11 with 5 variables. KW excluded in this step (78.6%).
Calculating model 12 with 4 variables. CW excluded in this step (85.7%).
Calculating model 13 with 3 variables. NKR excluded in this step (92.9%).
Calculating model 14 with 2 variables. TKW excluded in this step (100%).
Done!
-------------------------------------------------------------------------

Summary of the adjusted models
-------------------------------------------------------------------------
    Model excluded cophenetic remaining cormantel    pvmantel
  Model 1        -  0.8656190        15 1.0000000 0.000999001
  Model 2       EH  0.8656191        14 1.0000000 0.000999001
  Model 3       EP  0.8656191        13 1.0000000 0.000999001
  Model 4     CDED  0.8656191        12 1.0000000 0.000999001
  Model 5       PH  0.8656189        11 1.0000000 0.000999001
  Model 6       CL  0.8655939        10 0.9999996 0.000999001
  Model 7       NR  0.8656719         9 0.9999982 0.000999001
  Model 8     PERK  0.8657259         8 0.9999977 0.000999001
```

```
  Model 9       EL  0.8657904        7 0.9999972 0.000999001
 Model 10       CD  0.8658997        6 0.9999964 0.000999001
 Model 11       ED  0.8658274        5 0.9999931 0.000999001
 Model 12       KW  0.8643556        4 0.9929266 0.000999001
 Model 13       CW  0.8640355        3 0.9927593 0.000999001
 Model 14      NKR  0.8648384        2 0.9925396 0.000999001
--------------------------------------------------------------------------
Suggested variables to be used in the analysis
--------------------------------------------------------------------------
The clustering was calculated with the  Model 10
The variables included in this model were...
 ED CW KW NKR TKW NKE
--------------------------------------------------------------------------
```

One of the problems with hierarchical clustering is that it does not tell us how many clusters there are or where to cut out the dendrogram to form the clusters. You can cut the hierarchical tree at a certain height, say, on average distances, however, this decision is purely unhelpful. For example, if the cutoff is too high, we tend to group genotypes that may, in fact, not be similar. A very low cutoff, on the other hand, may result in selection failures, as we consider that genotypes are distinct where, in fact, they are not. By default, the function gives the cutoff calculated by the method of Mojena (1977). Bootstrap resampling-based procedures that allow estimating p-values for each junction can also be used (Suzuki & Shimodaira, 2006).

```r
library(pvclust)
pv_clust <- pvclust(t(sel_var$data), nboot = 100, method.dist = "euclidean")
```

```
Bootstrap (r = 0.5)... Done.
Bootstrap (r = 0.67)... Done.
Bootstrap (r = 0.83)... Done.
Bootstrap (r = 1.0)... Done.
Bootstrap (r = 1.17)... Done.
Bootstrap (r = 1.33)... Done.
```

```r
plot(pv_clust, hang = -1, cex = 0.5)
pvrect(pv_clust, alpha = 0.95)
```

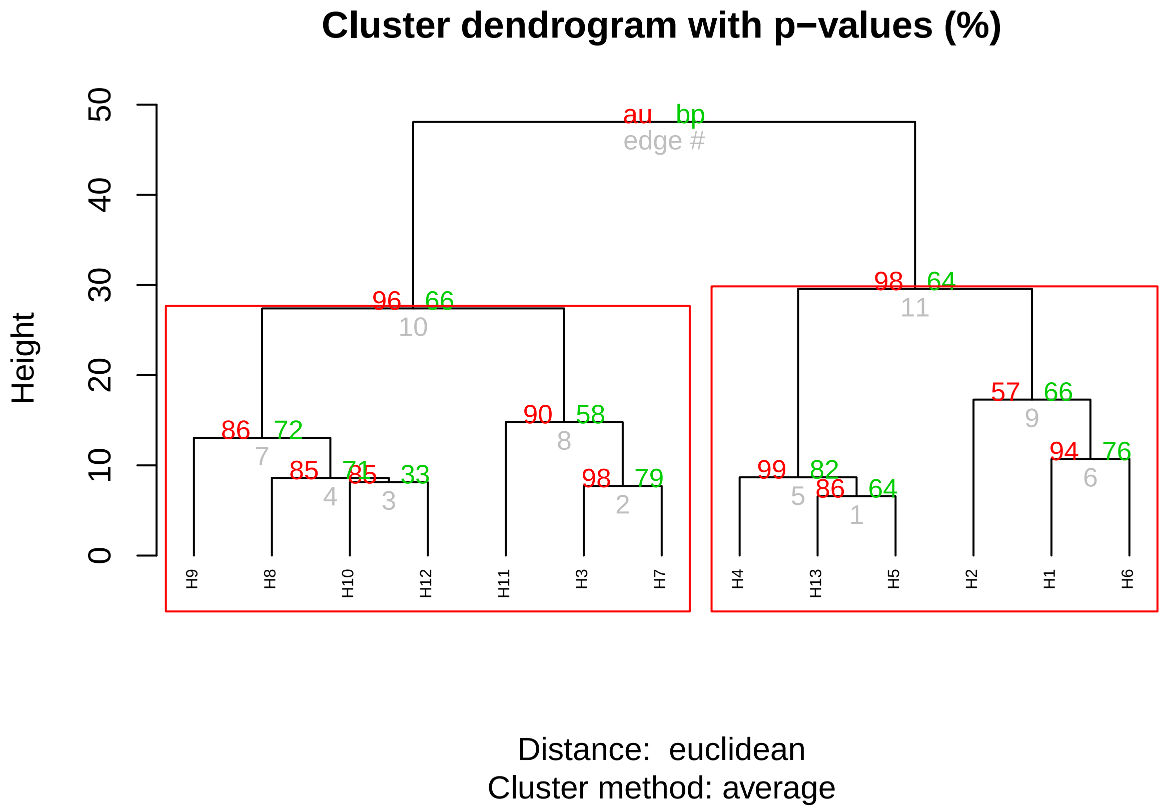

The residual (co)variance matrix obtained in the function function `covcor_design()` can be used as arguments in the function `mahala()` to compute the Mahalanobis distance. Note that in this example we use the argument `type = "means"` to get the means of the traits for each genotype

```
data = subset(data_ge2, ENV == "A1")
means = data %>%
      covcor_design(GEN, REP,
                    resp = c(PH, EH, NKE, TKW),
                    type = "means")

D2 <- mahala(.means = means, covar = rcov)
print(D2, digits = 2)
```

| | H1 | H10 | H11 | H12 | H13 | H2 | H3 | H4 | H5 | H6 | H7 | H8 | H9 |
|---|---|---|---|---|---|---|---|---|---|---|---|---|---|
| H1 | 0.0 | 2.18 | 3.83 | 5.91 | 3.1 | 19.0 | 10.0 | 3.6 | 4.73 | 4.7 | 4.97 | 5.9 | 6.3 |
| H10 | 2.2 | 0.00 | 0.62 | 2.59 | 4.0 | 10.5 | 4.6 | 2.8 | 0.66 | 6.5 | 1.32 | 2.0 | 3.9 |
| H11 | 3.8 | 0.62 | 0.00 | 0.73 | 5.4 | 8.1 | 4.1 | 3.4 | 0.69 | 9.8 | 0.25 | 1.2 | 7.2 |
| H12 | 5.9 | 2.59 | 0.73 | 0.00 | 7.2 | 7.8 | 5.3 | 4.7 | 2.37 | 13.5 | 0.54 | 1.9 | 12.4 |
| H13 | 3.1 | 4.02 | 5.40 | 7.23 | 0.0 | 11.9 | 6.0 | 2.3 | 4.68 | 2.0 | 6.30 | 5.1 | 6.7 |
| H2 | 19.0 | 10.46 | 8.07 | 7.77 | 11.9 | 0.0 | 5.0 | 11.6 | 6.75 | 17.6 | 7.87 | 5.0 | 16.5 |
| H3 | 10.0 | 4.61 | 4.07 | 5.26 | 6.0 | 5.0 | 0.0 | 2.6 | 2.04 | 11.8 | 3.28 | 5.2 | 8.4 |
| H4 | 3.6 | 2.82 | 3.38 | 4.74 | 2.3 | 11.6 | 2.6 | 0.0 | 2.53 | 7.1 | 3.04 | 5.8 | 7.1 |

```
H5   4.7  0.66 0.69  2.37  4.7  6.8  2.0  2.5 0.00  8.3  0.89 1.8  4.6
H6   4.7  6.54 9.75 13.54  2.0 17.6 11.8  7.1 8.32  0.0 11.93 8.2  4.8
H7   5.0  1.32 0.25  0.54  6.3  7.9  3.3  3.0 0.89 11.9  0.00 2.1  8.8
H8   5.9  2.00 1.24  1.90  5.1  5.0  5.2  5.8 1.78  8.2  2.10 0.0  7.9
H9   6.3  3.90 7.22 12.42  6.7 16.5  8.4  7.1 4.57  4.8  8.84 7.9  0.0
```

A dendrogram can then be created by coercing the Mahalanobis's distance matrix `D2` to an object of class `hclust`, as follows

```
d <- D2 %>% dist() %>% hclust()
plot(d, hang = -1)
```

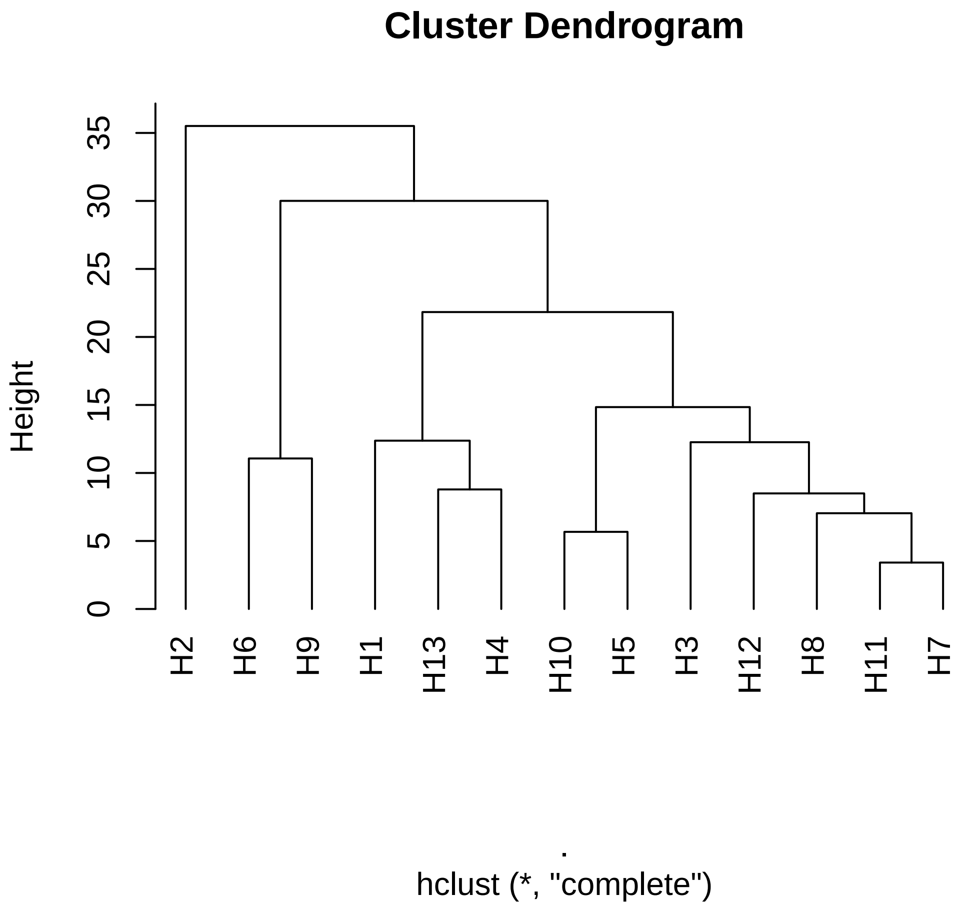

#### 10 Data visualization

`metan` provides useful functions for quickly creating typical plots used in the analysis of two-way data from MET analysis. This section is focused on creating plots using treatments of type (i) qualitative *vs* qualitative; (ii) qualitative *vs* quantitative; and (iii) quantitative *vs* quantitative

#### 10.1 Qualitative *vs* qualitative

Using the function `plot_factbars()` we will create a bar plot to show the EH of genotypes in the environments. For simplicity, we will subset the data example `data_ge2` to show only three hybrids (H1-H3) and two environments.

In function `plot_factbars()` the required arguments are only the data, the two factors, and the response variable. An error bar showing the standard error is shown by default. It is also possible to choose other statistics to show in the error bar such as standard deviation and confidence interval. To control specific aspects of the plot theme, use the function `ggplot2::theme()`

```
data_plot <- subset(data_ge2,
                    GEN %in% c("H1", "H2", "H3") &
                    ENV  %in% c("A1", "A2"))

a <- plot_factbars(data_plot, GEN, ENV, resp = EH)
b <-  plot_factbars(data_plot,
                     GEN,
                     ENV,
                     resp = EH,
                     xlab = "Genotypes",
                     ylab = "Ear height (m)",
                     col = FALSE,
                     # Letters only for teaching purposes
                     lab.bar = c("aA", "aA", "bB", "aA", "aA", "aA"),
                     lab.bar.vjust = -2,
                     size.text.bar = 3,
                     y.expand = 1.5,
                     width.bar = 0.6,
                     width.erbar = 0.2,
                     fontfam = "serif")
library(ggplot2)
c <-  a +
       theme(panel.grid.major = element_line(color = "gray90"),
             panel.background = element_rect(fill = "gray"))+
             ggtitle("My custom plot theme")

arrange_ggplot(a, b, c, ncol = 3, labels = letters[1:3])
```

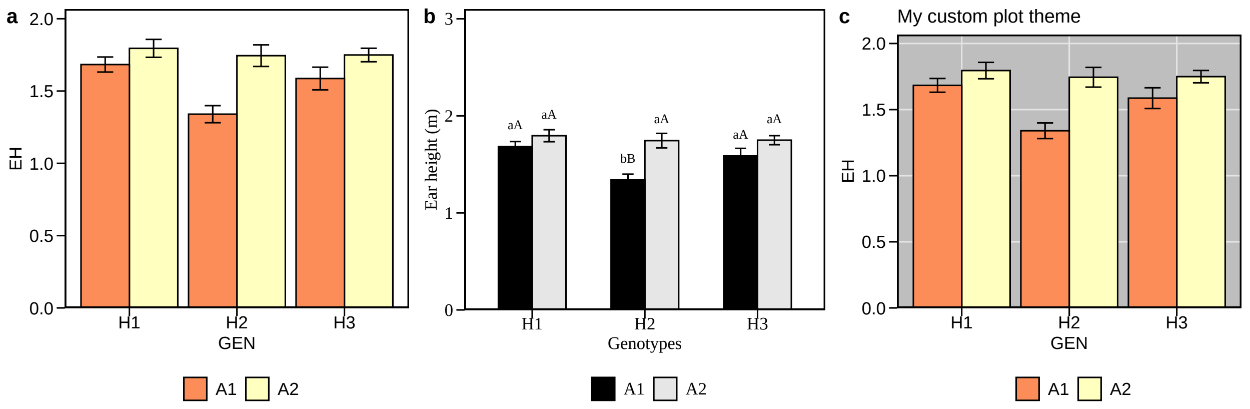

#### 10.2 Qualitative *vs* quantitative

Line plots containing one qualitative (e.g., genotype) and one quantitative factor (e.g., nitrogen rate) can be easily created with the function `plot_factlines()`. The data used to produce the plots in this and the next section have been achieved at https://doi.org/10.5281/zenodo.3526636 in folder `data` as `data_R.xlsx`. In this section, the data in sheet FAT1_CI is used. It has one qualitative factor (HIBRIDO) one quantitative factor (DOSEN) one column for blocks and one column for grain yield (RG).

In function `plot_factlines()` the required arguments are `.data` (the data set), `x` (the variable in the x-axis), `y` (the variable in the y-axis), `group` (a grouping variable), and `fit` to indicate the polynomial degree for regression lines. If a regression for each level of the grouping variable needs to be fitted, then, `fit` must be a numeric vector with the same length of the levels in `group`. If `fit` has length 1, then one regression is fitted for all levels.

In the next example, we will fit four regression lines, (linear for NUPEC_1 and NUPEC_4, and quadratic for NUPEC_2 and NUPEC_3)

```
library(readxl)
FAT1_CI <- read_excel("D:/Desktop/data_R.xlsx", sheet = "FAT1_CI")
str(FAT1_CI)
```

```
Classes 'tbl_df', 'tbl' and 'data.frame':   80 obs. of  4 variables:
 $ HIBRIDO: chr  "NUPEC_1" "NUPEC_1" "NUPEC_1" "NUPEC_1" ...
 $ DOSEN  : num  0 0 0 0 25 25 25 25 50 50 ...
 $ BLOCO  : num  1 2 3 4 1 2 3 4 1 2 ...
 $ RG     : num  8.2 8.5 8.8 8.76 8.94 ...
```

```
a <- plot_factlines(FAT1_CI,
                     DOSEN,
                     RG,
                     HIBRIDO,
                     fit = c(1, 2, 2, 1))
b <- plot_factlines(FAT1_CI,
                     DOSEN,
```

```
                          RG,
                          HIBRIDO,
                          fit = c(1, 2, 2, 1),
                          grid = TRUE)
c <- plot_factlines(FAT1_CI,
                          DOSEN,
                          RG,
                          HIBRIDO,
                          fit = 1,
                          col = FALSE)
arrange_ggplot(a, b, c, nrow = 1, labels = letters[1:3])
```

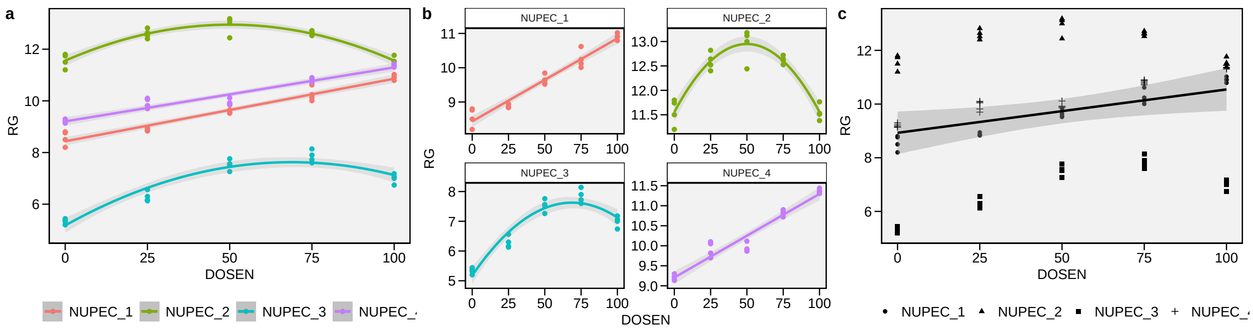

> **Quick tip**
>
> - Use the function `plot_lines()` to make a graph similar to the previous one.

#### 10.3 Quantitative *vs* quantitative

In this section the data in sheet FAT3 of `data_R` is used. It has two quantitative factors (DOSEN and DOSEK) one column for blocks and one column for grain yield (RG). The function `resp_surf()` can be used to fit the following surface response model

$$Y_i = \beta_0 + \beta_1 A_i + \beta_2 D_i + \beta_3 A_i^2 + \beta_4 D_i^2 + \beta_5 A_i D_i + \epsilon_i$$

where $A$ and $D$ are the quantitative factors.

The stationary point, i.e., the combination of $A$ and $D$ that provides the highest value of response variable is estimated as

$$-0.5 \times (\boldsymbol{A}^{-1}\boldsymbol{X})$$

where

$$\boldsymbol{A} = \begin{pmatrix} \beta_3 & \beta_5/2 \\ \beta_5/2 & \beta_4 \end{pmatrix}$$

and

$$\boldsymbol{X} = \begin{pmatrix} \beta_1 \\ \beta_2 \end{pmatrix}$$

```
FAT3 <- read_excel("D:/Desktop/data_R.xlsx", sheet = "FAT3")
str(FAT3)
```

```
Classes 'tbl_df', 'tbl' and 'data.frame':   80 obs. of  4 variables:
 $ DOSEN: num  45 45 45 45 45 45 45 45 45 45 ...
 $ DOSEK: num  0 0 0 0 25 25 25 25 50 50 ...
 $ BLOCO: num  1 2 3 4 1 2 3 4 1 2 ...
 $ RG   : num  120 125 129 130 140 147 144 142 150 153 ...
```

```
rs_model <- resp_surf(FAT3, DOSEN, DOSEK, BLOCO, RG)
```

```
-----------------------------------------------------------------
Result for the analysis of variance
Model: Y = m + bk + Ai + Dj + (AD)ij + eijk
-----------------------------------------------------------------
            Df Sum Sq Mean Sq  F value   Pr(>F)
BLOCO        3    158      53    3.621   0.0183 *
DOSEN        3  65978   21993 1515.063  < 2e-16 ***
DOSEK        4  11817    2954  203.513  < 2e-16 ***
DOSEN:DOSEK 12   2363     197   13.563 1.21e-12 ***
Residuals   57    827      15
---
Signif. codes:  0 '***' 0.001 '**' 0.01 '*' 0.05 '.' 0.1 ' ' 1
-----------------------------------------------------------------
Shapiro-Wilk's test for normality of residuals:
-----------------------------------------------------------------
W =  0.946194 p-valor =  0.002100403
-----------------------------------------------------------------
Anova table for the response surface model
-----------------------------------------------------------------
Analysis of Variance Table

Response: RG
            Df Sum Sq Mean Sq  F value    Pr(>F)
DOSEN        1   3215    3215  15.4854  0.000186 ***
DOSEK        1   6538    6538  31.4899 3.301e-07 ***
I(DOSEN^2)   1  50925   50925 245.2693 < 2.2e-16 ***
I(DOSEK^2)   1   5098    5098  24.5541 4.440e-06 ***
DOSEN:DOSEK  1      1       1   0.0053  0.942298
Residuals   74  15365     208
---
```

```
Signif. codes:  0 '***' 0.001 '**' 0.01 '*' 0.05 '.' 0.1 ' ' 1
-----------------------------------------------------------------
Model equation for response surface model
Y = B0 + B1*A + B2*D + B3*A^2 + B4*D^2 + B5*A*D
-----------------------------------------------------------------
Estimated parameters
B0: -317.8340786
B1: 14.7502633
B2: 1.0056943
B3: -0.1121344
B4: -0.0076331
B5: 0.0001973
-----------------------------------------------------------------
Matrix of parameters (A)
-----------------------------------------------------------------
-0.1121344    9.87e-05
9.87e-05    -0.0076331
-----------------------------------------------------------------
Inverse of the matrix A (invA)
-8.9179679    -0.1152744
-0.1152744    -131.0091259
-----------------------------------------------------------------
Vetor of parameters B1 e B2 (X)
-----------------------------------------------------------------
B1: 14.7502633
B2: 1.0056943
-----------------------------------------------------------------
Equation for the optimal points (A and D)
-----------------------------------------------------------------
-0.5*(invA*X)
Eigenvalue 1: -0.007633
Eigenvalue 2: -0.112135
Stacionary point is maximum!
-----------------------------------------------------------------
Stacionary point obtained with the following original units:
-----------------------------------------------------------------
Optimal dose (DOSEN): 65.8292
Optimal dose (DOSEK): 66.7277
-----------------------------------------------------------------
Fitted model
-----------------------------------------------------------------
A = DOSEN
D = DOSEK
y = -317.83408+14.75026A+1.00569D+-0.11213A^2+-0.00763D^2+2e-04A*D
-----------------------------------------------------------------
Shapiro-Wilk normality test
p-value:  0.4522241
According to Shapiro-Wilk normality test at 5% of significance, residuals can be considered no
```

```
----------------------------------------------------------------------
```

In our example

$$\boldsymbol{A} = \begin{pmatrix} -0.11213 & 9.865e-05 \\ 9.865e-05 & -0.00763 \end{pmatrix}; \boldsymbol{A}^{-1} = \begin{pmatrix} -8.91796 & -0.1152 \\ -0.11527 & -131.009 \end{pmatrix}$$

and

$$\boldsymbol{X} = \begin{pmatrix} 14.7502 \\ 1.00569 \end{pmatrix}$$

Thus

$$-0.5 \times \left[\begin{pmatrix} -8.91796 & -0.1152 \\ -0.11527 & -131.009 \end{pmatrix} \times \begin{pmatrix} 14.7502 \\ 1.00569 \end{pmatrix}\right] = \begin{pmatrix} 65.8292 \\ 66.7277 \end{pmatrix}$$

A contour plot can then be obtained by typing

```r
a <- plot(rs_model)
b <- plot(rs_model,
          region = FALSE,
          xlab = "Nitrogen rate",
          ylab = "Potassium rate")
arrange_ggplot(a, b, labels = letters[1:2])
```

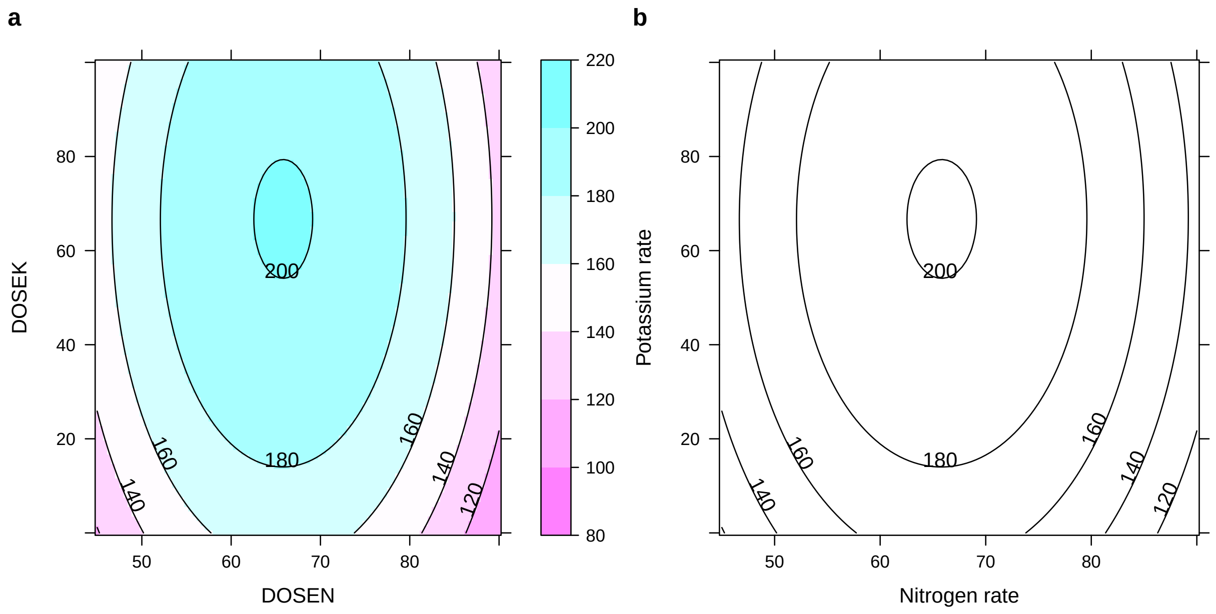
